# A 3D molecular atlas of the chick embryonic heart

**DOI:** 10.1101/609032

**Authors:** Claire Anderson, Bill Hill, Hui-Chun Lu, Adam Moverley, Youwen Yang, Nidia M.M. Oliveira, Richard A. Baldock, Claudio D. Stern

**Affiliations:** Department of Cell and Developmental Biology, University College London, London, WC1E 6BT, UK; MRC Human Genetics Unit, IGMM, University of Edinburgh, EH4 2XU, UK

**Keywords:** RNAseq, transcriptome, molecular anatomy, cardiac, cardiovascular, chick embryo

## Abstract

We present a detailed analysis of gene expression in the 2-day (HH12) embryonic chick heart. RNA-seq of 13 micro-dissected regions reveals regionalised expression of 15,570 genes. Of these, 132 were studied by *in situ* hybridisation and a subset (38 genes) was mapped by Optical Projection Tomography or serial sectioning to build a detailed 3-dimensional atlas of expression. We display this with a novel interactive 3-D viewer and as stacks of sections, revealing the boundaries of expression domains and regions of overlap. Analysis of the expression domains also defines some sub-regions distinct from those normally recognised by anatomical criteria at this stage of development, such as a previously undescribed subdivision of the atria into two orthogonal sets of domains (dorsoventral and left-right). We also include a detailed comparison of expression in the chick with the mouse and other species.

## Introduction

At stage 12 of chick embryo development (HH12; Hamburger and Hamilton, 1951) (2-days’ incubation), the territories that will give rise to most major regions of the mature heart can already be defined (apart from the definitive outflow tract and right ventricle, which undergoes extensive subsequent development from secondary/anterior heart field progenitors (de la Cruz et al., 1977; Mjaatvedt et al., 2001; Waldo et al., 2001)). Growth from localized proliferation zones (Soufan et al., 2006; van den Berg et al., 2009), as described by the ballooning model (Christoffels et al., 2000; Moorman and Christoffels, 2003), accounts for subsequent expansion and elaboration of the mature heart. Stage HH12 is equivalent to mouse E8.5-8.75 (Theiler stage 13-14) (de Boer et al., 2012), and human Carnegie stage 10-11 (20-26 days) (Sizarov et al., 2011). Specific molecular markers are known for several of these regions, but not for others, such as the early left pacemaker. Given current interest in identifying cells in organoids and cultures of embryonic stem cells and iPS cells (Sahara et al., 2015), as well as in single-cell RNA-seq studies, there is an urgent need for a comprehensive molecular map of the embryonic heart at stages when progenitors can be defined.

The chick embryo is an excellent model system because it is an amniote (like mouse and human) and specific heart regions can be accurately dissected. To identify regional markers, we micro-dissected 13 territories of the HH12 chick heart and analysed them by RNA-seq. These territories were defined by morphology, fate mapping, physiology and known gene expression patterns (Figure 1; see Supplementary Methods) as corresponding to: rostral (RDM) and caudal (CDM) dorsal mesocardia, outflow tract (OFT), prospective right (RV) and left (LV) ventricles, ventricular endocardium (En), future atria (At), the presumed prospective right pacemaker (RPC) and the corresponding tissue on the left side (“left-RPC”), the left pacemaker (LPC), right (RPE) and left (LPE) proepicardia and the sinus venosus (SV). The transcriptomes of these 13 regions were compared to identify unique regional markers for each. The spatial patterns of expression of 132 of these markers were then studied using *in situ* hybridisation (ISH), Optical Projection Tomography (OPT) and histological sections. The clearest regionally restricted markers (including many not previously described) are used to generate a 3D map; the patterns of expression also define some new sub-regions. We also relate our findings to the mouse embryonic heart, confirming conservation of markers.

**Figure 1.**
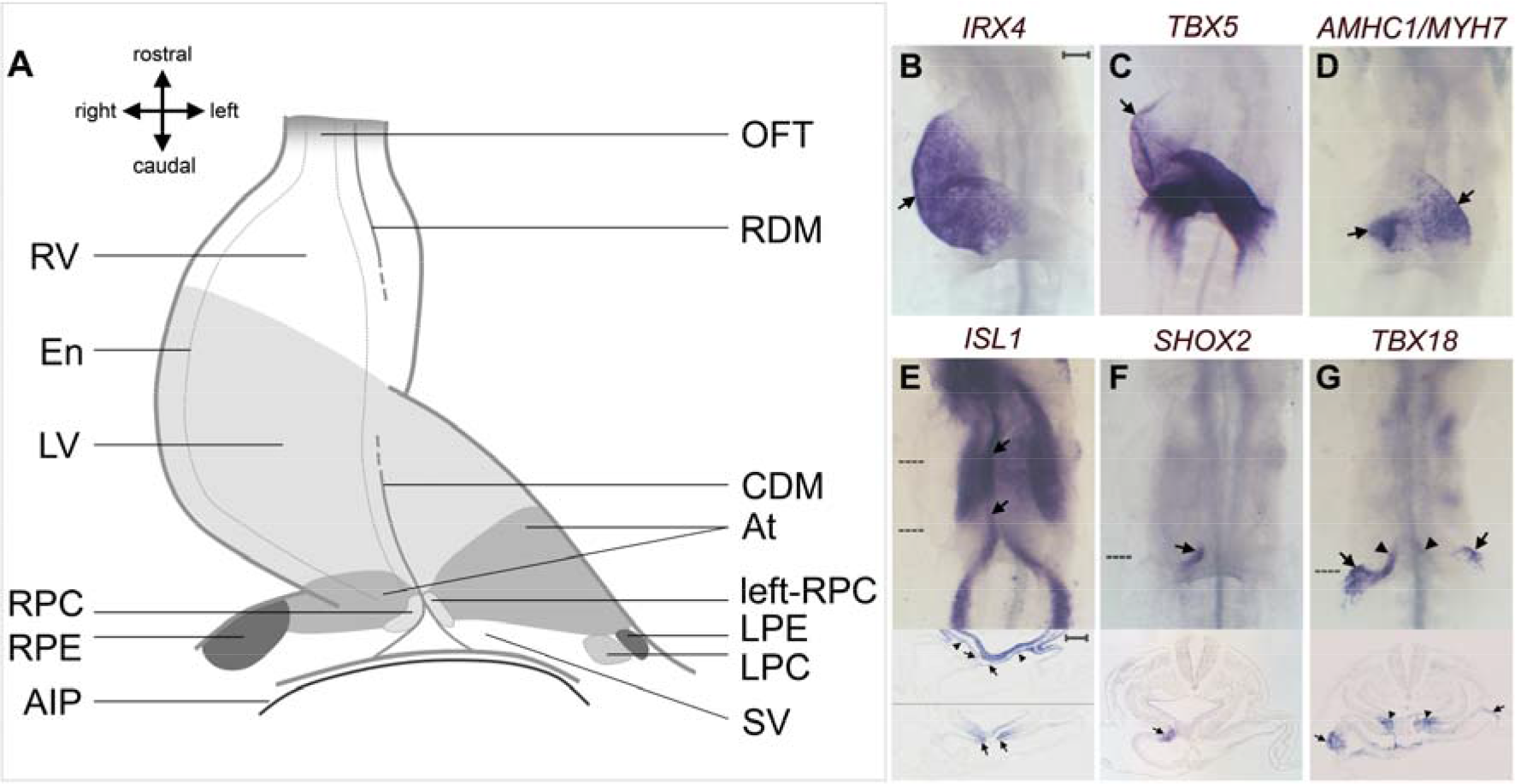
Gene expression criteria used to define different regions of the HH12 chick heart (see Supplementary Methods). **A.** Cartoon of heart regions dissected for RNA-seq based on gene expression patterns. Atrium, At; caudal dorsal mesocardium, CDM; endocardium, En; left pacemaker, LPC; prospective left ventricle, LV; left proepicardium, LPE; outflow tract, OFT; rostral dorsal mesocardium, RDM; right proepicardium, RPE; presumed prospective right pacemaker, RPC, and the corresponding tissue on the left side, “left-RPC”; prospective right ventricle, RV; sinus venosus, SV. **B-G**. Examples of markers. **B**. Prospective ventricle (*IRX4*). **C**. The rostral boundary of *TBX5* expression marking the border between prospective RV and LV (arrow). **D**. Atrium (*AMHC1*/*MYL7*). **E**. *ISL1* in both the RDM and CDM (arrows) and the rostral splanchnic mesoderm (SM; arrowheads). **F**. RPC (*SHOX2*). **G**. *TBX18* in proepicardia (arrows) and caudal-most DM (arrowheads). Scale bars: 100μm.

## Materials and Methods

For full methods, see “Supplementary Methods”. Briefly, fertile hens’ eggs were incubated at 38°C to HH12 (Hamburger and Hamilton, 1951). Thirteen regions of the heart were dissected (Figure 1A) and RNA extracted using RNAqueous Micro-kit (Ambion). Criteria for defining the regions to be dissected are described in detail in the Supplementary Methods. Libraries were constructed using NEBNext RNA Ultra II with Poly A+ selection (NEB) and sequenced. Sequence reads were aligned to the chick genome (galGal5) (Supplementary Dataset 1; the full dataset has been deposited in ArrayExpress with the name ‘A 3D molecular atlas of the chick embryonic heart’, accession E-MTAB-7663). Whole mount ISH was performed as described (Streit and Stern, 2001). Stained embryos were either processed for OPT (Sharpe et al., 2002) or paraffin-embedded and sectioned. Selected OPT assays were mapped onto a reference model of the HH12 chick heart through constrained distance transforms (Hill and Baldock, 2015) established using the Woolz image processing system (https://github.com/ma-tech/).

## Results

Supplementary Dataset 1 compares the expression (in Fragments Per Kilobase of transcript per Million mapped reads, FPKM) of 15,570 genes in 13 micro-dissected heart regions. In most cases, FPKM ≥20 and a fold change (FC) ≥1.5 were used to select differentially expressed genes uniquely (ie. with minimal overlap with other regions) expressed in one of the 13 regions (see Supplementary Methods). From these, we selected 132 (Supplementary Figure 1) putative regional markers for study by ISH. Note that as a result of the selection criteria applied (unique expression in one region), genes with left-right differential expression as well as some known markers were not selected for ISH because they are expressed in more than one region. However the datasets provided (Supplementary Dataset 1) can be interrogated in a number of different ways, for example to reveal genes expressed in more than one region.

### Outflow tract

Eleven candidate markers for the OFT were studied by ISH. Useful, novel markers for this region include *PRSS23* (Figure 2A) and *CR385930/LBH-LIKE* (Figure 3A). *CR385930/LBH-LIKE* is expressed only in the non-myocardial part of the OFT. Four other genes are localised to the OFT: *FGF8*, *MSX2*, *GDF3/cVg1* and *HPGDS* (Supplementary Figure 2A). Both *FGF8* and *MSX2* are involved in formation of the mouse OFT (Brown et al., 2004; Chen et al., 2007). The remaining five genes (*ACTA2*, *RASSF5*, *OLFML2B*, *SPRY2* and *PKIG*) are not OFT-specific (Supplementary Figure 3A).

**Figure 2.**
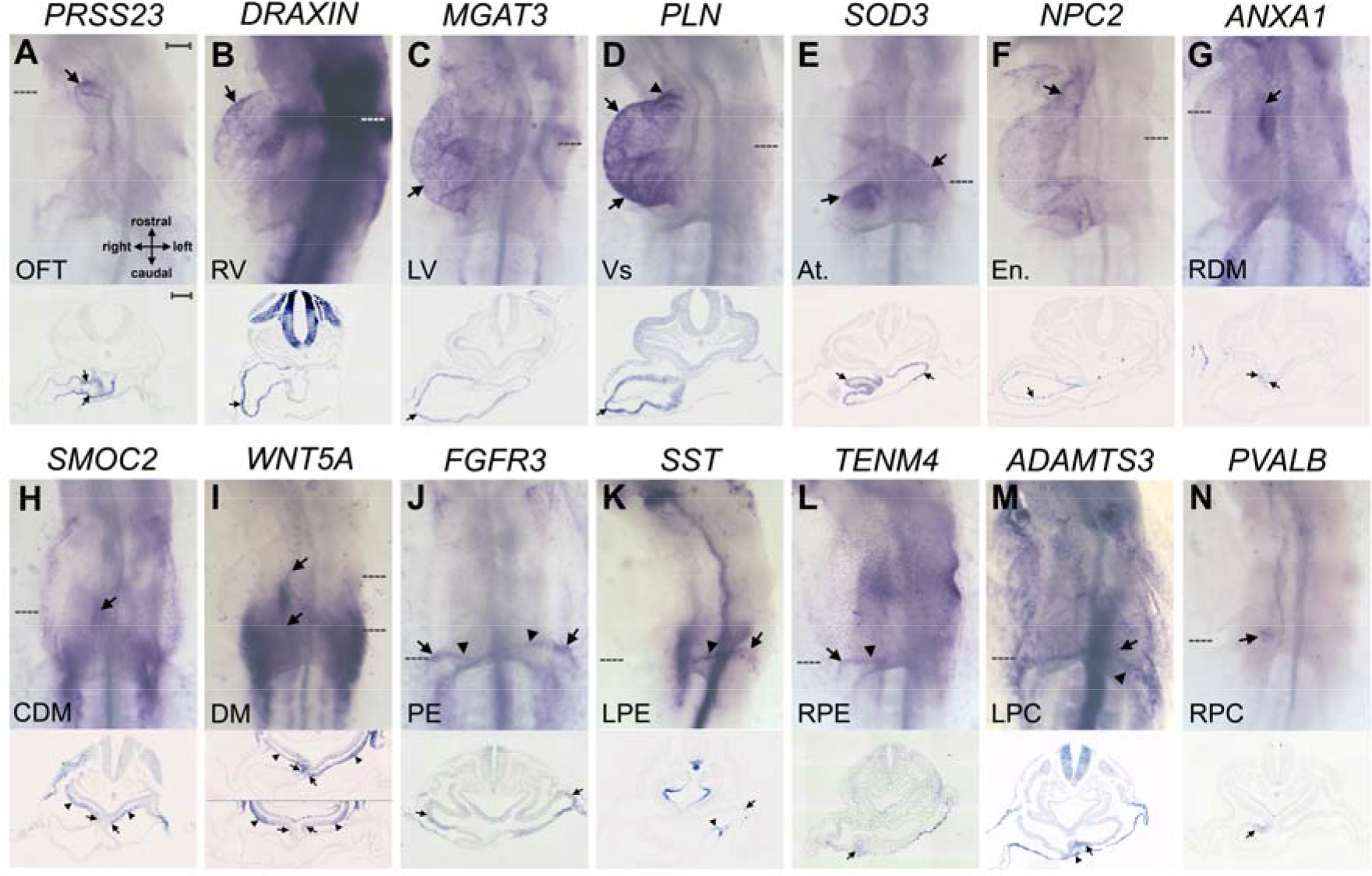
A selection of markers for 14 territories of the chick heart. **A**. *PRSS23* in the OFT (arrows). **B**. *DRAXIN* in RV (arrows). **C**. *MGAT3* in prospective LV (arrows). **D**. *PLN* in RV, LV (arrows) and OFT (arrowhead). **E**. *SOD3* in atria (arrows). **F**. *NPC2* in endocardium especially within the OFT (arrow in whole mount). **G**. *ANXA1* in RDM (arrows). **H**. *SMOC2* in CDM (arrows) and SM (arrowheads). **I**. *WNT5A* in RDM, CDM (arrows) and SM (arrowheads). **J**. *FGFR3* in RPE and LPE (arrows). **K**. *SST* in LPE (arrows). **L**. *TENM4* in RPE (arrows) (J-L are detected in proepicardial serosa; arrowheads). **M**. *ADAMTS3* in putative early LPC (arrows) and endoderm (arrowheads). **N**. *PVALB* in RPC (arrows). Scale bars: 100μm.

**Figure 3.**
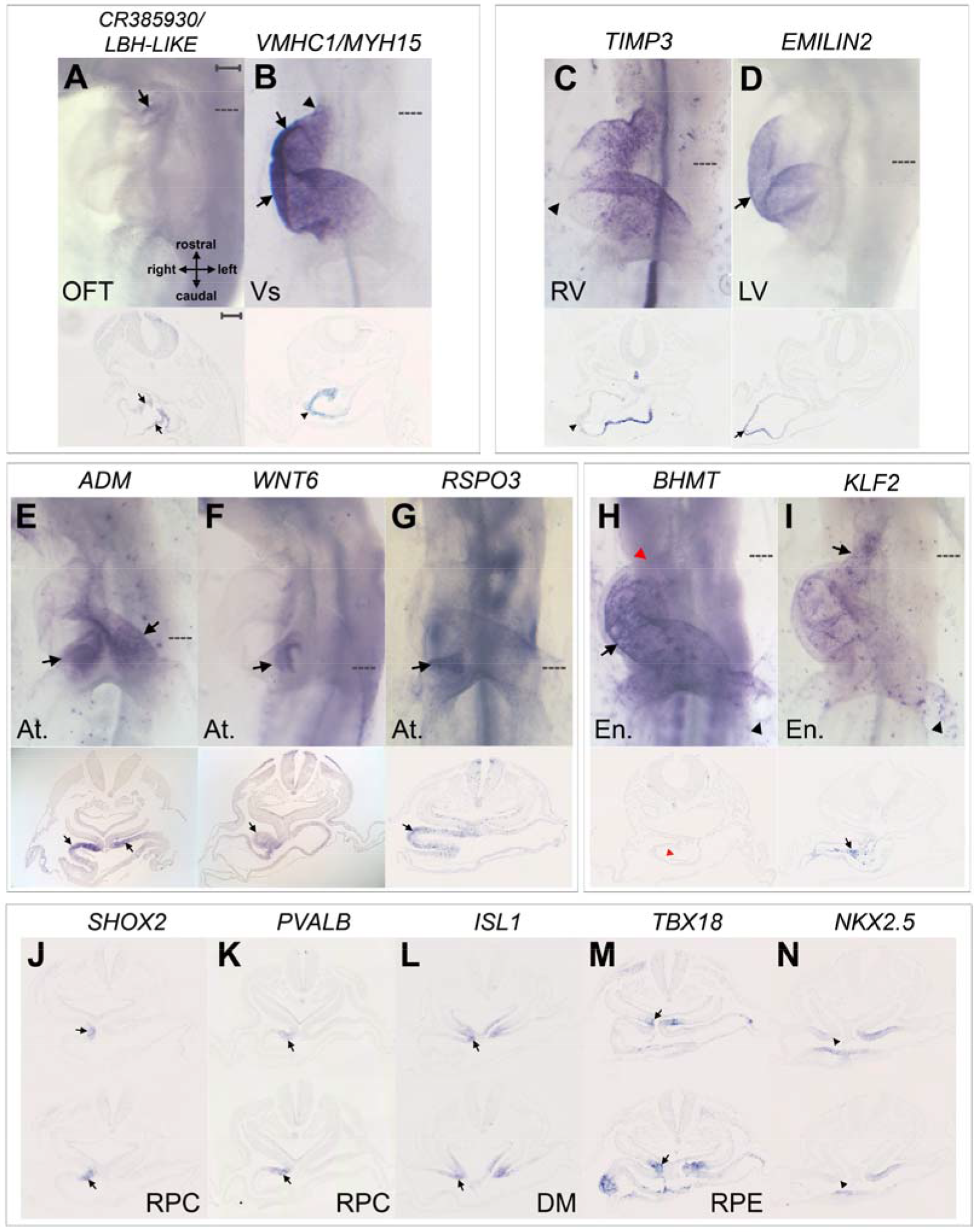
Sub-regions within heart territories. **A-B**. *CR385930/LBH-LIKE* in the non-myocardial OFT (arrows, A), and *VMHC1/MYH15* in the OFT myocardium (arrowheads) and ventricles (arrows, B). **C-D**. *TIMP3* in the myocardium except in LV (arrowhead, C), whereas *EMILIN2* is in LV (arrow, D). **E**. *ADM* in dorsal atria (arrows). **F-G**. *WNT6* and *RSPO3* on the right of the atrial region (arrows). **H-I**. *BHMT* in the endocardium (arrow, H) but not in the rostral pole of the heart (red arrowheads), whereas *KLF2* expression in endocardium is strongest in the OFT (arrows, I); both are also detected in endothelial cells (black arrowheads). **J-N**. RPC markers *SHOX2* (J) and *PVALB* (K) overlap *ISL1* (arrows, L), partially overlap *TBX18* (arrows, M) and do not overlap *NKX2.5* (arrowheads, N) expression. Scale bars: 100μm.

### Right ventricle

Differences in gene expression between the prospective RV and LV have not been described at this stage of heart development; our RNA-seq analysis reveals several differentially expressed genes. Seven RV-enriched genes were studied further. The best markers are *DRAXIN* (Figure 2B), *BMP10* and *TGFBI* (Supplementary Figure 2B), expressed particularly in the outer curvature. *NPPA*/*NPPB* expression is localised to the inner and outer curvatures of the prospective ventricles (Supplementary Figure 2B). *FBXL22*, *AMY2A* (Supplementary Figure 3B) and *TIMP3* (Figure 3C) are not sufficiently restricted to be useful RV markers. However, *TIMP3* is expressed throughout the myocardium excluding the LV (Figure 3C; cf. *EMILIN2*, see below). Several of these genes have been implicated in heart development, adult physiology, regeneration or repair, including *BMP10* (Chen et al., 2004), *TGFBI* (Schwanekamp et al., 2017), *FBXL22* (Spaich et al., 2012) and *TIMP3* (Fedak et al., 2004).

### Left ventricle

Seven markers for the prospective LV were analysed by ISH. *MGAT3* (Figure 2C), *MB* (Supplementary Figure 2C) and *EMILIN2* (Figure 3D) localise to the prospective LV. *PACAP*/*ADCYAP1* extends into the OFT, RV and atrial region, while *HS6ST3*, *PLBD1* and *ZNF106* are either too faint or strong to be useful LV markers (Supplementary Figure 3C).

### Both ventricles

In addition to the above, RNA-seq analyses for the RV and LV were combined to find markers for the prospective ventricle, uncovering known markers *IRX4*(Bao et al., 1999) (Figure 1B) and *VMHC1/MYH15* (Bisaha and Bader, 1991) (Figure 3B). Fourteen other genes were analysed: *PLN* extends rostrally into the myocardial part of the OFT (Figure 2D) like *VMHC1/MYH15* (Figure 3B); *LDB3* extends into the atrium, while *MYL3* and *SSPN* extend to both the OFT and atrium (Supplementary Figure 2D). The remaining 11 markers are too faint (*LOX*, *PLPPR1*, *VCAM1*) or too strongly expressed (*CMYA5, EXFABP, MYL1, SORBS2, TNNI2, KCNH6, LYPD1*) to be useful ventricular markers (Supplementary Figure 3D). Therefore only *IRX4* is restricted to the prospective ventricles.

### Atria

Known atrial marker *AMHC1*/*MYL7* (Yutzey et al., 1994) (Figure 1D) was revealed in the RNA-seq comparisons; 11 other genes were investigated by ISH. Our analysis also uncovers several other good atrial (At) markers, including *SOD3* (Figure 2E). Some of these markers are localised to different subdomains within the atrium: *ADM* (Figure 3E), *TUFT1, VIM* and *STBD1* (Supplementary Figure 2E) transcripts are enriched dorsally and *WNT6* and *RSPO3* are enriched on the right (Figure 3F,G). Four genes (*GJD2*, *FAM3B*, *UGT1A1, BORCS8*) had faint or strong expression throughout the heart (Supplementary Figure 3E).

### Endocardium

Many differentially expressed genes were revealed in the Endocardium (En) comparison; increasing the FC to ≥4.0 reveals 51 candidates, 14 of which were analysed further. *NPC2* is a good endocardial marker (Figure 2F), while *RARRES1*, *EDN1*, *NFATC1*, *STAB1*, *TIE1*, *PLVAP* (Supplementary Figure 2F), *BHMT and KLF2* (Figure 3H, I) are detected in endothelial cells as well as endocardium. *BHMT* is not expressed in the endocardium of the OFT or RV, whereas *NPC2* and *KLF2* expression is strongest in these regions (Figure 2F, Figure 3H, I), suggesting that the endocardium is regionally-subdivided. *FOXO1*, *HEG1*, *POSTN*, *TAL1* and *IQSEC1* were not detected in the HH12 endocardium (Supplementary Figure 3F). Several of these genes have known roles in endocardium and/or endothelium.

### Rostral mesocardium

Despite both being derived from splanchnic mesoderm (SM), the rostral (RDM) and caudal (CDM) parts of the dorsal mesocardium are molecularly distinct, perhaps reflecting their different roles: RDM generates secondary heart field progenitors for the OFT and RV (Waldo et al., 2001), while CDM contributes to SV and atria (van den Berg et al., 2009). Thirteen RDM-enriched genes were analysed, the best of which are *ANXA1* (Figure 2G) and *TNC* (Supplementary Figure 2G). Most others are continuous with the SM (*PKDCCB*, *LUM*, *BMP4*, *LOC417741*) and myocardium (*EBF3*, *KAZALD1*) or not detected (*PROM1*, *CCDC3*, *SIX1*, *SLIT2*) (Supplementary Figure 3G). *ISL1* transcripts are enriched in the RDM in the RNA-seq analysis, but are detected in the rostral SM and throughout the mesocardium by ISH (Figure 1E) (Yuan and Schoenwolf, 2000).

### Caudal mesocardium

Of 10 CDM markers assessed by ISH, most are weak and continuous with the SM, including *SMOC2* (Figure 2H), *PTCH2* (Supplementary Figure 2H), *FOXF2, APCDD1, STRA6* and *HOXA2*, whereas *MYCN*, *HOXA1*, *LINGO1* and *NGFR* are not significantly detected (Supplementary Figure 3H). Therefore none of these are useful CDM-specific markers. *PTCH2* expression is of interest, as Ptch1-mediated signalling within the DM is involved in atrioventricular septation in mouse (Goddeeris et al., 2008).

### Dorsal mesocardium

The RNA-seq analyses for RDM and CDM were combined to reveal markers for the DM; 7 genes were assessed by ISH. Most are continuous with the SM (*WNT5A*, Figure 2I; *IRX1*, Supplementary Figure 2I) and myocardium (*RDH10*, *SALL4)* or are not significantly detected in the DM (*LIX1*, *IRX3*, *ER81/ETV1*, Supplementary Figure 3I), therefore none of the seven genes analysed by ISH are useful DM markers, leaving only *ISL1* (Yuan and Schoenwolf, 2000) (see above).

### Proepicardium

The LPE and RPE RNA-seq analyses (below) were combined to reveal PE-enriched transcripts, including known marker *WT1*, previously observed only in the RPE (Schlueter et al., 2006) (Supplementary Figure 2J). Nine genes were analysed further. *FGFR3* (Figure 2J), *MAB21L1* and *MSX1* (Supplementary Figure 2J) are good markers. *GAL* and *FGFR2* are faint in the PE (Supplementary Figure 3J). Many of these are also detected in SV cells ventromedial to the PE (the “proepicardial serosa”). The remaining markers are haemoglobin-β-related (*HBZ*, *HBM* and *HBAA*); although not PE-specific (Supplementary Figure 3J), their enrichment in the RNA-seq suggests that the PE contains haematopoietic cells, as in mouse embryos (Jankowska-Steifer et al., 2018).

### Left proepicardium

No LPE-specific markers are known. Nine genes were analysed by ISH; *SST* was the best marker, localising to the LPE and left ventromedial cells in the SV (Figure 2K). *LHX2* is detected in both PE, but unusually, expression is stronger in the LPE than in the RPE (Supplementary Figure 2K). *ANXA2* transcripts are in both PE, whereas *RBP4A*, *APOC3*, *HOXB5*, *LMO2*, *PAMR1* and *HOXB4* are not detected (Supplementary Figure 3K). SST signalling may be involved in LPE regression in chick embryos (Schulte et al., 2007), as it is implicated in cell cycle arrest and/or apoptosis in cell lines (Lamberts et al., 2002).

### Right proepicardium

Of five RPE markers investigated by ISH, only *TENM4* is detected there (Figure 2L). Transcripts for *LSP1* (Supplementary Figure 2L) and known PE marker *TBX18* (Haenig and Kispert, 2004) (Figure 1G), are enriched in the RPE comparison but are also detected in LPE. *GEM*, *OPRM1* and *AGPAT2* are strongly expressed throughout the heart (Supplementary Figure 3L). Later in development, the epicardium undergoes *WT1*-mediated epithelial-mesenchymal transition (EMT) to form coronary vessels (Martinez-Estrada et al., 2010); *TENM4* may play a role in this as *Tenm4* mutant mouse embryos fail to undergo EMT (Nakamura et al., 2013).

### Left pacemaker

Despite being the early pacemaker in the heart (Bressan et al., 2013), only three genes were identified in the LPC-region (FC ≤1.5). Both *ADAMTS3* (Figure 2M) and *TTR* (Supplementary Figure 2M) are detected in the left side of the SV region in sections, overlapping ventromedial LPE cells (cf. *SST*, Figure 2K). Therefore, *ADAMTS3* and *TTR* could either be new markers for the LPC, for ventromedial LPE cells, or both. *GUCA2A* is not detected (Supplementary Figure 3M).

### Right pacemaker

RNA-seq shows enrichment of known sinoatrial node marker, *SHOX2*(Espinoza-Lewis et al., 2009) (Figure 1F). Two genes were studied by ISH. *PVALB* (Figure 2N) is detected in the dorso-caudal right atrium, overlapping *SHOX2. Pvalb*, implicated in calcium binding in rodent skeletal muscles, had not been detected in the heart (Celio and Heizmann, 1982), but is being explored as therapy for impaired ventricular relaxation (Szatkowski et al., 2001). *ISL1* and *TBX18*, but not *NKX2.5*, are detected in the *SHOX2*-*PVALB*-expressing region (Figure 3J-N), consistent with expression and lineage analyses of sinoatrial node progenitors in mouse embryos (Christoffels et al., 2010). *CENPW* expression is not localised to the RPC (Supplementary Figure 3N).

### “Left-RPC”

Although the left side corresponding to the RPC was taken to ensure enrichment of RPC transcripts, some genes were found to be “left-RPC”-specific by RNA-seq. *JAG1* and *RAI14* were studied further but are detected on both the left and right side (Supplementary Figure 3O).

### Pacemaker regions

The LPC and putative RPC were analysed to find common transcripts. Only one gene was revealed, *FGB*, which was not specifically expressed (Supplementary Figure 3P).

### Sinus venosus

RNA-seq analysis of isolated SV tissue only reveals two genes, *MSI2* and *TNR*, which have strong and widespread expression. Neither is SV-specific (Supplementary Figure 3Q).

### Resources

The results of RNA-seq (Supplementary Dataset 1; ArrayExpress accession number E-MTAB-7663), ISH in whole-mounts and sections (Figures 2-3, Supplementary Figures 2-3 and Supplementary Dataset 2) and 3-dimensional OPT scans of selected markers (Supplementary Datasets 3-4) reveal a number of good markers, both known and novel. Their known salient features are described in Supplementary Table 1, highlighting five markers (*BMP10*, *SOD3*, *NPC2*, *SHOX2* and *PVALB*) that are not expressed elsewhere outside of the heart, and not previously described as restricted within their region in chick or other species.

As resources for the community, we provide Supplementary Dataset 2 as a series of registered sections for 40 markers, and a selection of 28 markers mapped onto a HH12 chick heart as a novel, interactive 3D viewer (http://www.biomedatlas.org/ChickHeartAtlas/HH12/chick_heart_gene_expression.html; see Figure 4). This 3D viewer can also be accessed locally from Supplementary Dataset 5. Image data have been submitted to Image Data Resource, IDR (https://idr.openmicroscopy.org; accession number: idr0059).

**Figure 4.**
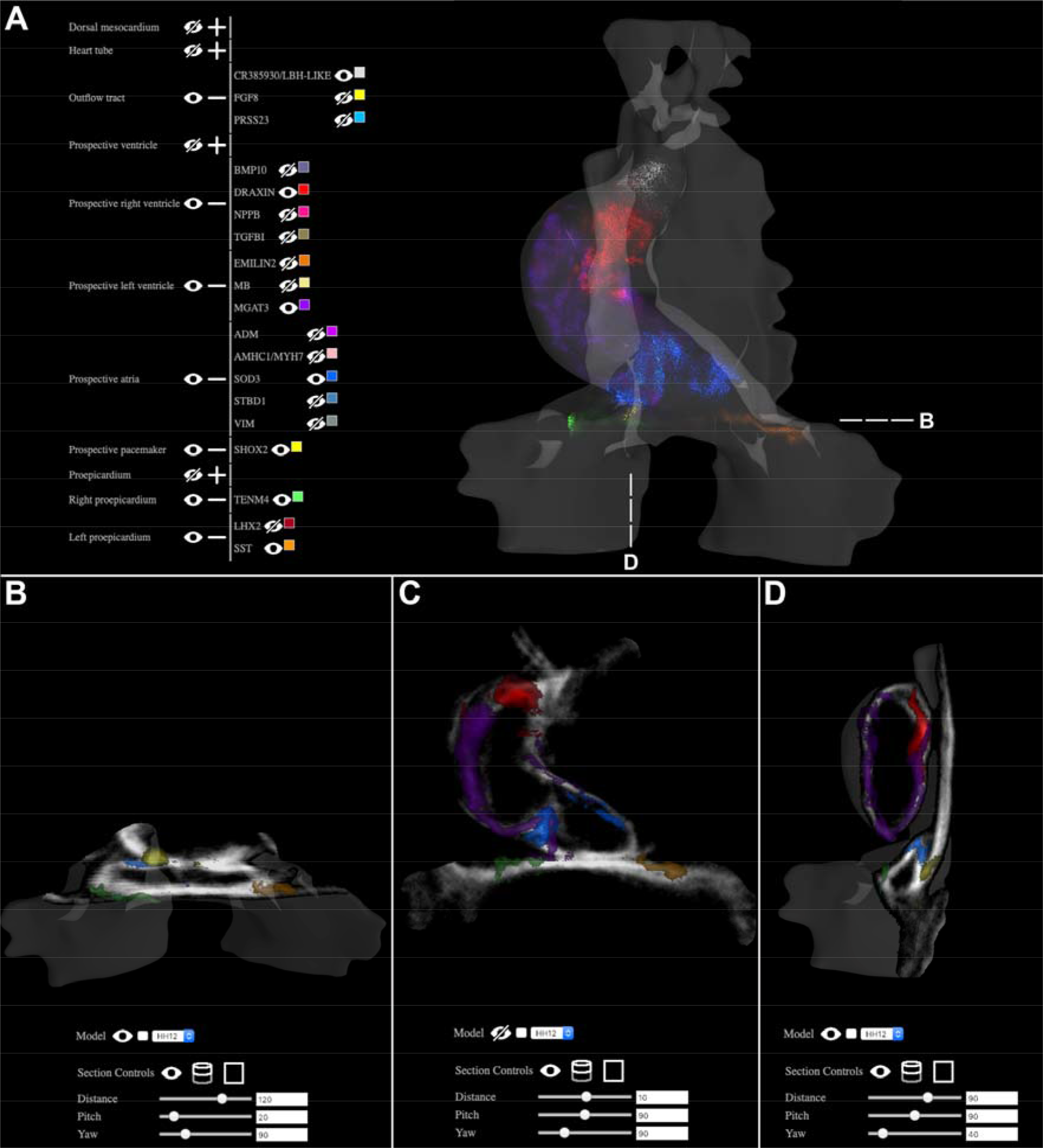
3D mapping of localised heart markers onto a HH12 chick heart model. **A**. Overview of the interactive 3D browser with selected heart markers visible as ‘points-clouds’, which allows multiple overlapping and graded patterns of expression to be observed (http://www.biomedatlas.org/ChickHeartAtlas/HH12/chick_heart_gene_expression.html). **B-D**. Transverse (B), coronal (C) and sagittal (D) section views.

## Discussion

Our study provides a comprehensive dataset of genome-wide mRNA expression revealing regionalised expression for 15,570 genes among 13 heart regions. Comparative analysis of their transcriptomes reveals hundreds of differentially expressed genes. Of these, 132 were studied further by ISH. We constructed a detailed 3-dimensional atlas of regional expression of 38 excellent markers in the HH12 chick embryonic heart, including an interactive tool to visualise expression and regions of overlap. Several markers had not been previously described. This analysis helps to define sub-regions distinct from those normally recognised by anatomical criteria at this stage of development, including a novel atrial subdivision into two orthogonal sets of domains (dorsoventral and left-right).

The 13 HH12 chick heart territories chosen for RNAseq were defined based on morphological criteria, fate maps and known physiological characteristics and previously described gene expression patterns.

Expression of some of these markers is largely conserved in mouse embryos at an equivalent stage of heart development (E8.5-8.75; de Boer et al., 2012) (Supplementary Table 1). Our analysis complements other RNA-seq analyses of mouse and human heart, combining dissection and single-cell RNA-seq to predict spatial patterns of gene expression (Li et al., 2016) or define a spatiotemporal transcriptome for heart cell types (e.g. cardiomyocytes, fibroblasts, endothelium and cardiac progenitor cells) (DeLaughter et al., 2016; Sahara et al., 2019). The closest analysis to the present study was performed in zebrafish, where RNA-seq was coupled with sections (“Tomo-seq”) (Burkhard and Bakkers, 2018) to reconstruct a 3D transcriptomic map. Our study, together with Tomo-seq of the zebrafish heart, relates the transcriptome of cells to their general position within the heart, revealing the state of cells at a particular time of development. A comprehensive molecular map of the embryonic heart will be helpful for future identification of heart progenitor cells generated in single-cell RNA-seq studies and in *in vitro* organoids, embryonic stem cells and iPS cells. However it is worth emphasizing that most developmentally-regulated genes are expressed in multiple cell types and/or at multiple stages in development, therefore gene expression patterns should not be considered as reliable indicators of cell *fates*, but rather of cell *states* (Stern and Canning, 1990; Izpisúa-Belmonte et al., 1993; Joubin and Stern, 1999; Stern, 2009) Therefore single-cell RNAseq is a useful tool to predict cell lineage relationships, but the interpretation of the results is highly dependent on the value of the markers selected and on the depth of prior knowledge that influenced both the selection of markers and the experimental design (selection of the cell populations to analyse and time points). Examining the relationships between multiple gene expression patterns in *space*, as in the present study, especially looking at regions of overlap (co-expression) and mosaicism (salt-and-pepper patterns), can help to define the degree to which different combinations of genes might be useful as regional or cell type specific markers.

## Acknowledgements

We are grateful to BHF (PG/17/2/32737) and BBSRC (BB/R003432/1) for funding and to K. Trevers, T. Solovieva, H.C. Lee and I. Almeida for comments and advice.

