## Supplementary material for "A 3D molecular atlas of the chick embryonic heart": Dataset legends

**Supplementary Table 1** List of genes analysed by *in situ* hybridisation with references to published expression patterns. Genes highlighted in red are specific to the region indicated by the RNAseq analysis, they are not expressed elsewhere within the embryo and have not been previously described as restricted to that region in chick or in other species.

**Supplementary Dataset 1** RNAseq readings and comparisons between thirteen HH12 chick heart regions; FPKM (Fragments Per Kilobase of transcript per Million mapped reads); red text indicates a fold change <1.5; bold text indicates genes tested by mRNA *in situ* hybridisation.

**Supplementary Dataset 2** Histological sections of 40 markers as registered stacks of images.

**Supplementary Dataset 3** Reconstructed Optical Projection Tomography scans of 28 markers. The *in situ* hybridisation signal is red and the anatomy signal is blue.

**Supplementary Dataset 4** Optical Projection Tomography scans of 28 markers that have been mapped onto a HH12 chick heart reference model. Mapped assays were used to construct the 3D-interactive browser. NIfTI files are grey-scale *in situ* hybridisation signal only.

**Supplementary Dataset 5** Local 3D interactive browser. The README file contains instructions on how to locally load the 3D interactive browser.
