## Supplementary Figures for "A 3D molecular atlas of the chick embryonic heart"

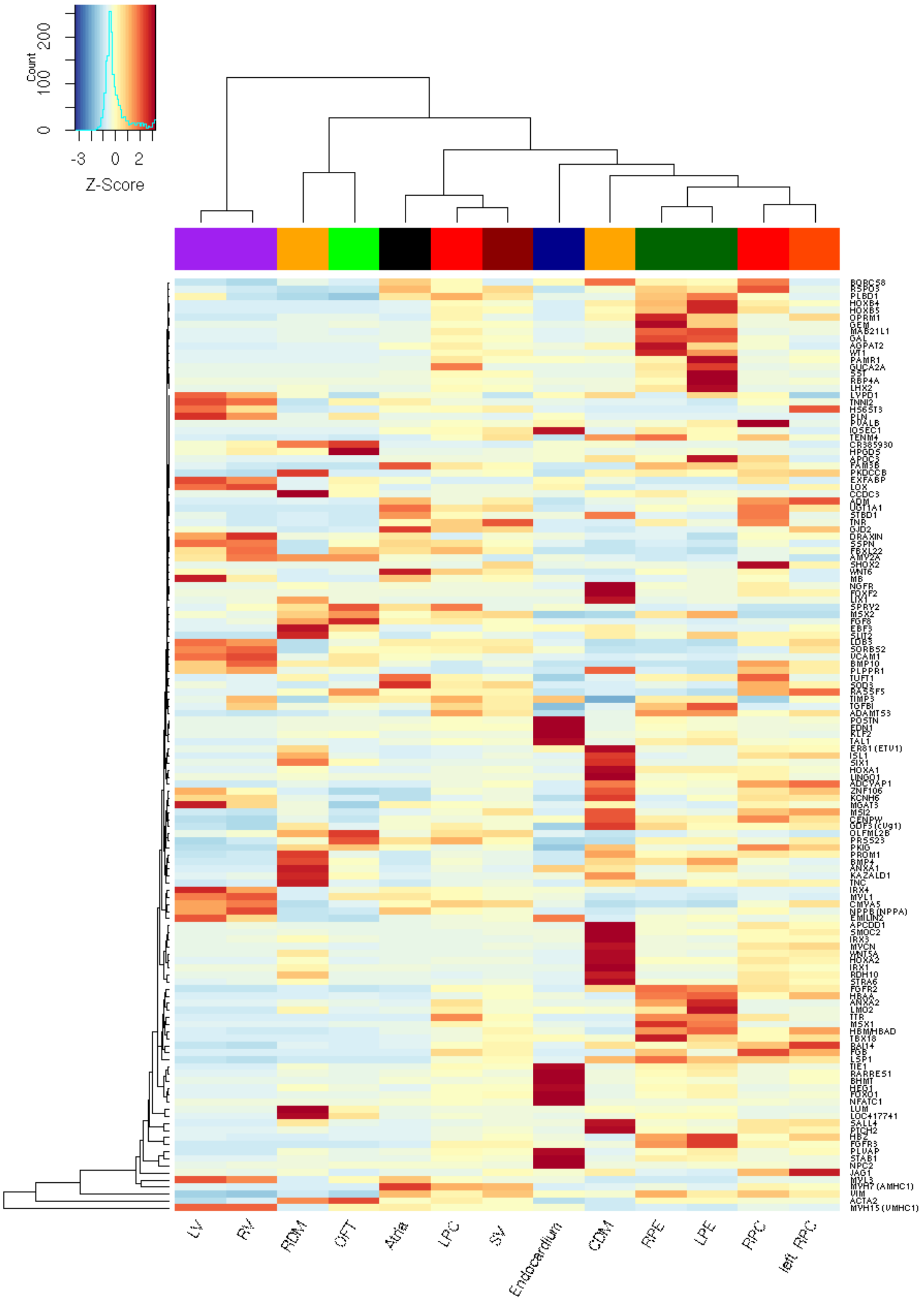


**Supplementary figure 1.** Hierarchical clustering of RNAseq readings for 132 genes tested by mRNA *in situ* hybridisation. Enriched RNAs are displayed in red, depleted RNA in blue. LV, prospective left ventricle; RV, prospective right ventricle; RDM, rostral dorsal mesocardium; OFT, outflow tract; SV, sinus venosus; LPC, left pacemaker; CDM, caudal dorsal mesocardium; LPE, left proepicardium; RPE, right proepicardium; left-RPC, left side corresponding to the position of the RPC; RPC, right pacemaker.

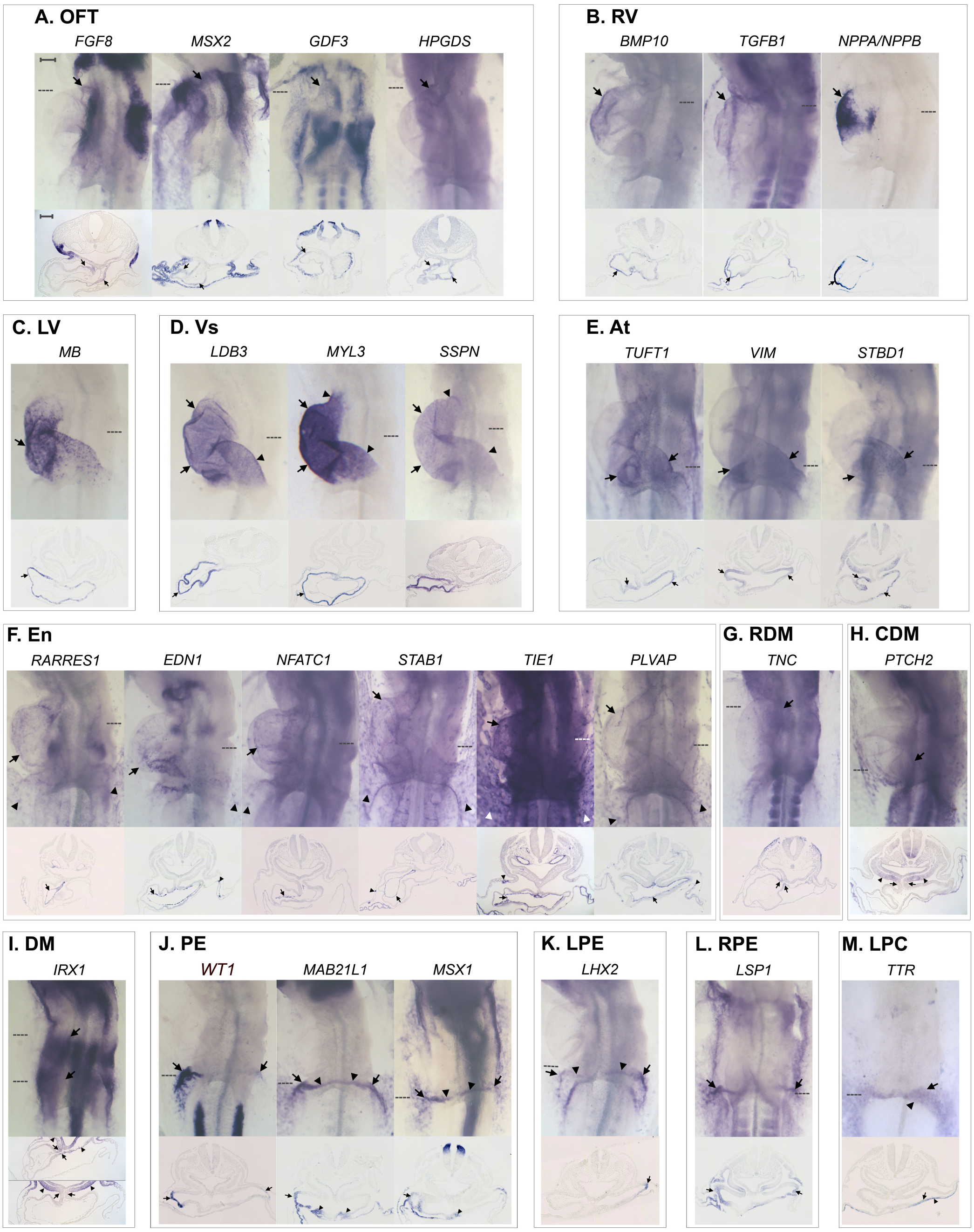


**Supplementary figure 2**. A further selection of regionalised heart markers. **A**. *FGF8*, *PRSS23, MSX2*, *GDF3/cVg1* and *HPGDS* are detected in the OFT (arrows). **B**. *BMP10* and *TGFBI* expression is restricted to the outer curvature of the prospective RV (arrows). *NPPA*/*NPPB* expression is observed in the RV (arrow), but also in the inner and outer curvatures of the prospective ventricle. **C**. *MB* is detected in the prospective LV. **D**. *LDB3, MYL3* and *SSPN* expression extends beyond the prospective ventricular region (arrows) into the atrial and OFT regions (arrowheads). **E**. *TUFT1, VIM* and *STBD1* localise to the atria (arrows). **F**. *RARRES1, EDN1, NFATC1, STAB1, TIE1* and *PLVAP* are detected in the endocardium (arrows) and endothelial cells (arrowheads); **G**. *TNC* expression is faint but detectable in the RDM (arrows). **H.** *PTCH2* is weakly detected in the CDM (arrows) and expression is continuous with the splanchnic mesoderm (arrowheads). **I.** *IRX1* in both the rostral and caudal DM (arrows) and the splanchnic mesoderm (arrowheads). **J.** *WT1* transcripts are detected in both the right and the left PE (arrows), as are *MAB21L1*

and *MSX1* (arrows) and also in ventro-medial cells (arrowheads). **K**. LPE marker *LHX2* is detected in both PE (arrows) and ventro-medial cells (arrowheads). **L**. RPE marker *LSP1* is detected in both PEs (arrows). **M**. *TTR,* a putative marker for the early LPC (arrows), is also expressed in the endoderm (arrowheads). At. Atria; En. Endocardium. Scale bars 100µm.

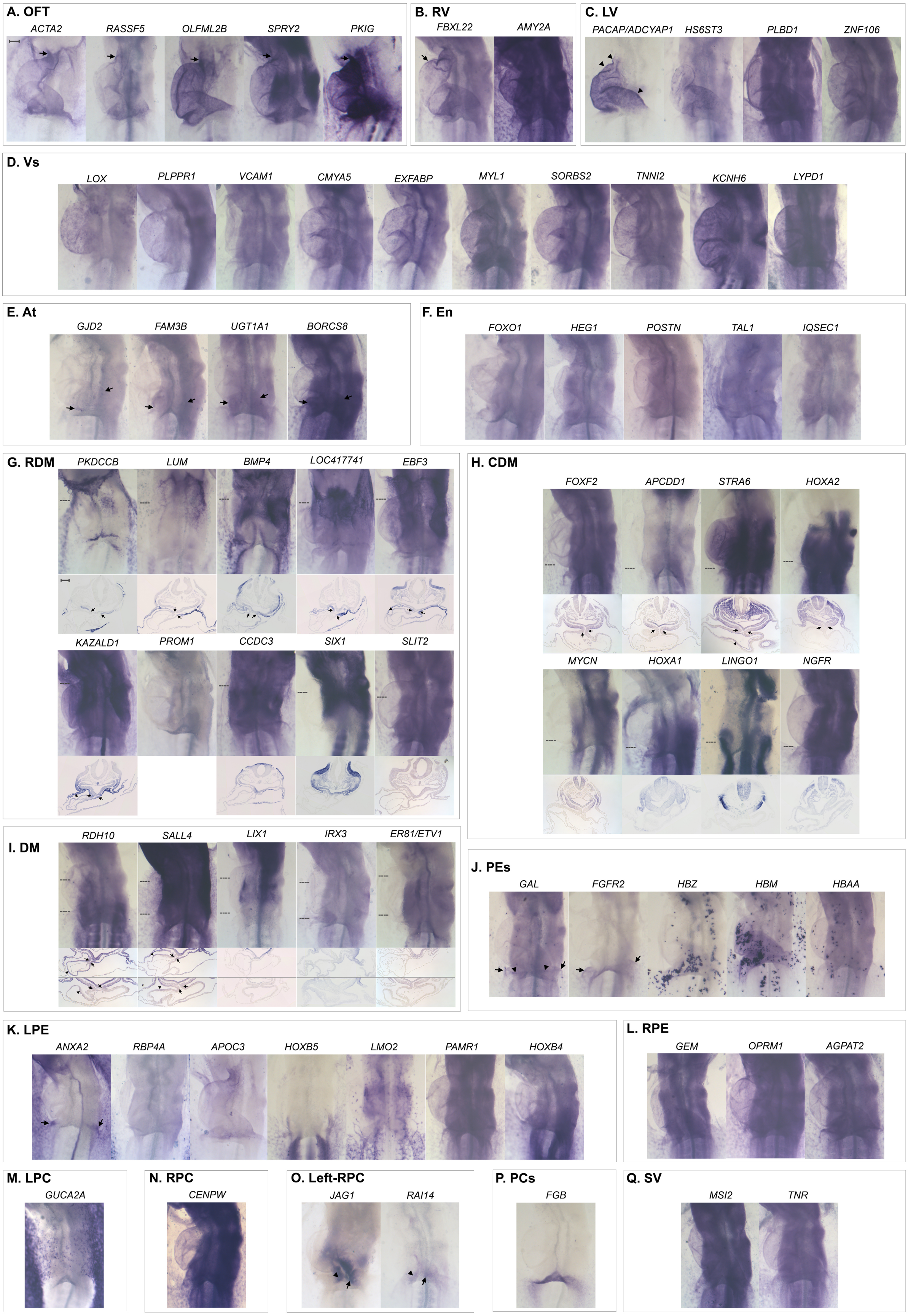


**<u>Supplementary figure 3</u>**. Markers with less regional specificity. **A**. *ACTA2*, *RASSF5*, *OLFML2B*, *SPRY2* and *PKIG* transcripts are enriched in the OFT (arrows), but their expression is not restricted to this region. **B**. *FBXL22* transcripts are enriched (arrows) but not restricted to the prospective RV. *AMY2A* expression is strong throughout the heart. **C**. *PACAP/ADCYAP1* expression is not restricted to the prospective LV. *HS6ST3* expression is faint and *PLBD1* and *ZNF106* expression is strong throughout the heart. **D**. *LOX* expression is faint in both the prospective right and left ventricles. *PLPPR1* and *VCAM1* expression is faint, whereas, *CMYA5, EXFABP, MYL1, SORBS2, TNNI2, KCNH6* and *LYPD1* are strong in the heart and/or embryo. **E**. *GJD2, FAM3B* and *UGT1A1* expression is faint in the atrial region (arrows). *BORCS8* expression is strong within the embryo and atrial region (arrows). **F**. Endocardial markers *FOXO1, HEG1, POSTN, TAL1* and *IQSEC1* are not detected. **G**. *PKDCCB, LUM, BMP4, LOC417741, EBF3* and *KAZALD1* expression in the RDM (arrows) is continuous with the splanchnic mesoderm. *KAZALD1* and *EBF3* are also in the myocardium (arrowheads). *PROM1*, *CCDC3, SIX1* and *SLIT2* are faint or not detected in the RDM. **H**. Expression of CDM markers *FOXF2, APCDD1, STRA6* and *HOXA2* are faint (arrows). *STRA6* is also detected in the endocardium (arrowhead). *MYCN, HOXA1, LINGO1* and *NGFR* are not detected in the CDM. **I.** *RDH10* and *SALL4* expression in the DM (arrows) is continuous with the splanchnic mesoderm and the myocardium (arrowheads). *LIX1*, *IRX3*, and *ER81/ETV1* are not detected in the DM. **J**. Faint expression of *GAL* and *FGFR2* is detected in both right and left PE (arrows), and *GAL* is detected in ventro-medial cells (arrowheads). *HBZ*, *HBM* and *HBAA* are not restricted to the PE. **K**. LPE marker *ANXA2* is detected in both PE (arrows). *RBP4A, APOC3, HOXB5, LMO2, PAMR1* and *HOXB4* are either faint or not expressed. **L**. *GEM, OPRM1* and *AGPAT2* are strongly expressed throughout the heart and are not specific to the RPE. **M**. *GUCA2A* transcripts are not detected in the LPC region. **N**. *CENPW* expression is strong throughout the heart and not localised to the RPC region. **O**. *JAG1* and *RAI14* are detected on both the left (arrow) and right (arrowhead) and is not restricted to the Left-RPC. **P**. Putative PC marker *FGB* is expressed in the AIP endoderm and not detected in either the early LPC or RPC. **Q**. *MSI2* and *TNR* are not restricted to the SV region. Scale bars 100μm.
