## Supplementary Methods for "A 3D molecular atlas of the chick embryonic heart"

**Embryos and dissection**

Fertile Brown Bovan Gold hens' eggs (Henry Stewart, UK) were incubated at 38°C for 45-50 hours to obtain Hamburger and Hamilton^1^ (HH) stage 12 (16 somites) embryos. Embryos were harvested either into ice cold Ca^2+^Mg^2+^-free PBS (CMF-PBS) containing 0.2mM EDTA for tissue separation, or into ice-cold Tyrode’s saline for tissue dissection. To separate the endocardium from ventricular tissue, 10µg/ml hyaluronidase from bovine testes (Sigma-Aldrich) was used. HH12 hearts were separated from surrounding splanchnic mesoderm (SM), foregut and anterior-intestinal portal (AIP) endoderm. Dissected tissues were collected and pooled in 200µl of ice-cold RNA stabilisation solution, RNAlater (Invitrogen) and stored at 4°C. Each sample comprised between 10-27 explants for RNA extraction, depending on the size of the region being collected.

**Definitions of 13 regions of the HH12 heart for tissue collection**

The following 13 tissue samples were collected (Figure 1A):

Outflow tract (OFT) - The myocardial part of the OFT anterior to the distal ventricular groove^2^ was dissected and the endocardium was removed. Addition of cells from the secondary/anterior heart fields to form definitive OFT occurs at looping stages^3-5^ (HH12-18).

Prospective ventricles (Vs) – The prospective right and left ventricles were collectively characterised by the expression of prospective ventricular marker *IRX4*^6^ (Figure 1B). Further growth of the prospective ventricular myocardium occurs after HH12^7, 8^.

Right ventricle (RV) – Fate maps indicate the position of the prospective RV at HH12^3, 9^, before major contribution from the anterior/secondary heart fields^3-5, 11^, and before the RV becomes clearly defined from left ventricle (LV) with the formation of the interventricular septum after HH17^10^. This was used to define the position of the RV

Left ventricle (LV) – Retrospective labelling using mouse embryos and fate maps in chick embryos indicate that most of the prospective ventricle at heart tube stages corresponds to the LV^12, 13^. The anterior limit of the LV was roughly defined by the anterior limit of *TBX5* expression^14^ (arrow, Figure 1C). The prospective atrioventricular canal (AVC) lies between the LV and atria^9^, but markers (e.g. *MSX2* and *BMP2*) are not regionally restricted at HH12 and therefore the AVC could not be taken for this study.

Endocardium (En) – The endocardial lining of both right and left prospective ventricles was collected.

Atria (At) – The selection of tissue selected to represent prospective atrium was guided partly by fate maps of the atrioventricular groove at HH12^9^ and *AMHC1/MYH7* expression (Figure 1D), which coincides with the atrial compartment throughout development^15^. Further growth of the atria is thought to derive from the *ISL1*-expressing caudal dorsal mesocardia (CDM) growth zone^7^.

Dorsal mesocardium (DM) – The DM initially links the heart tube to the dorsal body wall and is continuous with the splanchnic mesoderm (SM). At HH12, the DM ruptures in the mid-region and the heart tube is then joined to the SM by the rostral (RDM) and caudal (CDM) dorsal mesocardia^16^. The DM is characterised by the expression of secondary heart field marker, *ISL1*^17, 18^ (arrows, Figure 1E). The RDM and CDM were excised from both the SM and myocardium before being collected.

Left pacemaker (LPC) – Pacemaker activity originates from the left sinus venosus (SV) region in the HH10-17 chick heart^19^. To date there is no known specific marker for the early left pacemaker. The region dissected and used in this study was defined based on published DiI labelling and electrophysiological maps^20^.

Right pacemaker (RPC) – Pacemaking activity switches from the early left pacemaker to the right side at HH17-18^19^, eventually localising to the dorsal wall of the right atrium which forms the sinoatrial node (SAN)^20^. Expression of *Shox2*, a known marker of the SAN, coincides with SAN development in mouse embryos^21^. SAN precursors in chick embryos have been mapped to the right lateral plate mesoderm at HH8 (the tertiary heart field)^20^, but their location at HH12 is not known. The *SHOX2*-expressing domain in the dorso-caudal right atria at HH12 was taken as prospective RPC (arrows, Figure 1F), but a fate map of this region to confirm this is lacking.

Left-RPC – the left side corresponding to the position of the RPC (i.e. the mirror image of the RPC on the left side on the body) was collected to uncover possible right-specific RPC genes.

Proepicardium (PE) – In chick embryos, there is initially a right (RPE) and a left proepicardium^22^ (LPE) marked by the expression of *TBX18*^23^ (arrows, Figure 1G). The LPE is smaller than the RPE and disappears by HH18-19^22, 24^. RPE and LPE were collected separately.

Sinus venosus (SV) - The AIP endoderm and other tissues from the inflow tract of the heart tube (At, RPC, left-RPC, LPC, RPE and LPE) were removed and the remaining SV tissue collected. The SV region undergoes further growth from a caudal proliferation zone^7^.

**RNA extraction and sequencing**

RNA was extracted from collected tissue using the RNAqueous Micro-kit (Ambion, AM1931) with a slight modification. Briefly, an equal volume of cold PBS was added to the tissues in RNAlater and centrifuged at 500xg for 5 minutes. RNAlater and PBS were removed and tissues were lysed in 100µl lysis buffer with gentle pipetting, 125µl of 100% ethanol were added. The following steps were as per the manufacturer’s instructions, with the exception of the DNase treatment, which was extended to 60 minutes. Quality of the purified RNA samples was determined using Agilent 2200 TapeStation. Samples with an RNA Integrity number >7.8 were used to make libraries.

Libraries were constructed from 100ng total RNA using the NEBNext® Ultra™ II Directional RNA Library Prep Kit with NEBNext® Poly(A) mRNA Magnetic Isolation Module (NEB, E7760 and E7490) according to the manufacturer’s instructions. Libraries to be multiplexed in the same run were pooled in equimolar quantities, calculated from Qubit and Bioanalyzer fragment analysis. Samples were sequenced with the Illumina NextSeq 500 System (San Diego, US) (86bp single end, except for SV and left-RPC, which were paired-end-sequenced) which yielded >38 million reads per sample (Supplementary Dataset 1).

**Analysis**

Sequence reads were trimmed using trimmomatic-0.36^25^ and aligned to the galGal5 genome using TopHat2^26^, alignment rates were 83.9%±1.9%. Transcripts were counted and normalised using Cufflinks programmes *cuffquant* and *cuffnorm* respectively^27^. Data analysis was performed in the R environment. The matrix of transcript FPKMs (Fragments Per Kilobase of transcript per Million mapped reads), which contains expression of 15,570 genes in 13 samples was used to calculate differential gene expression (Supplementary Dataset 1). For most comparisons, a FPKM ≥20 and fold change (FC) ≥1.5 was used to select genes that are differentially expressed in a sample compared to the other 12 samples. Exceptions were made for either long or short lists of candidate genes, where the FC was increased or decreased respectively (red text in Supplementary Dataset 1 for FC values ≤1.5). To identify candidate markers that are differentially expressed in two regions (e.g. RDM and CDM), genes were selected with FPKM ≥20 and FC ≥1.5 in both regions compared to the other 11 samples. A total of 132 differentially expressed genes were selected for analysis using mRNA *in situ* hybridisation (ISH) (bold text in Supplementary Dataset 1). The FPKM of these selected genes were used to generate a hierarchical clustering heat map using R package *gplots* (Supplementary Figure 1).

The full dataset has been deposited in ArrayExpress ‘A 3D molecular atlas of the chick embryonic heart’, accession number E-MTAB-7663.

***In situ* hybridisation and image processing**

Embryos were fixed, processed and whole mount ISH was performed as described^28^. Most antisense probes were synthesised from EST clones^29^, others were kindly supplied from other sources (Table 1). Stained embryos were imaged on an Olympus SZH10 stereo-microscope and QImaging Retiga 2000R camera, then either processed for Optical Projection Tomography^30^ (OPT) or histological sectioning. OPT was performed using a 3001M OPT scanner and images reconstructed using NRecon (Bioptonics). 10µm paraffin sections were prepared on a Microm HM 315 microtome and scanned using Zeiss AxioScan Z1 slide scanner. Images were processed with Photoshop CS2 (Adobe) and ImageJ StackReg^31^.

Markers with restricted expression to the region of the heart collected for RNA-seq compared to the other 12 were considered ‘good’ markers for that region.

**Table 1.**

| **Ensembl ID** | **GENE** | **EST** | **Source** | **Comparison** |
| --- | --- | --- | --- | --- |
| ENSGALG00000006343 | ACTA2 | ChEST852h6 | Source Bioscience | OFT |
| ENSGALG00000011623 | ADAMTS3 | ChEST246g9 | Source Bioscience | LPC |
| ENSGALG00000014858 | ADCYAP1 | ChEST47n14 | Source Bioscience | LV |
| ENSGALG00000046593 | ADM | ChEST371m13 | Source Bioscience | A |
| ENSGALG00000023517 | AGPAT2 | ChEST730h13 | Source Bioscience | RPE |
| ENSGALG00000038740 | AMY2A | ChEST615c16 | Source Bioscience | RV |
| ENSGALG00000015148 | ANXA1 | ChEST160h21 | Source Bioscience | RDM |
| ENSGALG00000003770 | ANXA2 | ChEST175n21 | Source Bioscience | LPE |
| ENSGALG00000000894 | APCDD1 | ChEST872k3 | Source Bioscience | CDM |
| ENSGALG00000030920 | APOC3 | ChEST140j19 | Source Bioscience | LPE |
| ENSGALG00000004518 | BHMT | ChEST230l10 | Source Bioscience | E |
| ENSGALG00000000120 | BMP10 | ChEST1005c18 | Source Bioscience | RV |
| ENSGALG00000012429 | BMP4 |  | K Liem | RDM |
| ENSGALG00000014581 | BORCS8 | ChEST115h21 | Source Bioscience | A |
| ENSGALG00000030140 | CCDC3 | ChEST871g16 | Source Bioscience | RDM |
| ENSGALG00000027454 | CENPW | ChEST828o5 | Source Bioscience | RPC |
| ENSGALG00000026553 | CMYA5 | ChEST990b12 | Source Bioscience | Vs |
| ENSGALG00000004631 | DRAXIN | ChEST545l1 | Source Bioscience | RV |
| ENSGALG00000010461 | EBF3 |  | A Vincent | RDM |
| ENSGALG00000012735 | EDN1 | ChEST789p9 | Source Bioscience | E |
| ENSGALG00000032997 | EMILIN2 | ChEST359g19 | Source Bioscience | LV |
| ENSGALG00000035504 | ER81 (ETV1) |  | K Storey/J Lin | DMs |
| ENSGALG00000043064 | EXFABP | ChEST242g23 | Source Bioscience | Vs |
| ENSGALG00000016140 | FAM3B | ChEST757l22 | Source Bioscience | A |
| ENSGALG00000003401 | FBXL22 | ChEST64k4 | Source Bioscience | RV |
| ENSGALG00000009262 | FGB | ChEST159e17 | Source Bioscience | PCs |
| ENSGALG00000007706 | FGF8 |  | S Noji | OFT |
| ENSGALG00000009495 | FGFR2 |  | E Pasquale | PEs |
| ENSGALG00000015708 | FGFR3 |  | E Pasquale | PEs |
| ENSGALG00000000894 | FOXF2 | ChEST39c1 | Source Bioscience | CDM |
| ENSGALG00000017034 | FOXO1 | ChEST232b16 | Source Bioscience | E |
| ENSGALG00000007047 | GAL | ChEST825b20 | Source Bioscience | PEs |
| ENSGALG00000043371 | GDF3 (cVg1) |  |  | OFT |
| ENSGALG00000032506 | GEM | ChEST624g12 | Source Bioscience | RPE |
| ENSGALG00000009845 | GJD2 |  | V Berthoud | A |
| ENSGALG00000027269 | GUCA2A | ChEST152d9 | Source Bioscience | LPC |
| ENSGALG00000043234 | HBAA | ChEST973g24 | Source Bioscience | PEs |
| ENSGALG00000031597 | HBM/HBAD | ChEST382l19 | Source Bioscience | PEs |
| ENSGALG00000023740 | HBZ | ChEST53k6 | Source Bioscience | PEs |
| ENSGALG00000011813 | HEG1 | ChEST476a7 | Source Bioscience | E |
| ENSGALG00000028095 | HOXA1 | ChEST1010d14 | Source Bioscience | CDM |
| ENSGALG00000040521 | HOXA2 | ChEST671c8 | Source Bioscience | CDM |
| ENSGALG00000000284 | HOXB4 |  | R Krumlauf | LPE |
| ENSGALG00000034907 | HOXB5 |  |  | LPE |
| ENSGALG00000010402 | HPGDS | ChEST153g19 | Source Bioscience | OFT |
| ENSGALG00000025906 | HS6ST3 | ChEST935c22 | Source Bioscience | LV |
| ENSGALG00000036916 | IQSEC1 | ChEST666n9 | Source Bioscience | E |
| ENSGALG00000030991 | IRX1 |  | K Storey | DMs |
| ENSGALG00000042045 | IRX3 |  | K Shimamura | DMs |
| ENSGALG00000022896 | IRX4 |  | ZZ Bao | Vs |
| ENSGALG00000014884 | ISL1 |  | T Jessell/L Gunhaga | RDM |
| ENSGALG00000009020 | JAG1 |  | HC Lee | Left-RPC |
| ENSGALG00000007837 | KAZALD1 | ChEST70d11 | Source Bioscience | RDM |
| ENSGALG00000000505 | KCNH6 | ChEST674p14 | Source Bioscience | Vs |
| ENSGALG00000003939 | KLF2 |  | P Antin | E |
| ENSGALG00000044919 | CR385930 | ChEST350d20 | Source Bioscience | OFT |
| ENSGALG00000001977 | LDB3 | ChEST299f15 | Source Bioscience | Vs |
| ENSGALG00000001124 | LHX2 |  | T Nohno | LPE |
| ENSGALG00000002708 | LINGO1 | ChEST905c4 | Source Bioscience | CDM |
| ENSGALG00000015290 | LIX1 |  | S Faure | DMs |
| ENSGALG00000029892 | LMO2 |  | T Jaffredo | LPE |
| ENSGALG00000008635 | LOC417741 | ChEST714o24 | Source Bioscience | RDM |
| ENSGALG00000028063 | LOX | ChEST374f21 | Source Bioscience | Vs |
| ENSGALG00000006583 | LSP1 | ChEST978m2 | Source Bioscience | RPE |
| ENSGALG00000011271 | LUM | ChEST85i2 | Source Bioscience | RDM |
| ENSGALG00000012176 | LYPD1 | ChEST272k7 | Source Bioscience | Vs |
| ENSGALG00000045052 | MAB21L1 | ChEST386j5 | Source Bioscience | PEs |
| ENSGALG00000012541 | MB | ChEST501h20 | Source Bioscience | LV |
| ENSGALG00000036743 | MGAT3 | ChEST45g15 | Source Bioscience | LV |
| ENSGALG00000005567 | MSI2 | ChEST593o8 | Source Bioscience | SV |
| ENSGALG00000015013 | MSX1 |  |  | PEs |
| ENSGALG00000038848 | MSX2 | ChEST410j10 | Ark Genomics | OFT |
| ENSGALG00000016462 | MYCN | ChEST442N13 | Ark Genomics | CDM |
| ENSGALG00000015358 | MYH15 (VMHC1) |  | K Yutzey / D Bader | Vs |
| ENSGALG00000035594 | MYH7 (AMHC1) |  | T Brand | A |
| ENSGALG00000002907 | MYL1 | ChEST910o13 | Source Bioscience | Vs |
| ENSGALG00000005448 | MYL3 | ChEST220a4 | Source Bioscience | Vs |
| ENSGALG00000042534 | NFATC1 | ChEST526a21 | Source Bioscience | E |
| ENSGALG00000035950 | NGFR | ChEST998b1 | Source Bioscience | CDM |
| ENSGALG00000010237 | NPC2 | ChEST354l19 | Source Bioscience | E |
| ENSGALG00000004574 | NPPB (NPPA) | ChEST509m15 | Source Bioscience | RV |
| ENSGALG00000002667 | OLFML2B | ChEST64m16 | Source Bioscience | OFT |
| ENSGALG00000013616 | OPRM1 | ChEST659n20 | Source Bioscience | RPE |
| ENSGALG00000007886 | PAMR1 | ChEST911n19 | Source Bioscience | LPE |
| ENSGALG00000011166 | PKDCCB | ChEST195h16 | Source Bioscience | RDM |
| ENSGALG00000029583 | PLBD1 | ChEST631m10 | Source Bioscience | LV |
| ENSGALG00000027583 | PLN | ChEST53o9 | Source Bioscience | Vs |
| ENSGALG00000015542 | PLPPR1 | ChEST302e18 | Source Bioscience | Vs |
| ENSGALG00000045684 | PLVAP | ChEST353n11 | Source Bioscience | E |
| ENSGALG00000017046 | POSTN | ChEST61a21 | Source Bioscience | E |
| ENSGALG00000014496 | PROM1 | ChEST671b22 | Ark Genomics | RDM |
| ENSGALG00000017244 | PRSS23 | ChEST621i14 | Source Bioscience | OFT |
| ENSGALG00000010133 | PTCH2 |  | M Davey | CDM |
| ENSGALG00000012522 | PVALB | ChEST446a17 | Source Bioscience | RPC |
| ENSGALG00000003353 | RAI14 |  | C Tabin | Left-RPC |
| ENSGALG00000009594 | RARRES1 | ChEST790i12 | Source Bioscience | E |
| ENSGALG00000038943 | RASSF5 | ChEST453k15 | Source Bioscience | OFT |
| ENSGALG00000006629 | RBP4A | ChEST158a6 | Source Bioscience | LPE |
| ENSGALG00000034346 | RDH10 |  | M Maden | DMs |
| ENSGALG00000042607 | RSPO3 | ChEST784h18 | Ark Genomics | A |
| ENSGALG00000041365 | SALL4 |  | M Bronner | DMs |
| ENSGALG00000039728 | SHOX2 |  | G Rappold | RPC |
| ENSGALG00000029401 | SIX1 |  | P Gruss | RDM |
| ENSGALG00000041121 | SLIT2 | ChEST1007n17 | Source Bioscience | RDM |
| ENSGALG00000011205 | SMOC2 | ChEST527i5 | Source Bioscience | CDM |
| ENSGALG00000018557 | SOD3 | ChEST111b2 | Source Bioscience | A |
| ENSGALG00000030769 | SORBS2 | ChEST387n4 | Source Bioscience | Vs |
| ENSGALG00000016906 | SPRY2 |  | A Streit | OFT |
| ENSGALG00000014042 | SSPN | ChEST860p7 | Source Bioscience | Vs |
| ENSGALG00000007361 | SST | ChEST114e9 | A Streit | LPE |
| ENSGALG00000043672 | STAB1 | ChEST267o12 | Source Bioscience | E |
| ENSGALG00000027872 | STBD1 | ChEST728f8 | Source Bioscience | A |
| ENSGALG00000001449 | STRA6 |  | M Maden | CDM |
| ENSGALG00000023296 | TAL1 | ChEST56l11 | Source Bioscience | E |
| ENSGALG00000032789 | TBX18 |  | M Maroto | RPE |
| ENSGALG00000043815 | TENM4 | ChEST262c14 | Source Bioscience | RPE |
| ENSGALG00000029429 | TGFBI | ChEST551m21 | Source Bioscience | RV |
| ENSGALG00000009957 | TIE1 |  | Rusty Lansford | E |
| ENSGALG00000028627 | TIMP3 | ChEST177g6 | Source Bioscience | RV |
| ENSGALG00000039990 | TNC | ChEST681l9 | Source Bioscience | RDM |
| ENSGALG00000006591 | TNNI2 | ChEST162i22 | Source Bioscience | Vs |
| ENSGALG00000004526 | TNR | ChEST844g10 | Source Bioscience | SV |
| ENSGALG00000015143 | TTR | ChEST578j20 | Source Bioscience | LPC |
| ENSGALG00000032955 | TUFT1 | ChEST517o13 | Source Bioscience | A |
| ENSGALG00000004196 | UGT1A1 | ChEST90b1 | Source Bioscience | A |
| ENSGALG00000005257 | VCAM1 | ChEST77f24 | Source Bioscience | Vs |
| ENSGALG00000008677 | VIM | ChEST79m10 | Source Bioscience | A |
| ENSGALG00000034168 | WNT5A |  | T Brown | DMs |
| ENSGALG00000011358 | WNT6 |  |  | A |
| ENSGALG00000012115 | WT1 |  | T Mikawa | PEs |
| ENSGALG00000009081 | ZNF106 | ChEST224l1 | Source Bioscience | LV |

**Generating a 3D atlas of expression**

The atlas consists of a 3D reference model of the HH12 chick heart, onto which a collection of OPT assays were mapped. The model itself has three components; a segmented volumetric image corresponding to the heart territories domain, a surface mesh and a volumetric mesh. The volumetric image was segmented from a representative HH12 embryo that had undergone ISH without an antisense probe. The heart of this embryo was then used to define the meshes. Because of the complex spatial mapping transformations needed for the assays and the confounding nature of the expression signal it was not possible to use an automated image registration approach, a manually guided method was used based on Constrained Distance Transform (CDT)^32^. In the CDT, transformations are computed using distances evaluated along paths constrained to the model domain a using a conforming mesh. This approach allows OPT assays with variations in pose and requiring large deformation to be mapped. The mapping tool, WlzWarp, ([https://github.com/ma-tech/WlzQtApps](https://github.com/ma-tech/WlzQtApps" \t "_blank)) was used to define CDTs for each of the assays, spatially mapping them to the atlas reference model. Following image registration, the assay images were segmented and processed to remove noise and extraneous features. For visualization, point clouds were extracted from the mapped assay images, with the point value and density being determined by the assay image value. The Woolz image processing system ([https://github.com/ma-tech/Woolz](https://github.com/ma-tech/Woolz" \t "_blank)) and associated tools was used for segmentation, mesh and point cloud generation.
