## Supplementary Table 1 for "A 3D molecular atlas of the chick embryonic heart"

| Region from RNAseq | Transcript | Specific to OFT within heart? | Expressed outside of the heart? | Novel marker to OFT? | Published expression in OFT | Published expression in heart outside of the OFT |
|---|---|---|---|---|---|---|
| Outflow Tract (OFT) | *ACTA2* | no | yes | no | Expressed in 56 hours post fertilisation (hpf) zebrafish OFT[1] | Expressed in inflow and outflow tract in Hamburger Hamilton[2] (HH) stage 15 chick embryos[3] |
| OFT | *FGF8* | yes | yes | yes | | mRNA detected in embryonic day (E)8.5-9 mouse splanchnic mesoderm[4]. *fgf8/ace* is expressed throughout the zebrafish ventricle at 36hpf[5]. *FGF8* mRNA is not detected in the HH16 chick embryo OFT[6] |
| OFT | *GDF3/ cVg1* | yes | yes | yes | | Expressed in prospective ventricle in HH13 chick[7] |
| OFT | *HPGDS* | yes | yes | yes | | |
| OFT | *CR385930 / LBH-LIKE* | yes | yes | yes | | |
| OFT | *MSX2* | yes | yes | no | Detected in E9.5 mouse OFT[8] | |
| OFT | *OLFML2B* | no | yes | no | | |
| OFT | *PRSS23* | yes | yes | yes | | mRNA detected in the zebrafish atrium, ventricle and atrioventricular canal at 48hpf[9] |
| OFT | *RASSF5* | no | yes | no | | |
| OFT | *SPRY2* | no | yes | no | Expressed in splanchnic mesoderm and OFT in E8.5 mouse embryos[10] | |
| OFT | *PKIG* | no | yes | no | | |
| **Region from RNAseq** | **Transcript** | **Specific to RV within heart?** | **Expressed outside of the heart at HH12?** | **Novel marker to RV?** | **Published expression in RV** | **Published expression in heart outside of the RV** |
| Right ventricle (RV) | *AMY2A* | no | yes | no | | |
| RV | *BMP10* | yes | no | yes | | Transcripts detected throughout the prospective ventricle at HH10, and in OFT, Vs, and embryonic atria in HH14 chick embryos[11]. Expressed in E9.0 mouse ventricle and bulbus cordis (OFT region)[12] |

| | | | | | | |
|---|---|---|---|---|---|---|
| RV | *DRAXIN* | yes | yes | yes | Expressed in the rostral part of the prospective ventricle in HH15 chick heart[13] | |
| RV | *FBXL22* | no | yes | no | | |
| RV | *NPPA/ NPPB* | no | no | no | *NPPB*[14] expressed in prospective ventricle of E8.5 mouse and the rostral part of the HH12-13 chick prospective ventricle[7, 15] | Expressed throughout stage 32 *Xenopus* heart tube[16] |
| RV | *TGFBI* | yes | yes | yes | | mRNA is expressed in E10.0-10.5 mouse OFT and endocardial cushions of the AV canal[17, 18] |
| RV | *TIMP3* | no | yes | no | | |

| **Region from RNAseq** | **Transcript** | **Specific to LV within heart?** | **Expressed outside of the heart at HH12?** | **Novel marker to LV?** | **Published expression in LV** | **Published expression in heart outside of the LV** |
|---|---|---|---|---|---|---|
| Left ventricle (LV) | *ADCYAP1 / PACAP* | no | no | no | | |
| LV | *EMILIN2* | yes | no | no | *emilin2b* is expressed in the zebrafish ventricle at 40hpf[19]. Expression is detected in the E10.5 mouse ventricle[20] | *emilin2a* is expressed in the zebrafish atrium at 48hpf[19] |
| LV | *HS6ST3* | yes | yes | yes | | |
| LV | *MB* | yes | no | no | Detected in E9.5 mouse ventricles[21] and 48hpf zebrafish heart[22] but regionalisation is not clear in either | |
| LV | *MGAT3* | yes | yes | yes | | |
| LV | *PLBD1* | no | yes | yes | | |
| LV | *ZNF106* | no | yes | yes | | |

| **Region from RNAseq** | **Transcript** | **Specific to Vs within heart?** | **Expressed outside of the heart at HH12?** | **Novel marker to Vs?** | **Published expression in Vs** | **Published expression in heart outside of the Vs** |
|---|---|---|---|---|---|---|
| Ventricles (Vs) | *CMYA5* | no | yes | no | | Expressed in both the atria and ventricles of E10.5 mouse[23] |
| Vs | *EXFABP* | no | yes | no | | |
| Vs | *IRX4* | yes | no | no | Expression is detected in prospective ventricles of HH9+ chick[24] and E8.5 mouse embryos[25] | |

| | | | | | | |
|---|---|---|---|---|---|---|
| Vs | *KCNH6/ ERG* | no | yes | no | Weak expression detected in HH11 quail heart[26] but regionalisation is not clear | |
| Vs | *LDB3* | no | no | no | | |
| Vs | *LOX* | no | yes | no | Expressed in E11.5 mouse ventricle[27] | |
| Vs | *LYPD1* | no | yes | no | | |
| Vs | *MYH15/ VMHC1* | no | no | no | A *MYH15/VMHC1* specific mRNA in situ hybridisation probe is restricted to the prospective ventricle of HH12 chick embryos[28]. Zebrafish *vmhc* is detected in the 24-48hpf prospective ventricle[29, 30] | mRNA in situ hybridisation probes that recognise both *MYH15/VMHC1* and *MYH7b* are detected throughout the HH12 chick heart[28, 31, 32]. |
| Vs | *MYL1/ MLC3f* | no | yes | no | | Expressed in both the ventricle and inflow region (atria and sinus venosus) at E8.5 in mouse embryos[33] |
| Vs | *MYL3/ MLC1V* | no | no | no | | Expression is detected throughout the E8.0 mouse heart tube and in atria and ventricles at E9.5[34] |
| Vs | *PLN* | no | no | no | Expressed in HH15 chick ventricle[35] but regionalisation is not clear | |
| Vs | *PLPPR1* | no | yes | no | | |
| Vs | *SORBS2* | no | yes | no | | |
| Vs | *SSPN* | no | yes | no | | |
| Vs | *TNNI2* | no | yes | no | | |
| Vs | *VCAM1* | no | yes | no | Protein is detected in the prospective ventricle of E8.75 mouse embryos[36] | |
| **Region from RNAseq** | **Transcript** | **Specific to At. within heart?** | **Expressed outside of the heart at HH12?** | **Novel marker to At?** | **Published expression in At.** | **Published expression in heart outside of the At.** |
| Atria (At) | *ADM* | no | no | no | | Protein is detected in both the atria and ventricles of E9 mouse embryos[37] |
| At | *BORCS8* | no | yes | no | | |
| At | *FAM3B* | no | yes | no | | |
| At | *GJD2* | no | no | no | | |
| At | *MYH7/ AMHC1* | yes | no | no | Expressed in prospective atria in HH14 chick embryos[32]. *MHCa/Myh6* in mouse is expressed in the caudal part of the E8.0 heart tube (atrioventricular junction)[34]. Zebrafish *amhc* is expressed in the prospective atrium at 24-48hpf[29] | By E9.5 in mouse embryos, *MHCα/Myh6* transcripts are also detected in the ventricle as well as the atrium[34] |

| | | | | | | |
|---|---|---|---|---|---|---|
| At | *RSPO3* | yes | yes | no | *RSPO3* transcripts are detected in the caudal pole of the E8.5 mouse heart[38], but it's localisation to the atria is unclear | By E9.5-E10.5 of mouse heart development expression is observed in the atrioventricular canal and OFT[38] |
| At | *SOD3* | yes | no | yes | | |
| At | *STBD1* | yes | yes | yes | | |
| At | *TUFT1* | yes | yes | yes | | |
| At | *UGT1A1* | no | yes | no | | |
| At | *VIM* | yes | yes | no | Protein detected in HH12 chick myocardium, but regionalisation is unclear[39] | Protein detected in atrial myocardium but also OFT endocardium and epicardium in HH21 chick heart[40] |
| At | *WNT6* | yes | yes | yes | | Expressed in the atrioventricular junction in chick embryos at HH17[41]. XWnt6 protein is detected in the endocardium at stage 32, and in atrial and ventricular myocardium, as well as the epicardium, in stage 38-42 *Xenopus* embryos[42]. |
| **Region from RNAseq** | **Transcript** | **Specific to En. within heart?** | **Expressed outside of the heart at HH12?** | **Novel marker to En?** | **Published expression in En.** | **Published expression in heart outside of the En.** |
| Endo-cardium (En) | *BHMT* | yes | yes | yes | | |
| En | *EDN1* | yes | yes | yes | | *Edn1* expressed in OFT of E10.0 mouse heart[43] |
| En | *FOXO1* | no | yes | no | | Protein is detected in the myocardium as well as the endocardium of E10.5 mouse embryos[44] |
| En | *HEG1* | no | yes | no | Expressed in E10.5 mouse endocardium[45] | |
| En | *IQSEC1* | no | yes | no | | |
| En | *KLF2* | yes | no | no | Expression is detected in the endocardium of E8.5 mouse embryos[46] | |
| En | *NFATC1* | yes | yes | no | Expressed in the endocardium of E8.5 mouse embryos[47] | In addition to the endocardium, expression is detected in the vitelline vein (sinus venosus) region of E8.5 mouse embryos[47] |
| En | *NPC2* | yes | no | yes | | |
| En | *PLVAP* | yes | yes | yes | | |
| En | *POSTN* | no | yes | no | Expression starts to be detectable in the RV endocardium from HH17 in chick embryos[48] | Expressed in E10.0-10.5 mouse embryo OFT and endocardial cushions of the AV canal[17, 18] |
| En | *RARRES1* | yes | yes | yes | | |
| En | *STAB1* | yes | yes | no | Endocardial expression is detected in E10.0-5 mouse embryos[17, 18] | |

| | | | | | | |
|---|---|---|---|---|---|---|
| En | *TAL1* | no | yes | no | | *TAL1* is detected in blood Islands and blood cells at HH10-11, and in endothelial cells of the aorta at ~HH18, but localisation in the heart is not clear[49, 50] |
| En | *TIE1* | yes | yes | no | Expressed in HH12 chick endothelial cells and endocardium[51] and the endocardium in 48hpf zebrafish embryos[52] | |

| **Region from RNAseq** | **Transcript** | **Specific to RDM?** | **Expressed outside of the heart at HH12?** | **Novel marker to RDM?** | **Published expression in RDM** | **Published expression in heart outside of the RDM** |
|---|---|---|---|---|---|---|
| Rostral dorsal meso-cardium (RDM) | *ANXA1* | yes | yes | yes | | |
| RDM | *BMP4* | no | yes | no | *BMP4* expression is detected in the RDM in HH16 chick embryos[6]. *Bmp4* transcripts are present in the dorsal mesocardium of E9.5 mouse embryos[53] | In addition to the RDM, *BMP4* is detected in the caudal dorsal mesocardium and is continuous with the splanchnic mesoderm in HH16 chick embryos[6]. *XBmp4* transcripts are detected throughout the heart tube at stage 32 and in the dorsal presumptive atrium and ventricle at stage 35[54]. In 24hpf zebrafish embryos, *bmp4* is expressed on the left side of the atrium and ventricle[55] |
| RDM | *CCDC3* | no | yes | no | | |
| RDM | *EBF3* | no | yes | no | | |
| RDM | *ISL1* | no | yes | no | Expression is detected in the RDM of HH13 chick embryos[56] | Expression is detected in both RDM and CDM of HH13 chick embryos[56]. Expressed in the splanchnic mesoderm of E8.5 mouse embryos[57]. |
| RDM | *KAZALD1* | no | yes | no | | |
| RDM | *LOC417741* | no | yes | no | | |
| RDM | *LUM* | no | yes | no | | |
| RDM | *PKDCCB* | no | yes | no | | |
| RDM | *PROM1* | no | yes | no | | |
| RDM | *SIX1* | no | yes | no | | *Six1* transcripts are detected in the splanchnic mesoderm of E8.5 mouse embryos[58] |
| RDM | *SLIT2* | no | yes | no | | |
| RDM | *TNC* | yes | yes | yes | | *Tnc* is observed in the E9.5 mouse proepicardium[59] |

| Region from RNAseq | Transcript | Specific to CDM? | Expressed outside of the heart at HH12? | Novel marker to CDM? | Published expression in CDM | Published expression in heart outside of the CDM |
|---|---|---|---|---|---|---|
| Caudal dorsal meso-cardium (CDM) | *APCDD1* | no | yes | no | | |
| CDM | *FOXF2* | no | yes | no | | mRNA detected in hindgut splanchnic mesoderm at E8.5, and foregut and hindgut splanchnic mesoderm at E9.5 in mouse embryos[60] |
| CDM | *HOXA1* | no | yes | no | | *Hoxa1* transcripts are detected in the splanchnic mesoderm adjacent to the CDM in E8.5 mouse embryos[61] |
| CDM | *HOXA2* | no | yes | no | | |
| CDM | *LINGO1* | no | yes | no | | |
| CDM | *MYCN* | no | yes | no | | Detected in the myocardium of E9.5 mouse embryos[62] and HH20 chick embryos[63] |
| CDM | *NGFR* | no | yes | no | | |
| CDM | *PTCH2* | no | yes | no | | Expressed in the splanchnic mesoderm and dorsal mesocardia of E8.5-9.5 mouse embryos[64] |
| CDM | *SMOC2* | no | yes | no | | Smoc2 protein is detected in E14.5 mouse atrial and ventricular myocardium[65] |
| CDM | *STRA6* | no | yes | no | | |

| Region from RNAseq | Transcript | Specific to DM? | Expressed outside of the heart at HH12? | Novel marker to DM? | Published expression in DM | Published expression in heart outside of the DM |
|---|---|---|---|---|---|---|
| Dorsal meso-cardium (DM) | *ER81/ ETV1* | no | yes | no | | |
| DM | *IRX1* | no | yes | no | | *Irx1* mRNA expression first detected in E10.5-11.5 mouse ventricular septum[66] |

| | | | | | | |
|---|---|---|---|---|---|---|
| DM | *IRX3* | no | yes | no | | *Irx3* expression is first detected in E9.5 mouse ventricles[66] |
| DM | *LIX1* | no | yes | no | | |
| DM | *RDH10* | no | yes | no | | |
| DM | *SALL4* | no | yes | no | | |
| DM | *WNT5A* | no | yes | no | | Detected in the ventricle of HH20 chick embryos[67]. *Wnt5a* transcripts are observed in the OFT and RV myocardium in E8.5 mouse embryos[68] |
| **Region from RNAseq** | **Transcript** | **Specific to PEs?** | **Expressed outside of the heart at HH12?** | **Novel marker to PE?** | **Published expression in PEs** | **Published expression in heart outside of the PEs** |
| Proepi-cardium (PE) | *FGFR2* | yes | yes | no | *FGFR2* mRNA is detected in the RPE of HH17-18 chicken embryos[69] (by this stage the LPE has degenerated[69, 70]) | |
| PE | *FGFR3* | yes | yes | yes | | *FGFR3* mRNA not detected in the RPE of HH17-18 chicken embryos[69] |
| PE | *GAL* | yes | yes | yes | | |
| PE | *HBAA* | no | yes | yes | | |
| PE | *HBM /HBAD* | no | yes | no | | |
| PE | *HBZ* | no | yes | no | | |
| PE | *MAB21L1* | yes | yes | yes | | *MAB21L1* transcripts are not detected in mouse embryonic heart[71] |
| PE | *MSX1* | yes | yes | no | Transcripts are detected in the epicardium of HH23 chick embryos[72] | Expressed in endocardial cushions of the atrioventricular canal in HH15 chick embryos[72] |
| PE | *WT1* | yes | yes | no | | Found to be expressed only in RPE of HH13 chick embryos[73] |
| **Region from RNAseq** | **Transcript** | **Specific to LPE?** | **Expressed outside of the heart at HH12?** | **Novel marker to LPE?** | **Published expression in LPE** | **Published expression in heart outside of the LPE** |
| Left | *ANXA2* | no | yes | no | | Expression is detected in the endocardium of HH15 chick embryos[74]. mRNA is |

| | | | | | | |
|---|---|---|---|---|---|---|
| proepi-cardium (LPE) | | | | | | detected in the proepicardium of HH19-20 chick embryos[75] (by this stage the LPE has degenerated[69, 70]). |
| LPE | *APOC3* | no | yes | no | | |
| LPE | *HOXB4* | no | yes | no | | |
| LPE | *HOXB5* | no | yes | no | | |
| LPE | *LHX2* | no | yes | no | | *LHX2* expression is detected in the caudal pole of the HH18-20 chick heart[76], although localisation is unclear. LHX2 localises to the E9.5 mouse PE[77]. Transcripts are not detected in *Xenopus* PE at stages 32-46[78]. |
| LPE | *LMO2* | no | yes | no | | *LMO2* is detected in blood Islands at HH10-11 endothelial cells of the aorta at ~HH18[49, 50], but localisation in the heart is not clear |
| LPE | *PAMR1* | no | yes | no | | |
| LPE | *RBP4A* | no | no | no | | |
| LPE | *SST* | yes | yes | yes | | |

| **Region from RNAseq** | **Transcript** | **Specific to RPE?** | **Expressed outside of the heart at HH12?** | **Novel marker to RPE?** | **Published expression in RPE** | **Published expression in heart outside of the RPE** |
|---|---|---|---|---|---|---|
| Right proepi-cardium (RPE) | *AGPAT2* | no | yes | no | | |
| RPE | *GEM* | no | yes | no | | |
| RPE | *LSP1* | no | yes | no | mRNA is detected in the proepicardium of HH19-20 chick embryos[75] | Expression is detected in cardiac fibroblasts in E14.5 mouse embryos[79] |
| RPE | *OPRM1* | no | yes | no | | |
| RPE | *TBX18* | no | yes | no | *TBX18* is detected in the RPE of HH11-13 chick embryos[73] | Transcripts are detected in both the right and the left PE in HH11-12 chick embryos[80] and in E8.5 mouse embryos[70]. In another study, *TBX18* is not only detected in the RPE but also throughout the prospective atrial region in HH11-13 chick embryos[73]. |
| RPE | *TENM4* | yes | yes | yes | | mRNA expression in E8.5 mouse embryos is detected in the head folds, posterior somites and presomitic mesoderm [81] |

| Region from RNAseq | Transcript | Specific to LPC? | Expressed outside of the heart at HH12? | Novel marker to LPC? | Published expression in LPC | Published expression in heart outside of the LPC |
|---|---|---|---|---|---|---|
| Left pace-maker (LPC) | *ADAMTS3* | yes? | yes | yes | | |
| LPC | *GUCA2A* | no | yes | no | | |
| LPC | *TTR* | yes? | yes | yes | | Detected in endoderm of the anterior intestinal portal in HH8-14 chick embryos[82, 83] and in the myocardium of HH14 check embryos[83], although regionalisation is unclear. |
| **Region from RNAseq** | **Transcript** | **Specific to RPC?** | **Expressed outside of the heart at HH12?** | **Novel marker to RPC at HH12?** | **Published expression in RPC** | **Published expression in heart outside of the RPC** |
| Right pace-maker (RPC) | *CENPW* | no | yes | no | | |
| RPC | *PVALB* | yes | no | yes | | |
| RPC | *SHOX2* | yes | no | yes | | Initially expressed in the E8.5 mouse sinus venosus eventually restricting to the right dorsal atria (SAN region)[84]. Detected in the sinus venosus region of HH19 chick embryos[85]. *shox2* is detected in the 48hpf zebrafish sinoatrial region[86] |
| **Region from RNAseq** | **Transcript** | **Specific to Left-RPC?** | **Expressed outside of the heart at HH12?** | **Novel marker to Left-RPC?** | **Published expression in Left-RPC** | **Published expression in heart outside of the Left-RPC** |
| Left-RPC | *JAG1* | no | yes | no | | *Jag1* expression is detected in the E10.5 mouse pharyngeal endoderm and mesenchyme, OFT[87], atria and RV[88] |
| Left-RPC | *RAI14* | no | yes | no | | |

| Region from RNAseq | Transcript | Specific to PCs? | Expressed outside of the heart at HH12? | Novel marker to PCs? | Published expression in PC | Published expression in heart outside of the PC |
|---|---|---|---|---|---|---|
| Pace-makers (PCs) | *FGB* | no | yes | no | | |
| **Region from RNAseq** | **Transcript** | **Specific to SV?** | **Expressed outside of the heart at HH12?** | **Novel marker to SV?** | **Published expression in SV** | **Published expression in heart outside of the SV** |
| Sinus venosus (SV) | *MSI2* | no | yes | no | | |
| SV | *TNR* | no | yes | no | | |

75. Bressan M, Henley T, Louie JD, Liu G, Christodoulou D, Bai X, Taylor J, Seidman CE, Seidman JG, Mikawa T. Dynamic cellular integration drives functional assembly of the heart's pacemaker complex. *Cell Rep*. 2018;23:2283-2291
76. Nohno T, Kawakami Y, Wada N, Ishikawa T, Ohuchi H, Noji S. Differential expression of the two closely related lim-class homeobox genes lh-2a and lh-2b during limb development. *Biochem Biophys Res Commun*. 1997;238:506-511
77. Smagulova FO, Manuylov NL, Leach LL, Tevosian SG. Gata4/fog2 transcriptional complex regulates lhx9 gene expression in murine heart development. *BMC Dev Biol*. 2008;8:67
78. Tandon P, Wilczewski CM, Williams CE, Conlon FL. The lhx9-integrin pathway is essential for positioning of the proepicardial organ. *Development*. 2016;143:831-840
79. Dupays L, Shang C, Wilson R, Kotecha S, Wood S, Towers N, Mohun T. Sequential binding of meis1 and nkx2-5 on the popdc2 gene: A mechanism for spatiotemporal regulation of enhancers during cardiogenesis. *Cell Rep*. 2015;13:183-195
80. Haenig B, Kispert A. Analysis of tbx18 expression in chick embryos. *Dev Genes Evol*. 2004;214:407-411
81. Lossie AC, Nakamura H, Thomas SE, Justice MJ. Mutation of l7rn3 shows that odz4 is required for mouse gastrulation. *Genetics*. 2005;169:285-299
82. Yanai M, Tatsumi N, Endo F, Yokouchi Y. Analysis of gene expression patterns in the developing chick liver. *Dev Dyn*. 2005;233:1116-1122
83. Barron M, McAllister D, Smith SM, Lough J. Expression of retinol binding protein and transthyretin during early embryogenesis. *Dev Dyn*. 1998;212:413-422
84. Espinoza-Lewis RA, Yu L, He F, Liu H, Tang R, Shi J, Sun X, Martin JF, Wang D, Yang J, Chen Y. Shox2 is essential for the differentiation of cardiac pacemaker cells by repressing nkx2-5. *Dev Biol*. 2009;327:376-385
85. Tiecke E, Bangs F, Blaschke R, Farrell ER, Rappold G, Tickle C. Expression of the short stature homeobox gene shox is restricted by proximal and distal signals in chick limb buds and affects the length of skeletal elements. *Dev Biol*. 2006;298:585-596
86. Burkhard SB, Bakkers J. Spatially resolved rna-sequencing of the embryonic heart identifies a role for wnt/beta-catenin signaling in autonomic control of heart rate. *Elife*. 2018;7
87. High FA, Jain R, Stoller JZ, Antonucci NB, Lu MM, Loomes KM, Kaestner KH, Pear WS, Epstein JA. Murine jagged1/notch signaling in the second heart field orchestrates fgf8 expression and tissue-tissue interactions during outflow tract development. *J Clin Invest*. 2009;119:1986-1996
88. Watanabe Y, Kokubo H, Miyagawa-Tomita S, Endo M, Igarashi K, Aisaki K, Kanno J, Saga Y. Activation of notch1 signaling in cardiogenic mesoderm induces abnormal heart morphogenesis in mouse. *Development*. 2006;133:1625-1634
