## Supplementary Dataset 5 for "A 3D molecular atlas of the chick embryonic heart": chickheart-pointclouds-help.html

Help Information

This interface allows expression patterns to be viewed within the context of
a representative model. The patterns may be viewed as 3D point clouds
in which the size and intensity of points increases with expression level
and/or using arbitrary sections through the model with the patterns shown as
colour washes over the sections.

#### Interface Controls

- left button - rotate view
- scroll wheel - zoom (in and out)
- right button - pan view (side to side, up and down)
- hover (pause with cursor over icon) - icon explanation

The above controls will only function if the interface window has 'focus'.
You can give the window focus by clicking inside it.

Displaying too many point clouds can cause your computer to become slow
or un-responsive.

#### Colours

The colour used to display an object can be changed by clicking on the
button next to the objects name and visibility control.
A colour swatch is then displayed. Selecting a colour in the swatch
will change the object's colour to that selected. The swatch can be
canceled by clicking on the associated  icon.
Colour changes can be undone using the interface undo button.

#### Sectioning

The sectioning controls allow an arbitrary section through the 3D model
to be viewed. This section can also be used to clip the model surface.
The section distance controls the distance of the section from the
centre of the 3D model. The pitch angle may be varied between 0 degrees
(transverse) through 90 degrees (sagittal and coronal) to 180 degrees
(back to transverse).The yaw angle may be varied between 0 and 360 degrees
with 0 being coronal and 90 sagittal (when pitch is 90 degrees).
When expression patterns are selected and a section is viewed the patterns
are shown as colour washes over the section. When compositing the section
there is a precedence with patterns lower in the list mostly being
dominant.

#### Icons

- - undo last change.
- - redo last change.
- - save the current state of the interface to
  your computer.
- - load an interface state from your computer.
- - share your view via a browser url
- - toggle this help text
- - object or group displayed, click to un-display
- - object, section or group un-displayed, click to un-display
  (displaying a group will display all object in the group)
- - model surface not clipped by section, click to clip model
- - model surface clipped by section, click to clip opposite side
- - model surface clipped by section (opposite side),
  click to clip un-clip model
- - all point clouds hidden (click to un-hide)
- - all point clouds not hidden (click to hide)
- - open a group
- - close a group
- - return to the current home view
- - set the home view
- - decrease point cloud point size
- - increase point cloud point size
- - decrease surface opacity
- - increase surface opacity
- - toggle the display of controls
