## Supplementary Dataset 5 for "A 3D molecular atlas of the chick embryonic heart": BasicShader.js

```
/**
 * @author mrdoob / http://www.mrdoob.com
 *
 * Simple test shader
 */

THREE.BasicShader = {

	uniforms: {},

	vertexShader: [

		"void main() {",

			"gl_Position = projectionMatrix * modelViewMatrix * vec4( position, 1.0 );",

		"}"

	].join("\n"),

	fragmentShader: [

		"void main() {",

			"gl_FragColor = vec4( 1.0, 0.0, 0.0, 0.5 );",

		"}"

	].join("\n")

};
```
