## Supplementary Dataset 5 for "A 3D molecular atlas of the chick embryonic heart": BlendShader.js

```
/**
 * @author alteredq / http://alteredqualia.com/
 *
 * Blend two textures
 */

THREE.BlendShader = {

	uniforms: {

		"tDiffuse1": { type: "t", value: null },
		"tDiffuse2": { type: "t", value: null },
		"mixRatio":  { type: "f", value: 0.5 },
		"opacity":   { type: "f", value: 1.0 }

	},

	vertexShader: [

		"varying vec2 vUv;",

		"void main() {",

			"vUv = uv;",
			"gl_Position = projectionMatrix * modelViewMatrix * vec4( position, 1.0 );",

		"}"

	].join("\n"),

	fragmentShader: [

		"uniform float opacity;",
		"uniform float mixRatio;",

		"uniform sampler2D tDiffuse1;",
		"uniform sampler2D tDiffuse2;",

		"varying vec2 vUv;",

		"void main() {",

			"vec4 texel1 = texture2D( tDiffuse1, vUv );",
			"vec4 texel2 = texture2D( tDiffuse2, vUv );",
			"gl_FragColor = opacity * mix( texel1, texel2, mixRatio );",

		"}"

	].join("\n")

};
```
