## Supplementary Dataset 5 for "A 3D molecular atlas of the chick embryonic heart": Detector.js

```
/**
 * @author alteredq / http://alteredqualia.com/
 * @author mr.doob / http://mrdoob.com/
 */

var Detector = {

	canvas: !! window.CanvasRenderingContext2D,
	webgl: ( function () { try { var canvas = document.createElement( 'canvas' ); return !! window.WebGLRenderingContext && ( canvas.getContext( 'webgl' ) || canvas.getContext( 'experimental-webgl' ) ); } catch( e ) { return false; } } )(),
	workers: !! window.Worker,
	fileapi: window.File && window.FileReader && window.FileList && window.Blob,

	getWebGLErrorMessage: function () {

		var element = document.createElement( 'div' );
		element.id = 'webgl-error-message';
		element.style.fontFamily = 'monospace';
		element.style.fontSize = '13px';
		element.style.fontWeight = 'normal';
		element.style.textAlign = 'center';
		element.style.background = '#fff';
		element.style.color = '#000';
		element.style.padding = '1.5em';
		element.style.width = '400px';
		element.style.margin = '5em auto 0';

		if ( ! this.webgl ) {

			element.innerHTML = window.WebGLRenderingContext ? [
				'Your graphics card does not seem to support <a href="http://khronos.org/webgl/wiki/Getting_a_WebGL_Implementation" style="color:#000">WebGL</a>.<br />',
				'Find out how to get it <a href="http://get.webgl.org/" style="color:#000">here</a>.'
			].join( '\n' ) : [
				'Your browser does not seem to support <a href="http://khronos.org/webgl/wiki/Getting_a_WebGL_Implementation" style="color:#000">WebGL</a>.<br/>',
				'Find out how to get it <a href="http://get.webgl.org/" style="color:#000">here</a>.'
			].join( '\n' );

		}

		return element;

	},

	addGetWebGLMessage: function ( parameters ) {

		var parent, id, element;

		parameters = parameters || {};

		parent = parameters.parent !== undefined ? parameters.parent : document.body;
		id = parameters.id !== undefined ? parameters.id : 'oldie';

		element = Detector.getWebGLErrorMessage();
		element.id = id;

		parent.appendChild( element );

	}

};
```
