## Supplementary Dataset 5 for "A 3D molecular atlas of the chick embryonic heart": gxp-pointcloud.js

```
/* == GUI access constants. == */
const obj_ctl_model   = 0
const obj_ctl_section = 1
const obj_ctl_assays  = 2

/* == Globals variables. == */

var prm = null;
var ren = null;
var state = undefined;
var base_url = '';
var config_file = '';
var show_gui = true;
var cancel_img = undefined;
var swatch_img = undefined;
var swatch_canvas = undefined;
var swatch_target = undefined;
var undoStack = [];
var redoStack = [];
var maxSavedStates = 100;

/* == Small utility functions. == */

debug = function(str) {
  if(state.debug) {
    console.log('DEBUG: ' + str);
  }
}

cloneStruct = function(s) {
  return(JSON.parse(JSON.stringify(s)));
}

tripleHexColor = function(v) {
  var c = '';
  for(var idx = 0; idx < 3; ++idx) {
    if(v[idx] < 0x10) {
      c = c + '0';
    }
    c = c + v[idx].toString(16);
  }
  return(c);
}

sensibleFloatStr = function(f) {
  f = Math.round(f * 1000000.0) / 1000000.0;
  var s = '' + f;
  return(s);
}

rgbColor = function(n, a) {
  var r = (n >> 16) & 255;
  var g = (n >> 8) & 255;
  var b = n & 255;
  var rgba = '' + r + ',' + g + ',' + b + ',' + a;
  return(rgba);
}

/* == Small functions for GUI control. == */
getButtonVisible = function(b) {
  var noteye = b.src.match(/noteye\.png$/);
  var eye = ! Boolean(noteye);
  return(eye);
}

getButtonPointclouds = function(b) {
  var notpc = b.src.match(/notpointclouds\.png$/);
  var pc = ! Boolean(notpc);
  return(pc);
}

setButtonVisible = function(b, v) {
  b.src = (v)? 'icons/eye.png': 'icons/noteye.png';
}

setButtonClipping = function(b, c, f) {
  if(c) {
    b.src = (f)? 'icons/sectionclip2.png': 'icons/sectionclip1.png';
  } else {
    b.src = 'icons/notsectionclip.png';
  }
}

setButtonPointclouds = function(b, p) {
  b.src = (p)? 'icons/pointclouds.png': 'icons/notpointclouds.png';
}

setButtonOpen = function(b, o) {
  b.src = (o)? 'icons/minus.png': 'icons/plus.png';
}

setAssayListOpen = function(but, tbl, visible) {
  setButtonOpen(but, visible);
  tbl.style.display = (visible)? 'table': 'none';
}

/* == Functions called from GUI elements of the application. == */

redoAction = function() {
  debug('redoAction()');
  pushState();
  var s = redoStack.pop();
  if(Boolean(s)) {
    setState(s);
  }
  debug('redoAction() undoStack ' + undoStack.length +
        ' redoStack ' + redoStack.length)
}

undoAction = function() {
  debug('undoAction()');
  pushRedoState();
  var s = popState();
  if(Boolean(s)) {
    setState(s);
  }
  debug('undoAction() undoStack ' + undoStack.length +
        ' redoStack ' + redoStack.length)
}

saveStateToFileAction = function() {
  debug('saveStateToFileAction()');
  var jsn = JSON.stringify(state, null, 2);
  var elm = document.createElement('a');
  elm.setAttribute('href',
                   'data:text/plain;charset=utf-8,' + encodeURIComponent(jsn));
  elm.setAttribute('download', 'state.json');
  elm.style.display = 'none';
  document.body.appendChild(elm);
  elm.click();
  document.body.removeChild(elm);
}

loadStateFromFileAction = function() {
  debug('loadStateFromFileAction()');
  pushState();
  var elm = document.createElement('input');
  elm.setAttribute('type', 'file');
  elm.setAttribute('accept', 'application/json');
  elm.onchange = function(evt) {
    debug('loadStateFromFileAction() onchange = true');
    var files = evt.target.files;
    if(Boolean(files) && (files.length === 1)) {
      debug('loadStateFromFileAction() file = ' + files[0].name);
      var rdr = new FileReader();
      rdr.onload = function(evt) {
        var jsn = rdr.result;
	var newState = JSON.parse(jsn);
	setState(newState);
      }
      rdr.onerror = function(evt) {
        debug('loadStateFromFileAction() error');
      }
      rdr.readAsText(files[0]);
    }
    else {
      debug('loadStateFromFileAction() invalid file type');
    }
    document.body.removeChild(elm);
  }
  elm.click();
  document.body.appendChild(elm);
}

shareViewAction = function() {
  debug('shareViewAction()');
  var url = base_url + '?config=' + config_file;
  var idx = state.model_idx;
  url += '&model=' + idx + ',' + state.models[idx].color + ',' +
         ((state.model_visible)? 't': 'f');
  url += '&assays';
  var sep = '=';
  var assays = state.models[state.model_idx].assays;
  for(idx = 0; idx < assays.length; ++idx) {
    var asy = assays[idx];
    url += sep + idx + ',' + asy.color  + ',' + asy.visible;
    sep = ',';
  }
  url += '&home=' + sensibleFloatStr(ren.center.x) + ','
                  + sensibleFloatStr(ren.center.y) + ','
		  + sensibleFloatStr(ren.center.z);
  url += ',' + sensibleFloatStr(ren.nearPlane) + ','
             + sensibleFloatStr(ren.farPlane);
  url += ',' + sensibleFloatStr(ren.camera.position.x) + ','
             + sensibleFloatStr(ren.camera.position.y) + ','
             + sensibleFloatStr(ren.camera.position.z);
  url += ',' + sensibleFloatStr(ren.camera.up.x) + ','
             + sensibleFloatStr(ren.camera.up.y) + ','
             + sensibleFloatStr(ren.camera.up.z);
  if(state.sectioning) {
    var sec = state.models[state.model_idx].section;
    if(sec !== undefined) {
      url += '&section=' +
        sensibleFloatStr(sec.dst) + ',' +
        sensibleFloatStr(sec.pit) + ',' +
        sensibleFloatStr(sec.yaw) + ',' +
        sensibleFloatStr(sec.fxp[0]) + ',' +
        sensibleFloatStr(sec.fxp[1]) + ',' +
        sensibleFloatStr(sec.fxp[2]);
    }
  }
  debug('shareViewAction() url = ' + url);
  window.alert('Share this view using the URL:\n' + url);
}

clippingAction = function(but) {
  debug('clippingAction() ' + but + ' ' + but.id);
  var clip = 0;
  if(but.src.indexOf('notsectionclip') >= 0) {
    clip = 1;
    but.src = 'icons/sectionclip1.png';
  } else if(but.src.indexOf('sectionclip1') >= 0) {
    clip = 2;
    but.src = 'icons/sectionclip2.png';
  } else {
    clip = 0;
    but.src = 'icons/notsectionclip.png';
  }
  if(state.sectioning) {
    var sec = state.models[state.model_idx].section;
    switch(clip)
    {
      case 1:
        sec.clipping = true;
	sec.flip = false;
	break;
      case 2:
        sec.clipping = true;
	sec.flip = true;
	break;
      default:
        sec.clipping = false;
	break;
    }
    updateSection();
  }
}

pointcloudAction = function(but) {
  debug('pointcloudAction() ' + but + ' ' + but.id);
  if(but.src.indexOf('notpointclouds.png') >= 0) {
    but.src = 'icons/pointclouds.png';
  } else {
    but.src = 'icons/notpointclouds.png';
  }
  if(state.sectioning) {
    var sec = state.models[state.model_idx].section;
    sec.pointclouds = ! sec.pointclouds;
    assays = state.models[state.model_idx].assays;
    for(var asy_idx = 0; asy_idx < assays.length; ++asy_idx) {
      setAssay(asy_idx);
    }
  }
}

toggleVisibleAction = function(but) {
  debug('toggleVisibleAction() ' + but + ' ' + but.id);
  var make_visible = !getButtonVisible(but);
  setButtonVisible(but, make_visible);
  if(but.id === 'modelvisible') {
    setModelVisible(make_visible);
  } else if(but.id === 'sectionvisible') {
    setSectionVisible(make_visible);
  } else {
    var mat = but.id.match(/\d+/g);
    var asy_sel_idx = parseInt(mat[0]);
    var asy_idx = parseInt(mat[1]);
    setAssayVisible(asy_idx, make_visible);
  }
}

toggleAssayListVisibleAction = function(but) {
  debug('toggleAssayListVisibleAction() ' + but + ' ' + but.id);
  var make_visible = !getButtonVisible(but);
  setButtonVisible(but, make_visible);
  var mat = but.id.match(/\d+/g);
  var asy_sel_idx = parseInt(mat[0]);
  setAssayListVisible(asy_sel_idx, make_visible);
}

toggleAssayListOpenAction = function(but) {
  debug('toggleAssayListOpenAction() ' + but);
  var idx = parseInt(but.id.match(/\d+$/));
  var tbl = document.getElementById('assaytable' + idx);
  var visible = (Boolean(but.src.match(/plus\.png$/)) === false);
  setAssayListOpen(but, tbl, !visible);
}

toggleGUI = function() {
  debug('toggleGUI()');
  var table_ids = ['statecontrol', 'objcontrol', 'viewcontrol'];
  var elm_ids = ['info'];
  show_gui = !show_gui;
  debug('toggleGUI() ' + show_gui);
  for(var tid of table_ids) {
    var tbl = document.getElementById(tid);
    tbl.style.display = (show_gui)? 'block': 'none';
  }
  for(var eid of elm_ids) {
    var elm = document.getElementById(eid);
    elm.style.display = (show_gui)? 'block': 'none';
  }
}

helpAction = function() {
  debug('helpAction()');
  var ht = document.getElementById('helptext');
  if(ht.style.display == 'none') {
    ht.style.display = 'block';
  } else {
    ht.style.display = 'none';
  }
}

setModelColorAction = function(but) {
  debug('setModelColorActiont() ' + but + ' ' + but.id);
  swatch_target = but.id;
  showSwatchAction(true);
}

setModelAction = function(sel) {
  debug('setModelAction() ' + sel.selectedIndex);
  pushState();
  initModelSelect();
  setModel(sel.selectedIndex, true);
}

setAssayColorAction = function(but) {
  debug('setAssayColorAction() ' + but + ' ' + but.id);
  swatch_target = but.id;
  showSwatchAction(true);
}

setAssayAction = function(sel) {
  debug('setAssayAction() ' + sel.selectedIndex);
  pushState();
  var x = sel.id.replace(/\D+/g, '');
  var sel_idx = parseInt(sel.id.replace(/\D+/g, ''));
  var n_asy = state.models[state.model_idx].assays.length;
  if((sel_idx >= 0) && (sel_idx < n_asy)) {
    pushState();
    setAssay(sel.selectedIndex);
  }
}

goHomeViewAction = function() {
  debug('goHomeViewAction()');
  pushState();
  goHomeView();
}

setHomeViewAction = function() {
  debug('setHomeViewAction()');
  pushState();
  ren.setHome();
}

viewDotAction = function(inc) {
  debug('viewDotAction() ' + inc);
  pushState();
  ren.pointSizeIncrement(inc);
}

viewSurfAction = function(inc) {
  debug('viewSurfAction() ' + inc);
  pushState();
  ren.opacityIncrement(inc);
}

swatchDoneAction = function(img, ev) {
  debug('swatchDoneAction() ' + img + ' ' + ev);
  if(Boolean(swatch_target) && (img.id == 'swatch')) {
    pushState();
    var p = [ev.layerX, ev.layerY];
    debug('swatchDoneAction() ' + p[0] + ' ' + p[1]);
    var v = swatch_canvas.getContext('2d').getImageData(p[0], p[1], 1, 1,).data;
    debug('swatchDoneAction() ' + v[0] + ' ' + v[1] + ' ' + v[2]);
    var what = swatch_target.replace(/\d+.*$/, '');
    var color = tripleHexColor(v);
    if(what === 'modelbutton') {
      var mod = state.models[state.model_idx];
      mod.color = color;
      updateModel(mod);
    } else if(what == 'assaybutton') {
      var asy_idx;
      var asy_sel_idx;
      var mat = swatch_target.match(/\d+/g);
      var asy_sel_idx = parseInt(mat[0]);
      asy_idx = parseInt(mat[1]);
      state.models[state.model_idx].assays[asy_idx].color = color;
      updateAssay(asy_idx);
    }
  }
  swatch_target = undefined;
  showSwatchAction(false);
}

showSwatchAction = function(show) {
  debug('showSwatchAction() ' + show);
  if(Boolean(show)) {
    document.body.appendChild(swatch_img);
    document.body.appendChild(cancel_img);
  } else {
    swatch.parentNode.removeChild(swatch_img);
    cancel_img.parentNode.removeChild(cancel_img);
  }
}

/* == Rendering functions. == */

goHomeView = function() {
  ren.setCamera();
  ren.goHome();
}

clearRenModels = function() {
  ren.removeModel('reference');
  ren.removeModel('section');
  asys = state.models[state.model_idx].assays
  for(var idx = 0; idx < asys.length; ++idx) {
    var ren_obj_name = 'assay' + idx;
    ren.removeModel(ren_obj_name);
  }
}

/* == State management functions. == */

pushState = function() {
  debug('pushState()');
  if(undoStack.length >= maxSavedStates) {
    undoStack.length = maxSavedStates - 1;
  }
  undoStack.push(cloneStruct(state));
}

popState = function() {
  debug('popState()');
  return(undoStack.pop());
}

pushRedoState = function() {
  debug('pushRedoState()');
  if(redoStack.length >= maxSavedStates) {
    redoStack.length = maxSavedStates - 1;
  }
  redoStack.push(cloneStruct(state));
}

popRedoState = function() {
  debug('popRedoState()');
  return(redoStack.pop());
}

setState = function(newState) {
  debug('setState()');
  state = cloneStruct(newState);
  window.document.title = state.title;
  document.getElementById('info').innerHTML = state.info;
  if(state.sectioning === undefined) {
    state.sectioning = false;
  }
  initModelSelect();
  ren.pointSizeSet(state.point_size);
  setModel(state.model_idx, true);
  for(var asy_idx = 0; asy_idx < state.assay_sel.length; ++asy_idx) {
    setAssay(asy_idx);
  }
} 

/* == Functions which initialise the interface and state. == */

initHelpText = function() {
  debug('initHelpText()');
  var ht = document.getElementById('helptext');
  if(Boolean(state.help_file)) {
    var xhr= new XMLHttpRequest();
    xhr.open('GET', state.help_file, true);
    xhr.onreadystatechange= function() {
	if ((this.readyState === 4) && (this.status === 200)) {
          ht.innerHTML  = '<div class=margin5>' + 
              this.responseText + '</div>' +
	      '<p></p><p></p>' +
	      '<div class="aligncentre">' +
	      '<button type="button" onclick="helpAction()">' +
	      'Close</button></div><p></p>'
        }
    };
    xhr.send();
  } else {
    ht.innerHTML = '<div class="aligncentre"><u>' +
        'Help Information</u></div><p></p>'
    ht.innerHTML += '<p><div class=margin5>' +
        'No application specific help has been provided, for ' +
        'generic help using this interface please visit: ' +
	'<br>  <a href="https://github.com/ma-tech">ma-tech</a></div></p>'
    ht.innerHTML += '<p></p><p></p>' +
		    '<div class="aligncentre">' +
		    '<button type="button" onclick="helpAction()">' +
		    'Close</button></div><p></p>'
  }
  ht.style.display = 'none';
  debug('initHelpText() text = ' + ht.innerHTML)
}

initColorSwatch = function() {
  debug('initColorSwatch()');
  swatch_canvas = document.createElement('canvas');
  swatch_img = document.createElement('img');
  swatch_img.id = 'swatch';
  swatch_img.onload = function() {
    debug('initColorSwatch() swatch loaded');
    swatch_canvas.width = swatch_img.width;
    swatch_canvas.height = swatch_img.height;
    swatch_canvas.getContext('2d').drawImage(swatch_img, 0, 0);
    swatch_img.setAttribute('onclick', 'swatchDoneAction(this, event)');
  }
  cancel_img = document.createElement('img');
  cancel_img.id = 'cancelswatch';
  cancel_img.onload = function() {
    debug('initColorSwatch() cancel loaded');
    cancel_img.setAttribute('onclick', 'swatchDoneAction(this, event)');
  }
  swatch_img.src = 'icons/swatch.png';
  cancel_img.src = 'icons/cancel.png';
}

initModelSelect = function() {
  debug('initModelSelect()');
  var tbl = document.getElementById('objcontrol');
  while(tbl.hasChildNodes())
  {
    tbl.removeChild(tbl.firstChild);
  }
  var tr;
  var td;
  tr = tbl.insertRow(obj_ctl_model);
  td = tr.insertCell(0);
  tbl = document.createElement('table');
  td.appendChild(tbl);
  tr = tbl.insertRow(0);
  td = tr.insertCell(0);
  td.innerHTML = state.model_sel_txt;
  td = tr.insertCell(1);
  td.setAttribute('width', '100%');
  td.setAttribute('align', 'rigth');
  tbl = document.createElement('table');
  td.appendChild(tbl);
  tr = tbl.insertRow(0);
  td = tr.insertCell(0);
  td.setAttribute('width', '33%');
  var but = document.createElement('img');
  but.align = 'middle';
  but.setAttribute('id', 'modelvisible');
  but.title='Toggle object visability';
  but.setAttribute('onclick', 'toggleVisibleAction(this)');
  setButtonVisible(but, true);
  td.appendChild(but);
  td = tr.insertCell(1);
  td.setAttribute('width', '33%');
  but = document.createElement('button');
  but.align = 'middle';
  but.setAttribute('id', 'modelbutton');
  but.setAttribute('class', 'smbutton');
  but.title='Select reference model colour';
  but.setAttribute('onclick', 'setModelColorAction(this)');
  td.appendChild(but);
  td = tr.insertCell(2);
  td.setAttribute('width', '33%');
  var sel = document.createElement('select');
  sel.setAttribute('id', 'modelselect');
  sel.title='Select stage';
  td.appendChild(sel);
  for(var mod_idx = 0; mod_idx < state.models.length; ++mod_idx) {
    var mod = state.models[mod_idx];        
    var opt = document.createElement('option');
    opt.text = mod.name;
    opt.value = mod_idx;
    sel.add(opt);
  }
  sel.onchange = function(){setModelAction(sel)};
}

setSectionAction = function(ip, prm) {
  switch(ip.id) {
    case 'sec_' + prm + '_number':
      var op = document.getElementById('sec_' + prm + '_range');
      op.value = ip.value;
      break;
    case 'sec_' + prm + '_range':
      var op = document.getElementById('sec_' + prm + '_number');
      op.value = ip.value;
      break;
  }
  if(state.sectioning) {
    var sec = state.models[state.model_idx].section;
    if(prm === 'dst') {
      sec.dst = ip.value;
    } else if(prm == 'pit') {
      sec.pit = ip.value;
    } else if(prm == 'yaw') {
      sec.yaw = ip.value;
    }
    updateSection();
  }
}

initSectionValue = function(tr, label, prm, min, max, val) {
  var td = tr.insertCell(0);
  td.innerHTML = label;
  var rng = document.createElement('input');
  rng.setAttribute('type', 'range');
  rng.id = 'sec_' + prm + '_range';
  var hint = 'Set section ' + label.toLowerCase();
  rng.title = hint;
  rng.oninput = function(){setSectionAction(rng, prm)};
  rng.onmouseup = function(){setSectionAction(rng, prm)};
  td = tr.insertCell(1);
  td.appendChild(rng);
  var num = document.createElement('input');
  num.setAttribute('type', 'number');
  num.id = 'sec_' + prm + '_number';
  num.title = hint;
  num.onchange = function(){setSectionAction(num, prm)};
  td = tr.insertCell(2);
  td.appendChild(num);
  setSectionValue(prm, min, max, val);
}

setSectionValue = function(prm, min, max, val) {
  var rng = document.getElementById('sec_' + prm + '_range');
  var num = document.getElementById('sec_' + prm + '_number');
  rng.min = num.min = min;
  rng.max = num.max = max;
  var ln = min.toString().length;
  var lx = max.toString().length;
  if(lx > ln) {
    ln = lx;
  }
  if(6 > ln) {
    ln = 6;
  }
  num.style.width = ln.toString() + 'em';
  rng.value = num.value = val;
}

baseURL = function() {
  var sec = state.models[state.model_idx].section;        
  var url = null;
  if(Boolean(sec.iip3dsrv) && Boolean(sec.wlzobj)) {
    var url = sec.iip3dsrv + '?' +
	      'wlz=' + sec.wlzobj + '&' +
	      'mod=zeta&' +
	      'fxp=' + sec.fxp[0] + ',' + sec.fxp[1] + ',' + sec.fxp[2] + '&' +
	      'pit=' + sec.pit + '&' +
	      'yaw=' + sec.yaw + '&' +
	      'dst=' + sec.dst;
  }
  return(url);
}

initSectioning = function() {
  debug('initSectioning()');
  var objtbl = document.getElementById('objcontrol');
  var sec_row = objtbl.insertRow(obj_ctl_section);
  // sec_row.style.backgroundColor = '#ffff00'; // HACK TODO debug
  if((state.sectioning === undefined) || (state.sectioning === false)) {
    sec_row.style.display = 'none';
  } else {
    // Get section distance range from IIP3D server
    var dst_min = -1000;
    var dst_max = 1000;
    var url = baseURL();
    var mod = state.models[state.model_idx];
    var sec = mod.section;
    if(Boolean(url)) {
      var req = new XMLHttpRequest();
      req.open('GET', url + '&OBJ=Wlz-distance-range', false);
      req.send(null);
      if(req.status === 200) {
	var rsp = req.responseText.split(':')[1].split(' ');
	dst_min = Number(rsp[0]);
	dst_max = Number(rsp[1]);
      }
    }
    // Create GUI control elements
    var tr;
    td = sec_row.insertCell(0);
    var sec_tbl = document.createElement('table');
    sec_tbl.setAttribute('width', '100%');
    td.appendChild(sec_tbl);
    tr = sec_tbl.insertRow(0);
    td = tr.insertCell(0);
    td.innerHTML = 'Section Controls';
    td = tr.insertCell(1);
    var but_tbl = document.createElement('table');
    but_tbl.setAttribute('width', '100%');
    td.appendChild(but_tbl);
    tr = but_tbl.insertRow(0);
    td = tr.insertCell(0);
    td.setAttribute('width', '33%');
    var but = document.createElement('img');
    but.align = 'middle';
    but.setAttribute('id', 'sectionvisible');
    but.title='Toggle section visability';
    but.setAttribute('onclick', 'toggleVisibleAction(this)');
    setButtonVisible(but, sec.visible);
    td.appendChild(but);
    td = tr.insertCell(1);
    td.setAttribute('width', '33%');
    var but = document.createElement('img');
    but.align = 'middle';
    but.setAttribute('id', 'sectionclip');
    but.title='Toggle section clipping';
    but.setAttribute('onclick', 'clippingAction(this)');
    setButtonClipping(but, sec.clipping, sec.flip);
    td.appendChild(but);
    td = tr.insertCell(2);
    td.setAttribute('width', '33%');
    var but = document.createElement('img');
    but.align = 'middle';
    but.setAttribute('id', 'sectionpointclouds');
    but.title='Toggle pointcloud visability';
    but.setAttribute('onclick', 'pointcloudAction(this)');
    setButtonPointclouds(but, sec.pointclouds);
    td.appendChild(but);
    tr = sec_tbl.insertRow(1);
    initSectionValue(tr, 'Distance', 'dst', dst_min, dst_max, sec.dst);
    tr = sec_tbl.insertRow(2);
    initSectionValue(tr, 'Pitch', 'pit', 0, 180, sec.pit);
    tr = sec_tbl.insertRow(3);
    initSectionValue(tr, 'Yaw', 'yaw', 0, 360, sec.yaw);
    sec_row.style.display = 'block';
    // First render of section object
    var vtx = ren.getIIP3DBBVertices(url,
	new THREE.Vector3(sec.voxelsz[0], sec.voxelsz[1],
			  sec.voxelsz[2]));
    var tex = setSectionTexture();
    ren.addModel({name:        'section',
		  mode:        MARenderMode.SECTION,
		  transparent: true,
		  opacity:     0.5,
		  visible:     sec.visible,
		  vertices:    vtx,
		  texture:     tex});
  }
}

initAssaySelect = function() {
  debug('initAssaySelect()');
  var tbl = document.getElementById('objcontrol');
  var assays = state.models[state.model_idx].assays;
  var tr = tbl.insertRow(obj_ctl_assays);
  var td = tr.insertCell(0);
  tbl = document.createElement('table');
  td.appendChild(tbl);
  for(var asy_sel_idx = 0; asy_sel_idx < state.assay_sel.length;
      ++asy_sel_idx) {
    var sel_id = 'assay' + asy_sel_idx;
    var sel = document.getElementById(sel_id);
    var asytbl = undefined;
    if(Boolean(sel) === false)
    {
      var tr = tbl.insertRow(asy_sel_idx);
      tr.setAttribute('id', 'catagory' + asy_sel_idx);
      var td = tr.insertCell(0);
      td.innerHTML = state.assay_sel[asy_sel_idx].name;
      var td = tr.insertCell(1);
      var but = document.createElement('img');
      but.align = 'middle';
      but.setAttribute('id', 'assaylistvisiblebutton' + asy_sel_idx);
      but.title ='Assays visible';
      setButtonVisible(but, false);
      but.setAttribute('onclick', 'toggleAssayListVisibleAction(this)');
      td.appendChild(but);
      td = tr.insertCell(2);
      but = document.createElement('img');
      but.align = 'middle';
      but.setAttribute('id', 'assaylistbutton' + asy_sel_idx);
      but.title ='Toggle assay list';
      but.setAttribute('onclick', 'toggleAssayListOpenAction(this)');
      td.appendChild(but);
      td = tr.insertCell(3);
      td.colSpan = 2;
      var asy_sel = state.assay_sel[asy_sel_idx];
      setButtonOpen(but, asy_sel.visible);
      asytbl = document.createElement('table');
      asytbl.setAttribute('id', 'assaytable' + asy_sel_idx);
      asytbl.style.display = (asy_sel.visible)? 'table': 'none';
      asytbl.style.padding = '0px';
      td.appendChild(asytbl);
      td.style.borderStyle = 'none none none solid';
    }
    for(var idx = 0; idx < state.assay_sel.length; ++idx) {
      var asy_sel = state.assay_sel[idx];
      if(asy_sel.visible === undefined) {
        asy_sel['visible'] = false;
      }
    }
    for(var idx = 0; idx < assays.length; ++idx) {
      var asy = assays[idx];
      if(asy.visible === undefined) {
        asy['visible'] = false;
      }
    }
    var lstidx = 0;
    for(var idx = 0; idx < assays.length; ++idx) {
      var asy = assays[idx];
      if(asy.cat === undefined) {
	asy.cat = [];
      }
      if(asy.cat.indexOf(asy_sel_idx) >= 0) {
	var tr = asytbl.insertRow(lstidx);
	var td = tr.insertCell(0);
	td.innerHTML = asy.name;
	td = tr.insertCell(1);
	td.style.padding = '0px';
	var but = document.createElement('img');
	but.title='Toggle object visability';
	but.align = 'middle';
	but.setAttribute('id', 'assayvisible' + asy_sel_idx + '-' + idx);
	but.setAttribute('onclick', 'toggleVisibleAction(this)');
	setButtonVisible(but, false);
	td.appendChild(but);
	td.style.padding = '0px';
	but = document.createElement('button');
	but.align = 'middle';
	but.setAttribute('id', 'assaybutton' + asy_sel_idx + '-' + idx);
	but.setAttribute('class', 'smbutton');
	but.title='Select object colour';
	but.setAttribute('onclick', 'setAssayColorAction(this)');
	td.appendChild(but);
	td.style.padding = '0px';
	++lstidx;
      }
    }
  }
  for(var asy_idx = 0; asy_idx < assays.length; ++asy_idx) {
    setAssay(asy_idx);
    updateAssayListsVisible(asy_idx);
  }
}

/* == Model management functions. == */

setModel = function(new_model_idx, force) {
  debug('setModel() ' + new_model_idx + ' ' + force);
  var new_model = state.models[new_model_idx].name;
  if(force || (new_model_idx !== state.model_idx)) {
    state.model_idx  = new_model_idx;
    var mod = state.models[state.model_idx];
    for(var asy_sel_idx = 0; asy_sel_idx < state.assay_sel.length;
        ++asy_sel_idx) {
      var asy_sel = state.assay_sel[asy_sel_idx];
      if(asy_sel.sel_idx == undefined) {
        asy_sel['sel_idx'] = 0;
      }
    }
    state.model_init = true;
    checkModelSection();
    clearRenModels();
    if(mod.name !== 'TS19') {
      var cam = mod.camera_home;
      ren.setCamera(new THREE.Vector3(cam.cen[0], cam.cen[1], cam.cen[2]),
      		    cam.near, cam.far,
		    new THREE.Vector3(cam.pos[0], cam.pos[1], cam.pos[2]));
      ren.setHome(new THREE.Vector3(cam.pos[0], cam.pos[1], cam.pos[2]),
		  new THREE.Vector3(cam.up[0], cam.up[1], cam.up[2]));
    }
    var path = state.model_obj_dir + '/' + mod.reference_obj;
    var color = '0x' + mod.color;
    var vis = state.model_visible;
    ren.addModel({name:        'reference',
		  mode:        MARenderMode.PHONG,
		  path:        path,
		  color:       parseInt(color),
		  visible:     vis,
		  transparent: true,
		  clipping:    null,
		  opacity:     mod.opacity});
    var select = document.getElementById('modelselect');
    select.value = new_model_idx;
    initSectioning();
    updateModel(mod);
    initAssaySelect();
    goHomeView();
  }
}

checkModelSection = function() {
  if(state.sectioning) {
    var mod = state.models[state.model_idx];
    if(mod.section === undefined) {
      mod.section = [];
    }
    var sec = mod.section;
    for(var p in ['dst', 'pit', 'yaw']) {
      if(sec[p] === undefined) {
        sec[p] = 0.0;
      }
    } 
    for(var p in ['flip', 'visible']) {
      if(sec[p] === undefined) {
	sec[p] = false;
      }
    }
    if(sec.fxp === undefined) {
      sec.fxp = [0.0, 0.0, 0.0];
    }
    if(sec.voxelsz === undefined) {
      sec.voxelsz = [1.0, 1.0, 1.0];
    }
    if(sec.iip3dsrv === undefined) {
      sec.iip3dsrv = null;
    }
    if(sec.wlzobj === undefined) {
      sec.wlzobj = null;
    }
  }
}

updateModel = function(mod) {
  debug('updateModel() ' + mod);
  var but = document.getElementById('modelbutton');
  but.style.background = '#' + mod.color;
  var color = '0x' + mod.color;
  var vis = state.model_visible;
  ren.updateModel({name: 'reference',
		   color: parseInt(color),
		   visible: vis,
		   transparent: true,
		   opacity:     mod.opacity});
  updateSection();
}

updateSection = function() {
  debug('updateSection()');
  if(state.sectioning) {
    var mod = state.models[state.model_idx];
    var sec = mod.section;
    var vis = sec.visible;
    if(vis) {
      var url = baseURL();
      var vtx = ren.getIIP3DBBVertices(url,
	  new THREE.Vector3(sec.voxelsz[0], sec.voxelsz[1],
			    sec.voxelsz[2]));
      var tex = setSectionTexture();
      var pln = ren.makePlaneFromVertices(vtx);
      if(sec.flip) {
        pln.normal = pln.normal.negate();
        pln.constant = -pln.constant;
      }
      ren.updateModel({name:     'section',
		    transparent: true,
		    visible:     sec.visible,
		    opacity:     0.5,
		    vertices:    vtx,
		    texture:     tex});
      if(sec.clipping) {
	ren.updateModel({name: 'reference',
			 clipping:    pln,
			 transparent: true});
      } else {
	ren.updateModel({name: 'reference',
			 clipping:    null,
			 transparent: true});
      }
    } else {
      ren.updateModel({name: 'section',
		       visible: false,
		       transparent: true});
      ren.updateModel({name: 'reference',
                      clipping: null,
		      transparent: true});
    }
  }
}

setSectionTexture = function() {
  var mod = state.models[state.model_idx];
  var sec = mod.section;
  var tex = baseURL();
  var assays = mod.assays;
  // Add model
  tex = tex + '&qlt=50&sel=0,' + rgbColor(parseInt(mod.color, 16), 255) + '&';
  // Add assays
  for(var asy_idx = 0; asy_idx < assays.length; ++asy_idx) {
    var asy = assays[asy_idx];
    if(asy.visible && (asy.iip3d_idx !== undefined)) {
      tex = tex + 'sel=' + asy.iip3d_idx + ',' +
            rgbColor(parseInt(asy.color, 16), 192) + '&';
    }
  }
  tex = tex + 'cvt=png';
  return(tex);
}

setSectionVisible = function(make_visible) {
  if(state.sectioning) {
    var mod = state.models[state.model_idx];
    mod.section['visible'] = make_visible;
    updateSection();
  }
}

setModelVisible = function(make_visible) {
  debug('setModelVisible() ' + make_visible);
  state.model_visible = make_visible;
  var mod = state.models[state.model_idx];
  updateModel(mod);
}

/* == Assay management functions. == */

updateAssayListsVisible = function(gvn_asy_idx) {
  debug('updateAssayListsVisible() ' + gvn_asy_idx);
  var assays = state.models[state.model_idx].assays;
  var cat = assays[gvn_asy_idx].cat;
  // For each assay selection list that the assay is in
  for(var idx = 0; idx < cat.length; ++idx) {
    var visible = false;
    var asy_sel_idx = cat[idx];
    for(var asy_idx = 0; asy_idx < assays.length; ++asy_idx) {
      var asy = assays[asy_idx];
      if(asy.cat.indexOf(asy_sel_idx) >= 0) {
        visible = asy.visible;
	if(visible) {
	  break;
	}
      }
    }
    var but = document.getElementById('assaylistvisiblebutton' + asy_sel_idx);
    setButtonVisible(but, visible);
  }
}

/* \fn setAssayListVisible
 * Sets the visibility of all the assays of the assay selection list.
 * \param asy_sel_idx       The index of the assay selection list.
 * \param v		    True if assay to be set visible.
 */
setAssayListVisible = function(asy_sel_idx, v) {
  debug('setAssayListVisible() ' + asy_sel_idx + ' ' + v);
  assays = state.models[state.model_idx].assays;
  for(var asy_idx = 0; asy_idx < assays.length; ++asy_idx) {
    var asy = assays[asy_idx];
    if(asy.cat.indexOf(asy_sel_idx) >= 0) {
      setAssayVisible(asy_idx, v);
    }
  }
}

/* \fn setAssayVisible
 * Sets the visibility of the assay with the given assay index.
 * \param asy_idx           The assay to be set.
 * \param v		    True if assay to be set visible.
 */
setAssayVisible = function(asy_idx, v) {
  debug('setAssayVisible() ' + asy_idx + ' ' + v);
  state.models[state.model_idx].assays[asy_idx].visible = v;
  setAssay(asy_idx);
  updateAssayListsVisible(asy_idx);
}

/* \fn setAssay
 * Sets the assay in the state and then updates the GUI.
 * \param new_asy_idx           The assay to be set.
 */
setAssay = function(new_asy_idx) {
  debug('setAssay() ' + new_asy_idx);
  var mod = state.models[state.model_idx];
  var asy = mod.assays[new_asy_idx];
  var new_asy_obj = asy.obj;
  var new_asy_name = asy.name;
  var ren_obj_name = 'assay' + new_asy_idx;
  var vis = asy.visible;
  if(state.sectioning) {
    var sec = mod.section;
    vis = vis && sec.pointclouds;
  }
  ren.removeModel(ren_obj_name);
  if((new_asy_obj !== 'null') && vis) {
    var color = '0x' + asy.color;
    var path = state.assay_obj_dir + '/' + new_asy_obj;
    ren.addModel({name: ren_obj_name,
		  path: path,
		  color: parseInt(color),
		  visible: vis,
		  transparent: true,
		  opacity: 0.1,
		  mode: MARenderMode.POINT});
  }
  updateAssay(new_asy_idx);
}

/* \fn updateAssay
 * \param asy_idx               The assay to be updated.
 * Updates the valid assay GUI components and assay model from the state.
 */
updateAssay = function(asy_idx) {
  debug('updateAssay() ' + asy_idx);
  var but = false;
  var col = false;
  var vis = false;
  var mod = state.models[state.model_idx];
  var asy = mod.assays[asy_idx];
  // Assay may be in multiple catagories so run through them
  col = asy.color;
  vis = asy.visible;
  but = false;
  for(var asy_cat_idx = 0; asy_cat_idx < asy.cat.length; ++asy_cat_idx) {
    var asy_sel_idx = asy.cat[asy_cat_idx];
    var at = document.getElementById('assaytable' + asy_sel_idx);
    if(Boolean(at)) {
      var tr_list = at.rows;
      if(Boolean(tr_list)) {
	for(var tr_idx = 0; tr_idx < tr_list.length; ++tr_idx) {
	  var td_cells = tr_list[tr_idx].cells;
	  if(td_cells[0].innerText === asy.name) {
	    but = td_cells[1].children[0];
	    if(Boolean(but)) {
	      setButtonVisible(but, vis);
	    }
	    but = td_cells[1].children[1];
	    if(Boolean(but)) {
	      but.style.background = '#' + col;
	    }
	  }
	}
      }
    }
  }
  if(col) {
    var color = '0x' + col;
    var ren_obj_name = 'assay' + asy_idx;
    ren.updateModel({name: ren_obj_name,
		     color: parseInt(color),
		     visible: vis,
		     transparent: true,
		     opacity: 0.1});
  }
  updateSection();
}

/* == Main application functions. == */

/*!
 * \fn parseURL
 * Parses the query string of the URL for paramters
 * (see https://en.wikipedia.org/wiki/Query_string).
 * On return the global parameter map prm is set with key/value pairs.
 */
parseURL = function() {
  prm = {};
  var href = new String(location.href);
  var seg = href.split('?');
  base_url = seg[0];
  if(seg.length === 2) {
    seg = seg[1].split('&');
    var len = seg.length;
    for(var i = 0; i < len; ++i) {
      var s = seg[i].split('=');
      prm[s[0]] = s[1];
    }
  }
}

/*! \fn updateStateFromParam
 * Parse given parameters and use their to update the state.
 * See shareViewAction().
 */
updateStateFromParam = function() {
  var err = null;
  if(prm['debug']) {
    /* debug = [tTfF]
     */
    state.debug = prm['debug'].match(/^[tT]/);
  }
  if(!err && prm['model']) {
    /* model   = <mod_spec>
     * mod_spec = <idx>,<colour>,<visible>
     * idx      = integer
     * colour   = 6 digit hex
     * visible  = [tTfF]
     */
    var md = prm['model'].split(',');
    if(md.length !== 3) {
      err = 'model';
    } else {
      var mi = md[0];
      var mc = md[1];
      var mv = md[2];
      mi = parseInt(mi);
      mc = mc.toLowerCase().replace(/[^0-9a-f]+/g, '');
      mv = mv.match(/^[tT]/);
      if((isNaN(mi) !== false) || (mi < 0) || (mi >= state.models.length) ||
         (mc.length !== 6)) {
        err = 'model';
      } else {
        state.model_idx = mi;
	state.models[mi].color = mc;
	state.models[mi].visible = mv;
      }
    }
  }
  if(!err && prm['assays']) {
    /* assays   = <asy_spec>(,<asy_spec>)*
     * asy_spec = <idx>,<colour>,<visible>
     * idx      = integer
     * colour   = 6 digit hex
     * visible  = [tTfF]
     */
    var assays = state.models[state.model_idx].assays;
    var ga = prm['assays'].split(',');
    var ln = ga.length / 3;
    for(var idx = 0; (idx < ln); ++idx) {
      var i3 = idx * 3;
      var ai = ga[i3 + 0];
      var ac = ga[i3 + 1];
      var av = ga[i3 + 2];
      ai = parseInt(ai);
      ac = ac.toLowerCase().replace(/[^0-9a-f]+/g, '');
      av = av.match(/^[tT]/);
      if((isNaN(ai) !== false) || (ac.length !== 6)) {
	err = 'assays';
        break;
      } else if((ai > 0) && (ai < assays.length)) {
        var asy = assays[ai];
	asy.color = ac;
	asy.visible = av;
      }
    }
  }
  if(!err && prm['home']) {
    /* home    = <cen>,<near>,<far>,<cam_pos><cam_up>
     * cen     = float,float,float
     * near    = float
     * far     = float
     * cam_pos = float,float,float
     * cam_up  = float,float,float
     */
    var gh = prm['home'].split(',');
    if(gh.length !== 11) {
      err = 'home';
    } else {
      for(var idx = 0; idx < gh.length; ++idx) {
        gh[idx] = parseFloat(gh[idx]);
	if(isNaN(gh[idx])) {
	  err = 'home';
	  break;
	}
      }
    }
    if(!err) {
      var ch = state.models[state.model_idx].camera_home;
      ch.cen[0] = gh[ 0];
      ch.cen[1] = gh[ 1];
      ch.cen[2] = gh[ 2];
      ch.near   = gh[ 3];
      ch.far    = gh[ 4];
      ch.pos[0] = gh[ 5];
      ch.pos[1] = gh[ 6];
      ch.pos[2] = gh[ 7];
      ch.up[0]  = gh[ 8];
      ch.up[1]  = gh[ 9];
      ch.up[2]  = gh[10];
    }
  }
  if(!err) {
    if(prm['sectioning']) {
      state['sectioning'] = (prm['sectioning'] === 'true');
    } else {
      state['sectioning'] = Boolean(state['sectioning']);
    }
  }
  if(!err && prm['section']) {
    /* section = <dst>,<pit>,<yaw>,<fxp>
     * dst     = float
     * pit     = float
     * yaw     = float
     * fxp     = float,float,float
     */
    var gs = prm['section'].split(',');
    if(gs.length !== 6) {
      err = 'section';
    }
    if(!err && state.sectioning) {
      var sec = state.models[state.model_idx].section;
      if((sec !== undefined) &&
         (sec.dst !== undefined) && (sec.pit !== undefined) &&
         (sec.yaw !== undefined) && (sec.fxp !== undefined)) {
	sec.dst = gs[0]
	sec.pit = gs[1]
	sec.yaw = gs[2]
	sec.fxp[0] = gs[3];
	sec.fxp[1] = gs[4];
	sec.fxp[2] = gs[5];
      }
    }
  }
  if(err) {
    alert('Failed to parse url for ' + err);
  }
}

/*! \fn main
 * Given a valid state object already read when parsing the URL query string
 * this function completes the application. */
main = function() {
  var ren_container = document.getElementById('three');
  window.document.title = state.title;
  document.body.appendChild(ren_container);
  document.getElementById('info').innerHTML = state.info;
  initHelpText();
  initColorSwatch();
  initModelSelect();
  ren = new MARenderer(window, ren_container);
  ren.init();
  if(Boolean(prm['background'])) {
    var c = parseInt(prm['background']);
    if(!isNaN(c)) {
      ren.renderer.setClearColor(c, 1 );
    }
  }
  ren.setLocalClipping(true);
  ren.pointSizeSet(state.point_size);
  ren.win.removeEventListener('keypress', ren._keyPressed);
  setModel(state.model_idx, true);
  ren.setHomeOnLoad = false;
  ren.animate();
}

/* Check for WebGL and if there is none issue a message.
 * Then provided WebGL is supported parse the URL, create a state object
 * from the JSON config file, set parameters in the state from URL and
 * call main(). */
if((Detector.webgl)) {
  parseURL();
} else {
  alert('Your web browser does not appear to support WebGL, ' +
        'which is essential to this application. ' +
        'Please see: ' +
	'https://get.webgl.org or ' + 
	'https://en.wikipedia.org/wiki/WebGL#Support');
}
if(prm['config'] === undefined) {
  alert('No parameters provided,' +
        'as a minimum a configuration file is required.');
} else {
  config_file = prm['config'];
  var req = new XMLHttpRequest();
  req.open('GET', prm['config'], false);
  req.send(null);
  if(req.status === 200) {
    state = JSON.parse(req.responseText);
    if(Boolean(state)) {
      updateStateFromParam();
      main();
    } else {
      alert('Invalid configuration file provided.');
    }
  }
}
```
