## Supplementary Dataset 5 for "A 3D molecular atlas of the chick embryonic heart": MARender.js

```
/*!
* \file         MARender.js
* \author       Bill Hill
* \date         June 2015
* \version      $Id$
* \par
* Address:
*               MRC Human Genetics Unit,
*               MRC Institute of Genetics and Molecular Medicine,
*               University of Edinburgh,
*               Western General Hospital,
*               Edinburgh, EH4 2XU, UK.
* \par
* Copyright (C), [2015],
* The University Court of the University of Edinburgh,
* Old College, Edinburgh, UK.
* 
* This program is free software; you can redistribute it and/or
* modify it under the terms of the GNU General Public License
* as published by the Free Software Foundation; either version 2
* of the License, or (at your option) any later version.
*
* This program is distributed in the hope that it will be
* useful but WITHOUT ANY WARRANTY; without even the implied
* warranty of MERCHANTABILITY or FITNESS FOR A PARTICULAR
* PURPOSE.  See the GNU General Public License for more
* details.
*
* You should have received a copy of the GNU General Public
* License along with this program; if not, write to the Free
* Software Foundation, Inc., 51 Franklin Street, Fifth Floor,
* Boston, MA  02110-1301, USA.
* \brief	A simple renderer based on three.js which is aimed
* 		at displaying anatomy and gene expression domains in 3D.
*/

/**
* Rendering modes
*/
MARenderMode = {
  BASIC:                0,
  WIREFRAME:            1,
  LAMBERT:              2,
  PHONG:                3,
  EMISSIVE:             4,
  POINT:		5,
  SECTION:		6
}

/**
* Properties which may be set when adding or updating a model.
*/
MARenderItem = function() {
  this.name             = undefined;
  this.path             = undefined;
  this.color            = 0xffffff;
  this.side		= THREE.FrontSide; /* Double sided can cause artifacts
                                              so only use if needed. */
  this.transparent      = false;
  this.opacity          = 1.0;
  this.mode             = MARenderMode.PHONG;
  this.vertices		= undefined;
  this.texture		= undefined;
  this.visible          = true;
  this.clipplane        = undefined;
}

/**
* Enough of a camera state to restore position.
*/
MARenderCameraState = function() {
  this.up  = new THREE.Vector3(0, 0, 1);
  this.pos = new THREE.Vector3(0, 0, 0);
  this.target = new THREE.Vector3(0, 0, 0);
}

/**
* Look up tables.
*/
MARenderColormap = {
  CUSTOM:		0,
  GREYSCALE:		1,
  COOLTOWARM:           2,
  BLACKBODY:		3,
  RAINBOW:		4
}

MARenderLookUpTable = function(colormap) {
  this.type = 'MARenderer';
  this.colormap = colormap;
  this.map = new Array(256);
  var mappts = undefined;

  switch(self.colormap)
  {
    case MARenderColormap.GREYSCALE:
      mappts = [[0x00, 0x00, 0x00],
                [0x40, 0x40, 0x40]
                [0x7f, 0xf0, 0xf0]
                [0xbf, 0xbf, 0xbf]
                [0xff, 0xff, 0xff]];
      break;
    case MARenderColormap.COOLTOWARM:
      mappts = [[0x00, 0x00, 0x00],
                [0x40, 0x40, 0x40]
                [0x7f, 0xf0, 0xf0]
                [0xbf, 0xbf, 0xbf]
                [0xff, 0xff, 0xff]];
      break;
    case MARenderColormap.BLACKBODY:
      mappts = [[0x00, 0x00, 0x00],
                [0x80, 0x00, 0x00]
                [0xff, 0x40, 0x00]
                [0xff, 0xff, 0x00]
                [0xff, 0xff, 0xff]];
      break;
    case MARenderColormap.RAINBOW:
      mappts = [[0x00, 0x00, 0xff],
                [0x00, 0xff, 0xff]
                [0x00, 0xff, 0x00]
                [0xff, 0xff, 0x00]
                [0xff, 0x00, 0xff]];
      break;
    default:
      break;
  }
  if(mappts !== undefined) {
    for(var idx = 0; idx < 256; ++idx) {
      var p = (idx / 64) | 0;
      var p1 = p + 1;
      var q = idx - (p * 64);
      var q1 = 64 - q;
      var n = 64.0 * 255.0;
      var r = ((q1 * mappts[p][0]) + (q * mappts[p1][0])) / n;
      var g = ((q1 * mappts[p][1]) + (q * mappts[p1][1])) / n;
      var b = ((q1 * mappts[p][2]) + (q * mappts[p1][2])) / n;
      map.append(new THREE.Color(r, g, b));
    }
  }
}

/**
* Point rendering material
*/
AlphaPointsMaterial = function(params) {
  var vertexShader = [
    //AlphaPoint vertex shader
    'attribute float sizes;',
    'attribute float opacities;',
    'attribute vec3 colors;',
    'uniform float scale;',
    'varying vec3 vColor;',
    'varying float vOpacity;',
    '#include <common>',
    //'#include <logdepthbuf_pars_vertex>',
    '#include <clipping_planes_pars_vertex>',
    'void main() {',
      'vColor = colors;',
      'vOpacity = opacities;',
      '#include <begin_vertex>',
      '#include <project_vertex>',
      'gl_PointSize = 512.0 * sizes * scale / length(mvPosition.xyz);',
      '#include <clipping_planes_vertex>',
    '}'
  ].join('\n');
  var fragmentShader = [
    //AlphaPoint fragment shader
    'uniform vec3 diffuse;',
    'uniform float opacity;',
    'uniform sampler2D map;',
    'varying vec3 vColor;',
    'varying float vOpacity;',
    '#include <common>',
    '#include <clipping_planes_pars_fragment>',
    'void main() {',
      '#include <clipping_planes_fragment>',
      'gl_FragColor = vec4(diffuse * vColor, opacity * vOpacity);',
      'gl_FragColor = gl_FragColor * texture2D(map, gl_PointCoord);',
      'if(gl_FragColor.a < ALPHATEST) {',
        'discard;',
      '}',
    '}'
  ].join('\n');
  var splitParam = [ {
    uniforms: THREE.UniformsUtils.clone(THREE.UniformsLib['points']),
    vertexShader: vertexShader,
    fragmentShader: fragmentShader
  }, {}];
  for(var p in params) {
    switch(p) {
      case 'map':
      case 'color':
	splitParam[1][p] = params[p];
	break
      case 'scale':
      case 'size':
        splitParam[1]['scale'] = params[p];
	break;
      default:
	splitParam[0][p] = params[p];
	break;
    }
  }
  var material = new THREE.ShaderMaterial(splitParam[0]);
  material.type = 'AlphaPointsMaterial';
  for(var p in splitParam[1]) {
    switch(p) {
      case 'color':
	material.uniforms.diffuse.value = new THREE.Color(
	                                      splitParam[1]['color']);
	break;
      case 'map':
	material.uniforms.map.value = splitParam[1]['map'];
	break;
      case 'scale':
	material.uniforms.scale.value = splitParam[1]['scale'];
	break;
    }
  }
  return(material);
}

/*!
* \class	MARenderer
* \constructor
* \brief 	Creates a new renderer.
* \param win	The window used by the renderer.
* \param con	The container used by the renderer. A new div for the
* 		renderer will be created within this container.
*/
MARenderer = function(win, con) {
  var self = this;
  this.type = 'MARenderer';
  Object.defineProperty(self, 'version', {value: '1.3.1', writable: false});
  this.win = win;
  this.con = con;
  this.scene;
  this.ambLight;
  this.dirLight;
  this.pntLight;
  this.camera;
  this.cameraControls;
  this.renderer;
  this.animCount = 0;    // Used to count animation frames since mouse movement
  this.pointSize = 10;
  this.mousePos = new THREE.Vector2(0,0);
  this.pickOfs = 0;
  this.nearPlane = 1;
  this.farPlane = 10000;
  this.setCamOnLoad = true;
  this.setHomeOnLoad = true;
  this.useCameraControl = true;
  this.conOffset = new THREE.Vector2(0,0);
  this.cameraPos = new THREE.Vector3(0, 0, 10000);
  this.center   = new THREE.Vector3(0, 0, 0);
  this.homeCameraView = new MARenderCameraState();
  this.saveCameraView = new MARenderCameraState();
  this.eventHandler = new THREE.EventDispatcher();
  this.noClipPlanes = Object.freeze([]);
  this.globalClipping = false;
  this.globalClipPlanes = this.noClipPlanes;


  THREE.ImageUtils.crossOrigin = ''; // To allow CORS textures

  this.init = function() {

    this.scene = new THREE.Scene();

    this.setConOffset();
    this.camera = new THREE.PerspectiveCamera(25,
				   this.con.clientWidth / this.con.clientHeight,
				   this.nearPlane, this.farPlane);

    this.camera.updateProjectionMatrix();
    this.cameraControls = this._makeCameraControls();

    this.renderer = new THREE.WebGLRenderer({antialias: true});
    this.renderer.setSize(this.con.clientWidth, this.con.clientHeight);
    this.con.appendChild(this.renderer.domElement);

    this.scene.add(this.camera);
    
    this.ambLight = new THREE.AmbientLight(0x777777);
    this.dirLight = new THREE.DirectionalLight(0x777777);
    this.dirLight.position.set(0, 0, 1);
    this.pntLight = new THREE.PointLight(0x333333, 1, 10000);
    this.pntLight.position.set(0, 0.5, 0);
    this.scene.add(this.pntLight);
    this.camera.add(this.ambLight);
    this.camera.add(this.dirLight);

    this.raycaster = new THREE.Raycaster();

    self.win.addEventListener('mousemove', self._trackMouse, false);
    self.win.addEventListener('keypress', self._keyPressed, false);
    self.win.addEventListener('resize', self._windowResize, false);
  }

  /*!
   * \class	MARenderer
   * \return	True if local clipping is on or false if it's off.
   * \function	setClipping
   * \brief	Sets the local (per model) clipping on or off.
   */
  this.setLocalClipping = function(state) {
    self.renderer.localClippingEnabled = Boolean(state);
  }

  /*!
   * \class	MARenderer
   * \function	setClipping
   * \brief	Gets the state of local (per model) clipping (on or off).
   * \param state	Local clipping on if true or off if false.
   */
  this.getLocalClipping = function() {
    return(self.renderer.localClippingEnabled);
  }

  /*!
   * \class	MARenderer
   * \function	setClipping
   * \brief	Sets the global clipping plane on or off.
   * \param state	Clipping plane on if true or off if false.
   */
  this.setGlobalClipping = function(state) {
    self.globalClipping = Boolean(state);
    if(self.globalClipping) {
      self.renderer.clippingPlanes = this.globalClipPlanes;
    } else {
      self.renderer.clippingPlanes = this.noClipPlanes;
    }
    this.render();
  }

  /*!
   * \class	MARenderer
   * \function	setClipping
   * \brief	Sets the global clipping plane.
   * \param pln		New clipping plane.
   */
  this.setGlobalClipPlane = function(pln) {
    if(Boolean(pln) && (pln instanceof THREE.Plane)) {
      self.globalClipPlanes = [pln];
      if(self.globalClipping) {
        self.renderer.clippingPlanes = self.globalClipPlanes;
      }
      this.render();
    }
  }

  /*!
   * \class     MARenderer
   * \function	setPickOfs
   * \brief	Adds an offset to the mouse coords to compensate for title
   *            div above the canvas. For backwards compatability only.
   * \param ofs	height of title div in pixels
   */
  this.setPickOfs = function(ofs) {

     var iofs;
     var innerH;

     innerH = self.win.innerHeight;
     if(ofs !== undefined) {
        iofs = parseInt(ofs, 10);
        self.pickOfs = (iofs / self.win.innerHeight) * 2;
     }
  }

  /*!
   * \class     MARenderer
   * \function	addModel
   * \brief	Adds a new model to the renderer using the given properties.
   * \param gProp	Model properties which conform to MARenderItem and
   * 			must include a unique name.
   */
  this.addModel = function(gProp) {
    var loader;
    var itm = this._makeRenderItem(gProp);
    if(itm) {
      switch(Number(itm.mode)) {
	case MARenderMode.BASIC:
	case MARenderMode.WIREFRAME:
	case MARenderMode.LAMBERT:
	case MARenderMode.PHONG:
	case MARenderMode.EMISSIVE:
	case MARenderMode.POINT:
	  if(itm.path) {
	    var ext = itm.path.split('.').pop();
	    if(ext === 'stl') {
	      loader = new THREE.STLLoader();
	    } else if(ext === 'vtk') {
	      loader = new THREE.VTKLoader();
	    } else {
	      console.log('MARenderer.addModel() unknown file type: ' + ext);
	    }
	    loader.load(itm.path,
	      function(geom) {
		var mat = self._makeMaterial(geom, itm);
		if(mat) {
		  switch(Number(itm.mode)) {
		    case MARenderMode.POINT:
		      var pnts = new THREE.Points(geom, mat);
		      pnts.name = itm.name;
		      pnts.sortParticles = true;
		      self.scene.add(pnts);
		      break;
		    default:
		      var mesh = new THREE.Mesh(geom, mat);
		      mesh.name = itm.name;
		      self.scene.add(mesh);
		      break;
		  }
		  if(self.setCamOnLoad) {
		    self._computeCenter();
		    self.setCamera();
		  }
		  if(self.setHomeOnLoad) {
		    self.setHome();
		  }
		  self.makeLive();
		}
	      },
	      function(){},
	      function() {
		console.log('MARenderer.addModel() geometry load failed: ' +
		            itm.path);
	      });
	  }
	  break;
	case MARenderMode.SECTION:
	  if(itm.texture) {
	    var secLoader = new THREE.TextureLoader();
	    secLoader.crossOrigin = '';
	    secLoader.load(itm.texture,
		 function(tx) {
		   var geom = self.makeSectionGeometry(itm.vertices);
		   var mat = self._makeMaterial(geom, itm);
		   tx.flipY = false;
		   tx.minFilter = THREE.LinearFilter;
		   tx.needsUpdate = true;
		   mat.map = tx;
		   var pln = new THREE.Mesh(geom, mat);
		   pln.name = itm.name;
		   self.scene.add(pln);
		   self.makeLive();
		 },
		 function() {
		   console.log('MARenderer.addModel() texture load failed: ' +
			       itm.texture);
		 });
	  }
	  break;
	default:
	  break;
      }
    }
  }

  /*!
   * \class     MARenderer
   * \function	updateModel
   * \brief	Updates an existing model of the renderer using the given
   * 		properties.
   * \param gProp	Model properties which conform to MARenderItem and
   * 			must include a unique name.
   */
  this.updateModel = function(gProp) {
    if(gProp['name']) {
      var name = gProp['name'];
      var obj = this.scene.getObjectByName(name, true);
      if(obj) {
	this._updateObj(obj, gProp);
      }
    }
    this.render();
  }

  /*!
   * \class	MARenderer
   * \function	makeSectionGeometry
   * \return	New geometry.
   * \brief	Create a geometry for use in rendering planar sections.
   * \param vertices	Array of four ordered 3D vertices at the position
   * 			of the rectangular sections corners.
   */
  this.makeSectionGeometry = function(vertices) {
    var geom = new THREE.Geometry();
    if(vertices) {
      geom.vertices = vertices.slice(0);
    } else {
      geom.vertices = [new THREE.Vector3(0, 0, 0),
		       new THREE.Vector3(1, 0, 0),
		       new THREE.Vector3(1, 1, 0),
		       new THREE.Vector3(0, 1, 0)];
    }
    geom.faces = [new THREE.Face3(0, 1, 2),
		  new THREE.Face3(2, 3, 0)];
    geom.faceVertexUvs[0] = [[new THREE.Vector2(0, 0),
			      new THREE.Vector2(1, 0),
			      new THREE.Vector2(1, 1)],
			    [new THREE.Vector2(1, 1),
			     new THREE.Vector2(0, 1),
			     new THREE.Vector2(0, 0)]];
    geom.computeFaceNormals();
    geom.computeBoundingBox();
    return(geom);
  }

  /*!
   * \class	MARenderer
   * \function	makePlaneFromAngles
   * \return	New plane.
   * \brief	Create a plane using sectioning angles, a fixed point
   * 		and a distance from the fixed point ass as used in
   * 		Woolz and WlzIIPSrv.
   * \param pitch	Pitch angle.
   * \param yaw		Yaw angle.
   * \param fixed 	Fixed point.
   * \param dst		Prependicular distance from the fixed point to the
   * 			plane.
   */
  this.makePlaneFromAngles = function(pit, yaw, fxd, dst) {
    var cp = Math.cos(pit);
    var cy = Math.cos(yaw);
    var sp = Math.sin(pit);
    var sy = Math.sin(yaw);
    var nrm = new THREE.Vector3(sp * cy, sp * sy, cp);
    dst += (fxd.x * nrm.x) + (fxd.y * nrm.y) + (fxd.z * nrm.z);
    var pln = new THREE.Plane(nrm, -dst);
    return(pln);
  }

  /*!
   * \class	MARenderer
   * \function	setConOffset
   * \brief	Sets the offset of the container from the window.
   */
  this.setConOffset = function() {
    var c = self.con;
    var o = new THREE.Vector2(0, 0);
    while(c) {
      o.x += c.offsetLeft;
      o.y += c.offsetTop;
      c = c.offsetParent;
    }
    self.conOffset.copy(o);
  }


  /*!
   * \class	MARenderer
   * \function	makePlaneFromVertices
   * \return	New plane.
   * \brief	Create a plane using the first three vertices of the given
   * 		array of 3D vertices.
   * \param vtx		Array of vertices.
   */
  this.makePlaneFromVertices = function(vtx) {
    var pln;
    if(vtx && (vtx instanceof Array) && (vtx.length >= 3)) {
      var pln = new THREE.Plane();
      pln.setFromCoplanarPoints (vtx[0], vtx[1], vtx[2]);
    }
    return(pln);
  }

  /*!
   * \class	MARenderer
   * \function	setCamera
   * \brief	Sets the camera for the renderer.
   * \param cen		Centre of the scene.
   * \param near	Camera near plane.
   * \param far		Camera far plane.
   * \param pos		Camera position.
   */
  this.setCamera = function(cen, near, far, pos) {
    if(cen || near || far || pos) {
      this.setCamOnLoad = false;
      if(cen) {
	this.center.copy(cen);
      }
      if(near) {
	this.nearPlane = near;
      }
      if(far) {
	this.farPlane = far;
      }
      if(pos) {
	this.cameraPos.copy(pos);
      }
    } else {
      this._computeCenter();
    }
    this.camera.near = this.nearPlane;
    this.camera.far = this.farPlane;
    this.camera.updateProjectionMatrix();
    this.camera.position.copy(this.cameraPos);
  }

  /*!
   * \class	MARenderer
   * \function	setHome
   * \brief	Sets the camera home position.
   * \param pos		Given camera home position, default is the current
   * 			camera trackball position.
   * \param up		Given camera up vector, default is the current
   * 			camera trackball up vector.
   */
  this.setHome = function(pos, up) {
    if(this.cameraControls) {
      if(pos || up) {
	this.setHomeOnLoad = false;
      }
      if(pos === undefined) {
	pos = this.cameraControls.object.position.clone();
      }
      if(up === undefined) {
	up = this.cameraControls.object.up.clone();
      }
      this.homeCameraView.up.copy(up);
      this.homeCameraView.pos.copy(pos);
      this.goHome();
    }
  }

  /*!
   * \class	MARenderer
   * \function	goHome
   * \brief	Moves the camera trackball to the current home position.
   */
  this.goHome = function() {
    if(this.cameraControls) {
      this.cameraControls.up0.copy(this.homeCameraView.up);
      this.cameraControls.position0.copy(this.homeCameraView.pos);
      this.cameraControls.target0.copy(this.center);
      this.cameraControls.reset();
    }
  }

  /*!
   * \class	MARenderer
   * \function	removeModel
   * \brief	Removes the named model and frees its resources.
   * \param name	Given model name.
   */
  this.removeModel = function(name) {
    var obj = this.scene.getObjectByName(name, true);
    if(obj) {
      this.scene.remove(obj);
      if(obj.material) {
        obj.material.dispose();
      }
      if(obj.geometry) {
        obj.geometry.dispose();
      }
      this.render();
    }
  }

  /*!
   * \class	MARenderer
   * \function	getChildren
   * \brief	Gets the children of the scene.
   */
  this.getChildren = function() {
    return this.scene.children;
  }

  /*!
   * \class     MARenderer
   * \return	Clamped point size.
   * \function  _pointSizeClamp
   * \brief	Clamps the given point size to the allowed range.
   * \param	sz	Given point size.
   */
  this._opacityClamp = function(op) {
    if(op > 1.0) {
      op = 1.0;
    }
    else if(op < 0.0) {
      op = 0.0;
    }
    return(op);
  }

  /*!
   * \class	MARenderer
   * \function	opacityIncrement
   * \brief	Increments the opacity of all transparent models.
   * \param inc		Opacity increment which may be positive or
   * 			negative.
   */
  this.opacityIncrement = function(inc) {
    var update = false;
    for(var i = 0, l = this.scene.children.length; i < l; i ++ ) {
      var child = this.scene.children[i];
      if(child && (child.type === 'Mesh')) {
        if(child.material && child.material.transparent &&
	   (child.material.opacity != undefined)) {
	  var op = child.material.opacity;
	  var tr = child.material.transparent;
	  if((inc > 0.0) && (op < inc)) {
	    op = inc;
	  }
	  else {
	    op = op * (1.0 + inc);
	  }
	  op = this._opacityClamp(op);
	  this._setMaterialOpacity(child.material, tr, op);
	  child.material.needsUpdate = true;
	}
      }
    }
    if(update) {
      this.render();
    }
  }

  /*!
   * \class     MARenderer
   * \return	Clamped point size.
   * \function  _pointSizeClamp
   * \brief	Clamps the given point size to the allowed range.
   * \param	sz	Given point size.
   */
  this._pointSizeClamp = function(sz) {
    if(sz > 99.9) {
      sz = 99.9;
    }
    else if(sz < 0.1) {
      sz = 0.1;
    }
    return(sz);
  }

  /*!
   * \class	MARenderer
   * \function	pointSizeSet
   * \brief	Sets the current default point size and the point size of
   * 		all point cloud models.
   * \param sz		Given point size. By default the current point size
   * 			is used.
   */
  this.pointSizeSet = function(sz) {
    var update = false;
    if(sz !== undefined) {
      this.pointSize = this._pointSizeClamp(sz);
    }
    for(var i = 0, l = this.scene.children.length; i < l; i ++ ) {
      var child = this.scene.children[i];
      if(child && (child.type === 'Points')) {
	var mat = child.material;
	if(mat) {
	  if(mat.type == 'PointsMaterial') {
	    update = true;
	    mat.needsUpdate = true;
	    mat.size = this.pointSize;
	  }
	  else if(mat.type =='AlphaPointsMaterial') {
	    update = true;
	    mat.needsUpdate = true;
	    mat.uniforms.scale.value = this.pointSize;
	  }
	}
      }
    }
    if(update) {
      this.render();
    }
  }

  /*!
   * \class	MARenderer
   * \function	pointSizeIncrement
   * \brief	Increments the point size of all point cloud models.
   * \param inc		Increment which may be positive or negative.
   */
  this.pointSizeIncrement = function(inc) {
    this.pointSize = this._pointSizeClamp(this.pointSize * (1.0 + inc));
    this.pointSizeSet();
  }

  /*!
   * \class	MARenderer
   * \function	render
   * \brief	Renders the scene now.
   */
  this.render = function() {
    this.renderer.render(self.scene, self.camera);
  }

  /*!
   * \class	MARenderer
   * \function	setCameraControl
   * \brief	Sets camera control by the trackball on or off.
   */
  this.setCameraControl = function(state) {
    var newState = Boolean(state);
    if(newState != this.useCameraControl) {
      if(this.useCameraControl) {
	if(this.cameraControls) {
	  this.saveCameraView.up.copy(this.cameraControls.object.up.clone());
	  this.saveCameraView.pos.copy(
		  this.cameraControls.object.position.clone());
	  this.saveCameraView.target.copy(this.center);
	  this.cameraControls.enabled = false;
	}
      } else {
	//this.cameraControls = this._makeCameraControls();
	this.cameraControls.enabled = true;
	this.cameraControls.up0.copy(this.saveCameraView.up);
	this.cameraControls.position0.copy(this.saveCameraView.pos);
	this.cameraControls.target0.copy(this.saveCameraView.target);
	this.cameraControls.reset();
      }
      this.useCameraControl = newState;
    }
  }

  /*!
   * \class	MARenderer
   * \function	setCameraControl
   * \brief	Gets camera control by the trackball state.
   */
  this.getCameraControl = function() {
    return(this.useCameraControl);
  }

  /*!
   * \class	MARenderer
   * \function	animate
   * \brief	Starts rendering the scene. The rendering will be repeated
   * 		under the control of a timer and following mouse movement
   * 		or scene updates.
   */
  this.animate = function() {
    var aid = self.win.requestAnimationFrame(self.animate);
    if(self.useCameraControl && self.cameraControls) {
      self.cameraControls.update();
    }
    self.render();
    if(++(self.animCount) > 400) {
      self.win.cancelAnimationFrame(aid);
    }
  }

  /*!
   * \class	MARenderer
   * \function	makeLive
   * \brief	Makes sure that the scene is rendered and resets the
   * 		rendering timer.
   */
  this.makeLive = function() {
    var count = this.animCount;
    this.animCount = 0;
    if(count > 200) {
      this.animate();
    }
  }

  /*!
   * \class 	MARenderer
   * \function	addEventListener
   * \brief	Adds the event listener.
   * \param type	Event type.
   * \param listener	The event listener function.
   */
  this.addEventListener = function(type, listener) {
    this.eventHandler.addEventListener(type, listener)
  }

  /*!
   * \class 	MARenderer
   * \function	removeEventListener
   * \brief	Removes the event listener.
   * \param type	Event type.
   * \param listener	The event listener function.
   */
  this.removeEventListener = function(type, listener) {
    this.eventHandler.removeEventListener(type, listener)
  }

  /*!
   * \class	MARenderer
   * \function	getIIP3DBBVertices
   * \brief	Get vertices of the section defined by the IIP3D request
   *            encoded in the given URL using voxel scaling.
   * \param url		The IIP3D URL which should define the section but
   *                    not include an OBJ, CVT or tile request.
   * \param vsz		The voxel size, supplied as a THREE.Vector3().
   */
  this.getIIP3DBBVertices = function(url, vsz) {
    var prmX = [0, 1, 1, 0];
    var prmY = [0, 0, 1, 1];
    var max = new THREE.Vector2();
    var vtx = [new THREE.Vector3(), new THREE.Vector3(),
	       new THREE.Vector3(), new THREE.Vector3()];
    var req = new XMLHttpRequest();
    req.open('GET', url + '&OBJ=Wlz-true-voxel-size', false);
    req.send(null);
    req.open('GET', url + '&OBJ=Max-size', false);
    req.send(null);
    if(req.status === 200) {
      // rsp = Max-size:321 173
      var rsp = req.responseText.split(':')[1].split(' ');
      max.x = rsp[0] - 1.0;
      max.y = rsp[1] - 1.0;
    }
    for(var idx = 0; idx < 4; ++idx) {
      req.open('GET', url + '&PRL=-1,' +
	       prmX[idx] * max.x + ',' +
	       prmY[idx] * max.y +
               '&OBJ=Wlz-coordinate-3D', false);
      req.send(null);
      if(req.status === 200) {
	var rsp = req.responseText.split(':')[1].split(' ');
	vtx[idx].set(rsp[0] * vsz.x, rsp[1] * vsz.y, rsp[2] * vsz.z);
      }
    }
    return(vtx);
  }

  /*!
   * \class	MARenderer
   * \return	New camera controls.
   * \function	_makeCameraControls
   * \brief	Initialises the camera trackball controls.
   */
  this._makeCameraControls = function() {
    var cc = new THREE.TrackballControls(this.camera, this.con);
    cc.panSpeed = 0.3;
    cc.dynamicDampingFactor = 0.7;
    return(cc);
  }

  /*!
   * \class	MARenderer
   * \function	_setMaterialOpacity
   * \brief	Sets the opacity of the given material.
   * \param mat		Given material.
   * \param tr		True if material is tranparent, if false function
   * 			simply returns.
   * \param op		Given opacity.
   */
  this._setMaterialOpacity = function(mat, tr, op) {
    if(mat && tr) {
      if(op < 0.01) {
	mat['opacity'] = 0.0;
      } else {
	if(op > 1.0) {
	  op = 1.0;
	}
	if(op < 0.9) {
	  mat['depthWrite'] = false;
	} else {
	  mat['depthWrite'] = true;
	}
	mat['opacity'] = op;
      }
      this.render();
    }
  }

  /*!
   * \class 	MARenderer
   * \function	_updateObj
   * \brief	Updates the given objects properties.
   * \param obj		Given object.
   * \param gProp	Properties to set in the given object.
   */
  this._updateObj = function(obj, gProp) {
    var itm = new MARenderItem();
    if(itm) {
      if(gProp['color']) {
	itm.color = gProp['color'];
      } else if(obj.material && obj.material.color) {
	itm.color = obj.material.color;
      }
      if(gProp['clipping'] !== undefined) {
        if(gProp['clipping'] instanceof THREE.Plane) {
	  var pln = new THREE.Plane().copy(gProp['clipping']);
	  itm.clipping = [ pln ];
        } else {
	  itm.clipping = this.noClipPlanes;
	}
      } else if(obj.material) {
	itm.clipping = obj.material.clippingPlanes;
      }
      if(gProp['opacity']) {
	itm.opacity = gProp['opacity'];
      } else if(obj.material && obj.material.opacity) {
	itm.opacity = obj.material.opacity;
      }
      if(gProp['transparent'] !== undefined) {
	itm.transparent = gProp['transparent'];
      } else if(obj.material && obj.material.transparent) {
	itm.transparent = obj.material.transparent;
      }
      if(gProp['side'] !== undefined) {
	itm.side = gProp['side'];
      } else if(obj.material) {
        itm.side = obj.material.side;
      }
      if(gProp['visible'] !== undefined) {
	itm.visible = gProp['visible'];
      } else if(obj.material) {
        itm.visible = obj.material.visible;
      }
      if(gProp['texture']) {
	itm.texture = gProp['texture'].slice(0);
      }
      if(gProp['vertices']) {
	itm.vertices = gProp['vertices'].slice(0);
      }
      if(gProp['mode']) {
	// Always set the mode/material type
	var mode = this._checkRenderMode(gProp['mode']);
	if(mode) {
	  itm.mode = mode;
	}
      } else {
        if(obj.type === 'Points') {
	  itm.mode = MARenderMode.POINT;
	} else if(obj.material && (obj.material.map)) {
	  itm.mode = MARenderMode.SECTION;
	}
      }
    }
    switch(Number(itm.mode)) {
      case MARenderMode.BASIC:
      case MARenderMode.WIREFRAME:
      case MARenderMode.LAMBERT:
      case MARenderMode.PHONG:
      case MARenderMode.EMISSIVE:
      case MARenderMode.POINT:
        var mat = this._makeMaterial(obj.geometry, itm);
	var oldmat = obj.material;
	obj.material = mat;
	if(oldmat) {
	  oldmat.dispose();
	}
	break;
      case MARenderMode.SECTION:
	{
	  obj.material.visible = itm.visible;
	  obj.material.opacity = itm.opacity;
	  if(itm.texture) {
	    THREE.ImageUtils.loadTexture(itm.texture,
		THREE.UVMapping,
		function(tex) {
		  // onLoad
		  var oldgeom = obj.geometry;
		  var oldtex = obj.material.map;
		  var geom = self.makeSectionGeometry(itm.vertices);
		  tex.flipY = false;
		  tex.minFilter = THREE.LinearFilter;
		  tex.needsUpdate = true;
		  obj.material.map = tex;
		  obj.geometry = geom;
		  oldtex.dispose();
		  oldgeom.dispose();
		  self.makeLive();
		},
		function() {
	          console.log('MARenderer.updateObj() texture load failed: ' +
		              itm.texture);
		});
	  }
	}
	break;
    }
  }

  /*!
   * \class	MARenderer
   * \function	_updateAllMesh
   * \brief	Updates the properties of all meshes.
   * \param gProp	Given properties to update.
   */
  this._updateAllMesh = function(gProp) {
    for(var i = 0, l = this.scene.children.length; i < l; i ++ ) {
      var child = this.scene.children[i];
      if(child && (child.type === 'Mesh')) {
        if(child.material && !(child.material.map)) {
	  this._updateObj(child, gProp);
          this.render();
        }
      }
    }
  }

  /*!
   * \class	MARenderer
   * \function	_makeRenderItem
   * \return	New render item.
   * \brief	Create a new render item the given properties set.
   * 		All properties not given have the default value.
   * \param gProp	Given properties.
   */
  this._makeRenderItem = function(gProp) {
    var ok = true;
    var itm = new MARenderItem();
    for(var p in gProp) {
      switch(p) {
        case 'name':
        case 'path':
	case 'side':
          itm[p] = gProp[p];
          break;
        case 'color':
        case 'opacity':
	case 'pitch':
	case 'yaw':
	case 'dist':
	  itm[p] = Number(gProp[p]);
          break;
        case 'transparent':
          itm[p] = Boolean(gProp[p]);
          break;
        case 'mode':
	  var mode = this._checkRenderMode(gProp[p]);
	  if(mode) {
	    itm[p] = mode;
	  }
	  break;
	case 'vertices':
	  itm[p] = gProp[p].slice(0);
	  break;
	case 'texture':
	  itm[p] = gProp[p].slice(0);
	  break;
	case 'visible':
	  itm[p] = Boolean(gProp[p]);
	  break;
	case 'clipping':
	  if(gProp[p] && (gProp[p] instanceof THREE.Plane)) {
	    var pln = new THREE.Plane().copy(gProp[p]);
	    itm.clipping = [ pln ];
	  } else {
	    itm.clipping = this.noClipPlanes;
	  }
	  break;
        default:
	  ok = false;
	  console.log('MARenderer._makeRenderItem() unknown property: ' + p);
	  break;
      }
    }
    if(!ok) {
      itm = undefined;
    }
    return(itm);
  }

  /*!
   * \class	MARenderer
   * \function	_checkRenderMode
   * \return	Given render mode or undefined if the render mode is invalid.
   * \brief	Checks that the given render mode is valid.
   * \param gMode	Given render mode.
   */
  this._checkRenderMode = function(gMode) {
    var rMode = undefined;
    if(gMode) {
      switch(Number(gMode)) {
	case MARenderMode.BASIC:
	case MARenderMode.WIREFRAME:
	case MARenderMode.LAMBERT:
	case MARenderMode.PHONG:
	case MARenderMode.EMISSIVE:
	case MARenderMode.POINT:
	case MARenderMode.SECTION:
	  rMode = gMode;
	  break;
	default:
	  console.log('MARenderer: Unknown mode: ' + gMode);
	  break;
      }
    }
    return(rMode)
  }

  /*!
   * \class	MARenderer
   * \function	_makeMaterial
   * \return	New material.
   * \brief	Makes a new material using the properties of the given render
   * 		item.
   * \param itm		Given render item.
   */
  this._makeMaterial = function(geom, itm) {
    var mat;
    var sProp = {};
    switch(itm.mode) {
      case MARenderMode.BASIC:
	sProp['color'] = itm.color;
	sProp['visible'] = itm.visible;
	sProp['clippingPlanes'] = itm.clipping;
	sProp['wiretrame'] = false;
	mat = new THREE.MeshBasicMaterial(sProp);
	break;
      case MARenderMode.WIREFRAME:
	sProp['color'] = itm.color;
	sProp['opacity'] = itm.opacity;
	sProp['visible'] = itm.visible;
	sProp['clippingPlanes'] = itm.clipping;
	sProp['transparent'] = itm.transparent;
	sProp['wireframe'] = true;
	sProp['wireframeLinewidth'] = 1;
	mat = new THREE.MeshLambertMaterial(sProp);
	break;
      case MARenderMode.LAMBERT:
	sProp['color'] = itm.color;
	sProp['opacity'] = itm.opacity;
	sProp['visible'] = itm.visible;
	sProp['clippingPlanes'] = itm.clipping;
	sProp['transparent'] = itm.transparent;
	sProp['wireframe'] = false;
	sProp['side'] = itm.side;
	mat = new THREE.MeshLambertMaterial(sProp);
	break;
      case MARenderMode.PHONG:
	sProp['color'] = itm.color;
	sProp['opacity'] = itm.opacity;
	sProp['visible'] = itm.visible;
	sProp['clippingPlanes'] = itm.clipping;
	sProp['transparent'] = itm.transparent;
	sProp['specular'] = 0x111111;
	sProp['wireframe'] = false;
	sProp['side'] = itm.side;
	sProp['emissive'] = 0x000000;
	sProp['shininess'] = 25;
	this._setMaterialOpacity(sProp, itm.transparent, itm.opacity);
	mat = new THREE.MeshPhongMaterial(sProp);
	break;
      case MARenderMode.EMISSIVE:
	sProp['color'] = itm.color;
	sProp['opacity'] = itm.opacity;
	sProp['visible'] = itm.visible;
	sProp['clippingPlanes'] = itm.clipping;
	sProp['transparent'] = itm.transparent;
	sProp['specular'] =0x777777;
	sProp['wireframe'] = false;
	sProp['emissive'] = itm.color;
	sProp['shininess'] = 15;
	sProp['side'] = itm.side;
	mat = new THREE.MeshPhongMaterial(sProp);
	break;
      case MARenderMode.POINT:
	sProp['color'] = itm.color;
	sProp['opacity'] = itm.opacity;
	sProp['visible'] = itm.visible;
	sProp['transparent'] = itm.transparent;
	sProp['clippingPlanes'] = itm.clipping;
	sProp['size'] = this.pointSize;
	sProp['blending'] = THREE.CustomBlending;
	sProp['blendSrc'] = THREE.SrcColorFactor;
	sProp['blendDst'] = THREE.DstAlphaFactor;
	sProp['blendEquation'] = THREE.AddEquation;
	sProp['alphaTest'] = 0.1;
	sProp['map'] =        THREE.ImageUtils.loadTexture(
	    'data:image/png;base64,' +
	    'iVBORw0KGgoAAAANSUhEUgAAAAgAAAAICAQAAABuBnYAAAAAAmJLR0QA/4eP' +
	    'zL8AAAAJcEhZcwAACxMAAAsTAQCanBgAAAAHdElNRQfgAg8MNSkRqlGqAAAA' +
	    'VElEQVQI113NQQ0DIRBA0TcEFYQTSVWsAHRUWXUgoDY4bbAxPfS2X8D7ATk1' +
	    'nFhU8u0ysLPFp+Z0mTpe5CmaoYNuaMWj4thucNtOjZWNP+obK57bH17lGKmO' +
	    'V2FkAAAAAElFTkSuQmCC')
	if((geom.attributes.colors !== undefined) &&
	   (geom.attributes.colors.array.length > 0)) {
	  mat = new AlphaPointsMaterial(sProp);
	  mat.vertexColors = THREE.VertexColors;
	} else {
	  mat = new THREE.PointsMaterial(sProp);
	}
	break;
      case MARenderMode.SECTION:
	sProp['color'] = itm.color;
	sProp['opacity'] = itm.opacity;
	sProp['transparent'] = itm.transparent;
	sProp['alphaTest'] = 0.2;
	sProp['visible'] = itm.visible;
	sProp['clippingPlanes'] = itm.clipping;
	sProp['wireframe'] = false;
	sProp['side'] = THREE.DoubleSide;
	mat = new THREE.MeshBasicMaterial(sProp);
	break;
    }
    return(mat);
  }

  /*!
   * \class	MARenderer
   * \function	_computeCenter
   * \brief	Computes the scene centre and sets the camera trackball to
   * 		view the centre.
   */
  this._computeCenter = function() {
    var n = 0;
    var box = new THREE.Box3();
    for(var i = 0, l = this.scene.children.length; i < l; i ++ ) {
      var child = this.scene.children[i];
      if(child && 
         ((child.type === 'Mesh') ||
	  (child.type === 'Points'))) {
	++n;
	var b = new THREE.Box3();
	b.setFromObject(child);
	b.min.add(child.position);
	b.max.add(child.position);
        if(n == 1) {
	  box.copy(b);
	}
	else {
	  box.union(b);
	}
      }
    }
    if(n > 0) {
      var d, min, max, dMax;
      min = box.min.x;
      max = box.max.x;
      dMax = box.max.x - box.min.x;
      d = box.max.y - box.min.y;
      if(d > dMax)
      {
        dMax = d;
      }
      if(min > box.min.y)
      {
        min = box.min.y;
      }
      if(max < box.max.y)
      {
        max = box.max.y;
      }
      d = box.max.z - box.min.z;
      if(d > dMax)
      {
        dMax = d;
      }
      if(min > box.min.z)
      {
        min = box.min.z;
      }
      if(max < box.max.z)
      {
        max = box.max.z;
      }
      this.center.copy(box.center());
      this.nearPlane = (min < 0.2)? 0.1: min * 0.5;
      this.farPlane =  (max < 1.0)? 10.0: max * 10.0;
      this.cameraPos.set(0, 0, this.center.z + (4.0 * dMax));
    }
  }

  /*!
   * \class	MARenderer
   * \function	_testCode
   * \brief	Prints debug output to the console.
   */
  this._testCode	= function() {
    console.log('ren.version = ' + self.version);
    console.log('cen = (' +
		self.center.x + ', ' +
		self.center.y + ', ' +
		self.center.z + ')');
    console.log('near = ' + self.nearPlane);
    console.log('far = ' + self.farPlane);
    console.log('pos = ' + 
                 self.camera.position.x + ', ' +
		 self.camera.position.y + ', ' +
		 self.camera.position.z + ')');
    console.log('up = (' + 
		self.camera.up.x + ', ' +
		self.camera.up.y + ', ' +
		self.camera.up.z + ')');
    console.log('ren.setCamera(new THREE.Vector3(' +
		self.center.x + ', ' +
		self.center.y + ', ' +
		self.center.z + '), ' +
		self.nearPlane + ', ' +
		self.farPlane + ', ' +
		'new THREE.Vector3(' +
		self.camera.position.x + ', ' +
		self.camera.position.y + ', ' +
		self.camera.position.z + '));\n' +
                'ren.setHome(new THREE.Vector3(' +
		self.cameraControls.object.position.x + ', ' +
		self.cameraControls.object.position.y + ', ' +
		self.cameraControls.object.position.z + '), ' +
		'new THREE.Vector3(' +
		self.camera.up.x + ', ' +
		self.camera.up.y + ', ' +
		self.camera.up.z + '));');
    console.log('mousePos = (' + self.mousePos.x +
                          ', ' + self.mousePos.y + ')');
  }

  /*!
   * \class	MARenderer
   * \function	_pick
   * \brief	Performs picking and then dispatches a pick event.
   */
  this._pick = function() {
    var pos = this.mousePos;
    pos.y = pos.y + self.pickOfs;
    self.raycaster.setFromCamera(pos, self.camera);
    var isct = self.raycaster.intersectObjects(self.scene.children, false);
    if(isct.length > 0) {
      self.eventHandler.dispatchEvent({type: 'pick',
                                       hitlist: isct});
    }
  }

  /*!
   * \class 	MARenderer
   * \function	_trackMouse
   * \brief	Tracks the mouse position and resets the animation timer.
   * \param e		The event.
   */
  this._trackMouse = function(e) {
    self.mousePos.x =  ((e.clientX - self.conOffset.x) /
                        self.con.clientWidth) *  2 - 1;
    self.mousePos.y = -((e.clientY - self.conOffset.y) /
                        self.con.clientHeight) * 2 + 1;
    self.makeLive();
  }

  /*! \class	MARenderer
   * \function	_keyPressed
   * \brief	Handles keypress events.
   * \param e		The event.
   */
  this._keyPressed = function(e) {
    switch(e.charCode) {
      case 33: // ! Test code
	self._testCode();
        break;
      case 60: // < opacity down
	self.opacityIncrement(-0.1);
	break;
      case 62: // > opacity up
	self.opacityIncrement(0.1);
	break;
      case 63: // ?
	self._pick();
        break;
      case 67: // C
        self.setCamera();
	self.goHome();
	break;
      case 72: // H
	self.setHome();
        break;
      case 104: // h
	self.goHome();
        break;
      case 112: // p
	self.pointSizeIncrement(+0.1);
        break;
      case 113: // q
	self.pointSizeIncrement(-0.1);
        break;
      case 115: // s
	self._updateAllMesh({mode: MARenderMode.PHONG});
        break;
      case 119: // w
	self._updateAllMesh({mode: MARenderMode.WIREFRAME});
        break;
      default:
        break;
    }
    console.log('MARender: charCode = ' + e.charCode);
  }

  /*!
   * \class	MARenderer
   * \function	_windowResize
   * \brief	Handles window resize events.
   */
  this._windowResize = function() {
    self.setConOffset();
    self.camera.aspect = self.con.clientWidth / self.con.clientHeight;
    self.camera.updateProjectionMatrix();
    self.renderer.setSize(self.con.clientWidth, self.con.clientHeight);
    if(self.cameraControls) {
      self.cameraControls.handleResize();
    }
    self.makeLive();
  }

}
```
