## Supplementary Dataset 5 for "A 3D molecular atlas of the chick embryonic heart": MAVTKLoader.js

```
/*!*
* \file         MAVTKLoader.js
* \author       mrdoob, Bill Hill
* \date         June 2015
* \version      $Id$
* \brief	A VTK file loader for three.js. This is based on the VTK
* 		loader by mrdoob. Differences (so far) from the original
* 		are:
*               * Slightly more flexible parsing.
*               * Only triangulated surfaces and point sets are supported.
*               * Bounding box and sphere set.
*               Both triangulated surfaces and point sets use POINTS to define
*               vertex locations. Triangulated surfaces are defined using
*               POLYGONS. Point properties using POINT_DATA, SCALARS and
*               LOOKUP_TABLE. The POLYGONS, POINT_DATA, SCALARS and LOOKUP_TABLE
*               are optional.
*/


THREE.VTKLoader = function(manager) {
  this.manager = (manager !== undefined)?
    manager: THREE.DefaultLoadingManager;
};

THREE.VTKLoader.prototype = {
constructor: THREE.VTKLoader,

	     load: function(url, onLoad, onProgress, onError) {
	       var scope = this;
	       var loader = new THREE.XHRLoader(scope.manager);
	       if((loader.crossOrigin !== undefined) &&
	          (loader.setCrossOrigin !== undefined)) {
		 loader.setCrossOrigin(this.crossOrigin);
	       }
	       var req = loader.load(url,
		   function (text) {
		   onLoad(scope.parse(text));
		   }, onProgress, onError);
	       return(req);
	     },

parse: function(data) {
	 var idx, pattern, reg_index, result;
	 var line = 0, n_points = 0, n_polys = 0; n_dpp = 0;
	 var point_type;
	 var geometry = new THREE.BufferGeometry();
	 var vertices = [];
	 var indices = [];
	 var colors = [];
	 var opacities = [];
	 var sizes = [];
	 var lut = [];
	 /*
	  * Expects to find a legacy format vtk file, such as:
	  *
	  * # vtk DataFile Version 1.0
	  * Some comment text
	  * ASCII
	  * DATASET POLYDATA
	  * POINTS 4 float
	  * 0 0 0
	  * 100 0 0
	  * 50 87 0
	  * 50 43 87
	  * POLYGONS 4 16
	  * 3 0 1 2
	  * 3 0 1 3
	  * 3 1 2 3
	  * 3 2 0 3
	  * POINT_DATA 4
	  * SCALARS name char
	  * LOOKUP_TABLE default
	  *
	  * Here both the POLYGONS and POINT_DATA sections
	  * are optional.
	  */
	 // # vtk DataFile Version <uint>.<uint>
	 pattern = /#[\s]+vtk[\s]+DataFile[\s]+Version[\s]+\d+\.\d+/gi;
	 result = pattern.exec(data);
	 if(result) {
	   // Some comment text
	   ++line;
	   // ASCII
	   ++line;
	   reg_index = pattern.lastIndex;
	   pattern = /ASCII/g;
	   pattern.lastIndex = reg_index;
	   result = pattern.exec(data);
	 }
	 if(result) {
	   // DATASET POLYDATA
	   ++line;
	   reg_index = pattern.lastIndex;
	   pattern = /DATASET[\s]+POLYDATA/g;
	   pattern.lastIndex = reg_index;
	   result = pattern.exec(data);
	 }
	 if(result) {
	   // POINTS 4 float
	   ++line;
	   pattern = /POINTS[\s]+([1-9]+[\d]*)[\s]+float/g;
	   pattern.lastIndex = reg_index;
	   if((result = pattern.exec(data)) !== null) {
	     n_points = parseInt(result[1]);
	   }
	 }
	 if(n_points > 0) {
	   // <float> <float> <float>
	   reg_index = pattern.lastIndex;
	   pattern = /(([+-]?[\d]+[.]?[\d\+\-eE]*)[\s]+([+-]?[\d]+[.]?[\d\+\-eE]*)[\s]+([+-]?[\d]+[.]?[\d\+\-eE]*))/g;
	   pattern.lastIndex = reg_index;
	   for(idx = 0; idx < n_points; ++idx) {
	     ++line;
	     result = pattern.exec(data);
	     if(!result) {
	       break;
	     }
	     vertices.push(
	         parseFloat(result[2]),
		 parseFloat(result[3]),
		 parseFloat(result[4]));
	   }
	 }
	 if((n_points > 0) && result) {
	   // POLYGONS <uint> <uint> or LINES <uint> <uint>
	   ++line;
	   pattern = /(POLYGONS)[\s]+([1-9]+[\d]*)[\s]+([1-9]+[\d]*)/g;
	   if((result = pattern.exec(data)) !== null) {
	     if(result[1] === 'POLYGONS') {
	       n_polys = parseInt(result[2]);
	       var n_poly_points = parseInt(result[3]);
	       n_dpp = n_poly_points / n_polys;
	     } else {
	       result = null;
	     }

	   } else {
	     result = true; // Allow just points, no polygons or lines
	   }
	 }
	 if((n_points > 0) && result) {
	   if(n_polys > 0) {
	     if(n_dpp === 4) {
	       // 3 <uint> <uint> <uint>
	       var reg_index = pattern.lastIndex;
	       pattern = /3[\s]+([\d]+)[\s]+([\d]+)[\s]+([\d]+)/g;
	       pattern.lastIndex = reg_index;
	       for(idx = 0; idx < n_polys; ++idx) {
		 ++line;
		 if((result = pattern.exec(data)) === null) {
		   break;
		 }
		 indices.push(
		     parseInt(result[1]),
		     parseInt(result[2]),
		     parseInt(result[3]));
	       }
	     } else if(n_dpp === 5) {
	       // 4 <uint> <uint> <uint> <uint>
	       var reg_index = pattern.lastIndex;
	       pattern = /4[\s]+([\d]+)[\s]+([\d]+)[\s]+([\d]+)[\s]+([\d]+)/g;
	       pattern.lastIndex = reg_index;
	       for(idx = 0; idx < n_polys; ++idx) {
		 ++line;
		 if((result = pattern.exec(data)) === null) {
		   break;
		 }
		 indices.push(
		     parseInt(result[1]),
		     parseInt(result[2]),
		     parseInt(result[4]));
		 indices.push(
		     parseInt(result[2]),
		     parseInt(result[3]),
		     parseInt(result[4]));
	       }
	     }
	   }
	   else {
	     // POINT_DATA <uint>
	     ++line;
	     pattern = /(POINT_DATA)[\s]+([1-9]+[\d]*)/g;
	     if((result = pattern.exec(data)) !== null) {
	       if((result[1] === 'POINT_DATA') && (Number(result[2]) == n_points)) {
		 // SCALARS name <type>
		 ++line;
		 var reg_index = pattern.lastIndex;
		 pattern = /(SCALARS)[\s]+([\S]+)[\s]+([\S]+)/g;
		 pattern.lastIndex = reg_index;
		 if(((result = pattern.exec(data)) === null) ||
		     (result[1] !== 'SCALARS') ||
		     ((result[3] !== 'char') && (result[3] !== 'float'))) {
		   result = null;
		 } else {
		   point_type = (result[3] === 'char')? 'C': 'F';
		   // LOOKUP_TABLE default
		   pattern = /(LOOKUP_TABLE)[\s]+(default)/g;
		   pattern.lastIndex = reg_index;
		   if(((result = pattern.exec(data)) === null) ||
		       (result[1] !== 'LOOKUP_TABLE') ||
		       (result[2] !== 'default')) {
		     result = null;
		   } else {
		     // <uint>|<float>
		     pattern = /([+-]?[\d]+[.]?[\d\+\-eE]*)/g;
		     pattern.lastIndex = reg_index;
		     for(idx = 0; idx < n_points; ++idx) {
		       ++line;
		       if((result = pattern.exec(data)) === null) {
			 break;
		       } else {
			 var a = (point_type === 'C')?
			   parseInt(result[1]) / 255.0:
			   parseFloat(result[1]);
			 if(a < 0.0) {
			   a = 0.0;
			 } else if(a > 1.0) {
			   a = 1.0;
			 }
			 colors.push(a, a, a);
			 var aa = ((a * a) + 0.1) / 1.1;
			 opacities.push(a);
			 sizes.push(aa);
		       }
		     }
		   }
		 }
	       } else {
		 result = null;
	       }

	     } else {
	       result = true; // Allow points with no point data
	     }
	   }
	 }
	 if(result === null) {
	   geometry = null;
	 } else {
	   geometry.addAttribute('position',
	                         new THREE.BufferAttribute(
				 new Float32Array(vertices), 3));
	   if(indices.length > 0) {
	     geometry.setIndex(
	         new THREE.BufferAttribute(
		 new Uint32Array(indices), 1));
	     geometry.computeVertexNormals();
	   }
	   if(colors.length > 0) {
	     geometry.addAttribute('colors',
	                           new THREE.BufferAttribute(
				   new Float32Array(colors), 3));
	     geometry.addAttribute('opacities',
	                           new THREE.BufferAttribute(
				   new Float32Array(opacities), 1));
	     geometry.addAttribute('sizes',
	                           new THREE.BufferAttribute(
				   new Float32Array(sizes), 1));
	     geometry.attributes.colors.needsUpdate = true;
	   }
	   geometry.computeBoundingBox();
	   geometry.computeBoundingSphere();
	 }
	 return geometry;
       }
}
```
