## Supplementary Dataset 5 for "A 3D molecular atlas of the chick embryonic heart": OBJLoader.js

```
/**
 * @author mrdoob / http://mrdoob.com/
 */

THREE.OBJLoader = function ( manager ) {

	this.manager = ( manager !== undefined ) ? manager : THREE.DefaultLoadingManager;

};

THREE.OBJLoader.prototype = {

	constructor: THREE.OBJLoader,

	load: function ( url, onLoad, onProgress, onError ) {

		var scope = this;

		var loader = new THREE.XHRLoader( scope.manager );
		loader.setCrossOrigin( this.crossOrigin );
		loader.load( url, function ( text ) {

			onLoad( scope.parse( text ) );

		}, onProgress, onError );

	},

	parse: function ( text ) {

		console.time( 'OBJLoader' );

		var object, objects = [];
		var geometry, material;

		function parseVertexIndex( value ) {

			var index = parseInt( value );

			return ( index >= 0 ? index - 1 : index + vertices.length / 3 ) * 3;

		}

		function parseNormalIndex( value ) {

			var index = parseInt( value );

			return ( index >= 0 ? index - 1 : index + normals.length / 3 ) * 3;

		}

		function parseUVIndex( value ) {

			var index = parseInt( value );

			return ( index >= 0 ? index - 1 : index + uvs.length / 2 ) * 2;

		}

		function addVertex( a, b, c ) {

			geometry.vertices.push(
				vertices[ a ], vertices[ a + 1 ], vertices[ a + 2 ],
				vertices[ b ], vertices[ b + 1 ], vertices[ b + 2 ],
				vertices[ c ], vertices[ c + 1 ], vertices[ c + 2 ]
			);

		}

		function addNormal( a, b, c ) {

			geometry.normals.push(
				normals[ a ], normals[ a + 1 ], normals[ a + 2 ],
				normals[ b ], normals[ b + 1 ], normals[ b + 2 ],
				normals[ c ], normals[ c + 1 ], normals[ c + 2 ]
			);

		}

		function addUV( a, b, c ) {

			geometry.uvs.push(
				uvs[ a ], uvs[ a + 1 ],
				uvs[ b ], uvs[ b + 1 ],
				uvs[ c ], uvs[ c + 1 ]
			);

		}

		function addFace( a, b, c, d,  ua, ub, uc, ud, na, nb, nc, nd ) {

			var ia = parseVertexIndex( a );
			var ib = parseVertexIndex( b );
			var ic = parseVertexIndex( c );
			var id;

			if ( d === undefined ) {

				addVertex( ia, ib, ic );

			} else {

				id = parseVertexIndex( d );

				addVertex( ia, ib, id );
				addVertex( ib, ic, id );

			}

			if ( ua !== undefined ) {

				ia = parseUVIndex( ua );
				ib = parseUVIndex( ub );
				ic = parseUVIndex( uc );

				if ( d === undefined ) {

					addUV( ia, ib, ic );

				} else {

					id = parseUVIndex( ud );

					addUV( ia, ib, id );
					addUV( ib, ic, id );

				}

			}

			if ( na !== undefined ) {

				ia = parseNormalIndex( na );
				ib = parseNormalIndex( nb );
				ic = parseNormalIndex( nc );

				if ( d === undefined ) {

					addNormal( ia, ib, ic );

				} else {

					id = parseNormalIndex( nd );

					addNormal( ia, ib, id );
					addNormal( ib, ic, id );

				}

			}

		}

		// create mesh if no objects in text

		if ( /^o /gm.test( text ) === false ) {

			geometry = {
				vertices: [],
				normals: [],
				uvs: []
			};

			material = {
				name: ''
			};

			object = {
				name: '',
				geometry: geometry,
				material: material
			};

			objects.push( object );

		}

		var vertices = [];
		var normals = [];
		var uvs = [];

		// v float float float

		var vertex_pattern = /v( +[\d|\.|\+|\-|e|E]+)( +[\d|\.|\+|\-|e|E]+)( +[\d|\.|\+|\-|e|E]+)/;

		// vn float float float

		var normal_pattern = /vn( +[\d|\.|\+|\-|e|E]+)( +[\d|\.|\+|\-|e|E]+)( +[\d|\.|\+|\-|e|E]+)/;

		// vt float float

		var uv_pattern = /vt( +[\d|\.|\+|\-|e|E]+)( +[\d|\.|\+|\-|e|E]+)/;

		// f vertex vertex vertex ...

		var face_pattern1 = /f( +-?\d+)( +-?\d+)( +-?\d+)( +-?\d+)?/;

		// f vertex/uv vertex/uv vertex/uv ...

		var face_pattern2 = /f( +(-?\d+)\/(-?\d+))( +(-?\d+)\/(-?\d+))( +(-?\d+)\/(-?\d+))( +(-?\d+)\/(-?\d+))?/;

		// f vertex/uv/normal vertex/uv/normal vertex/uv/normal ...

		var face_pattern3 = /f( +(-?\d+)\/(-?\d+)\/(-?\d+))( +(-?\d+)\/(-?\d+)\/(-?\d+))( +(-?\d+)\/(-?\d+)\/(-?\d+))( +(-?\d+)\/(-?\d+)\/(-?\d+))?/;

		// f vertex//normal vertex//normal vertex//normal ...

		var face_pattern4 = /f( +(-?\d+)\/\/(-?\d+))( +(-?\d+)\/\/(-?\d+))( +(-?\d+)\/\/(-?\d+))( +(-?\d+)\/\/(-?\d+))?/

		//

		var lines = text.split( '\n' );

		for ( var i = 0; i < lines.length; i ++ ) {

			var line = lines[ i ];
			line = line.trim();

			var result;

			if ( line.length === 0 || line.charAt( 0 ) === '#' ) {

				continue;

			} else if ( ( result = vertex_pattern.exec( line ) ) !== null ) {

				// ["v 1.0 2.0 3.0", "1.0", "2.0", "3.0"]

				vertices.push(
					parseFloat( result[ 1 ] ),
					parseFloat( result[ 2 ] ),
					parseFloat( result[ 3 ] )
				);

			} else if ( ( result = normal_pattern.exec( line ) ) !== null ) {

				// ["vn 1.0 2.0 3.0", "1.0", "2.0", "3.0"]

				normals.push(
					parseFloat( result[ 1 ] ),
					parseFloat( result[ 2 ] ),
					parseFloat( result[ 3 ] )
				);

			} else if ( ( result = uv_pattern.exec( line ) ) !== null ) {

				// ["vt 0.1 0.2", "0.1", "0.2"]

				uvs.push(
					parseFloat( result[ 1 ] ),
					parseFloat( result[ 2 ] )
				);

			} else if ( ( result = face_pattern1.exec( line ) ) !== null ) {

				// ["f 1 2 3", "1", "2", "3", undefined]

				addFace(
					result[ 1 ], result[ 2 ], result[ 3 ], result[ 4 ]
				);

			} else if ( ( result = face_pattern2.exec( line ) ) !== null ) {

				// ["f 1/1 2/2 3/3", " 1/1", "1", "1", " 2/2", "2", "2", " 3/3", "3", "3", undefined, undefined, undefined]

				addFace(
					result[ 2 ], result[ 5 ], result[ 8 ], result[ 11 ],
					result[ 3 ], result[ 6 ], result[ 9 ], result[ 12 ]
				);

			} else if ( ( result = face_pattern3.exec( line ) ) !== null ) {

				// ["f 1/1/1 2/2/2 3/3/3", " 1/1/1", "1", "1", "1", " 2/2/2", "2", "2", "2", " 3/3/3", "3", "3", "3", undefined, undefined, undefined, undefined]

				addFace(
					result[ 2 ], result[ 6 ], result[ 10 ], result[ 14 ],
					result[ 3 ], result[ 7 ], result[ 11 ], result[ 15 ],
					result[ 4 ], result[ 8 ], result[ 12 ], result[ 16 ]
				);

			} else if ( ( result = face_pattern4.exec( line ) ) !== null ) {

				// ["f 1//1 2//2 3//3", " 1//1", "1", "1", " 2//2", "2", "2", " 3//3", "3", "3", undefined, undefined, undefined]

				addFace(
					result[ 2 ], result[ 5 ], result[ 8 ], result[ 11 ],
					undefined, undefined, undefined, undefined,
					result[ 3 ], result[ 6 ], result[ 9 ], result[ 12 ]
				);

			} else if ( /^o /.test( line ) ) {

				geometry = {
					vertices: [],
					normals: [],
					uvs: []
				};

				material = {
					name: ''
				};

				object = {
					name: line.substring( 2 ).trim(),
					geometry: geometry,
					material: material
				};

				objects.push( object )

			} else if ( /^g /.test( line ) ) {

				// group

			} else if ( /^usemtl /.test( line ) ) {

				// material

				material.name = line.substring( 7 ).trim();

			} else if ( /^mtllib /.test( line ) ) {

				// mtl file

			} else if ( /^s /.test( line ) ) {

				// smooth shading

			} else {

				// console.log( "THREE.OBJLoader: Unhandled line " + line );

			}

		}

		var container = new THREE.Object3D();

		for ( var i = 0, l = objects.length; i < l; i ++ ) {

			object = objects[ i ];
			geometry = object.geometry;

			var buffergeometry = new THREE.BufferGeometry();

			buffergeometry.addAttribute( 'position', new THREE.BufferAttribute( new Float32Array( geometry.vertices ), 3 ) );

			if ( geometry.normals.length > 0 ) {
				buffergeometry.addAttribute( 'normal', new THREE.BufferAttribute( new Float32Array( geometry.normals ), 3 ) );
			}

			if ( geometry.uvs.length > 0 ) {
				buffergeometry.addAttribute( 'uv', new THREE.BufferAttribute( new Float32Array( geometry.uvs ), 2 ) );
			}

			material = new THREE.MeshLambertMaterial();
			material.name = object.material.name;

			var mesh = new THREE.Mesh( buffergeometry, material );
			mesh.name = object.name;

			container.add( mesh );

		}

		console.timeEnd( 'OBJLoader' );

		return container;

	}

};
```
