## Supplementary Dataset 5 for "A 3D molecular atlas of the chick embryonic heart": STLLoader.js

```
/**
 * @author aleeper / http://adamleeper.com/
 * @author mrdoob / http://mrdoob.com/
 * @author gero3 / https://github.com/gero3
 *
 * Description: A THREE loader for STL ASCII files, as created by Solidworks and other CAD programs.
 *
 * Supports both binary and ASCII encoded files, with automatic detection of type.
 *
 * Limitations:
 *  Binary decoding supports "Magics" color format (http://en.wikipedia.org/wiki/STL_(file_format)#Color_in_binary_STL).
 *  There is perhaps some question as to how valid it is to always assume little-endian-ness.
 *  ASCII decoding assumes file is UTF-8. Seems to work for the examples...
 *
 * Usage:
 *  var loader = new THREE.STLLoader();
 *  loader.load( './models/stl/slotted_disk.stl', function ( geometry ) {
 *    scene.add( new THREE.Mesh( geometry ) );
 *  });
 *
 * For binary STLs geometry might contain colors for vertices. To use it:
 *  // use the same code to load STL as above
 *  if (geometry.hasColors) {
 *    material = new THREE.MeshPhongMaterial({ opacity: geometry.alpha, vertexColors: THREE.VertexColors });
 *  } else { .... }
 *  var mesh = new THREE.Mesh( geometry, material );
 */


THREE.STLLoader = function ( manager ) {

	this.manager = ( manager !== undefined ) ? manager : THREE.DefaultLoadingManager;

};

THREE.STLLoader.prototype = {

	constructor: THREE.STLLoader,

	load: function ( url, onLoad, onProgress, onError ) {

		var scope = this;

		var loader = new THREE.XHRLoader( scope.manager );
		loader.setCrossOrigin( this.crossOrigin );
		loader.setResponseType('arraybuffer');
		loader.load( url, function ( text ) {

			onLoad( scope.parse( text ) );

		}, onProgress, onError );

	},
	
	parse: function ( data ) {

		var isBinary = function () {

			var expect, face_size, n_faces, reader;
			reader = new DataView( binData );
			face_size = (32 / 8 * 3) + ((32 / 8 * 3) * 3) + (16 / 8);
			n_faces = reader.getUint32(80, true);
			expect = 80 + (32 / 8) + (n_faces * face_size);
			
			if ( expect === reader.byteLength ) {
				
				return true;
				
			}

			// some binary files will have different size from expected,
			// checking characters higher than ASCII to confirm is binary
			var fileLength = reader.byteLength;
			for ( var index = 0; index < fileLength; index ++ ) {

				if ( reader.getUint8(index, false) > 127 ) {
					
					return true;
					
				}

			}

			return false;

		};

		var binData = this.ensureBinary( data );

		return isBinary()
			? this.parseBinary( binData )
			: this.parseASCII( this.ensureString( data ) );

	},

	parseBinary: function ( data ) {

		var reader = new DataView( data );
		var faces = reader.getUint32( 80, true );

		var r, g, b, hasColors = false, colors;
		var defaultR, defaultG, defaultB, alpha;

		// process STL header
		// check for default color in header ("COLOR=rgba" sequence).

		for ( var index = 0; index < 80 - 10; index ++ ) {

			if ((reader.getUint32(index, false) == 0x434F4C4F /*COLO*/) &&
				(reader.getUint8(index + 4) == 0x52 /*'R'*/) &&
				(reader.getUint8(index + 5) == 0x3D /*'='*/)) {

				hasColors = true;
				colors = new Float32Array( faces * 3 * 3);

				defaultR = reader.getUint8(index + 6) / 255;
				defaultG = reader.getUint8(index + 7) / 255;
				defaultB = reader.getUint8(index + 8) / 255;
				alpha = reader.getUint8(index + 9) / 255;
			}
		}

		var dataOffset = 84;
		var faceLength = 12 * 4 + 2;

		var offset = 0;

		var geometry = new THREE.BufferGeometry();

		var vertices = new Float32Array( faces * 3 * 3 );
		var normals = new Float32Array( faces * 3 * 3 );

		for ( var face = 0; face < faces; face ++ ) {

			var start = dataOffset + face * faceLength;
			var normalX = reader.getFloat32(start, true);
			var normalY = reader.getFloat32(start + 4, true);
			var normalZ = reader.getFloat32(start + 8, true);

			if (hasColors) {

				var packedColor = reader.getUint16(start + 48, true);

				if ((packedColor & 0x8000) === 0) { // facet has its own unique color

					r = (packedColor & 0x1F) / 31;
					g = ((packedColor >> 5) & 0x1F) / 31;
					b = ((packedColor >> 10) & 0x1F) / 31;
				} else {

					r = defaultR;
					g = defaultG;
					b = defaultB;
				}
			}

			for ( var i = 1; i <= 3; i ++ ) {

				var vertexstart = start + i * 12;

				vertices[ offset     ] = reader.getFloat32( vertexstart, true );
				vertices[ offset + 1 ] = reader.getFloat32( vertexstart + 4, true );
				vertices[ offset + 2 ] = reader.getFloat32( vertexstart + 8, true );

				normals[ offset     ] = normalX;
				normals[ offset + 1 ] = normalY;
				normals[ offset + 2 ] = normalZ;

				if (hasColors) {
					colors[ offset     ] = r;
					colors[ offset + 1 ] = g;
					colors[ offset + 2 ] = b;
				}

				offset += 3;

			}

		}

		geometry.addAttribute( 'position', new THREE.BufferAttribute( vertices, 3 ) );
		geometry.addAttribute( 'normal', new THREE.BufferAttribute( normals, 3 ) );

		if (hasColors) {
			geometry.addAttribute( 'color', new THREE.BufferAttribute( colors, 3 ) );
			geometry.hasColors = true;
			geometry.alpha = alpha;
		}

		return geometry;

	},

	parseASCII: function ( data ) {

		var geometry, length, normal, patternFace, patternNormal, patternVertex, result, text;
		geometry = new THREE.Geometry();
		patternFace = /facet([\s\S]*?)endfacet/g;

		while ( ( result = patternFace.exec( data ) ) !== null ) {

			text = result[0];
			patternNormal = /normal[\s]+([\-+]?[0-9]+\.?[0-9]*([eE][\-+]?[0-9]+)?)+[\s]+([\-+]?[0-9]*\.?[0-9]+([eE][\-+]?[0-9]+)?)+[\s]+([\-+]?[0-9]*\.?[0-9]+([eE][\-+]?[0-9]+)?)+/g;

			while ( ( result = patternNormal.exec( text ) ) !== null ) {

				normal = new THREE.Vector3( parseFloat( result[ 1 ] ), parseFloat( result[ 3 ] ), parseFloat( result[ 5 ] ) );

			}

			patternVertex = /vertex[\s]+([\-+]?[0-9]+\.?[0-9]*([eE][\-+]?[0-9]+)?)+[\s]+([\-+]?[0-9]*\.?[0-9]+([eE][\-+]?[0-9]+)?)+[\s]+([\-+]?[0-9]*\.?[0-9]+([eE][\-+]?[0-9]+)?)+/g;

			while ( ( result = patternVertex.exec( text ) ) !== null ) {

				geometry.vertices.push( new THREE.Vector3( parseFloat( result[ 1 ] ), parseFloat( result[ 3 ] ), parseFloat( result[ 5 ] ) ) );

			}

			length = geometry.vertices.length;

			geometry.faces.push( new THREE.Face3( length - 3, length - 2, length - 1, normal ) );

		}

		geometry.computeBoundingBox();
		geometry.computeBoundingSphere();

		return geometry;

	},

	ensureString: function ( buf ) {

		if (typeof buf !== "string") {
			var array_buffer = new Uint8Array(buf);
			var str = '';
			for (var i = 0; i < buf.byteLength; i ++) {
				str += String.fromCharCode(array_buffer[i]); // implicitly assumes little-endian
			}
			return str;
		} else {
			return buf;
		}

	},

	ensureBinary: function ( buf ) {

		if (typeof buf === "string") {
			var array_buffer = new Uint8Array(buf.length);
			for (var i = 0; i < buf.length; i ++) {
				array_buffer[i] = buf.charCodeAt(i) & 0xff; // implicitly assumes little-endian
			}
			return array_buffer.buffer || array_buffer;
		} else {
			return buf;
		}
		
	}

};

if ( typeof DataView === 'undefined') {

	DataView = function(buffer, byteOffset, byteLength) {

		this.buffer = buffer;
		this.byteOffset = byteOffset || 0;
		this.byteLength = byteLength || buffer.byteLength || buffer.length;
		this._isString = typeof buffer === "string";

	}

	DataView.prototype = {

		_getCharCodes:function(buffer,start,length) {
			start = start || 0;
			length = length || buffer.length;
			var end = start + length;
			var codes = [];
			for (var i = start; i < end; i ++) {
				codes.push(buffer.charCodeAt(i) & 0xff);
			}
			return codes;
		},

		_getBytes: function (length, byteOffset, littleEndian) {

			var result;

			// Handle the lack of endianness
			if (littleEndian === undefined) {

				littleEndian = this._littleEndian;

			}

			// Handle the lack of byteOffset
			if (byteOffset === undefined) {

				byteOffset = this.byteOffset;

			} else {

				byteOffset = this.byteOffset + byteOffset;

			}

			if (length === undefined) {

				length = this.byteLength - byteOffset;

			}

			// Error Checking
			if (typeof byteOffset !== 'number') {

				throw new TypeError('DataView byteOffset is not a number');

			}

			if (length < 0 || byteOffset + length > this.byteLength) {

				throw new Error('DataView length or (byteOffset+length) value is out of bounds');

			}

			if (this.isString) {

				result = this._getCharCodes(this.buffer, byteOffset, byteOffset + length);

			} else {

				result = this.buffer.slice(byteOffset, byteOffset + length);

			}

			if (!littleEndian && length > 1) {

				if (!(result instanceof Array)) {

					result = Array.prototype.slice.call(result);

				}

				result.reverse();
			}

			return result;

		},

		// Compatibility functions on a String Buffer

		getFloat64: function (byteOffset, littleEndian) {

			var b = this._getBytes(8, byteOffset, littleEndian),

				sign = 1 - (2 * (b[7] >> 7)),
				exponent = ((((b[7] << 1) & 0xff) << 3) | (b[6] >> 4)) - ((1 << 10) - 1),

			// Binary operators such as | and << operate on 32 bit values, using + and Math.pow(2) instead
				mantissa = ((b[6] & 0x0f) * Math.pow(2, 48)) + (b[5] * Math.pow(2, 40)) + (b[4] * Math.pow(2, 32)) +
							(b[3] * Math.pow(2, 24)) + (b[2] * Math.pow(2, 16)) + (b[1] * Math.pow(2, 8)) + b[0];

			if (exponent === 1024) {
				if (mantissa !== 0) {
					return NaN;
				} else {
					return sign * Infinity;
				}
			}

			if (exponent === -1023) { // Denormalized
				return sign * mantissa * Math.pow(2, -1022 - 52);
			}

			return sign * (1 + mantissa * Math.pow(2, -52)) * Math.pow(2, exponent);

		},

		getFloat32: function (byteOffset, littleEndian) {

			var b = this._getBytes(4, byteOffset, littleEndian),

				sign = 1 - (2 * (b[3] >> 7)),
				exponent = (((b[3] << 1) & 0xff) | (b[2] >> 7)) - 127,
				mantissa = ((b[2] & 0x7f) << 16) | (b[1] << 8) | b[0];

			if (exponent === 128) {
				if (mantissa !== 0) {
					return NaN;
				} else {
					return sign * Infinity;
				}
			}

			if (exponent === -127) { // Denormalized
				return sign * mantissa * Math.pow(2, -126 - 23);
			}

			return sign * (1 + mantissa * Math.pow(2, -23)) * Math.pow(2, exponent);
		},

		getInt32: function (byteOffset, littleEndian) {
			var b = this._getBytes(4, byteOffset, littleEndian);
			return (b[3] << 24) | (b[2] << 16) | (b[1] << 8) | b[0];
		},

		getUint32: function (byteOffset, littleEndian) {
			return this.getInt32(byteOffset, littleEndian) >>> 0;
		},

		getInt16: function (byteOffset, littleEndian) {
			return (this.getUint16(byteOffset, littleEndian) << 16) >> 16;
		},

		getUint16: function (byteOffset, littleEndian) {
			var b = this._getBytes(2, byteOffset, littleEndian);
			return (b[1] << 8) | b[0];
		},

		getInt8: function (byteOffset) {
			return (this.getUint8(byteOffset) << 24) >> 24;
		},

		getUint8: function (byteOffset) {
			return this._getBytes(1, byteOffset)[0];
		}

	 };

}
```
