## Supplementary Dataset 5 for "A 3D molecular atlas of the chick embryonic heart": three.js

```
// File:src/Three.js

/**
 * @author mrdoob / http://mrdoob.com/
 */

var THREE = { REVISION: '77' };

//

if ( typeof define === 'function' && define.amd ) {

	define( 'three', THREE );

} else if ( 'undefined' !== typeof exports && 'undefined' !== typeof module ) {

	module.exports = THREE;

}

// Polyfills

if ( Number.EPSILON === undefined ) {

	Number.EPSILON = Math.pow( 2, - 52 );

}

//

if ( Math.sign === undefined ) {

	// https://developer.mozilla.org/en-US/docs/Web/JavaScript/Reference/Global_Objects/Math/sign

	Math.sign = function ( x ) {

		return ( x < 0 ) ? - 1 : ( x > 0 ) ? 1 : + x;

	};

}

if ( Function.prototype.name === undefined ) {

	// Missing in IE9-11.
	// https://developer.mozilla.org/en-US/docs/Web/JavaScript/Reference/Global_Objects/Function/name

	Object.defineProperty( Function.prototype, 'name', {

		get: function () {

			return this.toString().match( /^\s*function\s*(\S*)\s*\(/ )[ 1 ];

		}

	} );

}

if ( Object.assign === undefined ) {

	// Missing in IE.
	// https://developer.mozilla.org/en-US/docs/Web/JavaScript/Reference/Global_Objects/Object/assign

	( function () {

		Object.assign = function ( target ) {

			'use strict';

			if ( target === undefined || target === null ) {

				throw new TypeError( 'Cannot convert undefined or null to object' );

			}

			var output = Object( target );

			for ( var index = 1; index < arguments.length; index ++ ) {

				var source = arguments[ index ];

				if ( source !== undefined && source !== null ) {

					for ( var nextKey in source ) {

						if ( Object.prototype.hasOwnProperty.call( source, nextKey ) ) {

							output[ nextKey ] = source[ nextKey ];

						}

					}

				}

			}

			return output;

		};

	} )();

}

//

Object.assign( THREE, {

	// https://developer.mozilla.org/en-US/docs/Web/API/MouseEvent.button

	MOUSE: { LEFT: 0, MIDDLE: 1, RIGHT: 2 },

	// GL STATE CONSTANTS

	CullFaceNone: 0,
	CullFaceBack: 1,
	CullFaceFront: 2,
	CullFaceFrontBack: 3,

	FrontFaceDirectionCW: 0,
	FrontFaceDirectionCCW: 1,

	// SHADOWING TYPES

	BasicShadowMap: 0,
	PCFShadowMap: 1,
	PCFSoftShadowMap: 2,

	// MATERIAL CONSTANTS

	// side

	FrontSide: 0,
	BackSide: 1,
	DoubleSide: 2,

	// shading

	FlatShading: 1,
	SmoothShading: 2,

	// colors

	NoColors: 0,
	FaceColors: 1,
	VertexColors: 2,

	// blending modes

	NoBlending: 0,
	NormalBlending: 1,
	AdditiveBlending: 2,
	SubtractiveBlending: 3,
	MultiplyBlending: 4,
	CustomBlending: 5,

	// custom blending equations
	// (numbers start from 100 not to clash with other
	// mappings to OpenGL constants defined in Texture.js)

	AddEquation: 100,
	SubtractEquation: 101,
	ReverseSubtractEquation: 102,
	MinEquation: 103,
	MaxEquation: 104,

	// custom blending destination factors

	ZeroFactor: 200,
	OneFactor: 201,
	SrcColorFactor: 202,
	OneMinusSrcColorFactor: 203,
	SrcAlphaFactor: 204,
	OneMinusSrcAlphaFactor: 205,
	DstAlphaFactor: 206,
	OneMinusDstAlphaFactor: 207,

	// custom blending source factors

	//ZeroFactor: 200,
	//OneFactor: 201,
	//SrcAlphaFactor: 204,
	//OneMinusSrcAlphaFactor: 205,
	//DstAlphaFactor: 206,
	//OneMinusDstAlphaFactor: 207,
	DstColorFactor: 208,
	OneMinusDstColorFactor: 209,
	SrcAlphaSaturateFactor: 210,

	// depth modes

	NeverDepth: 0,
	AlwaysDepth: 1,
	LessDepth: 2,
	LessEqualDepth: 3,
	EqualDepth: 4,
	GreaterEqualDepth: 5,
	GreaterDepth: 6,
	NotEqualDepth: 7,


	// TEXTURE CONSTANTS

	MultiplyOperation: 0,
	MixOperation: 1,
	AddOperation: 2,

	// Tone Mapping modes

	NoToneMapping: 0, // do not do any tone mapping, not even exposure (required for special purpose passes.)
	LinearToneMapping: 1, // only apply exposure.
	ReinhardToneMapping: 2,
	Uncharted2ToneMapping: 3, // John Hable
	CineonToneMapping: 4, // optimized filmic operator by Jim Hejl and Richard Burgess-Dawson

	// Mapping modes

	UVMapping: 300,

	CubeReflectionMapping: 301,
	CubeRefractionMapping: 302,

	EquirectangularReflectionMapping: 303,
	EquirectangularRefractionMapping: 304,

	SphericalReflectionMapping: 305,
	CubeUVReflectionMapping: 306,
	CubeUVRefractionMapping: 307,

	// Wrapping modes

	RepeatWrapping: 1000,
	ClampToEdgeWrapping: 1001,
	MirroredRepeatWrapping: 1002,

	// Filters

	NearestFilter: 1003,
	NearestMipMapNearestFilter: 1004,
	NearestMipMapLinearFilter: 1005,
	LinearFilter: 1006,
	LinearMipMapNearestFilter: 1007,
	LinearMipMapLinearFilter: 1008,

	// Data types

	UnsignedByteType: 1009,
	ByteType: 1010,
	ShortType: 1011,
	UnsignedShortType: 1012,
	IntType: 1013,
	UnsignedIntType: 1014,
	FloatType: 1015,
	HalfFloatType: 1025,

	// Pixel types

	//UnsignedByteType: 1009,
	UnsignedShort4444Type: 1016,
	UnsignedShort5551Type: 1017,
	UnsignedShort565Type: 1018,

	// Pixel formats

	AlphaFormat: 1019,
	RGBFormat: 1020,
	RGBAFormat: 1021,
	LuminanceFormat: 1022,
	LuminanceAlphaFormat: 1023,
	// THREE.RGBEFormat handled as THREE.RGBAFormat in shaders
	RGBEFormat: THREE.RGBAFormat, //1024;
	DepthFormat: 1026,

	// DDS / ST3C Compressed texture formats

	RGB_S3TC_DXT1_Format: 2001,
	RGBA_S3TC_DXT1_Format: 2002,
	RGBA_S3TC_DXT3_Format: 2003,
	RGBA_S3TC_DXT5_Format: 2004,

	// PVRTC compressed texture formats

	RGB_PVRTC_4BPPV1_Format: 2100,
	RGB_PVRTC_2BPPV1_Format: 2101,
	RGBA_PVRTC_4BPPV1_Format: 2102,
	RGBA_PVRTC_2BPPV1_Format: 2103,

	// ETC compressed texture formats

	RGB_ETC1_Format: 2151,

	// Loop styles for AnimationAction

	LoopOnce: 2200,
	LoopRepeat: 2201,
	LoopPingPong: 2202,

	// Interpolation

	InterpolateDiscrete: 2300,
	InterpolateLinear: 2301,
	InterpolateSmooth: 2302,

	// Interpolant ending modes

	ZeroCurvatureEnding: 2400,
	ZeroSlopeEnding: 2401,
	WrapAroundEnding: 2402,

	// Triangle Draw modes

	TrianglesDrawMode: 0,
	TriangleStripDrawMode: 1,
	TriangleFanDrawMode: 2,

	// Texture Encodings

	LinearEncoding: 3000, // No encoding at all.
	sRGBEncoding: 3001,
	GammaEncoding: 3007, // uses GAMMA_FACTOR, for backwards compatibility with WebGLRenderer.gammaInput/gammaOutput

	// The following Texture Encodings are for RGB-only (no alpha) HDR light emission sources.
	// These encodings should not specified as output encodings except in rare situations.
	RGBEEncoding: 3002, // AKA Radiance.
	LogLuvEncoding: 3003,
	RGBM7Encoding: 3004,
	RGBM16Encoding: 3005,
	RGBDEncoding: 3006, // MaxRange is 256.

	// Depth packing strategies

	BasicDepthPacking: 3200, // for writing to float textures for high precision or for visualizing results in RGB buffers
	RGBADepthPacking: 3201 // for packing into RGBA buffers.

} );

// File:src/math/Color.js

/**
 * @author mrdoob / http://mrdoob.com/
 */

THREE.Color = function ( r, g, b ) {

	if ( g === undefined && b === undefined ) {

		// r is THREE.Color, hex or string
		return this.set( r );

	}

	return this.setRGB( r, g, b );

};

THREE.Color.prototype = {

	constructor: THREE.Color,

	r: 1, g: 1, b: 1,

	set: function ( value ) {

		if ( value instanceof THREE.Color ) {

			this.copy( value );

		} else if ( typeof value === 'number' ) {

			this.setHex( value );

		} else if ( typeof value === 'string' ) {

			this.setStyle( value );

		}

		return this;

	},

	setScalar: function ( scalar ) {

		this.r = scalar;
		this.g = scalar;
		this.b = scalar;

	},

	setHex: function ( hex ) {

		hex = Math.floor( hex );

		this.r = ( hex >> 16 & 255 ) / 255;
		this.g = ( hex >> 8 & 255 ) / 255;
		this.b = ( hex & 255 ) / 255;

		return this;

	},

	setRGB: function ( r, g, b ) {

		this.r = r;
		this.g = g;
		this.b = b;

		return this;

	},

	setHSL: function () {

		function hue2rgb( p, q, t ) {

			if ( t < 0 ) t += 1;
			if ( t > 1 ) t -= 1;
			if ( t < 1 / 6 ) return p + ( q - p ) * 6 * t;
			if ( t < 1 / 2 ) return q;
			if ( t < 2 / 3 ) return p + ( q - p ) * 6 * ( 2 / 3 - t );
			return p;

		}

		return function ( h, s, l ) {

			// h,s,l ranges are in 0.0 - 1.0
			h = THREE.Math.euclideanModulo( h, 1 );
			s = THREE.Math.clamp( s, 0, 1 );
			l = THREE.Math.clamp( l, 0, 1 );

			if ( s === 0 ) {

				this.r = this.g = this.b = l;

			} else {

				var p = l <= 0.5 ? l * ( 1 + s ) : l + s - ( l * s );
				var q = ( 2 * l ) - p;

				this.r = hue2rgb( q, p, h + 1 / 3 );
				this.g = hue2rgb( q, p, h );
				this.b = hue2rgb( q, p, h - 1 / 3 );

			}

			return this;

		};

	}(),

	setStyle: function ( style ) {

		function handleAlpha( string ) {

			if ( string === undefined ) return;

			if ( parseFloat( string ) < 1 ) {

				console.warn( 'THREE.Color: Alpha component of ' + style + ' will be ignored.' );

			}

		}


		var m;

		if ( m = /^((?:rgb|hsl)a?)\(\s*([^\)]*)\)/.exec( style ) ) {

			// rgb / hsl

			var color;
			var name = m[ 1 ];
			var components = m[ 2 ];

			switch ( name ) {

				case 'rgb':
				case 'rgba':

					if ( color = /^(\d+)\s*,\s*(\d+)\s*,\s*(\d+)\s*(,\s*([0-9]*\.?[0-9]+)\s*)?$/.exec( components ) ) {

						// rgb(255,0,0) rgba(255,0,0,0.5)
						this.r = Math.min( 255, parseInt( color[ 1 ], 10 ) ) / 255;
						this.g = Math.min( 255, parseInt( color[ 2 ], 10 ) ) / 255;
						this.b = Math.min( 255, parseInt( color[ 3 ], 10 ) ) / 255;

						handleAlpha( color[ 5 ] );

						return this;

					}

					if ( color = /^(\d+)\%\s*,\s*(\d+)\%\s*,\s*(\d+)\%\s*(,\s*([0-9]*\.?[0-9]+)\s*)?$/.exec( components ) ) {

						// rgb(100%,0%,0%) rgba(100%,0%,0%,0.5)
						this.r = Math.min( 100, parseInt( color[ 1 ], 10 ) ) / 100;
						this.g = Math.min( 100, parseInt( color[ 2 ], 10 ) ) / 100;
						this.b = Math.min( 100, parseInt( color[ 3 ], 10 ) ) / 100;

						handleAlpha( color[ 5 ] );

						return this;

					}

					break;

				case 'hsl':
				case 'hsla':

					if ( color = /^([0-9]*\.?[0-9]+)\s*,\s*(\d+)\%\s*,\s*(\d+)\%\s*(,\s*([0-9]*\.?[0-9]+)\s*)?$/.exec( components ) ) {

						// hsl(120,50%,50%) hsla(120,50%,50%,0.5)
						var h = parseFloat( color[ 1 ] ) / 360;
						var s = parseInt( color[ 2 ], 10 ) / 100;
						var l = parseInt( color[ 3 ], 10 ) / 100;

						handleAlpha( color[ 5 ] );

						return this.setHSL( h, s, l );

					}

					break;

			}

		} else if ( m = /^\#([A-Fa-f0-9]+)$/.exec( style ) ) {

			// hex color

			var hex = m[ 1 ];
			var size = hex.length;

			if ( size === 3 ) {

				// #ff0
				this.r = parseInt( hex.charAt( 0 ) + hex.charAt( 0 ), 16 ) / 255;
				this.g = parseInt( hex.charAt( 1 ) + hex.charAt( 1 ), 16 ) / 255;
				this.b = parseInt( hex.charAt( 2 ) + hex.charAt( 2 ), 16 ) / 255;

				return this;

			} else if ( size === 6 ) {

				// #ff0000
				this.r = parseInt( hex.charAt( 0 ) + hex.charAt( 1 ), 16 ) / 255;
				this.g = parseInt( hex.charAt( 2 ) + hex.charAt( 3 ), 16 ) / 255;
				this.b = parseInt( hex.charAt( 4 ) + hex.charAt( 5 ), 16 ) / 255;

				return this;

			}

		}

		if ( style && style.length > 0 ) {

			// color keywords
			var hex = THREE.ColorKeywords[ style ];

			if ( hex !== undefined ) {

				// red
				this.setHex( hex );

			} else {

				// unknown color
				console.warn( 'THREE.Color: Unknown color ' + style );

			}

		}

		return this;

	},

	clone: function () {

		return new this.constructor( this.r, this.g, this.b );

	},

	copy: function ( color ) {

		this.r = color.r;
		this.g = color.g;
		this.b = color.b;

		return this;

	},

	copyGammaToLinear: function ( color, gammaFactor ) {

		if ( gammaFactor === undefined ) gammaFactor = 2.0;

		this.r = Math.pow( color.r, gammaFactor );
		this.g = Math.pow( color.g, gammaFactor );
		this.b = Math.pow( color.b, gammaFactor );

		return this;

	},

	copyLinearToGamma: function ( color, gammaFactor ) {

		if ( gammaFactor === undefined ) gammaFactor = 2.0;

		var safeInverse = ( gammaFactor > 0 ) ? ( 1.0 / gammaFactor ) : 1.0;

		this.r = Math.pow( color.r, safeInverse );
		this.g = Math.pow( color.g, safeInverse );
		this.b = Math.pow( color.b, safeInverse );

		return this;

	},

	convertGammaToLinear: function () {

		var r = this.r, g = this.g, b = this.b;

		this.r = r * r;
		this.g = g * g;
		this.b = b * b;

		return this;

	},

	convertLinearToGamma: function () {

		this.r = Math.sqrt( this.r );
		this.g = Math.sqrt( this.g );
		this.b = Math.sqrt( this.b );

		return this;

	},

	getHex: function () {

		return ( this.r * 255 ) << 16 ^ ( this.g * 255 ) << 8 ^ ( this.b * 255 ) << 0;

	},

	getHexString: function () {

		return ( '000000' + this.getHex().toString( 16 ) ).slice( - 6 );

	},

	getHSL: function ( optionalTarget ) {

		// h,s,l ranges are in 0.0 - 1.0

		var hsl = optionalTarget || { h: 0, s: 0, l: 0 };

		var r = this.r, g = this.g, b = this.b;

		var max = Math.max( r, g, b );
		var min = Math.min( r, g, b );

		var hue, saturation;
		var lightness = ( min + max ) / 2.0;

		if ( min === max ) {

			hue = 0;
			saturation = 0;

		} else {

			var delta = max - min;

			saturation = lightness <= 0.5 ? delta / ( max + min ) : delta / ( 2 - max - min );

			switch ( max ) {

				case r: hue = ( g - b ) / delta + ( g < b ? 6 : 0 ); break;
				case g: hue = ( b - r ) / delta + 2; break;
				case b: hue = ( r - g ) / delta + 4; break;

			}

			hue /= 6;

		}

		hsl.h = hue;
		hsl.s = saturation;
		hsl.l = lightness;

		return hsl;

	},

	getStyle: function () {

		return 'rgb(' + ( ( this.r * 255 ) | 0 ) + ',' + ( ( this.g * 255 ) | 0 ) + ',' + ( ( this.b * 255 ) | 0 ) + ')';

	},

	offsetHSL: function ( h, s, l ) {

		var hsl = this.getHSL();

		hsl.h += h; hsl.s += s; hsl.l += l;

		this.setHSL( hsl.h, hsl.s, hsl.l );

		return this;

	},

	add: function ( color ) {

		this.r += color.r;
		this.g += color.g;
		this.b += color.b;

		return this;

	},

	addColors: function ( color1, color2 ) {

		this.r = color1.r + color2.r;
		this.g = color1.g + color2.g;
		this.b = color1.b + color2.b;

		return this;

	},

	addScalar: function ( s ) {

		this.r += s;
		this.g += s;
		this.b += s;

		return this;

	},

	multiply: function ( color ) {

		this.r *= color.r;
		this.g *= color.g;
		this.b *= color.b;

		return this;

	},

	multiplyScalar: function ( s ) {

		this.r *= s;
		this.g *= s;
		this.b *= s;

		return this;

	},

	lerp: function ( color, alpha ) {

		this.r += ( color.r - this.r ) * alpha;
		this.g += ( color.g - this.g ) * alpha;
		this.b += ( color.b - this.b ) * alpha;

		return this;

	},

	equals: function ( c ) {

		return ( c.r === this.r ) && ( c.g === this.g ) && ( c.b === this.b );

	},

	fromArray: function ( array, offset ) {

		if ( offset === undefined ) offset = 0;

		this.r = array[ offset ];
		this.g = array[ offset + 1 ];
		this.b = array[ offset + 2 ];

		return this;

	},

	toArray: function ( array, offset ) {

		if ( array === undefined ) array = [];
		if ( offset === undefined ) offset = 0;

		array[ offset ] = this.r;
		array[ offset + 1 ] = this.g;
		array[ offset + 2 ] = this.b;

		return array;

	}

};

THREE.ColorKeywords = { 'aliceblue': 0xF0F8FF, 'antiquewhite': 0xFAEBD7, 'aqua': 0x00FFFF, 'aquamarine': 0x7FFFD4, 'azure': 0xF0FFFF,
'beige': 0xF5F5DC, 'bisque': 0xFFE4C4, 'black': 0x000000, 'blanchedalmond': 0xFFEBCD, 'blue': 0x0000FF, 'blueviolet': 0x8A2BE2,
'brown': 0xA52A2A, 'burlywood': 0xDEB887, 'cadetblue': 0x5F9EA0, 'chartreuse': 0x7FFF00, 'chocolate': 0xD2691E, 'coral': 0xFF7F50,
'cornflowerblue': 0x6495ED, 'cornsilk': 0xFFF8DC, 'crimson': 0xDC143C, 'cyan': 0x00FFFF, 'darkblue': 0x00008B, 'darkcyan': 0x008B8B,
'darkgoldenrod': 0xB8860B, 'darkgray': 0xA9A9A9, 'darkgreen': 0x006400, 'darkgrey': 0xA9A9A9, 'darkkhaki': 0xBDB76B, 'darkmagenta': 0x8B008B,
'darkolivegreen': 0x556B2F, 'darkorange': 0xFF8C00, 'darkorchid': 0x9932CC, 'darkred': 0x8B0000, 'darksalmon': 0xE9967A, 'darkseagreen': 0x8FBC8F,
'darkslateblue': 0x483D8B, 'darkslategray': 0x2F4F4F, 'darkslategrey': 0x2F4F4F, 'darkturquoise': 0x00CED1, 'darkviolet': 0x9400D3,
'deeppink': 0xFF1493, 'deepskyblue': 0x00BFFF, 'dimgray': 0x696969, 'dimgrey': 0x696969, 'dodgerblue': 0x1E90FF, 'firebrick': 0xB22222,
'floralwhite': 0xFFFAF0, 'forestgreen': 0x228B22, 'fuchsia': 0xFF00FF, 'gainsboro': 0xDCDCDC, 'ghostwhite': 0xF8F8FF, 'gold': 0xFFD700,
'goldenrod': 0xDAA520, 'gray': 0x808080, 'green': 0x008000, 'greenyellow': 0xADFF2F, 'grey': 0x808080, 'honeydew': 0xF0FFF0, 'hotpink': 0xFF69B4,
'indianred': 0xCD5C5C, 'indigo': 0x4B0082, 'ivory': 0xFFFFF0, 'khaki': 0xF0E68C, 'lavender': 0xE6E6FA, 'lavenderblush': 0xFFF0F5, 'lawngreen': 0x7CFC00,
'lemonchiffon': 0xFFFACD, 'lightblue': 0xADD8E6, 'lightcoral': 0xF08080, 'lightcyan': 0xE0FFFF, 'lightgoldenrodyellow': 0xFAFAD2, 'lightgray': 0xD3D3D3,
'lightgreen': 0x90EE90, 'lightgrey': 0xD3D3D3, 'lightpink': 0xFFB6C1, 'lightsalmon': 0xFFA07A, 'lightseagreen': 0x20B2AA, 'lightskyblue': 0x87CEFA,
'lightslategray': 0x778899, 'lightslategrey': 0x778899, 'lightsteelblue': 0xB0C4DE, 'lightyellow': 0xFFFFE0, 'lime': 0x00FF00, 'limegreen': 0x32CD32,
'linen': 0xFAF0E6, 'magenta': 0xFF00FF, 'maroon': 0x800000, 'mediumaquamarine': 0x66CDAA, 'mediumblue': 0x0000CD, 'mediumorchid': 0xBA55D3,
'mediumpurple': 0x9370DB, 'mediumseagreen': 0x3CB371, 'mediumslateblue': 0x7B68EE, 'mediumspringgreen': 0x00FA9A, 'mediumturquoise': 0x48D1CC,
'mediumvioletred': 0xC71585, 'midnightblue': 0x191970, 'mintcream': 0xF5FFFA, 'mistyrose': 0xFFE4E1, 'moccasin': 0xFFE4B5, 'navajowhite': 0xFFDEAD,
'navy': 0x000080, 'oldlace': 0xFDF5E6, 'olive': 0x808000, 'olivedrab': 0x6B8E23, 'orange': 0xFFA500, 'orangered': 0xFF4500, 'orchid': 0xDA70D6,
'palegoldenrod': 0xEEE8AA, 'palegreen': 0x98FB98, 'paleturquoise': 0xAFEEEE, 'palevioletred': 0xDB7093, 'papayawhip': 0xFFEFD5, 'peachpuff': 0xFFDAB9,
'peru': 0xCD853F, 'pink': 0xFFC0CB, 'plum': 0xDDA0DD, 'powderblue': 0xB0E0E6, 'purple': 0x800080, 'red': 0xFF0000, 'rosybrown': 0xBC8F8F,
'royalblue': 0x4169E1, 'saddlebrown': 0x8B4513, 'salmon': 0xFA8072, 'sandybrown': 0xF4A460, 'seagreen': 0x2E8B57, 'seashell': 0xFFF5EE,
'sienna': 0xA0522D, 'silver': 0xC0C0C0, 'skyblue': 0x87CEEB, 'slateblue': 0x6A5ACD, 'slategray': 0x708090, 'slategrey': 0x708090, 'snow': 0xFFFAFA,
'springgreen': 0x00FF7F, 'steelblue': 0x4682B4, 'tan': 0xD2B48C, 'teal': 0x008080, 'thistle': 0xD8BFD8, 'tomato': 0xFF6347, 'turquoise': 0x40E0D0,
'violet': 0xEE82EE, 'wheat': 0xF5DEB3, 'white': 0xFFFFFF, 'whitesmoke': 0xF5F5F5, 'yellow': 0xFFFF00, 'yellowgreen': 0x9ACD32 };

// File:src/math/Quaternion.js

/**
 * @author mikael emtinger / http://gomo.se/
 * @author alteredq / http://alteredqualia.com/
 * @author WestLangley / http://github.com/WestLangley
 * @author bhouston / http://clara.io
 */

THREE.Quaternion = function ( x, y, z, w ) {

	this._x = x || 0;
	this._y = y || 0;
	this._z = z || 0;
	this._w = ( w !== undefined ) ? w : 1;

};

THREE.Quaternion.prototype = {

	constructor: THREE.Quaternion,

	get x () {

		return this._x;

	},

	set x ( value ) {

		this._x = value;
		this.onChangeCallback();

	},

	get y () {

		return this._y;

	},

	set y ( value ) {

		this._y = value;
		this.onChangeCallback();

	},

	get z () {

		return this._z;

	},

	set z ( value ) {

		this._z = value;
		this.onChangeCallback();

	},

	get w () {

		return this._w;

	},

	set w ( value ) {

		this._w = value;
		this.onChangeCallback();

	},

	set: function ( x, y, z, w ) {

		this._x = x;
		this._y = y;
		this._z = z;
		this._w = w;

		this.onChangeCallback();

		return this;

	},

	clone: function () {

		return new this.constructor( this._x, this._y, this._z, this._w );

	},

	copy: function ( quaternion ) {

		this._x = quaternion.x;
		this._y = quaternion.y;
		this._z = quaternion.z;
		this._w = quaternion.w;

		this.onChangeCallback();

		return this;

	},

	setFromEuler: function ( euler, update ) {

		if ( euler instanceof THREE.Euler === false ) {

			throw new Error( 'THREE.Quaternion: .setFromEuler() now expects a Euler rotation rather than a Vector3 and order.' );

		}

		// http://www.mathworks.com/matlabcentral/fileexchange/
		// 	20696-function-to-convert-between-dcm-euler-angles-quaternions-and-euler-vectors/
		//	content/SpinCalc.m

		var c1 = Math.cos( euler._x / 2 );
		var c2 = Math.cos( euler._y / 2 );
		var c3 = Math.cos( euler._z / 2 );
		var s1 = Math.sin( euler._x / 2 );
		var s2 = Math.sin( euler._y / 2 );
		var s3 = Math.sin( euler._z / 2 );

		var order = euler.order;

		if ( order === 'XYZ' ) {

			this._x = s1 * c2 * c3 + c1 * s2 * s3;
			this._y = c1 * s2 * c3 - s1 * c2 * s3;
			this._z = c1 * c2 * s3 + s1 * s2 * c3;
			this._w = c1 * c2 * c3 - s1 * s2 * s3;

		} else if ( order === 'YXZ' ) {

			this._x = s1 * c2 * c3 + c1 * s2 * s3;
			this._y = c1 * s2 * c3 - s1 * c2 * s3;
			this._z = c1 * c2 * s3 - s1 * s2 * c3;
			this._w = c1 * c2 * c3 + s1 * s2 * s3;

		} else if ( order === 'ZXY' ) {

			this._x = s1 * c2 * c3 - c1 * s2 * s3;
			this._y = c1 * s2 * c3 + s1 * c2 * s3;
			this._z = c1 * c2 * s3 + s1 * s2 * c3;
			this._w = c1 * c2 * c3 - s1 * s2 * s3;

		} else if ( order === 'ZYX' ) {

			this._x = s1 * c2 * c3 - c1 * s2 * s3;
			this._y = c1 * s2 * c3 + s1 * c2 * s3;
			this._z = c1 * c2 * s3 - s1 * s2 * c3;
			this._w = c1 * c2 * c3 + s1 * s2 * s3;

		} else if ( order === 'YZX' ) {

			this._x = s1 * c2 * c3 + c1 * s2 * s3;
			this._y = c1 * s2 * c3 + s1 * c2 * s3;
			this._z = c1 * c2 * s3 - s1 * s2 * c3;
			this._w = c1 * c2 * c3 - s1 * s2 * s3;

		} else if ( order === 'XZY' ) {

			this._x = s1 * c2 * c3 - c1 * s2 * s3;
			this._y = c1 * s2 * c3 - s1 * c2 * s3;
			this._z = c1 * c2 * s3 + s1 * s2 * c3;
			this._w = c1 * c2 * c3 + s1 * s2 * s3;

		}

		if ( update !== false ) this.onChangeCallback();

		return this;

	},

	setFromAxisAngle: function ( axis, angle ) {

		// http://www.euclideanspace.com/maths/geometry/rotations/conversions/angleToQuaternion/index.htm

		// assumes axis is normalized

		var halfAngle = angle / 2, s = Math.sin( halfAngle );

		this._x = axis.x * s;
		this._y = axis.y * s;
		this._z = axis.z * s;
		this._w = Math.cos( halfAngle );

		this.onChangeCallback();

		return this;

	},

	setFromRotationMatrix: function ( m ) {

		// http://www.euclideanspace.com/maths/geometry/rotations/conversions/matrixToQuaternion/index.htm

		// assumes the upper 3x3 of m is a pure rotation matrix (i.e, unscaled)

		var te = m.elements,

			m11 = te[ 0 ], m12 = te[ 4 ], m13 = te[ 8 ],
			m21 = te[ 1 ], m22 = te[ 5 ], m23 = te[ 9 ],
			m31 = te[ 2 ], m32 = te[ 6 ], m33 = te[ 10 ],

			trace = m11 + m22 + m33,
			s;

		if ( trace > 0 ) {

			s = 0.5 / Math.sqrt( trace + 1.0 );

			this._w = 0.25 / s;
			this._x = ( m32 - m23 ) * s;
			this._y = ( m13 - m31 ) * s;
			this._z = ( m21 - m12 ) * s;

		} else if ( m11 > m22 && m11 > m33 ) {

			s = 2.0 * Math.sqrt( 1.0 + m11 - m22 - m33 );

			this._w = ( m32 - m23 ) / s;
			this._x = 0.25 * s;
			this._y = ( m12 + m21 ) / s;
			this._z = ( m13 + m31 ) / s;

		} else if ( m22 > m33 ) {

			s = 2.0 * Math.sqrt( 1.0 + m22 - m11 - m33 );

			this._w = ( m13 - m31 ) / s;
			this._x = ( m12 + m21 ) / s;
			this._y = 0.25 * s;
			this._z = ( m23 + m32 ) / s;

		} else {

			s = 2.0 * Math.sqrt( 1.0 + m33 - m11 - m22 );

			this._w = ( m21 - m12 ) / s;
			this._x = ( m13 + m31 ) / s;
			this._y = ( m23 + m32 ) / s;
			this._z = 0.25 * s;

		}

		this.onChangeCallback();

		return this;

	},

	setFromUnitVectors: function () {

		// http://lolengine.net/blog/2014/02/24/quaternion-from-two-vectors-final

		// assumes direction vectors vFrom and vTo are normalized

		var v1, r;

		var EPS = 0.000001;

		return function ( vFrom, vTo ) {

			if ( v1 === undefined ) v1 = new THREE.Vector3();

			r = vFrom.dot( vTo ) + 1;

			if ( r < EPS ) {

				r = 0;

				if ( Math.abs( vFrom.x ) > Math.abs( vFrom.z ) ) {

					v1.set( - vFrom.y, vFrom.x, 0 );

				} else {

					v1.set( 0, - vFrom.z, vFrom.y );

				}

			} else {

				v1.crossVectors( vFrom, vTo );

			}

			this._x = v1.x;
			this._y = v1.y;
			this._z = v1.z;
			this._w = r;

			return this.normalize();

		};

	}(),

	inverse: function () {

		return this.conjugate().normalize();

	},

	conjugate: function () {

		this._x *= - 1;
		this._y *= - 1;
		this._z *= - 1;

		this.onChangeCallback();

		return this;

	},

	dot: function ( v ) {

		return this._x * v._x + this._y * v._y + this._z * v._z + this._w * v._w;

	},

	lengthSq: function () {

		return this._x * this._x + this._y * this._y + this._z * this._z + this._w * this._w;

	},

	length: function () {

		return Math.sqrt( this._x * this._x + this._y * this._y + this._z * this._z + this._w * this._w );

	},

	normalize: function () {

		var l = this.length();

		if ( l === 0 ) {

			this._x = 0;
			this._y = 0;
			this._z = 0;
			this._w = 1;

		} else {

			l = 1 / l;

			this._x = this._x * l;
			this._y = this._y * l;
			this._z = this._z * l;
			this._w = this._w * l;

		}

		this.onChangeCallback();

		return this;

	},

	multiply: function ( q, p ) {

		if ( p !== undefined ) {

			console.warn( 'THREE.Quaternion: .multiply() now only accepts one argument. Use .multiplyQuaternions( a, b ) instead.' );
			return this.multiplyQuaternions( q, p );

		}

		return this.multiplyQuaternions( this, q );

	},

	premultiply: function ( q ) {

		return this.multiplyQuaternions( q, this );

	},

	multiplyQuaternions: function ( a, b ) {

		// from http://www.euclideanspace.com/maths/algebra/realNormedAlgebra/quaternions/code/index.htm

		var qax = a._x, qay = a._y, qaz = a._z, qaw = a._w;
		var qbx = b._x, qby = b._y, qbz = b._z, qbw = b._w;

		this._x = qax * qbw + qaw * qbx + qay * qbz - qaz * qby;
		this._y = qay * qbw + qaw * qby + qaz * qbx - qax * qbz;
		this._z = qaz * qbw + qaw * qbz + qax * qby - qay * qbx;
		this._w = qaw * qbw - qax * qbx - qay * qby - qaz * qbz;

		this.onChangeCallback();

		return this;

	},

	slerp: function ( qb, t ) {

		if ( t === 0 ) return this;
		if ( t === 1 ) return this.copy( qb );

		var x = this._x, y = this._y, z = this._z, w = this._w;

		// http://www.euclideanspace.com/maths/algebra/realNormedAlgebra/quaternions/slerp/

		var cosHalfTheta = w * qb._w + x * qb._x + y * qb._y + z * qb._z;

		if ( cosHalfTheta < 0 ) {

			this._w = - qb._w;
			this._x = - qb._x;
			this._y = - qb._y;
			this._z = - qb._z;

			cosHalfTheta = - cosHalfTheta;

		} else {

			this.copy( qb );

		}

		if ( cosHalfTheta >= 1.0 ) {

			this._w = w;
			this._x = x;
			this._y = y;
			this._z = z;

			return this;

		}

		var sinHalfTheta = Math.sqrt( 1.0 - cosHalfTheta * cosHalfTheta );

		if ( Math.abs( sinHalfTheta ) < 0.001 ) {

			this._w = 0.5 * ( w + this._w );
			this._x = 0.5 * ( x + this._x );
			this._y = 0.5 * ( y + this._y );
			this._z = 0.5 * ( z + this._z );

			return this;

		}

		var halfTheta = Math.atan2( sinHalfTheta, cosHalfTheta );
		var ratioA = Math.sin( ( 1 - t ) * halfTheta ) / sinHalfTheta,
		ratioB = Math.sin( t * halfTheta ) / sinHalfTheta;

		this._w = ( w * ratioA + this._w * ratioB );
		this._x = ( x * ratioA + this._x * ratioB );
		this._y = ( y * ratioA + this._y * ratioB );
		this._z = ( z * ratioA + this._z * ratioB );

		this.onChangeCallback();

		return this;

	},

	equals: function ( quaternion ) {

		return ( quaternion._x === this._x ) && ( quaternion._y === this._y ) && ( quaternion._z === this._z ) && ( quaternion._w === this._w );

	},

	fromArray: function ( array, offset ) {

		if ( offset === undefined ) offset = 0;

		this._x = array[ offset ];
		this._y = array[ offset + 1 ];
		this._z = array[ offset + 2 ];
		this._w = array[ offset + 3 ];

		this.onChangeCallback();

		return this;

	},

	toArray: function ( array, offset ) {

		if ( array === undefined ) array = [];
		if ( offset === undefined ) offset = 0;

		array[ offset ] = this._x;
		array[ offset + 1 ] = this._y;
		array[ offset + 2 ] = this._z;
		array[ offset + 3 ] = this._w;

		return array;

	},

	onChange: function ( callback ) {

		this.onChangeCallback = callback;

		return this;

	},

	onChangeCallback: function () {}

};

Object.assign( THREE.Quaternion, {

	slerp: function( qa, qb, qm, t ) {

		return qm.copy( qa ).slerp( qb, t );

	},

	slerpFlat: function(
			dst, dstOffset, src0, srcOffset0, src1, srcOffset1, t ) {

		// fuzz-free, array-based Quaternion SLERP operation

		var x0 = src0[ srcOffset0 + 0 ],
			y0 = src0[ srcOffset0 + 1 ],
			z0 = src0[ srcOffset0 + 2 ],
			w0 = src0[ srcOffset0 + 3 ],

			x1 = src1[ srcOffset1 + 0 ],
			y1 = src1[ srcOffset1 + 1 ],
			z1 = src1[ srcOffset1 + 2 ],
			w1 = src1[ srcOffset1 + 3 ];

		if ( w0 !== w1 || x0 !== x1 || y0 !== y1 || z0 !== z1 ) {

			var s = 1 - t,

				cos = x0 * x1 + y0 * y1 + z0 * z1 + w0 * w1,

				dir = ( cos >= 0 ? 1 : - 1 ),
				sqrSin = 1 - cos * cos;

			// Skip the Slerp for tiny steps to avoid numeric problems:
			if ( sqrSin > Number.EPSILON ) {

				var sin = Math.sqrt( sqrSin ),
					len = Math.atan2( sin, cos * dir );

				s = Math.sin( s * len ) / sin;
				t = Math.sin( t * len ) / sin;

			}

			var tDir = t * dir;

			x0 = x0 * s + x1 * tDir;
			y0 = y0 * s + y1 * tDir;
			z0 = z0 * s + z1 * tDir;
			w0 = w0 * s + w1 * tDir;

			// Normalize in case we just did a lerp:
			if ( s === 1 - t ) {

				var f = 1 / Math.sqrt( x0 * x0 + y0 * y0 + z0 * z0 + w0 * w0 );

				x0 *= f;
				y0 *= f;
				z0 *= f;
				w0 *= f;

			}

		}

		dst[ dstOffset ] = x0;
		dst[ dstOffset + 1 ] = y0;
		dst[ dstOffset + 2 ] = z0;
		dst[ dstOffset + 3 ] = w0;

	}

} );

// File:src/math/Vector2.js

/**
 * @author mrdoob / http://mrdoob.com/
 * @author philogb / http://blog.thejit.org/
 * @author egraether / http://egraether.com/
 * @author zz85 / http://www.lab4games.net/zz85/blog
 */

THREE.Vector2 = function ( x, y ) {

	this.x = x || 0;
	this.y = y || 0;

};

THREE.Vector2.prototype = {

	constructor: THREE.Vector2,

	get width() {

		return this.x;

	},

	set width( value ) {

		this.x = value;

	},

	get height() {

		return this.y;

	},

	set height( value ) {

		this.y = value;

	},

	//

	set: function ( x, y ) {

		this.x = x;
		this.y = y;

		return this;

	},

	setScalar: function ( scalar ) {

		this.x = scalar;
		this.y = scalar;

		return this;

	},

	setX: function ( x ) {

		this.x = x;

		return this;

	},

	setY: function ( y ) {

		this.y = y;

		return this;

	},

	setComponent: function ( index, value ) {

		switch ( index ) {

			case 0: this.x = value; break;
			case 1: this.y = value; break;
			default: throw new Error( 'index is out of range: ' + index );

		}

	},

	getComponent: function ( index ) {

		switch ( index ) {

			case 0: return this.x;
			case 1: return this.y;
			default: throw new Error( 'index is out of range: ' + index );

		}

	},

	clone: function () {

		return new this.constructor( this.x, this.y );

	},

	copy: function ( v ) {

		this.x = v.x;
		this.y = v.y;

		return this;

	},

	add: function ( v, w ) {

		if ( w !== undefined ) {

			console.warn( 'THREE.Vector2: .add() now only accepts one argument. Use .addVectors( a, b ) instead.' );
			return this.addVectors( v, w );

		}

		this.x += v.x;
		this.y += v.y;

		return this;

	},

	addScalar: function ( s ) {

		this.x += s;
		this.y += s;

		return this;

	},

	addVectors: function ( a, b ) {

		this.x = a.x + b.x;
		this.y = a.y + b.y;

		return this;

	},

	addScaledVector: function ( v, s ) {

		this.x += v.x * s;
		this.y += v.y * s;

		return this;

	},

	sub: function ( v, w ) {

		if ( w !== undefined ) {

			console.warn( 'THREE.Vector2: .sub() now only accepts one argument. Use .subVectors( a, b ) instead.' );
			return this.subVectors( v, w );

		}

		this.x -= v.x;
		this.y -= v.y;

		return this;

	},

	subScalar: function ( s ) {

		this.x -= s;
		this.y -= s;

		return this;

	},

	subVectors: function ( a, b ) {

		this.x = a.x - b.x;
		this.y = a.y - b.y;

		return this;

	},

	multiply: function ( v ) {

		this.x *= v.x;
		this.y *= v.y;

		return this;

	},

	multiplyScalar: function ( scalar ) {

		if ( isFinite( scalar ) ) {

			this.x *= scalar;
			this.y *= scalar;

		} else {

			this.x = 0;
			this.y = 0;

		}

		return this;

	},

	divide: function ( v ) {

		this.x /= v.x;
		this.y /= v.y;

		return this;

	},

	divideScalar: function ( scalar ) {

		return this.multiplyScalar( 1 / scalar );

	},

	min: function ( v ) {

		this.x = Math.min( this.x, v.x );
		this.y = Math.min( this.y, v.y );

		return this;

	},

	max: function ( v ) {

		this.x = Math.max( this.x, v.x );
		this.y = Math.max( this.y, v.y );

		return this;

	},

	clamp: function ( min, max ) {

		// This function assumes min < max, if this assumption isn't true it will not operate correctly

		this.x = Math.max( min.x, Math.min( max.x, this.x ) );
		this.y = Math.max( min.y, Math.min( max.y, this.y ) );

		return this;

	},

	clampScalar: function () {

		var min, max;

		return function clampScalar( minVal, maxVal ) {

			if ( min === undefined ) {

				min = new THREE.Vector2();
				max = new THREE.Vector2();

			}

			min.set( minVal, minVal );
			max.set( maxVal, maxVal );

			return this.clamp( min, max );

		};

	}(),

	clampLength: function ( min, max ) {

		var length = this.length();

		return this.multiplyScalar( Math.max( min, Math.min( max, length ) ) / length );

	},

	floor: function () {

		this.x = Math.floor( this.x );
		this.y = Math.floor( this.y );

		return this;

	},

	ceil: function () {

		this.x = Math.ceil( this.x );
		this.y = Math.ceil( this.y );

		return this;

	},

	round: function () {

		this.x = Math.round( this.x );
		this.y = Math.round( this.y );

		return this;

	},

	roundToZero: function () {

		this.x = ( this.x < 0 ) ? Math.ceil( this.x ) : Math.floor( this.x );
		this.y = ( this.y < 0 ) ? Math.ceil( this.y ) : Math.floor( this.y );

		return this;

	},

	negate: function () {

		this.x = - this.x;
		this.y = - this.y;

		return this;

	},

	dot: function ( v ) {

		return this.x * v.x + this.y * v.y;

	},

	lengthSq: function () {

		return this.x * this.x + this.y * this.y;

	},

	length: function () {

		return Math.sqrt( this.x * this.x + this.y * this.y );

	},

	lengthManhattan: function() {

		return Math.abs( this.x ) + Math.abs( this.y );

	},

	normalize: function () {

		return this.divideScalar( this.length() );

	},

	angle: function () {

		// computes the angle in radians with respect to the positive x-axis

		var angle = Math.atan2( this.y, this.x );

		if ( angle < 0 ) angle += 2 * Math.PI;

		return angle;

	},

	distanceTo: function ( v ) {

		return Math.sqrt( this.distanceToSquared( v ) );

	},

	distanceToSquared: function ( v ) {

		var dx = this.x - v.x, dy = this.y - v.y;
		return dx * dx + dy * dy;

	},

	setLength: function ( length ) {

		return this.multiplyScalar( length / this.length() );

	},

	lerp: function ( v, alpha ) {

		this.x += ( v.x - this.x ) * alpha;
		this.y += ( v.y - this.y ) * alpha;

		return this;

	},

	lerpVectors: function ( v1, v2, alpha ) {

		return this.subVectors( v2, v1 ).multiplyScalar( alpha ).add( v1 );

	},

	equals: function ( v ) {

		return ( ( v.x === this.x ) && ( v.y === this.y ) );

	},

	fromArray: function ( array, offset ) {

		if ( offset === undefined ) offset = 0;

		this.x = array[ offset ];
		this.y = array[ offset + 1 ];

		return this;

	},

	toArray: function ( array, offset ) {

		if ( array === undefined ) array = [];
		if ( offset === undefined ) offset = 0;

		array[ offset ] = this.x;
		array[ offset + 1 ] = this.y;

		return array;

	},

	fromAttribute: function ( attribute, index, offset ) {

		if ( offset === undefined ) offset = 0;

		index = index * attribute.itemSize + offset;

		this.x = attribute.array[ index ];
		this.y = attribute.array[ index + 1 ];

		return this;

	},

	rotateAround: function ( center, angle ) {

		var c = Math.cos( angle ), s = Math.sin( angle );

		var x = this.x - center.x;
		var y = this.y - center.y;

		this.x = x * c - y * s + center.x;
		this.y = x * s + y * c + center.y;

		return this;

	}

};

// File:src/math/Vector3.js

/**
 * @author mrdoob / http://mrdoob.com/
 * @author *kile / http://kile.stravaganza.org/
 * @author philogb / http://blog.thejit.org/
 * @author mikael emtinger / http://gomo.se/
 * @author egraether / http://egraether.com/
 * @author WestLangley / http://github.com/WestLangley
 */

THREE.Vector3 = function ( x, y, z ) {

	this.x = x || 0;
	this.y = y || 0;
	this.z = z || 0;

};

THREE.Vector3.prototype = {

	constructor: THREE.Vector3,

	set: function ( x, y, z ) {

		this.x = x;
		this.y = y;
		this.z = z;

		return this;

	},

	setScalar: function ( scalar ) {

		this.x = scalar;
		this.y = scalar;
		this.z = scalar;

		return this;

	},

	setX: function ( x ) {

		this.x = x;

		return this;

	},

	setY: function ( y ) {

		this.y = y;

		return this;

	},

	setZ: function ( z ) {

		this.z = z;

		return this;

	},

	setComponent: function ( index, value ) {

		switch ( index ) {

			case 0: this.x = value; break;
			case 1: this.y = value; break;
			case 2: this.z = value; break;
			default: throw new Error( 'index is out of range: ' + index );

		}

	},

	getComponent: function ( index ) {

		switch ( index ) {

			case 0: return this.x;
			case 1: return this.y;
			case 2: return this.z;
			default: throw new Error( 'index is out of range: ' + index );

		}

	},

	clone: function () {

		return new this.constructor( this.x, this.y, this.z );

	},

	copy: function ( v ) {

		this.x = v.x;
		this.y = v.y;
		this.z = v.z;

		return this;

	},

	add: function ( v, w ) {

		if ( w !== undefined ) {

			console.warn( 'THREE.Vector3: .add() now only accepts one argument. Use .addVectors( a, b ) instead.' );
			return this.addVectors( v, w );

		}

		this.x += v.x;
		this.y += v.y;
		this.z += v.z;

		return this;

	},

	addScalar: function ( s ) {

		this.x += s;
		this.y += s;
		this.z += s;

		return this;

	},

	addVectors: function ( a, b ) {

		this.x = a.x + b.x;
		this.y = a.y + b.y;
		this.z = a.z + b.z;

		return this;

	},

	addScaledVector: function ( v, s ) {

		this.x += v.x * s;
		this.y += v.y * s;
		this.z += v.z * s;

		return this;

	},

	sub: function ( v, w ) {

		if ( w !== undefined ) {

			console.warn( 'THREE.Vector3: .sub() now only accepts one argument. Use .subVectors( a, b ) instead.' );
			return this.subVectors( v, w );

		}

		this.x -= v.x;
		this.y -= v.y;
		this.z -= v.z;

		return this;

	},

	subScalar: function ( s ) {

		this.x -= s;
		this.y -= s;
		this.z -= s;

		return this;

	},

	subVectors: function ( a, b ) {

		this.x = a.x - b.x;
		this.y = a.y - b.y;
		this.z = a.z - b.z;

		return this;

	},

	multiply: function ( v, w ) {

		if ( w !== undefined ) {

			console.warn( 'THREE.Vector3: .multiply() now only accepts one argument. Use .multiplyVectors( a, b ) instead.' );
			return this.multiplyVectors( v, w );

		}

		this.x *= v.x;
		this.y *= v.y;
		this.z *= v.z;

		return this;

	},

	multiplyScalar: function ( scalar ) {

		if ( isFinite( scalar ) ) {

			this.x *= scalar;
			this.y *= scalar;
			this.z *= scalar;

		} else {

			this.x = 0;
			this.y = 0;
			this.z = 0;

		}

		return this;

	},

	multiplyVectors: function ( a, b ) {

		this.x = a.x * b.x;
		this.y = a.y * b.y;
		this.z = a.z * b.z;

		return this;

	},

	applyEuler: function () {

		var quaternion;

		return function applyEuler( euler ) {

			if ( euler instanceof THREE.Euler === false ) {

				console.error( 'THREE.Vector3: .applyEuler() now expects an Euler rotation rather than a Vector3 and order.' );

			}

			if ( quaternion === undefined ) quaternion = new THREE.Quaternion();

			return this.applyQuaternion( quaternion.setFromEuler( euler ) );

		};

	}(),

	applyAxisAngle: function () {

		var quaternion;

		return function applyAxisAngle( axis, angle ) {

			if ( quaternion === undefined ) quaternion = new THREE.Quaternion();

			return this.applyQuaternion( quaternion.setFromAxisAngle( axis, angle ) );

		};

	}(),

	applyMatrix3: function ( m ) {

		var x = this.x, y = this.y, z = this.z;
		var e = m.elements;

		this.x = e[ 0 ] * x + e[ 3 ] * y + e[ 6 ] * z;
		this.y = e[ 1 ] * x + e[ 4 ] * y + e[ 7 ] * z;
		this.z = e[ 2 ] * x + e[ 5 ] * y + e[ 8 ] * z;

		return this;

	},

	applyMatrix4: function ( m ) {

		// input: THREE.Matrix4 affine matrix

		var x = this.x, y = this.y, z = this.z;
		var e = m.elements;

		this.x = e[ 0 ] * x + e[ 4 ] * y + e[ 8 ]  * z + e[ 12 ];
		this.y = e[ 1 ] * x + e[ 5 ] * y + e[ 9 ]  * z + e[ 13 ];
		this.z = e[ 2 ] * x + e[ 6 ] * y + e[ 10 ] * z + e[ 14 ];

		return this;

	},

	applyProjection: function ( m ) {

		// input: THREE.Matrix4 projection matrix

		var x = this.x, y = this.y, z = this.z;
		var e = m.elements;
		var d = 1 / ( e[ 3 ] * x + e[ 7 ] * y + e[ 11 ] * z + e[ 15 ] ); // perspective divide

		this.x = ( e[ 0 ] * x + e[ 4 ] * y + e[ 8 ]  * z + e[ 12 ] ) * d;
		this.y = ( e[ 1 ] * x + e[ 5 ] * y + e[ 9 ]  * z + e[ 13 ] ) * d;
		this.z = ( e[ 2 ] * x + e[ 6 ] * y + e[ 10 ] * z + e[ 14 ] ) * d;

		return this;

	},

	applyQuaternion: function ( q ) {

		var x = this.x, y = this.y, z = this.z;
		var qx = q.x, qy = q.y, qz = q.z, qw = q.w;

		// calculate quat * vector

		var ix =  qw * x + qy * z - qz * y;
		var iy =  qw * y + qz * x - qx * z;
		var iz =  qw * z + qx * y - qy * x;
		var iw = - qx * x - qy * y - qz * z;

		// calculate result * inverse quat

		this.x = ix * qw + iw * - qx + iy * - qz - iz * - qy;
		this.y = iy * qw + iw * - qy + iz * - qx - ix * - qz;
		this.z = iz * qw + iw * - qz + ix * - qy - iy * - qx;

		return this;

	},

	project: function () {

		var matrix;

		return function project( camera ) {

			if ( matrix === undefined ) matrix = new THREE.Matrix4();

			matrix.multiplyMatrices( camera.projectionMatrix, matrix.getInverse( camera.matrixWorld ) );
			return this.applyProjection( matrix );

		};

	}(),

	unproject: function () {

		var matrix;

		return function unproject( camera ) {

			if ( matrix === undefined ) matrix = new THREE.Matrix4();

			matrix.multiplyMatrices( camera.matrixWorld, matrix.getInverse( camera.projectionMatrix ) );
			return this.applyProjection( matrix );

		};

	}(),

	transformDirection: function ( m ) {

		// input: THREE.Matrix4 affine matrix
		// vector interpreted as a direction

		var x = this.x, y = this.y, z = this.z;
		var e = m.elements;

		this.x = e[ 0 ] * x + e[ 4 ] * y + e[ 8 ]  * z;
		this.y = e[ 1 ] * x + e[ 5 ] * y + e[ 9 ]  * z;
		this.z = e[ 2 ] * x + e[ 6 ] * y + e[ 10 ] * z;

		return this.normalize();

	},

	divide: function ( v ) {

		this.x /= v.x;
		this.y /= v.y;
		this.z /= v.z;

		return this;

	},

	divideScalar: function ( scalar ) {

		return this.multiplyScalar( 1 / scalar );

	},

	min: function ( v ) {

		this.x = Math.min( this.x, v.x );
		this.y = Math.min( this.y, v.y );
		this.z = Math.min( this.z, v.z );

		return this;

	},

	max: function ( v ) {

		this.x = Math.max( this.x, v.x );
		this.y = Math.max( this.y, v.y );
		this.z = Math.max( this.z, v.z );

		return this;

	},

	clamp: function ( min, max ) {

		// This function assumes min < max, if this assumption isn't true it will not operate correctly

		this.x = Math.max( min.x, Math.min( max.x, this.x ) );
		this.y = Math.max( min.y, Math.min( max.y, this.y ) );
		this.z = Math.max( min.z, Math.min( max.z, this.z ) );

		return this;

	},

	clampScalar: function () {

		var min, max;

		return function clampScalar( minVal, maxVal ) {

			if ( min === undefined ) {

				min = new THREE.Vector3();
				max = new THREE.Vector3();

			}

			min.set( minVal, minVal, minVal );
			max.set( maxVal, maxVal, maxVal );

			return this.clamp( min, max );

		};

	}(),

	clampLength: function ( min, max ) {

		var length = this.length();

		return this.multiplyScalar( Math.max( min, Math.min( max, length ) ) / length );

	},

	floor: function () {

		this.x = Math.floor( this.x );
		this.y = Math.floor( this.y );
		this.z = Math.floor( this.z );

		return this;

	},

	ceil: function () {

		this.x = Math.ceil( this.x );
		this.y = Math.ceil( this.y );
		this.z = Math.ceil( this.z );

		return this;

	},

	round: function () {

		this.x = Math.round( this.x );
		this.y = Math.round( this.y );
		this.z = Math.round( this.z );

		return this;

	},

	roundToZero: function () {

		this.x = ( this.x < 0 ) ? Math.ceil( this.x ) : Math.floor( this.x );
		this.y = ( this.y < 0 ) ? Math.ceil( this.y ) : Math.floor( this.y );
		this.z = ( this.z < 0 ) ? Math.ceil( this.z ) : Math.floor( this.z );

		return this;

	},

	negate: function () {

		this.x = - this.x;
		this.y = - this.y;
		this.z = - this.z;

		return this;

	},

	dot: function ( v ) {

		return this.x * v.x + this.y * v.y + this.z * v.z;

	},

	lengthSq: function () {

		return this.x * this.x + this.y * this.y + this.z * this.z;

	},

	length: function () {

		return Math.sqrt( this.x * this.x + this.y * this.y + this.z * this.z );

	},

	lengthManhattan: function () {

		return Math.abs( this.x ) + Math.abs( this.y ) + Math.abs( this.z );

	},

	normalize: function () {

		return this.divideScalar( this.length() );

	},

	setLength: function ( length ) {

		return this.multiplyScalar( length / this.length() );

	},

	lerp: function ( v, alpha ) {

		this.x += ( v.x - this.x ) * alpha;
		this.y += ( v.y - this.y ) * alpha;
		this.z += ( v.z - this.z ) * alpha;

		return this;

	},

	lerpVectors: function ( v1, v2, alpha ) {

		return this.subVectors( v2, v1 ).multiplyScalar( alpha ).add( v1 );

	},

	cross: function ( v, w ) {

		if ( w !== undefined ) {

			console.warn( 'THREE.Vector3: .cross() now only accepts one argument. Use .crossVectors( a, b ) instead.' );
			return this.crossVectors( v, w );

		}

		var x = this.x, y = this.y, z = this.z;

		this.x = y * v.z - z * v.y;
		this.y = z * v.x - x * v.z;
		this.z = x * v.y - y * v.x;

		return this;

	},

	crossVectors: function ( a, b ) {

		var ax = a.x, ay = a.y, az = a.z;
		var bx = b.x, by = b.y, bz = b.z;

		this.x = ay * bz - az * by;
		this.y = az * bx - ax * bz;
		this.z = ax * by - ay * bx;

		return this;

	},

	projectOnVector: function () {

		var v1, dot;

		return function projectOnVector( vector ) {

			if ( v1 === undefined ) v1 = new THREE.Vector3();

			v1.copy( vector ).normalize();

			dot = this.dot( v1 );

			return this.copy( v1 ).multiplyScalar( dot );

		};

	}(),

	projectOnPlane: function () {

		var v1;

		return function projectOnPlane( planeNormal ) {

			if ( v1 === undefined ) v1 = new THREE.Vector3();

			v1.copy( this ).projectOnVector( planeNormal );

			return this.sub( v1 );

		};

	}(),

	reflect: function () {

		// reflect incident vector off plane orthogonal to normal
		// normal is assumed to have unit length

		var v1;

		return function reflect( normal ) {

			if ( v1 === undefined ) v1 = new THREE.Vector3();

			return this.sub( v1.copy( normal ).multiplyScalar( 2 * this.dot( normal ) ) );

		};

	}(),

	angleTo: function ( v ) {

		var theta = this.dot( v ) / ( Math.sqrt( this.lengthSq() * v.lengthSq() ) );

		// clamp, to handle numerical problems

		return Math.acos( THREE.Math.clamp( theta, - 1, 1 ) );

	},

	distanceTo: function ( v ) {

		return Math.sqrt( this.distanceToSquared( v ) );

	},

	distanceToSquared: function ( v ) {

		var dx = this.x - v.x, dy = this.y - v.y, dz = this.z - v.z;

		return dx * dx + dy * dy + dz * dz;

	},

	setFromSpherical: function( s ) {

		var sinPhiRadius = Math.sin( s.phi ) * s.radius;

		this.x = sinPhiRadius * Math.sin( s.theta );
		this.y = Math.cos( s.phi ) * s.radius;
		this.z = sinPhiRadius * Math.cos( s.theta );

		return this;

	},

	setFromMatrixPosition: function ( m ) {

		return this.setFromMatrixColumn( m, 3 );

	},

	setFromMatrixScale: function ( m ) {

		var sx = this.setFromMatrixColumn( m, 0 ).length();
		var sy = this.setFromMatrixColumn( m, 1 ).length();
		var sz = this.setFromMatrixColumn( m, 2 ).length();

		this.x = sx;
		this.y = sy;
		this.z = sz;

		return this;

	},

	setFromMatrixColumn: function ( m, index ) {

		if ( typeof m === 'number' ) {

			console.warn( 'THREE.Vector3: setFromMatrixColumn now expects ( matrix, index ).' );
			var temp = m
			m = index;
			index = temp;

		}

		return this.fromArray( m.elements, index * 4 );

	},

	equals: function ( v ) {

		return ( ( v.x === this.x ) && ( v.y === this.y ) && ( v.z === this.z ) );

	},

	fromArray: function ( array, offset ) {

		if ( offset === undefined ) offset = 0;

		this.x = array[ offset ];
		this.y = array[ offset + 1 ];
		this.z = array[ offset + 2 ];

		return this;

	},

	toArray: function ( array, offset ) {

		if ( array === undefined ) array = [];
		if ( offset === undefined ) offset = 0;

		array[ offset ] = this.x;
		array[ offset + 1 ] = this.y;
		array[ offset + 2 ] = this.z;

		return array;

	},

	fromAttribute: function ( attribute, index, offset ) {

		if ( offset === undefined ) offset = 0;

		index = index * attribute.itemSize + offset;

		this.x = attribute.array[ index ];
		this.y = attribute.array[ index + 1 ];
		this.z = attribute.array[ index + 2 ];

		return this;

	}

};

// File:src/math/Vector4.js

/**
 * @author supereggbert / http://www.paulbrunt.co.uk/
 * @author philogb / http://blog.thejit.org/
 * @author mikael emtinger / http://gomo.se/
 * @author egraether / http://egraether.com/
 * @author WestLangley / http://github.com/WestLangley
 */

THREE.Vector4 = function ( x, y, z, w ) {

	this.x = x || 0;
	this.y = y || 0;
	this.z = z || 0;
	this.w = ( w !== undefined ) ? w : 1;

};

THREE.Vector4.prototype = {

	constructor: THREE.Vector4,

	set: function ( x, y, z, w ) {

		this.x = x;
		this.y = y;
		this.z = z;
		this.w = w;

		return this;

	},

	setScalar: function ( scalar ) {

		this.x = scalar;
		this.y = scalar;
		this.z = scalar;
		this.w = scalar;

		return this;

	},

	setX: function ( x ) {

		this.x = x;

		return this;

	},

	setY: function ( y ) {

		this.y = y;

		return this;

	},

	setZ: function ( z ) {

		this.z = z;

		return this;

	},

	setW: function ( w ) {

		this.w = w;

		return this;

	},

	setComponent: function ( index, value ) {

		switch ( index ) {

			case 0: this.x = value; break;
			case 1: this.y = value; break;
			case 2: this.z = value; break;
			case 3: this.w = value; break;
			default: throw new Error( 'index is out of range: ' + index );

		}

	},

	getComponent: function ( index ) {

		switch ( index ) {

			case 0: return this.x;
			case 1: return this.y;
			case 2: return this.z;
			case 3: return this.w;
			default: throw new Error( 'index is out of range: ' + index );

		}

	},

	clone: function () {

		return new this.constructor( this.x, this.y, this.z, this.w );

	},

	copy: function ( v ) {

		this.x = v.x;
		this.y = v.y;
		this.z = v.z;
		this.w = ( v.w !== undefined ) ? v.w : 1;

		return this;

	},

	add: function ( v, w ) {

		if ( w !== undefined ) {

			console.warn( 'THREE.Vector4: .add() now only accepts one argument. Use .addVectors( a, b ) instead.' );
			return this.addVectors( v, w );

		}

		this.x += v.x;
		this.y += v.y;
		this.z += v.z;
		this.w += v.w;

		return this;

	},

	addScalar: function ( s ) {

		this.x += s;
		this.y += s;
		this.z += s;
		this.w += s;

		return this;

	},

	addVectors: function ( a, b ) {

		this.x = a.x + b.x;
		this.y = a.y + b.y;
		this.z = a.z + b.z;
		this.w = a.w + b.w;

		return this;

	},

	addScaledVector: function ( v, s ) {

		this.x += v.x * s;
		this.y += v.y * s;
		this.z += v.z * s;
		this.w += v.w * s;

		return this;

	},

	sub: function ( v, w ) {

		if ( w !== undefined ) {

			console.warn( 'THREE.Vector4: .sub() now only accepts one argument. Use .subVectors( a, b ) instead.' );
			return this.subVectors( v, w );

		}

		this.x -= v.x;
		this.y -= v.y;
		this.z -= v.z;
		this.w -= v.w;

		return this;

	},

	subScalar: function ( s ) {

		this.x -= s;
		this.y -= s;
		this.z -= s;
		this.w -= s;

		return this;

	},

	subVectors: function ( a, b ) {

		this.x = a.x - b.x;
		this.y = a.y - b.y;
		this.z = a.z - b.z;
		this.w = a.w - b.w;

		return this;

	},

	multiplyScalar: function ( scalar ) {

		if ( isFinite( scalar ) ) {

			this.x *= scalar;
			this.y *= scalar;
			this.z *= scalar;
			this.w *= scalar;

		} else {

			this.x = 0;
			this.y = 0;
			this.z = 0;
			this.w = 0;

		}

		return this;

	},

	applyMatrix4: function ( m ) {

		var x = this.x, y = this.y, z = this.z, w = this.w;
		var e = m.elements;

		this.x = e[ 0 ] * x + e[ 4 ] * y + e[ 8 ] * z + e[ 12 ] * w;
		this.y = e[ 1 ] * x + e[ 5 ] * y + e[ 9 ] * z + e[ 13 ] * w;
		this.z = e[ 2 ] * x + e[ 6 ] * y + e[ 10 ] * z + e[ 14 ] * w;
		this.w = e[ 3 ] * x + e[ 7 ] * y + e[ 11 ] * z + e[ 15 ] * w;

		return this;

	},

	divideScalar: function ( scalar ) {

		return this.multiplyScalar( 1 / scalar );

	},

	setAxisAngleFromQuaternion: function ( q ) {

		// http://www.euclideanspace.com/maths/geometry/rotations/conversions/quaternionToAngle/index.htm

		// q is assumed to be normalized

		this.w = 2 * Math.acos( q.w );

		var s = Math.sqrt( 1 - q.w * q.w );

		if ( s < 0.0001 ) {

			 this.x = 1;
			 this.y = 0;
			 this.z = 0;

		} else {

			 this.x = q.x / s;
			 this.y = q.y / s;
			 this.z = q.z / s;

		}

		return this;

	},

	setAxisAngleFromRotationMatrix: function ( m ) {

		// http://www.euclideanspace.com/maths/geometry/rotations/conversions/matrixToAngle/index.htm

		// assumes the upper 3x3 of m is a pure rotation matrix (i.e, unscaled)

		var angle, x, y, z,		// variables for result
			epsilon = 0.01,		// margin to allow for rounding errors
			epsilon2 = 0.1,		// margin to distinguish between 0 and 180 degrees

			te = m.elements,

			m11 = te[ 0 ], m12 = te[ 4 ], m13 = te[ 8 ],
			m21 = te[ 1 ], m22 = te[ 5 ], m23 = te[ 9 ],
			m31 = te[ 2 ], m32 = te[ 6 ], m33 = te[ 10 ];

		if ( ( Math.abs( m12 - m21 ) < epsilon ) &&
		     ( Math.abs( m13 - m31 ) < epsilon ) &&
		     ( Math.abs( m23 - m32 ) < epsilon ) ) {

			// singularity found
			// first check for identity matrix which must have +1 for all terms
			// in leading diagonal and zero in other terms

			if ( ( Math.abs( m12 + m21 ) < epsilon2 ) &&
			     ( Math.abs( m13 + m31 ) < epsilon2 ) &&
			     ( Math.abs( m23 + m32 ) < epsilon2 ) &&
			     ( Math.abs( m11 + m22 + m33 - 3 ) < epsilon2 ) ) {

				// this singularity is identity matrix so angle = 0

				this.set( 1, 0, 0, 0 );

				return this; // zero angle, arbitrary axis

			}

			// otherwise this singularity is angle = 180

			angle = Math.PI;

			var xx = ( m11 + 1 ) / 2;
			var yy = ( m22 + 1 ) / 2;
			var zz = ( m33 + 1 ) / 2;
			var xy = ( m12 + m21 ) / 4;
			var xz = ( m13 + m31 ) / 4;
			var yz = ( m23 + m32 ) / 4;

			if ( ( xx > yy ) && ( xx > zz ) ) {

				// m11 is the largest diagonal term

				if ( xx < epsilon ) {

					x = 0;
					y = 0.707106781;
					z = 0.707106781;

				} else {

					x = Math.sqrt( xx );
					y = xy / x;
					z = xz / x;

				}

			} else if ( yy > zz ) {

				// m22 is the largest diagonal term

				if ( yy < epsilon ) {

					x = 0.707106781;
					y = 0;
					z = 0.707106781;

				} else {

					y = Math.sqrt( yy );
					x = xy / y;
					z = yz / y;

				}

			} else {

				// m33 is the largest diagonal term so base result on this

				if ( zz < epsilon ) {

					x = 0.707106781;
					y = 0.707106781;
					z = 0;

				} else {

					z = Math.sqrt( zz );
					x = xz / z;
					y = yz / z;

				}

			}

			this.set( x, y, z, angle );

			return this; // return 180 deg rotation

		}

		// as we have reached here there are no singularities so we can handle normally

		var s = Math.sqrt( ( m32 - m23 ) * ( m32 - m23 ) +
		                   ( m13 - m31 ) * ( m13 - m31 ) +
		                   ( m21 - m12 ) * ( m21 - m12 ) ); // used to normalize

		if ( Math.abs( s ) < 0.001 ) s = 1;

		// prevent divide by zero, should not happen if matrix is orthogonal and should be
		// caught by singularity test above, but I've left it in just in case

		this.x = ( m32 - m23 ) / s;
		this.y = ( m13 - m31 ) / s;
		this.z = ( m21 - m12 ) / s;
		this.w = Math.acos( ( m11 + m22 + m33 - 1 ) / 2 );

		return this;

	},

	min: function ( v ) {

		this.x = Math.min( this.x, v.x );
		this.y = Math.min( this.y, v.y );
		this.z = Math.min( this.z, v.z );
		this.w = Math.min( this.w, v.w );

		return this;

	},

	max: function ( v ) {

		this.x = Math.max( this.x, v.x );
		this.y = Math.max( this.y, v.y );
		this.z = Math.max( this.z, v.z );
		this.w = Math.max( this.w, v.w );

		return this;

	},

	clamp: function ( min, max ) {

		// This function assumes min < max, if this assumption isn't true it will not operate correctly

		this.x = Math.max( min.x, Math.min( max.x, this.x ) );
		this.y = Math.max( min.y, Math.min( max.y, this.y ) );
		this.z = Math.max( min.z, Math.min( max.z, this.z ) );
		this.w = Math.max( min.w, Math.min( max.w, this.w ) );

		return this;

	},

	clampScalar: function () {

		var min, max;

		return function clampScalar( minVal, maxVal ) {

			if ( min === undefined ) {

				min = new THREE.Vector4();
				max = new THREE.Vector4();

			}

			min.set( minVal, minVal, minVal, minVal );
			max.set( maxVal, maxVal, maxVal, maxVal );

			return this.clamp( min, max );

		};

	}(),

	floor: function () {

		this.x = Math.floor( this.x );
		this.y = Math.floor( this.y );
		this.z = Math.floor( this.z );
		this.w = Math.floor( this.w );

		return this;

	},

	ceil: function () {

		this.x = Math.ceil( this.x );
		this.y = Math.ceil( this.y );
		this.z = Math.ceil( this.z );
		this.w = Math.ceil( this.w );

		return this;

	},

	round: function () {

		this.x = Math.round( this.x );
		this.y = Math.round( this.y );
		this.z = Math.round( this.z );
		this.w = Math.round( this.w );

		return this;

	},

	roundToZero: function () {

		this.x = ( this.x < 0 ) ? Math.ceil( this.x ) : Math.floor( this.x );
		this.y = ( this.y < 0 ) ? Math.ceil( this.y ) : Math.floor( this.y );
		this.z = ( this.z < 0 ) ? Math.ceil( this.z ) : Math.floor( this.z );
		this.w = ( this.w < 0 ) ? Math.ceil( this.w ) : Math.floor( this.w );

		return this;

	},

	negate: function () {

		this.x = - this.x;
		this.y = - this.y;
		this.z = - this.z;
		this.w = - this.w;

		return this;

	},

	dot: function ( v ) {

		return this.x * v.x + this.y * v.y + this.z * v.z + this.w * v.w;

	},

	lengthSq: function () {

		return this.x * this.x + this.y * this.y + this.z * this.z + this.w * this.w;

	},

	length: function () {

		return Math.sqrt( this.x * this.x + this.y * this.y + this.z * this.z + this.w * this.w );

	},

	lengthManhattan: function () {

		return Math.abs( this.x ) + Math.abs( this.y ) + Math.abs( this.z ) + Math.abs( this.w );

	},

	normalize: function () {

		return this.divideScalar( this.length() );

	},

	setLength: function ( length ) {

		return this.multiplyScalar( length / this.length() );

	},

	lerp: function ( v, alpha ) {

		this.x += ( v.x - this.x ) * alpha;
		this.y += ( v.y - this.y ) * alpha;
		this.z += ( v.z - this.z ) * alpha;
		this.w += ( v.w - this.w ) * alpha;

		return this;

	},

	lerpVectors: function ( v1, v2, alpha ) {

		return this.subVectors( v2, v1 ).multiplyScalar( alpha ).add( v1 );

	},

	equals: function ( v ) {

		return ( ( v.x === this.x ) && ( v.y === this.y ) && ( v.z === this.z ) && ( v.w === this.w ) );

	},

	fromArray: function ( array, offset ) {

		if ( offset === undefined ) offset = 0;

		this.x = array[ offset ];
		this.y = array[ offset + 1 ];
		this.z = array[ offset + 2 ];
		this.w = array[ offset + 3 ];

		return this;

	},

	toArray: function ( array, offset ) {

		if ( array === undefined ) array = [];
		if ( offset === undefined ) offset = 0;

		array[ offset ] = this.x;
		array[ offset + 1 ] = this.y;
		array[ offset + 2 ] = this.z;
		array[ offset + 3 ] = this.w;

		return array;

	},

	fromAttribute: function ( attribute, index, offset ) {

		if ( offset === undefined ) offset = 0;

		index = index * attribute.itemSize + offset;

		this.x = attribute.array[ index ];
		this.y = attribute.array[ index + 1 ];
		this.z = attribute.array[ index + 2 ];
		this.w = attribute.array[ index + 3 ];

		return this;

	}

};

// File:src/math/Euler.js

/**
 * @author mrdoob / http://mrdoob.com/
 * @author WestLangley / http://github.com/WestLangley
 * @author bhouston / http://clara.io
 */

THREE.Euler = function ( x, y, z, order ) {

	this._x = x || 0;
	this._y = y || 0;
	this._z = z || 0;
	this._order = order || THREE.Euler.DefaultOrder;

};

THREE.Euler.RotationOrders = [ 'XYZ', 'YZX', 'ZXY', 'XZY', 'YXZ', 'ZYX' ];

THREE.Euler.DefaultOrder = 'XYZ';

THREE.Euler.prototype = {

	constructor: THREE.Euler,

	get x () {

		return this._x;

	},

	set x ( value ) {

		this._x = value;
		this.onChangeCallback();

	},

	get y () {

		return this._y;

	},

	set y ( value ) {

		this._y = value;
		this.onChangeCallback();

	},

	get z () {

		return this._z;

	},

	set z ( value ) {

		this._z = value;
		this.onChangeCallback();

	},

	get order () {

		return this._order;

	},

	set order ( value ) {

		this._order = value;
		this.onChangeCallback();

	},

	set: function ( x, y, z, order ) {

		this._x = x;
		this._y = y;
		this._z = z;
		this._order = order || this._order;

		this.onChangeCallback();

		return this;

	},

	clone: function () {

		return new this.constructor( this._x, this._y, this._z, this._order );

	},

	copy: function ( euler ) {

		this._x = euler._x;
		this._y = euler._y;
		this._z = euler._z;
		this._order = euler._order;

		this.onChangeCallback();

		return this;

	},

	setFromRotationMatrix: function ( m, order, update ) {

		var clamp = THREE.Math.clamp;

		// assumes the upper 3x3 of m is a pure rotation matrix (i.e, unscaled)

		var te = m.elements;
		var m11 = te[ 0 ], m12 = te[ 4 ], m13 = te[ 8 ];
		var m21 = te[ 1 ], m22 = te[ 5 ], m23 = te[ 9 ];
		var m31 = te[ 2 ], m32 = te[ 6 ], m33 = te[ 10 ];

		order = order || this._order;

		if ( order === 'XYZ' ) {

			this._y = Math.asin( clamp( m13, - 1, 1 ) );

			if ( Math.abs( m13 ) < 0.99999 ) {

				this._x = Math.atan2( - m23, m33 );
				this._z = Math.atan2( - m12, m11 );

			} else {

				this._x = Math.atan2( m32, m22 );
				this._z = 0;

			}

		} else if ( order === 'YXZ' ) {

			this._x = Math.asin( - clamp( m23, - 1, 1 ) );

			if ( Math.abs( m23 ) < 0.99999 ) {

				this._y = Math.atan2( m13, m33 );
				this._z = Math.atan2( m21, m22 );

			} else {

				this._y = Math.atan2( - m31, m11 );
				this._z = 0;

			}

		} else if ( order === 'ZXY' ) {

			this._x = Math.asin( clamp( m32, - 1, 1 ) );

			if ( Math.abs( m32 ) < 0.99999 ) {

				this._y = Math.atan2( - m31, m33 );
				this._z = Math.atan2( - m12, m22 );

			} else {

				this._y = 0;
				this._z = Math.atan2( m21, m11 );

			}

		} else if ( order === 'ZYX' ) {

			this._y = Math.asin( - clamp( m31, - 1, 1 ) );

			if ( Math.abs( m31 ) < 0.99999 ) {

				this._x = Math.atan2( m32, m33 );
				this._z = Math.atan2( m21, m11 );

			} else {

				this._x = 0;
				this._z = Math.atan2( - m12, m22 );

			}

		} else if ( order === 'YZX' ) {

			this._z = Math.asin( clamp( m21, - 1, 1 ) );

			if ( Math.abs( m21 ) < 0.99999 ) {

				this._x = Math.atan2( - m23, m22 );
				this._y = Math.atan2( - m31, m11 );

			} else {

				this._x = 0;
				this._y = Math.atan2( m13, m33 );

			}

		} else if ( order === 'XZY' ) {

			this._z = Math.asin( - clamp( m12, - 1, 1 ) );

			if ( Math.abs( m12 ) < 0.99999 ) {

				this._x = Math.atan2( m32, m22 );
				this._y = Math.atan2( m13, m11 );

			} else {

				this._x = Math.atan2( - m23, m33 );
				this._y = 0;

			}

		} else {

			console.warn( 'THREE.Euler: .setFromRotationMatrix() given unsupported order: ' + order );

		}

		this._order = order;

		if ( update !== false ) this.onChangeCallback();

		return this;

	},

	setFromQuaternion: function () {

		var matrix;

		return function ( q, order, update ) {

			if ( matrix === undefined ) matrix = new THREE.Matrix4();

			matrix.makeRotationFromQuaternion( q );

			return this.setFromRotationMatrix( matrix, order, update );

		};

	}(),

	setFromVector3: function ( v, order ) {

		return this.set( v.x, v.y, v.z, order || this._order );

	},

	reorder: function () {

		// WARNING: this discards revolution information -bhouston

		var q = new THREE.Quaternion();

		return function ( newOrder ) {

			q.setFromEuler( this );
			
			return this.setFromQuaternion( q, newOrder );

		};

	}(),

	equals: function ( euler ) {

		return ( euler._x === this._x ) && ( euler._y === this._y ) && ( euler._z === this._z ) && ( euler._order === this._order );

	},

	fromArray: function ( array ) {

		this._x = array[ 0 ];
		this._y = array[ 1 ];
		this._z = array[ 2 ];
		if ( array[ 3 ] !== undefined ) this._order = array[ 3 ];

		this.onChangeCallback();

		return this;

	},

	toArray: function ( array, offset ) {

		if ( array === undefined ) array = [];
		if ( offset === undefined ) offset = 0;

		array[ offset ] = this._x;
		array[ offset + 1 ] = this._y;
		array[ offset + 2 ] = this._z;
		array[ offset + 3 ] = this._order;

		return array;

	},

	toVector3: function ( optionalResult ) {

		if ( optionalResult ) {

			return optionalResult.set( this._x, this._y, this._z );

		} else {

			return new THREE.Vector3( this._x, this._y, this._z );

		}

	},

	onChange: function ( callback ) {

		this.onChangeCallback = callback;

		return this;

	},

	onChangeCallback: function () {}

};

// File:src/math/Line3.js

/**
 * @author bhouston / http://clara.io
 */

THREE.Line3 = function ( start, end ) {

	this.start = ( start !== undefined ) ? start : new THREE.Vector3();
	this.end = ( end !== undefined ) ? end : new THREE.Vector3();

};

THREE.Line3.prototype = {

	constructor: THREE.Line3,

	set: function ( start, end ) {

		this.start.copy( start );
		this.end.copy( end );

		return this;

	},

	clone: function () {

		return new this.constructor().copy( this );

	},

	copy: function ( line ) {

		this.start.copy( line.start );
		this.end.copy( line.end );

		return this;

	},

	center: function ( optionalTarget ) {

		var result = optionalTarget || new THREE.Vector3();
		return result.addVectors( this.start, this.end ).multiplyScalar( 0.5 );

	},

	delta: function ( optionalTarget ) {

		var result = optionalTarget || new THREE.Vector3();
		return result.subVectors( this.end, this.start );

	},

	distanceSq: function () {

		return this.start.distanceToSquared( this.end );

	},

	distance: function () {

		return this.start.distanceTo( this.end );

	},

	at: function ( t, optionalTarget ) {

		var result = optionalTarget || new THREE.Vector3();

		return this.delta( result ).multiplyScalar( t ).add( this.start );

	},

	closestPointToPointParameter: function () {

		var startP = new THREE.Vector3();
		var startEnd = new THREE.Vector3();

		return function ( point, clampToLine ) {

			startP.subVectors( point, this.start );
			startEnd.subVectors( this.end, this.start );

			var startEnd2 = startEnd.dot( startEnd );
			var startEnd_startP = startEnd.dot( startP );

			var t = startEnd_startP / startEnd2;

			if ( clampToLine ) {

				t = THREE.Math.clamp( t, 0, 1 );

			}

			return t;

		};

	}(),

	closestPointToPoint: function ( point, clampToLine, optionalTarget ) {

		var t = this.closestPointToPointParameter( point, clampToLine );

		var result = optionalTarget || new THREE.Vector3();

		return this.delta( result ).multiplyScalar( t ).add( this.start );

	},

	applyMatrix4: function ( matrix ) {

		this.start.applyMatrix4( matrix );
		this.end.applyMatrix4( matrix );

		return this;

	},

	equals: function ( line ) {

		return line.start.equals( this.start ) && line.end.equals( this.end );

	}

};

// File:src/math/Box2.js

/**
 * @author bhouston / http://clara.io
 */

THREE.Box2 = function ( min, max ) {

	this.min = ( min !== undefined ) ? min : new THREE.Vector2( + Infinity, + Infinity );
	this.max = ( max !== undefined ) ? max : new THREE.Vector2( - Infinity, - Infinity );

};

THREE.Box2.prototype = {

	constructor: THREE.Box2,

	set: function ( min, max ) {

		this.min.copy( min );
		this.max.copy( max );

		return this;

	},

	setFromPoints: function ( points ) {

		this.makeEmpty();

		for ( var i = 0, il = points.length; i < il; i ++ ) {

			this.expandByPoint( points[ i ] );

		}

		return this;

	},

	setFromCenterAndSize: function () {

		var v1 = new THREE.Vector2();

		return function ( center, size ) {

			var halfSize = v1.copy( size ).multiplyScalar( 0.5 );
			this.min.copy( center ).sub( halfSize );
			this.max.copy( center ).add( halfSize );

			return this;

		};

	}(),

	clone: function () {

		return new this.constructor().copy( this );

	},

	copy: function ( box ) {

		this.min.copy( box.min );
		this.max.copy( box.max );

		return this;

	},

	makeEmpty: function () {

		this.min.x = this.min.y = + Infinity;
		this.max.x = this.max.y = - Infinity;

		return this;

	},

	isEmpty: function () {

		// this is a more robust check for empty than ( volume <= 0 ) because volume can get positive with two negative axes

		return ( this.max.x < this.min.x ) || ( this.max.y < this.min.y );

	},

	center: function ( optionalTarget ) {

		var result = optionalTarget || new THREE.Vector2();
		return result.addVectors( this.min, this.max ).multiplyScalar( 0.5 );

	},

	size: function ( optionalTarget ) {

		var result = optionalTarget || new THREE.Vector2();
		return result.subVectors( this.max, this.min );

	},

	expandByPoint: function ( point ) {

		this.min.min( point );
		this.max.max( point );

		return this;

	},

	expandByVector: function ( vector ) {

		this.min.sub( vector );
		this.max.add( vector );

		return this;

	},

	expandByScalar: function ( scalar ) {

		this.min.addScalar( - scalar );
		this.max.addScalar( scalar );

		return this;

	},

	containsPoint: function ( point ) {

		if ( point.x < this.min.x || point.x > this.max.x ||
		     point.y < this.min.y || point.y > this.max.y ) {

			return false;

		}

		return true;

	},

	containsBox: function ( box ) {

		if ( ( this.min.x <= box.min.x ) && ( box.max.x <= this.max.x ) &&
		     ( this.min.y <= box.min.y ) && ( box.max.y <= this.max.y ) ) {

			return true;

		}

		return false;

	},

	getParameter: function ( point, optionalTarget ) {

		// This can potentially have a divide by zero if the box
		// has a size dimension of 0.

		var result = optionalTarget || new THREE.Vector2();

		return result.set(
			( point.x - this.min.x ) / ( this.max.x - this.min.x ),
			( point.y - this.min.y ) / ( this.max.y - this.min.y )
		);

	},

	intersectsBox: function ( box ) {

		// using 6 splitting planes to rule out intersections.

		if ( box.max.x < this.min.x || box.min.x > this.max.x ||
		     box.max.y < this.min.y || box.min.y > this.max.y ) {

			return false;

		}

		return true;

	},

	clampPoint: function ( point, optionalTarget ) {

		var result = optionalTarget || new THREE.Vector2();
		return result.copy( point ).clamp( this.min, this.max );

	},

	distanceToPoint: function () {

		var v1 = new THREE.Vector2();

		return function ( point ) {

			var clampedPoint = v1.copy( point ).clamp( this.min, this.max );
			return clampedPoint.sub( point ).length();

		};

	}(),

	intersect: function ( box ) {

		this.min.max( box.min );
		this.max.min( box.max );

		return this;

	},

	union: function ( box ) {

		this.min.min( box.min );
		this.max.max( box.max );

		return this;

	},

	translate: function ( offset ) {

		this.min.add( offset );
		this.max.add( offset );

		return this;

	},

	equals: function ( box ) {

		return box.min.equals( this.min ) && box.max.equals( this.max );

	}

};

// File:src/math/Box3.js

/**
 * @author bhouston / http://clara.io
 * @author WestLangley / http://github.com/WestLangley
 */

THREE.Box3 = function ( min, max ) {

	this.min = ( min !== undefined ) ? min : new THREE.Vector3( + Infinity, + Infinity, + Infinity );
	this.max = ( max !== undefined ) ? max : new THREE.Vector3( - Infinity, - Infinity, - Infinity );

};

THREE.Box3.prototype = {

	constructor: THREE.Box3,

	set: function ( min, max ) {

		this.min.copy( min );
		this.max.copy( max );

		return this;

	},

	setFromArray: function ( array ) {

		var minX = + Infinity;
		var minY = + Infinity;
		var minZ = + Infinity;

		var maxX = - Infinity;
		var maxY = - Infinity;
		var maxZ = - Infinity;

		for ( var i = 0, l = array.length; i < l; i += 3 ) {

			var x = array[ i ];
			var y = array[ i + 1 ];
			var z = array[ i + 2 ];

			if ( x < minX ) minX = x;
			if ( y < minY ) minY = y;
			if ( z < minZ ) minZ = z;

			if ( x > maxX ) maxX = x;
			if ( y > maxY ) maxY = y;
			if ( z > maxZ ) maxZ = z;

		}

		this.min.set( minX, minY, minZ );
		this.max.set( maxX, maxY, maxZ );

	},

	setFromPoints: function ( points ) {

		this.makeEmpty();

		for ( var i = 0, il = points.length; i < il; i ++ ) {

			this.expandByPoint( points[ i ] );

		}

		return this;

	},

	setFromCenterAndSize: function () {

		var v1 = new THREE.Vector3();

		return function ( center, size ) {

			var halfSize = v1.copy( size ).multiplyScalar( 0.5 );

			this.min.copy( center ).sub( halfSize );
			this.max.copy( center ).add( halfSize );

			return this;

		};

	}(),

	setFromObject: function () {

		// Computes the world-axis-aligned bounding box of an object (including its children),
		// accounting for both the object's, and children's, world transforms

		var v1 = new THREE.Vector3();

		return function ( object ) {

			var scope = this;

			object.updateMatrixWorld( true );

			this.makeEmpty();

			object.traverse( function ( node ) {

				var geometry = node.geometry;

				if ( geometry !== undefined ) {

					if ( geometry instanceof THREE.Geometry ) {

						var vertices = geometry.vertices;

						for ( var i = 0, il = vertices.length; i < il; i ++ ) {

							v1.copy( vertices[ i ] );
							v1.applyMatrix4( node.matrixWorld );

							scope.expandByPoint( v1 );

						}

					} else if ( geometry instanceof THREE.BufferGeometry && geometry.attributes[ 'position' ] !== undefined ) {

						var positions = geometry.attributes[ 'position' ].array;

						for ( var i = 0, il = positions.length; i < il; i += 3 ) {

							v1.fromArray( positions, i );
							v1.applyMatrix4( node.matrixWorld );

							scope.expandByPoint( v1 );

						}

					}

				}

			} );

			return this;

		};

	}(),

	clone: function () {

		return new this.constructor().copy( this );

	},

	copy: function ( box ) {

		this.min.copy( box.min );
		this.max.copy( box.max );

		return this;

	},

	makeEmpty: function () {

		this.min.x = this.min.y = this.min.z = + Infinity;
		this.max.x = this.max.y = this.max.z = - Infinity;

		return this;

	},

	isEmpty: function () {

		// this is a more robust check for empty than ( volume <= 0 ) because volume can get positive with two negative axes

		return ( this.max.x < this.min.x ) || ( this.max.y < this.min.y ) || ( this.max.z < this.min.z );

	},

	center: function ( optionalTarget ) {

		var result = optionalTarget || new THREE.Vector3();
		return result.addVectors( this.min, this.max ).multiplyScalar( 0.5 );

	},

	size: function ( optionalTarget ) {

		var result = optionalTarget || new THREE.Vector3();
		return result.subVectors( this.max, this.min );

	},

	expandByPoint: function ( point ) {

		this.min.min( point );
		this.max.max( point );

		return this;

	},

	expandByVector: function ( vector ) {

		this.min.sub( vector );
		this.max.add( vector );

		return this;

	},

	expandByScalar: function ( scalar ) {

		this.min.addScalar( - scalar );
		this.max.addScalar( scalar );

		return this;

	},

	containsPoint: function ( point ) {

		if ( point.x < this.min.x || point.x > this.max.x ||
				 point.y < this.min.y || point.y > this.max.y ||
				 point.z < this.min.z || point.z > this.max.z ) {

			return false;

		}

		return true;

	},

	containsBox: function ( box ) {

		if ( ( this.min.x <= box.min.x ) && ( box.max.x <= this.max.x ) &&
			 ( this.min.y <= box.min.y ) && ( box.max.y <= this.max.y ) &&
			 ( this.min.z <= box.min.z ) && ( box.max.z <= this.max.z ) ) {

			return true;

		}

		return false;

	},

	getParameter: function ( point, optionalTarget ) {

		// This can potentially have a divide by zero if the box
		// has a size dimension of 0.

		var result = optionalTarget || new THREE.Vector3();

		return result.set(
			( point.x - this.min.x ) / ( this.max.x - this.min.x ),
			( point.y - this.min.y ) / ( this.max.y - this.min.y ),
			( point.z - this.min.z ) / ( this.max.z - this.min.z )
		);

	},

	intersectsBox: function ( box ) {

		// using 6 splitting planes to rule out intersections.

		if ( box.max.x < this.min.x || box.min.x > this.max.x ||
				 box.max.y < this.min.y || box.min.y > this.max.y ||
				 box.max.z < this.min.z || box.min.z > this.max.z ) {

			return false;

		}

		return true;

	},

	intersectsSphere: ( function () {

		var closestPoint;

		return function intersectsSphere( sphere ) {

			if ( closestPoint === undefined ) closestPoint = new THREE.Vector3();

			// Find the point on the AABB closest to the sphere center.
			this.clampPoint( sphere.center, closestPoint );

			// If that point is inside the sphere, the AABB and sphere intersect.
			return closestPoint.distanceToSquared( sphere.center ) <= ( sphere.radius * sphere.radius );

		};

	} )(),

	intersectsPlane: function ( plane ) {

		// We compute the minimum and maximum dot product values. If those values
		// are on the same side (back or front) of the plane, then there is no intersection.

		var min, max;

		if ( plane.normal.x > 0 ) {

			min = plane.normal.x * this.min.x;
			max = plane.normal.x * this.max.x;

		} else {

			min = plane.normal.x * this.max.x;
			max = plane.normal.x * this.min.x;

		}

		if ( plane.normal.y > 0 ) {

			min += plane.normal.y * this.min.y;
			max += plane.normal.y * this.max.y;

		} else {

			min += plane.normal.y * this.max.y;
			max += plane.normal.y * this.min.y;

		}

		if ( plane.normal.z > 0 ) {

			min += plane.normal.z * this.min.z;
			max += plane.normal.z * this.max.z;

		} else {

			min += plane.normal.z * this.max.z;
			max += plane.normal.z * this.min.z;

		}

		return ( min <= plane.constant && max >= plane.constant );

	},

	clampPoint: function ( point, optionalTarget ) {

		var result = optionalTarget || new THREE.Vector3();
		return result.copy( point ).clamp( this.min, this.max );

	},

	distanceToPoint: function () {

		var v1 = new THREE.Vector3();

		return function ( point ) {

			var clampedPoint = v1.copy( point ).clamp( this.min, this.max );
			return clampedPoint.sub( point ).length();

		};

	}(),

	getBoundingSphere: function () {

		var v1 = new THREE.Vector3();

		return function ( optionalTarget ) {

			var result = optionalTarget || new THREE.Sphere();

			result.center = this.center();
			result.radius = this.size( v1 ).length() * 0.5;

			return result;

		};

	}(),

	intersect: function ( box ) {

		this.min.max( box.min );
		this.max.min( box.max );

		// ensure that if there is no overlap, the result is fully empty, not slightly empty with non-inf/+inf values that will cause subsequence intersects to erroneously return valid values.
		if( this.isEmpty() ) this.makeEmpty();

		return this;

	},

	union: function ( box ) {

		this.min.min( box.min );
		this.max.max( box.max );

		return this;

	},

	applyMatrix4: function () {

		var points = [
			new THREE.Vector3(),
			new THREE.Vector3(),
			new THREE.Vector3(),
			new THREE.Vector3(),
			new THREE.Vector3(),
			new THREE.Vector3(),
			new THREE.Vector3(),
			new THREE.Vector3()
		];

		return function ( matrix ) {

			// transform of empty box is an empty box.
			if( this.isEmpty() ) return this;

			// NOTE: I am using a binary pattern to specify all 2^3 combinations below
			points[ 0 ].set( this.min.x, this.min.y, this.min.z ).applyMatrix4( matrix ); // 000
			points[ 1 ].set( this.min.x, this.min.y, this.max.z ).applyMatrix4( matrix ); // 001
			points[ 2 ].set( this.min.x, this.max.y, this.min.z ).applyMatrix4( matrix ); // 010
			points[ 3 ].set( this.min.x, this.max.y, this.max.z ).applyMatrix4( matrix ); // 011
			points[ 4 ].set( this.max.x, this.min.y, this.min.z ).applyMatrix4( matrix ); // 100
			points[ 5 ].set( this.max.x, this.min.y, this.max.z ).applyMatrix4( matrix ); // 101
			points[ 6 ].set( this.max.x, this.max.y, this.min.z ).applyMatrix4( matrix ); // 110
			points[ 7 ].set( this.max.x, this.max.y, this.max.z ).applyMatrix4( matrix );	// 111

			this.setFromPoints( points );

			return this;

		};

	}(),

	translate: function ( offset ) {

		this.min.add( offset );
		this.max.add( offset );

		return this;

	},

	equals: function ( box ) {

		return box.min.equals( this.min ) && box.max.equals( this.max );

	}

};

// File:src/math/Matrix3.js

/**
 * @author alteredq / http://alteredqualia.com/
 * @author WestLangley / http://github.com/WestLangley
 * @author bhouston / http://clara.io
 * @author tschw
 */

THREE.Matrix3 = function () {

	this.elements = new Float32Array( [

		1, 0, 0,
		0, 1, 0,
		0, 0, 1

	] );

	if ( arguments.length > 0 ) {

		console.error( 'THREE.Matrix3: the constructor no longer reads arguments. use .set() instead.' );

	}

};

THREE.Matrix3.prototype = {

	constructor: THREE.Matrix3,

	set: function ( n11, n12, n13, n21, n22, n23, n31, n32, n33 ) {

		var te = this.elements;

		te[ 0 ] = n11; te[ 1 ] = n21; te[ 2 ] = n31;
		te[ 3 ] = n12; te[ 4 ] = n22; te[ 5 ] = n32;
		te[ 6 ] = n13; te[ 7 ] = n23; te[ 8 ] = n33;

		return this;

	},

	identity: function () {

		this.set(

			1, 0, 0,
			0, 1, 0,
			0, 0, 1

		);

		return this;

	},

	clone: function () {

		return new this.constructor().fromArray( this.elements );

	},

	copy: function ( m ) {

		var me = m.elements;

		this.set(

			me[ 0 ], me[ 3 ], me[ 6 ],
			me[ 1 ], me[ 4 ], me[ 7 ],
			me[ 2 ], me[ 5 ], me[ 8 ]

		);

		return this;

	},

	setFromMatrix4: function( m ) {

		var me = m.elements;

		this.set(

			me[ 0 ], me[ 4 ], me[  8 ],
			me[ 1 ], me[ 5 ], me[  9 ],
			me[ 2 ], me[ 6 ], me[ 10 ]

		);

		return this;

	},

	applyToVector3Array: function () {

		var v1;

		return function ( array, offset, length ) {

			if ( v1 === undefined ) v1 = new THREE.Vector3();
			if ( offset === undefined ) offset = 0;
			if ( length === undefined ) length = array.length;

			for ( var i = 0, j = offset; i < length; i += 3, j += 3 ) {

				v1.fromArray( array, j );
				v1.applyMatrix3( this );
				v1.toArray( array, j );

			}

			return array;

		};

	}(),

	applyToBuffer: function () {

		var v1;

		return function applyToBuffer( buffer, offset, length ) {

			if ( v1 === undefined ) v1 = new THREE.Vector3();
			if ( offset === undefined ) offset = 0;
			if ( length === undefined ) length = buffer.length / buffer.itemSize;

			for ( var i = 0, j = offset; i < length; i ++, j ++ ) {

				v1.x = buffer.getX( j );
				v1.y = buffer.getY( j );
				v1.z = buffer.getZ( j );

				v1.applyMatrix3( this );

				buffer.setXYZ( v1.x, v1.y, v1.z );

			}

			return buffer;

		};

	}(),

	multiplyScalar: function ( s ) {

		var te = this.elements;

		te[ 0 ] *= s; te[ 3 ] *= s; te[ 6 ] *= s;
		te[ 1 ] *= s; te[ 4 ] *= s; te[ 7 ] *= s;
		te[ 2 ] *= s; te[ 5 ] *= s; te[ 8 ] *= s;

		return this;

	},

	determinant: function () {

		var te = this.elements;

		var a = te[ 0 ], b = te[ 1 ], c = te[ 2 ],
			d = te[ 3 ], e = te[ 4 ], f = te[ 5 ],
			g = te[ 6 ], h = te[ 7 ], i = te[ 8 ];

		return a * e * i - a * f * h - b * d * i + b * f * g + c * d * h - c * e * g;

	},

	getInverse: function ( matrix, throwOnDegenerate ) {

		if ( matrix instanceof THREE.Matrix4 ) {

			console.error( "THREE.Matrix3.getInverse no longer takes a Matrix4 argument." );

		}

		var me = matrix.elements,
			te = this.elements,

			n11 = me[ 0 ], n21 = me[ 1 ], n31 = me[ 2 ],
			n12 = me[ 3 ], n22 = me[ 4 ], n32 = me[ 5 ],
			n13 = me[ 6 ], n23 = me[ 7 ], n33 = me[ 8 ],

			t11 = n33 * n22 - n32 * n23,
			t12 = n32 * n13 - n33 * n12,
			t13 = n23 * n12 - n22 * n13,

			det = n11 * t11 + n21 * t12 + n31 * t13;

		if ( det === 0 ) {

			var msg = "THREE.Matrix3.getInverse(): can't invert matrix, determinant is 0";

			if ( throwOnDegenerate || false ) {

				throw new Error( msg );

			} else {

				console.warn( msg );

			}

			return this.identity();
		}
		
		var detInv = 1 / det;

		te[ 0 ] = t11 * detInv;
		te[ 1 ] = ( n31 * n23 - n33 * n21 ) * detInv;
		te[ 2 ] = ( n32 * n21 - n31 * n22 ) * detInv;

		te[ 3 ] = t12 * detInv;
		te[ 4 ] = ( n33 * n11 - n31 * n13 ) * detInv;
		te[ 5 ] = ( n31 * n12 - n32 * n11 ) * detInv;

		te[ 6 ] = t13 * detInv;
		te[ 7 ] = ( n21 * n13 - n23 * n11 ) * detInv;
		te[ 8 ] = ( n22 * n11 - n21 * n12 ) * detInv;

		return this;

	},

	transpose: function () {

		var tmp, m = this.elements;

		tmp = m[ 1 ]; m[ 1 ] = m[ 3 ]; m[ 3 ] = tmp;
		tmp = m[ 2 ]; m[ 2 ] = m[ 6 ]; m[ 6 ] = tmp;
		tmp = m[ 5 ]; m[ 5 ] = m[ 7 ]; m[ 7 ] = tmp;

		return this;

	},

	flattenToArrayOffset: function ( array, offset ) {

		console.warn( "THREE.Matrix3: .flattenToArrayOffset is deprecated " +
				"- just use .toArray instead." );

		return this.toArray( array, offset );

	},

	getNormalMatrix: function ( matrix4 ) {

		return this.setFromMatrix4( matrix4 ).getInverse( this ).transpose();

	},

	transposeIntoArray: function ( r ) {

		var m = this.elements;

		r[ 0 ] = m[ 0 ];
		r[ 1 ] = m[ 3 ];
		r[ 2 ] = m[ 6 ];
		r[ 3 ] = m[ 1 ];
		r[ 4 ] = m[ 4 ];
		r[ 5 ] = m[ 7 ];
		r[ 6 ] = m[ 2 ];
		r[ 7 ] = m[ 5 ];
		r[ 8 ] = m[ 8 ];

		return this;

	},

	fromArray: function ( array ) {

		this.elements.set( array );

		return this;

	},

	toArray: function ( array, offset ) {

		if ( array === undefined ) array = [];
		if ( offset === undefined ) offset = 0;

		var te = this.elements;

		array[ offset ] = te[ 0 ];
		array[ offset + 1 ] = te[ 1 ];
		array[ offset + 2 ] = te[ 2 ];

		array[ offset + 3 ] = te[ 3 ];
		array[ offset + 4 ] = te[ 4 ];
		array[ offset + 5 ] = te[ 5 ];

		array[ offset + 6 ] = te[ 6 ];
		array[ offset + 7 ] = te[ 7 ];
		array[ offset + 8 ]  = te[ 8 ];

		return array;

	}

};

// File:src/math/Matrix4.js

/**
 * @author mrdoob / http://mrdoob.com/
 * @author supereggbert / http://www.paulbrunt.co.uk/
 * @author philogb / http://blog.thejit.org/
 * @author jordi_ros / http://plattsoft.com
 * @author D1plo1d / http://github.com/D1plo1d
 * @author alteredq / http://alteredqualia.com/
 * @author mikael emtinger / http://gomo.se/
 * @author timknip / http://www.floorplanner.com/
 * @author bhouston / http://clara.io
 * @author WestLangley / http://github.com/WestLangley
 */

THREE.Matrix4 = function () {

	this.elements = new Float32Array( [

		1, 0, 0, 0,
		0, 1, 0, 0,
		0, 0, 1, 0,
		0, 0, 0, 1

	] );

	if ( arguments.length > 0 ) {

		console.error( 'THREE.Matrix4: the constructor no longer reads arguments. use .set() instead.' );

	}

};

THREE.Matrix4.prototype = {

	constructor: THREE.Matrix4,

	set: function ( n11, n12, n13, n14, n21, n22, n23, n24, n31, n32, n33, n34, n41, n42, n43, n44 ) {

		var te = this.elements;

		te[ 0 ] = n11; te[ 4 ] = n12; te[ 8 ] = n13; te[ 12 ] = n14;
		te[ 1 ] = n21; te[ 5 ] = n22; te[ 9 ] = n23; te[ 13 ] = n24;
		te[ 2 ] = n31; te[ 6 ] = n32; te[ 10 ] = n33; te[ 14 ] = n34;
		te[ 3 ] = n41; te[ 7 ] = n42; te[ 11 ] = n43; te[ 15 ] = n44;

		return this;

	},

	identity: function () {

		this.set(

			1, 0, 0, 0,
			0, 1, 0, 0,
			0, 0, 1, 0,
			0, 0, 0, 1

		);

		return this;

	},

	clone: function () {

		return new THREE.Matrix4().fromArray( this.elements );

	},

	copy: function ( m ) {

		this.elements.set( m.elements );

		return this;

	},

	copyPosition: function ( m ) {

		var te = this.elements;
		var me = m.elements;

		te[ 12 ] = me[ 12 ];
		te[ 13 ] = me[ 13 ];
		te[ 14 ] = me[ 14 ];

		return this;

	},

	extractBasis: function ( xAxis, yAxis, zAxis ) {

		xAxis.setFromMatrixColumn( this, 0 );
		yAxis.setFromMatrixColumn( this, 1 );
		zAxis.setFromMatrixColumn( this, 2 );

		return this;

	},

	makeBasis: function ( xAxis, yAxis, zAxis ) {

		this.set(
			xAxis.x, yAxis.x, zAxis.x, 0,
			xAxis.y, yAxis.y, zAxis.y, 0,
			xAxis.z, yAxis.z, zAxis.z, 0,
			0,       0,       0,       1
		);

		return this;

	},

	extractRotation: function () {

		var v1;

		return function ( m ) {

			if ( v1 === undefined ) v1 = new THREE.Vector3();

			var te = this.elements;
			var me = m.elements;

			var scaleX = 1 / v1.setFromMatrixColumn( m, 0 ).length();
			var scaleY = 1 / v1.setFromMatrixColumn( m, 1 ).length();
			var scaleZ = 1 / v1.setFromMatrixColumn( m, 2 ).length();

			te[ 0 ] = me[ 0 ] * scaleX;
			te[ 1 ] = me[ 1 ] * scaleX;
			te[ 2 ] = me[ 2 ] * scaleX;

			te[ 4 ] = me[ 4 ] * scaleY;
			te[ 5 ] = me[ 5 ] * scaleY;
			te[ 6 ] = me[ 6 ] * scaleY;

			te[ 8 ] = me[ 8 ] * scaleZ;
			te[ 9 ] = me[ 9 ] * scaleZ;
			te[ 10 ] = me[ 10 ] * scaleZ;

			return this;

		};

	}(),

	makeRotationFromEuler: function ( euler ) {

		if ( euler instanceof THREE.Euler === false ) {

			console.error( 'THREE.Matrix: .makeRotationFromEuler() now expects a Euler rotation rather than a Vector3 and order.' );

		}

		var te = this.elements;

		var x = euler.x, y = euler.y, z = euler.z;
		var a = Math.cos( x ), b = Math.sin( x );
		var c = Math.cos( y ), d = Math.sin( y );
		var e = Math.cos( z ), f = Math.sin( z );

		if ( euler.order === 'XYZ' ) {

			var ae = a * e, af = a * f, be = b * e, bf = b * f;

			te[ 0 ] = c * e;
			te[ 4 ] = - c * f;
			te[ 8 ] = d;

			te[ 1 ] = af + be * d;
			te[ 5 ] = ae - bf * d;
			te[ 9 ] = - b * c;

			te[ 2 ] = bf - ae * d;
			te[ 6 ] = be + af * d;
			te[ 10 ] = a * c;

		} else if ( euler.order === 'YXZ' ) {

			var ce = c * e, cf = c * f, de = d * e, df = d * f;

			te[ 0 ] = ce + df * b;
			te[ 4 ] = de * b - cf;
			te[ 8 ] = a * d;

			te[ 1 ] = a * f;
			te[ 5 ] = a * e;
			te[ 9 ] = - b;

			te[ 2 ] = cf * b - de;
			te[ 6 ] = df + ce * b;
			te[ 10 ] = a * c;

		} else if ( euler.order === 'ZXY' ) {

			var ce = c * e, cf = c * f, de = d * e, df = d * f;

			te[ 0 ] = ce - df * b;
			te[ 4 ] = - a * f;
			te[ 8 ] = de + cf * b;

			te[ 1 ] = cf + de * b;
			te[ 5 ] = a * e;
			te[ 9 ] = df - ce * b;

			te[ 2 ] = - a * d;
			te[ 6 ] = b;
			te[ 10 ] = a * c;

		} else if ( euler.order === 'ZYX' ) {

			var ae = a * e, af = a * f, be = b * e, bf = b * f;

			te[ 0 ] = c * e;
			te[ 4 ] = be * d - af;
			te[ 8 ] = ae * d + bf;

			te[ 1 ] = c * f;
			te[ 5 ] = bf * d + ae;
			te[ 9 ] = af * d - be;

			te[ 2 ] = - d;
			te[ 6 ] = b * c;
			te[ 10 ] = a * c;

		} else if ( euler.order === 'YZX' ) {

			var ac = a * c, ad = a * d, bc = b * c, bd = b * d;

			te[ 0 ] = c * e;
			te[ 4 ] = bd - ac * f;
			te[ 8 ] = bc * f + ad;

			te[ 1 ] = f;
			te[ 5 ] = a * e;
			te[ 9 ] = - b * e;

			te[ 2 ] = - d * e;
			te[ 6 ] = ad * f + bc;
			te[ 10 ] = ac - bd * f;

		} else if ( euler.order === 'XZY' ) {

			var ac = a * c, ad = a * d, bc = b * c, bd = b * d;

			te[ 0 ] = c * e;
			te[ 4 ] = - f;
			te[ 8 ] = d * e;

			te[ 1 ] = ac * f + bd;
			te[ 5 ] = a * e;
			te[ 9 ] = ad * f - bc;

			te[ 2 ] = bc * f - ad;
			te[ 6 ] = b * e;
			te[ 10 ] = bd * f + ac;

		}

		// last column
		te[ 3 ] = 0;
		te[ 7 ] = 0;
		te[ 11 ] = 0;

		// bottom row
		te[ 12 ] = 0;
		te[ 13 ] = 0;
		te[ 14 ] = 0;
		te[ 15 ] = 1;

		return this;

	},

	makeRotationFromQuaternion: function ( q ) {

		var te = this.elements;

		var x = q.x, y = q.y, z = q.z, w = q.w;
		var x2 = x + x, y2 = y + y, z2 = z + z;
		var xx = x * x2, xy = x * y2, xz = x * z2;
		var yy = y * y2, yz = y * z2, zz = z * z2;
		var wx = w * x2, wy = w * y2, wz = w * z2;

		te[ 0 ] = 1 - ( yy + zz );
		te[ 4 ] = xy - wz;
		te[ 8 ] = xz + wy;

		te[ 1 ] = xy + wz;
		te[ 5 ] = 1 - ( xx + zz );
		te[ 9 ] = yz - wx;

		te[ 2 ] = xz - wy;
		te[ 6 ] = yz + wx;
		te[ 10 ] = 1 - ( xx + yy );

		// last column
		te[ 3 ] = 0;
		te[ 7 ] = 0;
		te[ 11 ] = 0;

		// bottom row
		te[ 12 ] = 0;
		te[ 13 ] = 0;
		te[ 14 ] = 0;
		te[ 15 ] = 1;

		return this;

	},

	lookAt: function () {

		var x, y, z;

		return function ( eye, target, up ) {

			if ( x === undefined ) {

				x = new THREE.Vector3();
				y = new THREE.Vector3();
				z = new THREE.Vector3();

			}

			var te = this.elements;

			z.subVectors( eye, target ).normalize();

			if ( z.lengthSq() === 0 ) {

				z.z = 1;

			}

			x.crossVectors( up, z ).normalize();

			if ( x.lengthSq() === 0 ) {

				z.z += 0.0001;
				x.crossVectors( up, z ).normalize();

			}

			y.crossVectors( z, x );


			te[ 0 ] = x.x; te[ 4 ] = y.x; te[ 8 ] = z.x;
			te[ 1 ] = x.y; te[ 5 ] = y.y; te[ 9 ] = z.y;
			te[ 2 ] = x.z; te[ 6 ] = y.z; te[ 10 ] = z.z;

			return this;

		};

	}(),

	multiply: function ( m, n ) {

		if ( n !== undefined ) {

			console.warn( 'THREE.Matrix4: .multiply() now only accepts one argument. Use .multiplyMatrices( a, b ) instead.' );
			return this.multiplyMatrices( m, n );

		}

		return this.multiplyMatrices( this, m );

	},

	premultiply: function ( m ) {

		return this.multiplyMatrices( m, this );

	},

	multiplyMatrices: function ( a, b ) {

		var ae = a.elements;
		var be = b.elements;
		var te = this.elements;

		var a11 = ae[ 0 ], a12 = ae[ 4 ], a13 = ae[ 8 ], a14 = ae[ 12 ];
		var a21 = ae[ 1 ], a22 = ae[ 5 ], a23 = ae[ 9 ], a24 = ae[ 13 ];
		var a31 = ae[ 2 ], a32 = ae[ 6 ], a33 = ae[ 10 ], a34 = ae[ 14 ];
		var a41 = ae[ 3 ], a42 = ae[ 7 ], a43 = ae[ 11 ], a44 = ae[ 15 ];

		var b11 = be[ 0 ], b12 = be[ 4 ], b13 = be[ 8 ], b14 = be[ 12 ];
		var b21 = be[ 1 ], b22 = be[ 5 ], b23 = be[ 9 ], b24 = be[ 13 ];
		var b31 = be[ 2 ], b32 = be[ 6 ], b33 = be[ 10 ], b34 = be[ 14 ];
		var b41 = be[ 3 ], b42 = be[ 7 ], b43 = be[ 11 ], b44 = be[ 15 ];

		te[ 0 ] = a11 * b11 + a12 * b21 + a13 * b31 + a14 * b41;
		te[ 4 ] = a11 * b12 + a12 * b22 + a13 * b32 + a14 * b42;
		te[ 8 ] = a11 * b13 + a12 * b23 + a13 * b33 + a14 * b43;
		te[ 12 ] = a11 * b14 + a12 * b24 + a13 * b34 + a14 * b44;

		te[ 1 ] = a21 * b11 + a22 * b21 + a23 * b31 + a24 * b41;
		te[ 5 ] = a21 * b12 + a22 * b22 + a23 * b32 + a24 * b42;
		te[ 9 ] = a21 * b13 + a22 * b23 + a23 * b33 + a24 * b43;
		te[ 13 ] = a21 * b14 + a22 * b24 + a23 * b34 + a24 * b44;

		te[ 2 ] = a31 * b11 + a32 * b21 + a33 * b31 + a34 * b41;
		te[ 6 ] = a31 * b12 + a32 * b22 + a33 * b32 + a34 * b42;
		te[ 10 ] = a31 * b13 + a32 * b23 + a33 * b33 + a34 * b43;
		te[ 14 ] = a31 * b14 + a32 * b24 + a33 * b34 + a34 * b44;

		te[ 3 ] = a41 * b11 + a42 * b21 + a43 * b31 + a44 * b41;
		te[ 7 ] = a41 * b12 + a42 * b22 + a43 * b32 + a44 * b42;
		te[ 11 ] = a41 * b13 + a42 * b23 + a43 * b33 + a44 * b43;
		te[ 15 ] = a41 * b14 + a42 * b24 + a43 * b34 + a44 * b44;

		return this;

	},

	multiplyToArray: function ( a, b, r ) {

		var te = this.elements;

		this.multiplyMatrices( a, b );

		r[ 0 ] = te[ 0 ]; r[ 1 ] = te[ 1 ]; r[ 2 ] = te[ 2 ]; r[ 3 ] = te[ 3 ];
		r[ 4 ] = te[ 4 ]; r[ 5 ] = te[ 5 ]; r[ 6 ] = te[ 6 ]; r[ 7 ] = te[ 7 ];
		r[ 8 ]  = te[ 8 ]; r[ 9 ]  = te[ 9 ]; r[ 10 ] = te[ 10 ]; r[ 11 ] = te[ 11 ];
		r[ 12 ] = te[ 12 ]; r[ 13 ] = te[ 13 ]; r[ 14 ] = te[ 14 ]; r[ 15 ] = te[ 15 ];

		return this;

	},

	multiplyScalar: function ( s ) {

		var te = this.elements;

		te[ 0 ] *= s; te[ 4 ] *= s; te[ 8 ] *= s; te[ 12 ] *= s;
		te[ 1 ] *= s; te[ 5 ] *= s; te[ 9 ] *= s; te[ 13 ] *= s;
		te[ 2 ] *= s; te[ 6 ] *= s; te[ 10 ] *= s; te[ 14 ] *= s;
		te[ 3 ] *= s; te[ 7 ] *= s; te[ 11 ] *= s; te[ 15 ] *= s;

		return this;

	},

	applyToVector3Array: function () {

		var v1;

		return function ( array, offset, length ) {

			if ( v1 === undefined ) v1 = new THREE.Vector3();
			if ( offset === undefined ) offset = 0;
			if ( length === undefined ) length = array.length;

			for ( var i = 0, j = offset; i < length; i += 3, j += 3 ) {

				v1.fromArray( array, j );
				v1.applyMatrix4( this );
				v1.toArray( array, j );

			}

			return array;

		};

	}(),

	applyToBuffer: function () {

		var v1;

		return function applyToBuffer( buffer, offset, length ) {

			if ( v1 === undefined ) v1 = new THREE.Vector3();
			if ( offset === undefined ) offset = 0;
			if ( length === undefined ) length = buffer.length / buffer.itemSize;

			for ( var i = 0, j = offset; i < length; i ++, j ++ ) {

				v1.x = buffer.getX( j );
				v1.y = buffer.getY( j );
				v1.z = buffer.getZ( j );

				v1.applyMatrix4( this );

				buffer.setXYZ( v1.x, v1.y, v1.z );

			}

			return buffer;

		};

	}(),

	determinant: function () {

		var te = this.elements;

		var n11 = te[ 0 ], n12 = te[ 4 ], n13 = te[ 8 ], n14 = te[ 12 ];
		var n21 = te[ 1 ], n22 = te[ 5 ], n23 = te[ 9 ], n24 = te[ 13 ];
		var n31 = te[ 2 ], n32 = te[ 6 ], n33 = te[ 10 ], n34 = te[ 14 ];
		var n41 = te[ 3 ], n42 = te[ 7 ], n43 = te[ 11 ], n44 = te[ 15 ];

		//TODO: make this more efficient
		//( based on http://www.euclideanspace.com/maths/algebra/matrix/functions/inverse/fourD/index.htm )

		return (
			n41 * (
				+ n14 * n23 * n32
				 - n13 * n24 * n32
				 - n14 * n22 * n33
				 + n12 * n24 * n33
				 + n13 * n22 * n34
				 - n12 * n23 * n34
			) +
			n42 * (
				+ n11 * n23 * n34
				 - n11 * n24 * n33
				 + n14 * n21 * n33
				 - n13 * n21 * n34
				 + n13 * n24 * n31
				 - n14 * n23 * n31
			) +
			n43 * (
				+ n11 * n24 * n32
				 - n11 * n22 * n34
				 - n14 * n21 * n32
				 + n12 * n21 * n34
				 + n14 * n22 * n31
				 - n12 * n24 * n31
			) +
			n44 * (
				- n13 * n22 * n31
				 - n11 * n23 * n32
				 + n11 * n22 * n33
				 + n13 * n21 * n32
				 - n12 * n21 * n33
				 + n12 * n23 * n31
			)

		);

	},

	transpose: function () {

		var te = this.elements;
		var tmp;

		tmp = te[ 1 ]; te[ 1 ] = te[ 4 ]; te[ 4 ] = tmp;
		tmp = te[ 2 ]; te[ 2 ] = te[ 8 ]; te[ 8 ] = tmp;
		tmp = te[ 6 ]; te[ 6 ] = te[ 9 ]; te[ 9 ] = tmp;

		tmp = te[ 3 ]; te[ 3 ] = te[ 12 ]; te[ 12 ] = tmp;
		tmp = te[ 7 ]; te[ 7 ] = te[ 13 ]; te[ 13 ] = tmp;
		tmp = te[ 11 ]; te[ 11 ] = te[ 14 ]; te[ 14 ] = tmp;

		return this;

	},

	flattenToArrayOffset: function ( array, offset ) {

		console.warn( "THREE.Matrix3: .flattenToArrayOffset is deprecated " +
				"- just use .toArray instead." );

		return this.toArray( array, offset );

	},

	getPosition: function () {

		var v1;

		return function () {

			if ( v1 === undefined ) v1 = new THREE.Vector3();
			console.warn( 'THREE.Matrix4: .getPosition() has been removed. Use Vector3.setFromMatrixPosition( matrix ) instead.' );

			return v1.setFromMatrixColumn( this, 3 );

		};

	}(),

	setPosition: function ( v ) {

		var te = this.elements;

		te[ 12 ] = v.x;
		te[ 13 ] = v.y;
		te[ 14 ] = v.z;

		return this;

	},

	getInverse: function ( m, throwOnDegenerate ) {

		// based on http://www.euclideanspace.com/maths/algebra/matrix/functions/inverse/fourD/index.htm
		var te = this.elements,
			me = m.elements,

			n11 = me[ 0 ], n21 = me[ 1 ], n31 = me[ 2 ], n41 = me[ 3 ],
			n12 = me[ 4 ], n22 = me[ 5 ], n32 = me[ 6 ], n42 = me[ 7 ],
			n13 = me[ 8 ], n23 = me[ 9 ], n33 = me[ 10 ], n43 = me[ 11 ],
			n14 = me[ 12 ], n24 = me[ 13 ], n34 = me[ 14 ], n44 = me[ 15 ],

			t11 = n23 * n34 * n42 - n24 * n33 * n42 + n24 * n32 * n43 - n22 * n34 * n43 - n23 * n32 * n44 + n22 * n33 * n44,
			t12 = n14 * n33 * n42 - n13 * n34 * n42 - n14 * n32 * n43 + n12 * n34 * n43 + n13 * n32 * n44 - n12 * n33 * n44,
			t13 = n13 * n24 * n42 - n14 * n23 * n42 + n14 * n22 * n43 - n12 * n24 * n43 - n13 * n22 * n44 + n12 * n23 * n44,
			t14 = n14 * n23 * n32 - n13 * n24 * n32 - n14 * n22 * n33 + n12 * n24 * n33 + n13 * n22 * n34 - n12 * n23 * n34;

		var det = n11 * t11 + n21 * t12 + n31 * t13 + n41 * t14;

		if ( det === 0 ) {

			var msg = "THREE.Matrix4.getInverse(): can't invert matrix, determinant is 0";

			if ( throwOnDegenerate || false ) {

				throw new Error( msg );

			} else {

				console.warn( msg );

			}

			return this.identity();

		}
		
		var detInv = 1 / det;

		te[ 0 ] = t11 * detInv;
		te[ 1 ] = ( n24 * n33 * n41 - n23 * n34 * n41 - n24 * n31 * n43 + n21 * n34 * n43 + n23 * n31 * n44 - n21 * n33 * n44 ) * detInv;
		te[ 2 ] = ( n22 * n34 * n41 - n24 * n32 * n41 + n24 * n31 * n42 - n21 * n34 * n42 - n22 * n31 * n44 + n21 * n32 * n44 ) * detInv;
		te[ 3 ] = ( n23 * n32 * n41 - n22 * n33 * n41 - n23 * n31 * n42 + n21 * n33 * n42 + n22 * n31 * n43 - n21 * n32 * n43 ) * detInv;

		te[ 4 ] = t12 * detInv;
		te[ 5 ] = ( n13 * n34 * n41 - n14 * n33 * n41 + n14 * n31 * n43 - n11 * n34 * n43 - n13 * n31 * n44 + n11 * n33 * n44 ) * detInv;
		te[ 6 ] = ( n14 * n32 * n41 - n12 * n34 * n41 - n14 * n31 * n42 + n11 * n34 * n42 + n12 * n31 * n44 - n11 * n32 * n44 ) * detInv;
		te[ 7 ] = ( n12 * n33 * n41 - n13 * n32 * n41 + n13 * n31 * n42 - n11 * n33 * n42 - n12 * n31 * n43 + n11 * n32 * n43 ) * detInv;

		te[ 8 ] = t13 * detInv;
		te[ 9 ] = ( n14 * n23 * n41 - n13 * n24 * n41 - n14 * n21 * n43 + n11 * n24 * n43 + n13 * n21 * n44 - n11 * n23 * n44 ) * detInv;
		te[ 10 ] = ( n12 * n24 * n41 - n14 * n22 * n41 + n14 * n21 * n42 - n11 * n24 * n42 - n12 * n21 * n44 + n11 * n22 * n44 ) * detInv;
		te[ 11 ] = ( n13 * n22 * n41 - n12 * n23 * n41 - n13 * n21 * n42 + n11 * n23 * n42 + n12 * n21 * n43 - n11 * n22 * n43 ) * detInv;

		te[ 12 ] = t14 * detInv;
		te[ 13 ] = ( n13 * n24 * n31 - n14 * n23 * n31 + n14 * n21 * n33 - n11 * n24 * n33 - n13 * n21 * n34 + n11 * n23 * n34 ) * detInv;
		te[ 14 ] = ( n14 * n22 * n31 - n12 * n24 * n31 - n14 * n21 * n32 + n11 * n24 * n32 + n12 * n21 * n34 - n11 * n22 * n34 ) * detInv;
		te[ 15 ] = ( n12 * n23 * n31 - n13 * n22 * n31 + n13 * n21 * n32 - n11 * n23 * n32 - n12 * n21 * n33 + n11 * n22 * n33 ) * detInv;

		return this;

	},

	scale: function ( v ) {

		var te = this.elements;
		var x = v.x, y = v.y, z = v.z;

		te[ 0 ] *= x; te[ 4 ] *= y; te[ 8 ] *= z;
		te[ 1 ] *= x; te[ 5 ] *= y; te[ 9 ] *= z;
		te[ 2 ] *= x; te[ 6 ] *= y; te[ 10 ] *= z;
		te[ 3 ] *= x; te[ 7 ] *= y; te[ 11 ] *= z;

		return this;

	},

	getMaxScaleOnAxis: function () {

		var te = this.elements;

		var scaleXSq = te[ 0 ] * te[ 0 ] + te[ 1 ] * te[ 1 ] + te[ 2 ] * te[ 2 ];
		var scaleYSq = te[ 4 ] * te[ 4 ] + te[ 5 ] * te[ 5 ] + te[ 6 ] * te[ 6 ];
		var scaleZSq = te[ 8 ] * te[ 8 ] + te[ 9 ] * te[ 9 ] + te[ 10 ] * te[ 10 ];

		return Math.sqrt( Math.max( scaleXSq, scaleYSq, scaleZSq ) );

	},

	makeTranslation: function ( x, y, z ) {

		this.set(

			1, 0, 0, x,
			0, 1, 0, y,
			0, 0, 1, z,
			0, 0, 0, 1

		);

		return this;

	},

	makeRotationX: function ( theta ) {

		var c = Math.cos( theta ), s = Math.sin( theta );

		this.set(

			1, 0,  0, 0,
			0, c, - s, 0,
			0, s,  c, 0,
			0, 0,  0, 1

		);

		return this;

	},

	makeRotationY: function ( theta ) {

		var c = Math.cos( theta ), s = Math.sin( theta );

		this.set(

			 c, 0, s, 0,
			 0, 1, 0, 0,
			- s, 0, c, 0,
			 0, 0, 0, 1

		);

		return this;

	},

	makeRotationZ: function ( theta ) {

		var c = Math.cos( theta ), s = Math.sin( theta );

		this.set(

			c, - s, 0, 0,
			s,  c, 0, 0,
			0,  0, 1, 0,
			0,  0, 0, 1

		);

		return this;

	},

	makeRotationAxis: function ( axis, angle ) {

		// Based on http://www.gamedev.net/reference/articles/article1199.asp

		var c = Math.cos( angle );
		var s = Math.sin( angle );
		var t = 1 - c;
		var x = axis.x, y = axis.y, z = axis.z;
		var tx = t * x, ty = t * y;

		this.set(

			tx * x + c, tx * y - s * z, tx * z + s * y, 0,
			tx * y + s * z, ty * y + c, ty * z - s * x, 0,
			tx * z - s * y, ty * z + s * x, t * z * z + c, 0,
			0, 0, 0, 1

		);

		 return this;

	},

	makeScale: function ( x, y, z ) {

		this.set(

			x, 0, 0, 0,
			0, y, 0, 0,
			0, 0, z, 0,
			0, 0, 0, 1

		);

		return this;

	},

	compose: function ( position, quaternion, scale ) {

		this.makeRotationFromQuaternion( quaternion );
		this.scale( scale );
		this.setPosition( position );

		return this;

	},

	decompose: function () {

		var vector, matrix;

		return function ( position, quaternion, scale ) {

			if ( vector === undefined ) {

				vector = new THREE.Vector3();
				matrix = new THREE.Matrix4();

			}

			var te = this.elements;

			var sx = vector.set( te[ 0 ], te[ 1 ], te[ 2 ] ).length();
			var sy = vector.set( te[ 4 ], te[ 5 ], te[ 6 ] ).length();
			var sz = vector.set( te[ 8 ], te[ 9 ], te[ 10 ] ).length();

			// if determine is negative, we need to invert one scale
			var det = this.determinant();
			if ( det < 0 ) {

				sx = - sx;

			}

			position.x = te[ 12 ];
			position.y = te[ 13 ];
			position.z = te[ 14 ];

			// scale the rotation part

			matrix.elements.set( this.elements ); // at this point matrix is incomplete so we can't use .copy()

			var invSX = 1 / sx;
			var invSY = 1 / sy;
			var invSZ = 1 / sz;

			matrix.elements[ 0 ] *= invSX;
			matrix.elements[ 1 ] *= invSX;
			matrix.elements[ 2 ] *= invSX;

			matrix.elements[ 4 ] *= invSY;
			matrix.elements[ 5 ] *= invSY;
			matrix.elements[ 6 ] *= invSY;

			matrix.elements[ 8 ] *= invSZ;
			matrix.elements[ 9 ] *= invSZ;
			matrix.elements[ 10 ] *= invSZ;

			quaternion.setFromRotationMatrix( matrix );

			scale.x = sx;
			scale.y = sy;
			scale.z = sz;

			return this;

		};

	}(),

	makeFrustum: function ( left, right, bottom, top, near, far ) {

		var te = this.elements;
		var x = 2 * near / ( right - left );
		var y = 2 * near / ( top - bottom );

		var a = ( right + left ) / ( right - left );
		var b = ( top + bottom ) / ( top - bottom );
		var c = - ( far + near ) / ( far - near );
		var d = - 2 * far * near / ( far - near );

		te[ 0 ] = x;	te[ 4 ] = 0;	te[ 8 ] = a;	te[ 12 ] = 0;
		te[ 1 ] = 0;	te[ 5 ] = y;	te[ 9 ] = b;	te[ 13 ] = 0;
		te[ 2 ] = 0;	te[ 6 ] = 0;	te[ 10 ] = c;	te[ 14 ] = d;
		te[ 3 ] = 0;	te[ 7 ] = 0;	te[ 11 ] = - 1;	te[ 15 ] = 0;

		return this;

	},

	makePerspective: function ( fov, aspect, near, far ) {

		var ymax = near * Math.tan( THREE.Math.DEG2RAD * fov * 0.5 );
		var ymin = - ymax;
		var xmin = ymin * aspect;
		var xmax = ymax * aspect;

		return this.makeFrustum( xmin, xmax, ymin, ymax, near, far );

	},

	makeOrthographic: function ( left, right, top, bottom, near, far ) {

		var te = this.elements;
		var w = 1.0 / ( right - left );
		var h = 1.0 / ( top - bottom );
		var p = 1.0 / ( far - near );

		var x = ( right + left ) * w;
		var y = ( top + bottom ) * h;
		var z = ( far + near ) * p;

		te[ 0 ] = 2 * w;	te[ 4 ] = 0;	te[ 8 ] = 0;	te[ 12 ] = - x;
		te[ 1 ] = 0;	te[ 5 ] = 2 * h;	te[ 9 ] = 0;	te[ 13 ] = - y;
		te[ 2 ] = 0;	te[ 6 ] = 0;	te[ 10 ] = - 2 * p;	te[ 14 ] = - z;
		te[ 3 ] = 0;	te[ 7 ] = 0;	te[ 11 ] = 0;	te[ 15 ] = 1;

		return this;

	},

	equals: function ( matrix ) {

		var te = this.elements;
		var me = matrix.elements;

		for ( var i = 0; i < 16; i ++ ) {

			if ( te[ i ] !== me[ i ] ) return false;

		}

		return true;

	},

	fromArray: function ( array ) {

		this.elements.set( array );

		return this;

	},

	toArray: function ( array, offset ) {

		if ( array === undefined ) array = [];
		if ( offset === undefined ) offset = 0;

		var te = this.elements;

		array[ offset ] = te[ 0 ];
		array[ offset + 1 ] = te[ 1 ];
		array[ offset + 2 ] = te[ 2 ];
		array[ offset + 3 ] = te[ 3 ];

		array[ offset + 4 ] = te[ 4 ];
		array[ offset + 5 ] = te[ 5 ];
		array[ offset + 6 ] = te[ 6 ];
		array[ offset + 7 ] = te[ 7 ];

		array[ offset + 8 ]  = te[ 8 ];
		array[ offset + 9 ]  = te[ 9 ];
		array[ offset + 10 ] = te[ 10 ];
		array[ offset + 11 ] = te[ 11 ];

		array[ offset + 12 ] = te[ 12 ];
		array[ offset + 13 ] = te[ 13 ];
		array[ offset + 14 ] = te[ 14 ];
		array[ offset + 15 ] = te[ 15 ];

		return array;

	}

};

// File:src/math/Ray.js

/**
 * @author bhouston / http://clara.io
 */

THREE.Ray = function ( origin, direction ) {

	this.origin = ( origin !== undefined ) ? origin : new THREE.Vector3();
	this.direction = ( direction !== undefined ) ? direction : new THREE.Vector3();

};

THREE.Ray.prototype = {

	constructor: THREE.Ray,

	set: function ( origin, direction ) {

		this.origin.copy( origin );
		this.direction.copy( direction );

		return this;

	},

	clone: function () {

		return new this.constructor().copy( this );

	},

	copy: function ( ray ) {

		this.origin.copy( ray.origin );
		this.direction.copy( ray.direction );

		return this;

	},

	at: function ( t, optionalTarget ) {

		var result = optionalTarget || new THREE.Vector3();

		return result.copy( this.direction ).multiplyScalar( t ).add( this.origin );

	},

	lookAt: function ( v ) {

		this.direction.copy( v ).sub( this.origin ).normalize();

		return this;

	},

	recast: function () {

		var v1 = new THREE.Vector3();

		return function ( t ) {

			this.origin.copy( this.at( t, v1 ) );

			return this;

		};

	}(),

	closestPointToPoint: function ( point, optionalTarget ) {

		var result = optionalTarget || new THREE.Vector3();
		result.subVectors( point, this.origin );
		var directionDistance = result.dot( this.direction );

		if ( directionDistance < 0 ) {

			return result.copy( this.origin );

		}

		return result.copy( this.direction ).multiplyScalar( directionDistance ).add( this.origin );

	},

	distanceToPoint: function ( point ) {

		return Math.sqrt( this.distanceSqToPoint( point ) );

	},

	distanceSqToPoint: function () {

		var v1 = new THREE.Vector3();

		return function ( point ) {

			var directionDistance = v1.subVectors( point, this.origin ).dot( this.direction );

			// point behind the ray

			if ( directionDistance < 0 ) {

				return this.origin.distanceToSquared( point );

			}

			v1.copy( this.direction ).multiplyScalar( directionDistance ).add( this.origin );

			return v1.distanceToSquared( point );

		};

	}(),

	distanceSqToSegment: function () {

		var segCenter = new THREE.Vector3();
		var segDir = new THREE.Vector3();
		var diff = new THREE.Vector3();

		return function ( v0, v1, optionalPointOnRay, optionalPointOnSegment ) {

			// from http://www.geometrictools.com/GTEngine/Include/Mathematics/GteDistRaySegment.h
			// It returns the min distance between the ray and the segment
			// defined by v0 and v1
			// It can also set two optional targets :
			// - The closest point on the ray
			// - The closest point on the segment

			segCenter.copy( v0 ).add( v1 ).multiplyScalar( 0.5 );
			segDir.copy( v1 ).sub( v0 ).normalize();
			diff.copy( this.origin ).sub( segCenter );

			var segExtent = v0.distanceTo( v1 ) * 0.5;
			var a01 = - this.direction.dot( segDir );
			var b0 = diff.dot( this.direction );
			var b1 = - diff.dot( segDir );
			var c = diff.lengthSq();
			var det = Math.abs( 1 - a01 * a01 );
			var s0, s1, sqrDist, extDet;

			if ( det > 0 ) {

				// The ray and segment are not parallel.

				s0 = a01 * b1 - b0;
				s1 = a01 * b0 - b1;
				extDet = segExtent * det;

				if ( s0 >= 0 ) {

					if ( s1 >= - extDet ) {

						if ( s1 <= extDet ) {

							// region 0
							// Minimum at interior points of ray and segment.

							var invDet = 1 / det;
							s0 *= invDet;
							s1 *= invDet;
							sqrDist = s0 * ( s0 + a01 * s1 + 2 * b0 ) + s1 * ( a01 * s0 + s1 + 2 * b1 ) + c;

						} else {

							// region 1

							s1 = segExtent;
							s0 = Math.max( 0, - ( a01 * s1 + b0 ) );
							sqrDist = - s0 * s0 + s1 * ( s1 + 2 * b1 ) + c;

						}

					} else {

						// region 5

						s1 = - segExtent;
						s0 = Math.max( 0, - ( a01 * s1 + b0 ) );
						sqrDist = - s0 * s0 + s1 * ( s1 + 2 * b1 ) + c;

					}

				} else {

					if ( s1 <= - extDet ) {

						// region 4

						s0 = Math.max( 0, - ( - a01 * segExtent + b0 ) );
						s1 = ( s0 > 0 ) ? - segExtent : Math.min( Math.max( - segExtent, - b1 ), segExtent );
						sqrDist = - s0 * s0 + s1 * ( s1 + 2 * b1 ) + c;

					} else if ( s1 <= extDet ) {

						// region 3

						s0 = 0;
						s1 = Math.min( Math.max( - segExtent, - b1 ), segExtent );
						sqrDist = s1 * ( s1 + 2 * b1 ) + c;

					} else {

						// region 2

						s0 = Math.max( 0, - ( a01 * segExtent + b0 ) );
						s1 = ( s0 > 0 ) ? segExtent : Math.min( Math.max( - segExtent, - b1 ), segExtent );
						sqrDist = - s0 * s0 + s1 * ( s1 + 2 * b1 ) + c;

					}

				}

			} else {

				// Ray and segment are parallel.

				s1 = ( a01 > 0 ) ? - segExtent : segExtent;
				s0 = Math.max( 0, - ( a01 * s1 + b0 ) );
				sqrDist = - s0 * s0 + s1 * ( s1 + 2 * b1 ) + c;

			}

			if ( optionalPointOnRay ) {

				optionalPointOnRay.copy( this.direction ).multiplyScalar( s0 ).add( this.origin );

			}

			if ( optionalPointOnSegment ) {

				optionalPointOnSegment.copy( segDir ).multiplyScalar( s1 ).add( segCenter );

			}

			return sqrDist;

		};

	}(),

	intersectSphere: function () {

		var v1 = new THREE.Vector3();

		return function ( sphere, optionalTarget ) {

			v1.subVectors( sphere.center, this.origin );
			var tca = v1.dot( this.direction );
			var d2 = v1.dot( v1 ) - tca * tca;
			var radius2 = sphere.radius * sphere.radius;

			if ( d2 > radius2 ) return null;

			var thc = Math.sqrt( radius2 - d2 );

			// t0 = first intersect point - entrance on front of sphere
			var t0 = tca - thc;

			// t1 = second intersect point - exit point on back of sphere
			var t1 = tca + thc;

			// test to see if both t0 and t1 are behind the ray - if so, return null
			if ( t0 < 0 && t1 < 0 ) return null;

			// test to see if t0 is behind the ray:
			// if it is, the ray is inside the sphere, so return the second exit point scaled by t1,
			// in order to always return an intersect point that is in front of the ray.
			if ( t0 < 0 ) return this.at( t1, optionalTarget );

			// else t0 is in front of the ray, so return the first collision point scaled by t0
			return this.at( t0, optionalTarget );

		};

	}(),

	intersectsSphere: function ( sphere ) {

		return this.distanceToPoint( sphere.center ) <= sphere.radius;

	},

	distanceToPlane: function ( plane ) {

		var denominator = plane.normal.dot( this.direction );

		if ( denominator === 0 ) {

			// line is coplanar, return origin
			if ( plane.distanceToPoint( this.origin ) === 0 ) {

				return 0;

			}

			// Null is preferable to undefined since undefined means.... it is undefined

			return null;

		}

		var t = - ( this.origin.dot( plane.normal ) + plane.constant ) / denominator;

		// Return if the ray never intersects the plane

		return t >= 0 ? t :  null;

	},

	intersectPlane: function ( plane, optionalTarget ) {

		var t = this.distanceToPlane( plane );

		if ( t === null ) {

			return null;

		}

		return this.at( t, optionalTarget );

	},


	intersectsPlane: function ( plane ) {

		// check if the ray lies on the plane first

		var distToPoint = plane.distanceToPoint( this.origin );

		if ( distToPoint === 0 ) {

			return true;

		}

		var denominator = plane.normal.dot( this.direction );

		if ( denominator * distToPoint < 0 ) {

			return true;

		}

		// ray origin is behind the plane (and is pointing behind it)

		return false;

	},

	intersectBox: function ( box, optionalTarget ) {

		var tmin, tmax, tymin, tymax, tzmin, tzmax;

		var invdirx = 1 / this.direction.x,
			invdiry = 1 / this.direction.y,
			invdirz = 1 / this.direction.z;

		var origin = this.origin;

		if ( invdirx >= 0 ) {

			tmin = ( box.min.x - origin.x ) * invdirx;
			tmax = ( box.max.x - origin.x ) * invdirx;

		} else {

			tmin = ( box.max.x - origin.x ) * invdirx;
			tmax = ( box.min.x - origin.x ) * invdirx;

		}

		if ( invdiry >= 0 ) {

			tymin = ( box.min.y - origin.y ) * invdiry;
			tymax = ( box.max.y - origin.y ) * invdiry;

		} else {

			tymin = ( box.max.y - origin.y ) * invdiry;
			tymax = ( box.min.y - origin.y ) * invdiry;

		}

		if ( ( tmin > tymax ) || ( tymin > tmax ) ) return null;

		// These lines also handle the case where tmin or tmax is NaN
		// (result of 0 * Infinity). x !== x returns true if x is NaN

		if ( tymin > tmin || tmin !== tmin ) tmin = tymin;

		if ( tymax < tmax || tmax !== tmax ) tmax = tymax;

		if ( invdirz >= 0 ) {

			tzmin = ( box.min.z - origin.z ) * invdirz;
			tzmax = ( box.max.z - origin.z ) * invdirz;

		} else {

			tzmin = ( box.max.z - origin.z ) * invdirz;
			tzmax = ( box.min.z - origin.z ) * invdirz;

		}

		if ( ( tmin > tzmax ) || ( tzmin > tmax ) ) return null;

		if ( tzmin > tmin || tmin !== tmin ) tmin = tzmin;

		if ( tzmax < tmax || tmax !== tmax ) tmax = tzmax;

		//return point closest to the ray (positive side)

		if ( tmax < 0 ) return null;

		return this.at( tmin >= 0 ? tmin : tmax, optionalTarget );

	},

	intersectsBox: ( function () {

		var v = new THREE.Vector3();

		return function ( box ) {

			return this.intersectBox( box, v ) !== null;

		};

	} )(),

	intersectTriangle: function () {

		// Compute the offset origin, edges, and normal.
		var diff = new THREE.Vector3();
		var edge1 = new THREE.Vector3();
		var edge2 = new THREE.Vector3();
		var normal = new THREE.Vector3();

		return function ( a, b, c, backfaceCulling, optionalTarget ) {

			// from http://www.geometrictools.com/GTEngine/Include/Mathematics/GteIntrRay3Triangle3.h

			edge1.subVectors( b, a );
			edge2.subVectors( c, a );
			normal.crossVectors( edge1, edge2 );

			// Solve Q + t*D = b1*E1 + b2*E2 (Q = kDiff, D = ray direction,
			// E1 = kEdge1, E2 = kEdge2, N = Cross(E1,E2)) by
			//   |Dot(D,N)|*b1 = sign(Dot(D,N))*Dot(D,Cross(Q,E2))
			//   |Dot(D,N)|*b2 = sign(Dot(D,N))*Dot(D,Cross(E1,Q))
			//   |Dot(D,N)|*t = -sign(Dot(D,N))*Dot(Q,N)
			var DdN = this.direction.dot( normal );
			var sign;

			if ( DdN > 0 ) {

				if ( backfaceCulling ) return null;
				sign = 1;

			} else if ( DdN < 0 ) {

				sign = - 1;
				DdN = - DdN;

			} else {

				return null;

			}

			diff.subVectors( this.origin, a );
			var DdQxE2 = sign * this.direction.dot( edge2.crossVectors( diff, edge2 ) );

			// b1 < 0, no intersection
			if ( DdQxE2 < 0 ) {

				return null;

			}

			var DdE1xQ = sign * this.direction.dot( edge1.cross( diff ) );

			// b2 < 0, no intersection
			if ( DdE1xQ < 0 ) {

				return null;

			}

			// b1+b2 > 1, no intersection
			if ( DdQxE2 + DdE1xQ > DdN ) {

				return null;

			}

			// Line intersects triangle, check if ray does.
			var QdN = - sign * diff.dot( normal );

			// t < 0, no intersection
			if ( QdN < 0 ) {

				return null;

			}

			// Ray intersects triangle.
			return this.at( QdN / DdN, optionalTarget );

		};

	}(),

	applyMatrix4: function ( matrix4 ) {

		this.direction.add( this.origin ).applyMatrix4( matrix4 );
		this.origin.applyMatrix4( matrix4 );
		this.direction.sub( this.origin );
		this.direction.normalize();

		return this;

	},

	equals: function ( ray ) {

		return ray.origin.equals( this.origin ) && ray.direction.equals( this.direction );

	}

};

// File:src/math/Sphere.js

/**
 * @author bhouston / http://clara.io
 * @author mrdoob / http://mrdoob.com/
 */

THREE.Sphere = function ( center, radius ) {

	this.center = ( center !== undefined ) ? center : new THREE.Vector3();
	this.radius = ( radius !== undefined ) ? radius : 0;

};

THREE.Sphere.prototype = {

	constructor: THREE.Sphere,

	set: function ( center, radius ) {

		this.center.copy( center );
		this.radius = radius;

		return this;

	},

	setFromPoints: function () {

		var box = new THREE.Box3();

		return function ( points, optionalCenter ) {

			var center = this.center;

			if ( optionalCenter !== undefined ) {

				center.copy( optionalCenter );

			} else {

				box.setFromPoints( points ).center( center );

			}

			var maxRadiusSq = 0;

			for ( var i = 0, il = points.length; i < il; i ++ ) {

				maxRadiusSq = Math.max( maxRadiusSq, center.distanceToSquared( points[ i ] ) );

			}

			this.radius = Math.sqrt( maxRadiusSq );

			return this;

		};

	}(),

	clone: function () {

		return new this.constructor().copy( this );

	},

	copy: function ( sphere ) {

		this.center.copy( sphere.center );
		this.radius = sphere.radius;

		return this;

	},

	empty: function () {

		return ( this.radius <= 0 );

	},

	containsPoint: function ( point ) {

		return ( point.distanceToSquared( this.center ) <= ( this.radius * this.radius ) );

	},

	distanceToPoint: function ( point ) {

		return ( point.distanceTo( this.center ) - this.radius );

	},

	intersectsSphere: function ( sphere ) {

		var radiusSum = this.radius + sphere.radius;

		return sphere.center.distanceToSquared( this.center ) <= ( radiusSum * radiusSum );

	},

	intersectsBox: function ( box ) {

		return box.intersectsSphere( this );

	},

	intersectsPlane: function ( plane ) {

		// We use the following equation to compute the signed distance from
		// the center of the sphere to the plane.
		//
		// distance = q * n - d
		//
		// If this distance is greater than the radius of the sphere,
		// then there is no intersection.

		return Math.abs( this.center.dot( plane.normal ) - plane.constant ) <= this.radius;

	},

	clampPoint: function ( point, optionalTarget ) {

		var deltaLengthSq = this.center.distanceToSquared( point );

		var result = optionalTarget || new THREE.Vector3();

		result.copy( point );

		if ( deltaLengthSq > ( this.radius * this.radius ) ) {

			result.sub( this.center ).normalize();
			result.multiplyScalar( this.radius ).add( this.center );

		}

		return result;

	},

	getBoundingBox: function ( optionalTarget ) {

		var box = optionalTarget || new THREE.Box3();

		box.set( this.center, this.center );
		box.expandByScalar( this.radius );

		return box;

	},

	applyMatrix4: function ( matrix ) {

		this.center.applyMatrix4( matrix );
		this.radius = this.radius * matrix.getMaxScaleOnAxis();

		return this;

	},

	translate: function ( offset ) {

		this.center.add( offset );

		return this;

	},

	equals: function ( sphere ) {

		return sphere.center.equals( this.center ) && ( sphere.radius === this.radius );

	}

};

// File:src/math/Frustum.js

/**
 * @author mrdoob / http://mrdoob.com/
 * @author alteredq / http://alteredqualia.com/
 * @author bhouston / http://clara.io
 */

THREE.Frustum = function ( p0, p1, p2, p3, p4, p5 ) {

	this.planes = [

		( p0 !== undefined ) ? p0 : new THREE.Plane(),
		( p1 !== undefined ) ? p1 : new THREE.Plane(),
		( p2 !== undefined ) ? p2 : new THREE.Plane(),
		( p3 !== undefined ) ? p3 : new THREE.Plane(),
		( p4 !== undefined ) ? p4 : new THREE.Plane(),
		( p5 !== undefined ) ? p5 : new THREE.Plane()

	];

};

THREE.Frustum.prototype = {

	constructor: THREE.Frustum,

	set: function ( p0, p1, p2, p3, p4, p5 ) {

		var planes = this.planes;

		planes[ 0 ].copy( p0 );
		planes[ 1 ].copy( p1 );
		planes[ 2 ].copy( p2 );
		planes[ 3 ].copy( p3 );
		planes[ 4 ].copy( p4 );
		planes[ 5 ].copy( p5 );

		return this;

	},

	clone: function () {

		return new this.constructor().copy( this );

	},

	copy: function ( frustum ) {

		var planes = this.planes;

		for ( var i = 0; i < 6; i ++ ) {

			planes[ i ].copy( frustum.planes[ i ] );

		}

		return this;

	},

	setFromMatrix: function ( m ) {

		var planes = this.planes;
		var me = m.elements;
		var me0 = me[ 0 ], me1 = me[ 1 ], me2 = me[ 2 ], me3 = me[ 3 ];
		var me4 = me[ 4 ], me5 = me[ 5 ], me6 = me[ 6 ], me7 = me[ 7 ];
		var me8 = me[ 8 ], me9 = me[ 9 ], me10 = me[ 10 ], me11 = me[ 11 ];
		var me12 = me[ 12 ], me13 = me[ 13 ], me14 = me[ 14 ], me15 = me[ 15 ];

		planes[ 0 ].setComponents( me3 - me0, me7 - me4, me11 - me8, me15 - me12 ).normalize();
		planes[ 1 ].setComponents( me3 + me0, me7 + me4, me11 + me8, me15 + me12 ).normalize();
		planes[ 2 ].setComponents( me3 + me1, me7 + me5, me11 + me9, me15 + me13 ).normalize();
		planes[ 3 ].setComponents( me3 - me1, me7 - me5, me11 - me9, me15 - me13 ).normalize();
		planes[ 4 ].setComponents( me3 - me2, me7 - me6, me11 - me10, me15 - me14 ).normalize();
		planes[ 5 ].setComponents( me3 + me2, me7 + me6, me11 + me10, me15 + me14 ).normalize();

		return this;

	},

	intersectsObject: function () {

		var sphere = new THREE.Sphere();

		return function ( object ) {

			var geometry = object.geometry;

			if ( geometry.boundingSphere === null )
				geometry.computeBoundingSphere();

			sphere.copy( geometry.boundingSphere )
				.applyMatrix4( object.matrixWorld );

			return this.intersectsSphere( sphere );

		};

	}(),

	intersectsSprite: function () {

		var sphere = new THREE.Sphere();

		return function ( sprite ) {

			sphere.center.set( 0, 0, 0 );
			sphere.radius = 0.7071067811865476;
			sphere.applyMatrix4( sprite.matrixWorld );

			return this.intersectsSphere( sphere );

		};

	}(),

	intersectsSphere: function ( sphere ) {

		var planes = this.planes;
		var center = sphere.center;
		var negRadius = - sphere.radius;

		for ( var i = 0; i < 6; i ++ ) {

			var distance = planes[ i ].distanceToPoint( center );

			if ( distance < negRadius ) {

				return false;

			}

		}

		return true;

	},

	intersectsBox: function () {

		var p1 = new THREE.Vector3(),
			p2 = new THREE.Vector3();

		return function ( box ) {

			var planes = this.planes;

			for ( var i = 0; i < 6 ; i ++ ) {

				var plane = planes[ i ];

				p1.x = plane.normal.x > 0 ? box.min.x : box.max.x;
				p2.x = plane.normal.x > 0 ? box.max.x : box.min.x;
				p1.y = plane.normal.y > 0 ? box.min.y : box.max.y;
				p2.y = plane.normal.y > 0 ? box.max.y : box.min.y;
				p1.z = plane.normal.z > 0 ? box.min.z : box.max.z;
				p2.z = plane.normal.z > 0 ? box.max.z : box.min.z;

				var d1 = plane.distanceToPoint( p1 );
				var d2 = plane.distanceToPoint( p2 );

				// if both outside plane, no intersection

				if ( d1 < 0 && d2 < 0 ) {

					return false;

				}

			}

			return true;

		};

	}(),


	containsPoint: function ( point ) {

		var planes = this.planes;

		for ( var i = 0; i < 6; i ++ ) {

			if ( planes[ i ].distanceToPoint( point ) < 0 ) {

				return false;

			}

		}

		return true;

	}

};

// File:src/math/Plane.js

/**
 * @author bhouston / http://clara.io
 */

THREE.Plane = function ( normal, constant ) {

	this.normal = ( normal !== undefined ) ? normal : new THREE.Vector3( 1, 0, 0 );
	this.constant = ( constant !== undefined ) ? constant : 0;

};

THREE.Plane.prototype = {

	constructor: THREE.Plane,

	set: function ( normal, constant ) {

		this.normal.copy( normal );
		this.constant = constant;

		return this;

	},

	setComponents: function ( x, y, z, w ) {

		this.normal.set( x, y, z );
		this.constant = w;

		return this;

	},

	setFromNormalAndCoplanarPoint: function ( normal, point ) {

		this.normal.copy( normal );
		this.constant = - point.dot( this.normal );	// must be this.normal, not normal, as this.normal is normalized

		return this;

	},

	setFromCoplanarPoints: function () {

		var v1 = new THREE.Vector3();
		var v2 = new THREE.Vector3();

		return function ( a, b, c ) {

			var normal = v1.subVectors( c, b ).cross( v2.subVectors( a, b ) ).normalize();

			// Q: should an error be thrown if normal is zero (e.g. degenerate plane)?

			this.setFromNormalAndCoplanarPoint( normal, a );

			return this;

		};

	}(),

	clone: function () {

		return new this.constructor().copy( this );

	},

	copy: function ( plane ) {

		this.normal.copy( plane.normal );
		this.constant = plane.constant;

		return this;

	},

	normalize: function () {

		// Note: will lead to a divide by zero if the plane is invalid.

		var inverseNormalLength = 1.0 / this.normal.length();
		this.normal.multiplyScalar( inverseNormalLength );
		this.constant *= inverseNormalLength;

		return this;

	},

	negate: function () {

		this.constant *= - 1;
		this.normal.negate();

		return this;

	},

	distanceToPoint: function ( point ) {

		return this.normal.dot( point ) + this.constant;

	},

	distanceToSphere: function ( sphere ) {

		return this.distanceToPoint( sphere.center ) - sphere.radius;

	},

	projectPoint: function ( point, optionalTarget ) {

		return this.orthoPoint( point, optionalTarget ).sub( point ).negate();

	},

	orthoPoint: function ( point, optionalTarget ) {

		var perpendicularMagnitude = this.distanceToPoint( point );

		var result = optionalTarget || new THREE.Vector3();
		return result.copy( this.normal ).multiplyScalar( perpendicularMagnitude );

	},

	intersectLine: function () {

		var v1 = new THREE.Vector3();

		return function ( line, optionalTarget ) {

			var result = optionalTarget || new THREE.Vector3();

			var direction = line.delta( v1 );

			var denominator = this.normal.dot( direction );

			if ( denominator === 0 ) {

				// line is coplanar, return origin
				if ( this.distanceToPoint( line.start ) === 0 ) {

					return result.copy( line.start );

				}

				// Unsure if this is the correct method to handle this case.
				return undefined;

			}

			var t = - ( line.start.dot( this.normal ) + this.constant ) / denominator;

			if ( t < 0 || t > 1 ) {

				return undefined;

			}

			return result.copy( direction ).multiplyScalar( t ).add( line.start );

		};

	}(),

	intersectsLine: function ( line ) {

		// Note: this tests if a line intersects the plane, not whether it (or its end-points) are coplanar with it.

		var startSign = this.distanceToPoint( line.start );
		var endSign = this.distanceToPoint( line.end );

		return ( startSign < 0 && endSign > 0 ) || ( endSign < 0 && startSign > 0 );

	},

	intersectsBox: function ( box ) {

		return box.intersectsPlane( this );

	},

	intersectsSphere: function ( sphere ) {

		return sphere.intersectsPlane( this );

	},

	coplanarPoint: function ( optionalTarget ) {

		var result = optionalTarget || new THREE.Vector3();
		return result.copy( this.normal ).multiplyScalar( - this.constant );

	},

	applyMatrix4: function () {

		var v1 = new THREE.Vector3();
		var m1 = new THREE.Matrix3();

		return function ( matrix, optionalNormalMatrix ) {

			var referencePoint = this.coplanarPoint( v1 ).applyMatrix4( matrix );

			// transform normal based on theory here:
			// http://www.songho.ca/opengl/gl_normaltransform.html
			var normalMatrix = optionalNormalMatrix || m1.getNormalMatrix( matrix );
			var normal = this.normal.applyMatrix3( normalMatrix ).normalize();

			// recalculate constant (like in setFromNormalAndCoplanarPoint)
			this.constant = - referencePoint.dot( normal );

			return this;

		};

	}(),

	translate: function ( offset ) {

		this.constant = this.constant - offset.dot( this.normal );

		return this;

	},

	equals: function ( plane ) {

		return plane.normal.equals( this.normal ) && ( plane.constant === this.constant );

	}

};

// File:src/math/Spherical.js

/**
 * @author bhouston / http://clara.io
 * @author WestLangley / http://github.com/WestLangley
 *
 * Ref: https://en.wikipedia.org/wiki/Spherical_coordinate_system
 *
 * The poles (phi) are at the positive and negative y axis.
 * The equator starts at positive z.
 */

THREE.Spherical = function ( radius, phi, theta ) {

	this.radius = ( radius !== undefined ) ? radius : 1.0;
	this.phi = ( phi !== undefined ) ? phi : 0; // up / down towards top and bottom pole
	this.theta = ( theta !== undefined ) ? theta : 0; // around the equator of the sphere

	return this;

};

THREE.Spherical.prototype = {

	constructor: THREE.Spherical,

	set: function ( radius, phi, theta ) {

		this.radius = radius;
		this.phi = phi;
		this.theta = theta;

		return this;

	},

	clone: function () {

		return new this.constructor().copy( this );

	},

	copy: function ( other ) {

		this.radius.copy( other.radius );
		this.phi.copy( other.phi );
		this.theta.copy( other.theta );

		return this;

	},

	// restrict phi to be betwee EPS and PI-EPS
	makeSafe: function() {

		var EPS = 0.000001;
		this.phi = Math.max( EPS, Math.min( Math.PI - EPS, this.phi ) );

		return this;

	},

	setFromVector3: function( vec3 ) {

		this.radius = vec3.length();

		if ( this.radius === 0 ) {

			this.theta = 0;
			this.phi = 0;

		} else {

			this.theta = Math.atan2( vec3.x, vec3.z ); // equator angle around y-up axis
			this.phi = Math.acos( THREE.Math.clamp( vec3.y / this.radius, - 1, 1 ) ); // polar angle

		}

		return this;

	},

};

// File:src/math/Math.js

/**
 * @author alteredq / http://alteredqualia.com/
 * @author mrdoob / http://mrdoob.com/
 */

THREE.Math = {

	DEG2RAD: Math.PI / 180,
	RAD2DEG: 180 / Math.PI,

	generateUUID: function () {

		// http://www.broofa.com/Tools/Math.uuid.htm

		var chars = '0123456789ABCDEFGHIJKLMNOPQRSTUVWXYZabcdefghijklmnopqrstuvwxyz'.split( '' );
		var uuid = new Array( 36 );
		var rnd = 0, r;

		return function () {

			for ( var i = 0; i < 36; i ++ ) {

				if ( i === 8 || i === 13 || i === 18 || i === 23 ) {

					uuid[ i ] = '-';

				} else if ( i === 14 ) {

					uuid[ i ] = '4';

				} else {

					if ( rnd <= 0x02 ) rnd = 0x2000000 + ( Math.random() * 0x1000000 ) | 0;
					r = rnd & 0xf;
					rnd = rnd >> 4;
					uuid[ i ] = chars[ ( i === 19 ) ? ( r & 0x3 ) | 0x8 : r ];

				}

			}

			return uuid.join( '' );

		};

	}(),

	clamp: function ( value, min, max ) {

		return Math.max( min, Math.min( max, value ) );

	},

	// compute euclidian modulo of m % n
	// https://en.wikipedia.org/wiki/Modulo_operation

	euclideanModulo: function ( n, m ) {

		return ( ( n % m ) + m ) % m;

	},

	// Linear mapping from range <a1, a2> to range <b1, b2>

	mapLinear: function ( x, a1, a2, b1, b2 ) {

		return b1 + ( x - a1 ) * ( b2 - b1 ) / ( a2 - a1 );

	},

	// http://en.wikipedia.org/wiki/Smoothstep

	smoothstep: function ( x, min, max ) {

		if ( x <= min ) return 0;
		if ( x >= max ) return 1;

		x = ( x - min ) / ( max - min );

		return x * x * ( 3 - 2 * x );

	},

	smootherstep: function ( x, min, max ) {

		if ( x <= min ) return 0;
		if ( x >= max ) return 1;

		x = ( x - min ) / ( max - min );

		return x * x * x * ( x * ( x * 6 - 15 ) + 10 );

	},

	random16: function () {

		console.warn( 'THREE.Math.random16() has been deprecated. Use Math.random() instead.' );
		return Math.random();

	},

	// Random integer from <low, high> interval

	randInt: function ( low, high ) {

		return low + Math.floor( Math.random() * ( high - low + 1 ) );

	},

	// Random float from <low, high> interval

	randFloat: function ( low, high ) {

		return low + Math.random() * ( high - low );

	},

	// Random float from <-range/2, range/2> interval

	randFloatSpread: function ( range ) {

		return range * ( 0.5 - Math.random() );

	},

	degToRad: function ( degrees ) {

		return degrees * THREE.Math.DEG2RAD;

	},

	radToDeg: function ( radians ) {

		return radians * THREE.Math.RAD2DEG;

	},

	isPowerOfTwo: function ( value ) {

		return ( value & ( value - 1 ) ) === 0 && value !== 0;

	},

	nearestPowerOfTwo: function ( value ) {

		return Math.pow( 2, Math.round( Math.log( value ) / Math.LN2 ) );

	},

	nextPowerOfTwo: function ( value ) {

		value --;
		value |= value >> 1;
		value |= value >> 2;
		value |= value >> 4;
		value |= value >> 8;
		value |= value >> 16;
		value ++;

		return value;

	}

};

// File:src/math/Spline.js

/**
 * Spline from Tween.js, slightly optimized (and trashed)
 * http://sole.github.com/tween.js/examples/05_spline.html
 *
 * @author mrdoob / http://mrdoob.com/
 * @author alteredq / http://alteredqualia.com/
 */

THREE.Spline = function ( points ) {

	this.points = points;

	var c = [], v3 = { x: 0, y: 0, z: 0 },
	point, intPoint, weight, w2, w3,
	pa, pb, pc, pd;

	this.initFromArray = function ( a ) {

		this.points = [];

		for ( var i = 0; i < a.length; i ++ ) {

			this.points[ i ] = { x: a[ i ][ 0 ], y: a[ i ][ 1 ], z: a[ i ][ 2 ] };

		}

	};

	this.getPoint = function ( k ) {

		point = ( this.points.length - 1 ) * k;
		intPoint = Math.floor( point );
		weight = point - intPoint;

		c[ 0 ] = intPoint === 0 ? intPoint : intPoint - 1;
		c[ 1 ] = intPoint;
		c[ 2 ] = intPoint  > this.points.length - 2 ? this.points.length - 1 : intPoint + 1;
		c[ 3 ] = intPoint  > this.points.length - 3 ? this.points.length - 1 : intPoint + 2;

		pa = this.points[ c[ 0 ] ];
		pb = this.points[ c[ 1 ] ];
		pc = this.points[ c[ 2 ] ];
		pd = this.points[ c[ 3 ] ];

		w2 = weight * weight;
		w3 = weight * w2;

		v3.x = interpolate( pa.x, pb.x, pc.x, pd.x, weight, w2, w3 );
		v3.y = interpolate( pa.y, pb.y, pc.y, pd.y, weight, w2, w3 );
		v3.z = interpolate( pa.z, pb.z, pc.z, pd.z, weight, w2, w3 );

		return v3;

	};

	this.getControlPointsArray = function () {

		var i, p, l = this.points.length,
			coords = [];

		for ( i = 0; i < l; i ++ ) {

			p = this.points[ i ];
			coords[ i ] = [ p.x, p.y, p.z ];

		}

		return coords;

	};

	// approximate length by summing linear segments

	this.getLength = function ( nSubDivisions ) {

		var i, index, nSamples, position,
			point = 0, intPoint = 0, oldIntPoint = 0,
			oldPosition = new THREE.Vector3(),
			tmpVec = new THREE.Vector3(),
			chunkLengths = [],
			totalLength = 0;

		// first point has 0 length

		chunkLengths[ 0 ] = 0;

		if ( ! nSubDivisions ) nSubDivisions = 100;

		nSamples = this.points.length * nSubDivisions;

		oldPosition.copy( this.points[ 0 ] );

		for ( i = 1; i < nSamples; i ++ ) {

			index = i / nSamples;

			position = this.getPoint( index );
			tmpVec.copy( position );

			totalLength += tmpVec.distanceTo( oldPosition );

			oldPosition.copy( position );

			point = ( this.points.length - 1 ) * index;
			intPoint = Math.floor( point );

			if ( intPoint !== oldIntPoint ) {

				chunkLengths[ intPoint ] = totalLength;
				oldIntPoint = intPoint;

			}

		}

		// last point ends with total length

		chunkLengths[ chunkLengths.length ] = totalLength;

		return { chunks: chunkLengths, total: totalLength };

	};

	this.reparametrizeByArcLength = function ( samplingCoef ) {

		var i, j,
			index, indexCurrent, indexNext,
			realDistance,
			sampling, position,
			newpoints = [],
			tmpVec = new THREE.Vector3(),
			sl = this.getLength();

		newpoints.push( tmpVec.copy( this.points[ 0 ] ).clone() );

		for ( i = 1; i < this.points.length; i ++ ) {

			//tmpVec.copy( this.points[ i - 1 ] );
			//linearDistance = tmpVec.distanceTo( this.points[ i ] );

			realDistance = sl.chunks[ i ] - sl.chunks[ i - 1 ];

			sampling = Math.ceil( samplingCoef * realDistance / sl.total );

			indexCurrent = ( i - 1 ) / ( this.points.length - 1 );
			indexNext = i / ( this.points.length - 1 );

			for ( j = 1; j < sampling - 1; j ++ ) {

				index = indexCurrent + j * ( 1 / sampling ) * ( indexNext - indexCurrent );

				position = this.getPoint( index );
				newpoints.push( tmpVec.copy( position ).clone() );

			}

			newpoints.push( tmpVec.copy( this.points[ i ] ).clone() );

		}

		this.points = newpoints;

	};

	// Catmull-Rom

	function interpolate( p0, p1, p2, p3, t, t2, t3 ) {

		var v0 = ( p2 - p0 ) * 0.5,
			v1 = ( p3 - p1 ) * 0.5;

		return ( 2 * ( p1 - p2 ) + v0 + v1 ) * t3 + ( - 3 * ( p1 - p2 ) - 2 * v0 - v1 ) * t2 + v0 * t + p1;

	}

};

// File:src/math/Triangle.js

/**
 * @author bhouston / http://clara.io
 * @author mrdoob / http://mrdoob.com/
 */

THREE.Triangle = function ( a, b, c ) {

	this.a = ( a !== undefined ) ? a : new THREE.Vector3();
	this.b = ( b !== undefined ) ? b : new THREE.Vector3();
	this.c = ( c !== undefined ) ? c : new THREE.Vector3();

};

THREE.Triangle.normal = function () {

	var v0 = new THREE.Vector3();

	return function ( a, b, c, optionalTarget ) {

		var result = optionalTarget || new THREE.Vector3();

		result.subVectors( c, b );
		v0.subVectors( a, b );
		result.cross( v0 );

		var resultLengthSq = result.lengthSq();
		if ( resultLengthSq > 0 ) {

			return result.multiplyScalar( 1 / Math.sqrt( resultLengthSq ) );

		}

		return result.set( 0, 0, 0 );

	};

}();

// static/instance method to calculate barycentric coordinates
// based on: http://www.blackpawn.com/texts/pointinpoly/default.html
THREE.Triangle.barycoordFromPoint = function () {

	var v0 = new THREE.Vector3();
	var v1 = new THREE.Vector3();
	var v2 = new THREE.Vector3();

	return function ( point, a, b, c, optionalTarget ) {

		v0.subVectors( c, a );
		v1.subVectors( b, a );
		v2.subVectors( point, a );

		var dot00 = v0.dot( v0 );
		var dot01 = v0.dot( v1 );
		var dot02 = v0.dot( v2 );
		var dot11 = v1.dot( v1 );
		var dot12 = v1.dot( v2 );

		var denom = ( dot00 * dot11 - dot01 * dot01 );

		var result = optionalTarget || new THREE.Vector3();

		// collinear or singular triangle
		if ( denom === 0 ) {

			// arbitrary location outside of triangle?
			// not sure if this is the best idea, maybe should be returning undefined
			return result.set( - 2, - 1, - 1 );

		}

		var invDenom = 1 / denom;
		var u = ( dot11 * dot02 - dot01 * dot12 ) * invDenom;
		var v = ( dot00 * dot12 - dot01 * dot02 ) * invDenom;

		// barycentric coordinates must always sum to 1
		return result.set( 1 - u - v, v, u );

	};

}();

THREE.Triangle.containsPoint = function () {

	var v1 = new THREE.Vector3();

	return function ( point, a, b, c ) {

		var result = THREE.Triangle.barycoordFromPoint( point, a, b, c, v1 );

		return ( result.x >= 0 ) && ( result.y >= 0 ) && ( ( result.x + result.y ) <= 1 );

	};

}();

THREE.Triangle.prototype = {

	constructor: THREE.Triangle,

	set: function ( a, b, c ) {

		this.a.copy( a );
		this.b.copy( b );
		this.c.copy( c );

		return this;

	},

	setFromPointsAndIndices: function ( points, i0, i1, i2 ) {

		this.a.copy( points[ i0 ] );
		this.b.copy( points[ i1 ] );
		this.c.copy( points[ i2 ] );

		return this;

	},

	clone: function () {

		return new this.constructor().copy( this );

	},

	copy: function ( triangle ) {

		this.a.copy( triangle.a );
		this.b.copy( triangle.b );
		this.c.copy( triangle.c );

		return this;

	},

	area: function () {

		var v0 = new THREE.Vector3();
		var v1 = new THREE.Vector3();

		return function () {

			v0.subVectors( this.c, this.b );
			v1.subVectors( this.a, this.b );

			return v0.cross( v1 ).length() * 0.5;

		};

	}(),

	midpoint: function ( optionalTarget ) {

		var result = optionalTarget || new THREE.Vector3();
		return result.addVectors( this.a, this.b ).add( this.c ).multiplyScalar( 1 / 3 );

	},

	normal: function ( optionalTarget ) {

		return THREE.Triangle.normal( this.a, this.b, this.c, optionalTarget );

	},

	plane: function ( optionalTarget ) {

		var result = optionalTarget || new THREE.Plane();

		return result.setFromCoplanarPoints( this.a, this.b, this.c );

	},

	barycoordFromPoint: function ( point, optionalTarget ) {

		return THREE.Triangle.barycoordFromPoint( point, this.a, this.b, this.c, optionalTarget );

	},

	containsPoint: function ( point ) {

		return THREE.Triangle.containsPoint( point, this.a, this.b, this.c );

	},

	closestPointToPoint: function () {

		var plane, edgeList, projectedPoint, closestPoint;

		return function closestPointToPoint( point, optionalTarget ) {

			if ( plane === undefined ) {

				plane = new THREE.Plane();
				edgeList = [ new THREE.Line3(), new THREE.Line3(), new THREE.Line3() ];
				projectedPoint = new THREE.Vector3();
				closestPoint = new THREE.Vector3();

			}

			var result = optionalTarget || new THREE.Vector3();
			var minDistance = Infinity;

			// project the point onto the plane of the triangle

			plane.setFromCoplanarPoints( this.a, this.b, this.c );
			plane.projectPoint( point, projectedPoint );

			// check if the projection lies within the triangle

			if( this.containsPoint( projectedPoint ) === true ) {

				// if so, this is the closest point

				result.copy( projectedPoint );

			} else {

				// if not, the point falls outside the triangle. the result is the closest point to the triangle's edges or vertices

				edgeList[ 0 ].set( this.a, this.b );
				edgeList[ 1 ].set( this.b, this.c );
				edgeList[ 2 ].set( this.c, this.a );

				for( var i = 0; i < edgeList.length; i ++ ) {

					edgeList[ i ].closestPointToPoint( projectedPoint, true, closestPoint );

					var distance = projectedPoint.distanceToSquared( closestPoint );

					if( distance < minDistance ) {

						minDistance = distance;

						result.copy( closestPoint );

					}

				}

			}

			return result;

		};

	}(),

	equals: function ( triangle ) {

		return triangle.a.equals( this.a ) && triangle.b.equals( this.b ) && triangle.c.equals( this.c );

	}

};

// File:src/math/Interpolant.js

/**
 * Abstract base class of interpolants over parametric samples.
 *
 * The parameter domain is one dimensional, typically the time or a path
 * along a curve defined by the data.
 *
 * The sample values can have any dimensionality and derived classes may
 * apply special interpretations to the data.
 *
 * This class provides the interval seek in a Template Method, deferring
 * the actual interpolation to derived classes.
 *
 * Time complexity is O(1) for linear access crossing at most two points
 * and O(log N) for random access, where N is the number of positions.
 *
 * References:
 *
 * 		http://www.oodesign.com/template-method-pattern.html
 *
 * @author tschw
 */

THREE.Interpolant = function(
		parameterPositions, sampleValues, sampleSize, resultBuffer ) {

	this.parameterPositions = parameterPositions;
	this._cachedIndex = 0;

	this.resultBuffer = resultBuffer !== undefined ?
			resultBuffer : new sampleValues.constructor( sampleSize );
	this.sampleValues = sampleValues;
	this.valueSize = sampleSize;

};

THREE.Interpolant.prototype = {

	constructor: THREE.Interpolant,

	evaluate: function( t ) {

		var pp = this.parameterPositions,
			i1 = this._cachedIndex,

			t1 = pp[   i1   ],
			t0 = pp[ i1 - 1 ];

		validate_interval: {

			seek: {

				var right;

				linear_scan: {
//- See http://jsperf.com/comparison-to-undefined/3
//- slower code:
//-
//- 				if ( t >= t1 || t1 === undefined ) {
					forward_scan: if ( ! ( t < t1 ) ) {

						for ( var giveUpAt = i1 + 2; ;) {

							if ( t1 === undefined ) {

								if ( t < t0 ) break forward_scan;

								// after end

								i1 = pp.length;
								this._cachedIndex = i1;
								return this.afterEnd_( i1 - 1, t, t0 );

							}

							if ( i1 === giveUpAt ) break; // this loop

							t0 = t1;
							t1 = pp[ ++ i1 ];

							if ( t < t1 ) {

								// we have arrived at the sought interval
								break seek;

							}

						}

						// prepare binary search on the right side of the index
						right = pp.length;
						break linear_scan;

					}

//- slower code:
//-					if ( t < t0 || t0 === undefined ) {
					if ( ! ( t >= t0 ) ) {

						// looping?

						var t1global = pp[ 1 ];

						if ( t < t1global ) {

							i1 = 2; // + 1, using the scan for the details
							t0 = t1global;

						}

						// linear reverse scan

						for ( var giveUpAt = i1 - 2; ;) {

							if ( t0 === undefined ) {

								// before start

								this._cachedIndex = 0;
								return this.beforeStart_( 0, t, t1 );

							}

							if ( i1 === giveUpAt ) break; // this loop

							t1 = t0;
							t0 = pp[ -- i1 - 1 ];

							if ( t >= t0 ) {

								// we have arrived at the sought interval
								break seek;

							}

						}

						// prepare binary search on the left side of the index
						right = i1;
						i1 = 0;
						break linear_scan;

					}

					// the interval is valid

					break validate_interval;

				} // linear scan

				// binary search

				while ( i1 < right ) {

					var mid = ( i1 + right ) >>> 1;

					if ( t < pp[ mid ] ) {

						right = mid;

					} else {

						i1 = mid + 1;

					}

				}

				t1 = pp[   i1   ];
				t0 = pp[ i1 - 1 ];

				// check boundary cases, again

				if ( t0 === undefined ) {

					this._cachedIndex = 0;
					return this.beforeStart_( 0, t, t1 );

				}

				if ( t1 === undefined ) {

					i1 = pp.length;
					this._cachedIndex = i1;
					return this.afterEnd_( i1 - 1, t0, t );

				}

			} // seek

			this._cachedIndex = i1;

			this.intervalChanged_( i1, t0, t1 );

		} // validate_interval

		return this.interpolate_( i1, t0, t, t1 );

	},

	settings: null, // optional, subclass-specific settings structure
	// Note: The indirection allows central control of many interpolants.

	// --- Protected interface

	DefaultSettings_: {},

	getSettings_: function() {

		return this.settings || this.DefaultSettings_;

	},

	copySampleValue_: function( index ) {

		// copies a sample value to the result buffer

		var result = this.resultBuffer,
			values = this.sampleValues,
			stride = this.valueSize,
			offset = index * stride;

		for ( var i = 0; i !== stride; ++ i ) {

			result[ i ] = values[ offset + i ];

		}

		return result;

	},

	// Template methods for derived classes:

	interpolate_: function( i1, t0, t, t1 ) {

		throw new Error( "call to abstract method" );
		// implementations shall return this.resultBuffer

	},

	intervalChanged_: function( i1, t0, t1 ) {

		// empty

	}

};

Object.assign( THREE.Interpolant.prototype, {

	beforeStart_: //( 0, t, t0 ), returns this.resultBuffer
		THREE.Interpolant.prototype.copySampleValue_,

	afterEnd_: //( N-1, tN-1, t ), returns this.resultBuffer
		THREE.Interpolant.prototype.copySampleValue_

} );

// File:src/math/interpolants/CubicInterpolant.js

/**
 * Fast and simple cubic spline interpolant.
 *
 * It was derived from a Hermitian construction setting the first derivative
 * at each sample position to the linear slope between neighboring positions
 * over their parameter interval.
 *
 * @author tschw
 */

THREE.CubicInterpolant = function(
		parameterPositions, sampleValues, sampleSize, resultBuffer ) {

	THREE.Interpolant.call(
			this, parameterPositions, sampleValues, sampleSize, resultBuffer );

	this._weightPrev = -0;
	this._offsetPrev = -0;
	this._weightNext = -0;
	this._offsetNext = -0;

};

THREE.CubicInterpolant.prototype =
		Object.assign( Object.create( THREE.Interpolant.prototype ), {

	constructor: THREE.CubicInterpolant,

	DefaultSettings_: {

		endingStart: 	THREE.ZeroCurvatureEnding,
		endingEnd:		THREE.ZeroCurvatureEnding

	},

	intervalChanged_: function( i1, t0, t1 ) {

		var pp = this.parameterPositions,
			iPrev = i1 - 2,
			iNext = i1 + 1,

			tPrev = pp[ iPrev ],
			tNext = pp[ iNext ];

		if ( tPrev === undefined ) {

			switch ( this.getSettings_().endingStart ) {

				case THREE.ZeroSlopeEnding:

					// f'(t0) = 0
					iPrev = i1;
					tPrev = 2 * t0 - t1;

					break;

				case THREE.WrapAroundEnding:

					// use the other end of the curve
					iPrev = pp.length - 2;
					tPrev = t0 + pp[ iPrev ] - pp[ iPrev + 1 ];

					break;

				default: // ZeroCurvatureEnding

					// f''(t0) = 0 a.k.a. Natural Spline
					iPrev = i1;
					tPrev = t1;

			}

		}

		if ( tNext === undefined ) {

			switch ( this.getSettings_().endingEnd ) {

				case THREE.ZeroSlopeEnding:

					// f'(tN) = 0
					iNext = i1;
					tNext = 2 * t1 - t0;

					break;

				case THREE.WrapAroundEnding:

					// use the other end of the curve
					iNext = 1;
					tNext = t1 + pp[ 1 ] - pp[ 0 ];

					break;

				default: // ZeroCurvatureEnding

					// f''(tN) = 0, a.k.a. Natural Spline
					iNext = i1 - 1;
					tNext = t0;

			}

		}

		var halfDt = ( t1 - t0 ) * 0.5,
			stride = this.valueSize;

		this._weightPrev = halfDt / ( t0 - tPrev );
		this._weightNext = halfDt / ( tNext - t1 );
		this._offsetPrev = iPrev * stride;
		this._offsetNext = iNext * stride;

	},

	interpolate_: function( i1, t0, t, t1 ) {

		var result = this.resultBuffer,
			values = this.sampleValues,
			stride = this.valueSize,

			o1 = i1 * stride,		o0 = o1 - stride,
			oP = this._offsetPrev, 	oN = this._offsetNext,
			wP = this._weightPrev,	wN = this._weightNext,

			p = ( t - t0 ) / ( t1 - t0 ),
			pp = p * p,
			ppp = pp * p;

		// evaluate polynomials

		var sP =     - wP   * ppp   +         2 * wP    * pp    -          wP   * p;
		var s0 = ( 1 + wP ) * ppp   + (-1.5 - 2 * wP )  * pp    + ( -0.5 + wP ) * p     + 1;
		var s1 = (-1 - wN ) * ppp   + ( 1.5 +   wN   )  * pp    +    0.5        * p;
		var sN =       wN   * ppp   -           wN      * pp;

		// combine data linearly

		for ( var i = 0; i !== stride; ++ i ) {

			result[ i ] =
					sP * values[ oP + i ] +
					s0 * values[ o0 + i ] +
					s1 * values[ o1 + i ] +
					sN * values[ oN + i ];

		}

		return result;

	}

} );

// File:src/math/interpolants/DiscreteInterpolant.js

/**
 *
 * Interpolant that evaluates to the sample value at the position preceeding
 * the parameter.
 *
 * @author tschw
 */

THREE.DiscreteInterpolant = function(
		parameterPositions, sampleValues, sampleSize, resultBuffer ) {

	THREE.Interpolant.call(
			this, parameterPositions, sampleValues, sampleSize, resultBuffer );

};

THREE.DiscreteInterpolant.prototype =
		Object.assign( Object.create( THREE.Interpolant.prototype ), {

	constructor: THREE.DiscreteInterpolant,

	interpolate_: function( i1, t0, t, t1 ) {

		return this.copySampleValue_( i1 - 1 );

	}

} );

// File:src/math/interpolants/LinearInterpolant.js

/**
 * @author tschw
 */

THREE.LinearInterpolant = function(
		parameterPositions, sampleValues, sampleSize, resultBuffer ) {

	THREE.Interpolant.call(
			this, parameterPositions, sampleValues, sampleSize, resultBuffer );

};

THREE.LinearInterpolant.prototype =
		Object.assign( Object.create( THREE.Interpolant.prototype ), {

	constructor: THREE.LinearInterpolant,

	interpolate_: function( i1, t0, t, t1 ) {

		var result = this.resultBuffer,
			values = this.sampleValues,
			stride = this.valueSize,

			offset1 = i1 * stride,
			offset0 = offset1 - stride,

			weight1 = ( t - t0 ) / ( t1 - t0 ),
			weight0 = 1 - weight1;

		for ( var i = 0; i !== stride; ++ i ) {

			result[ i ] =
					values[ offset0 + i ] * weight0 +
					values[ offset1 + i ] * weight1;

		}

		return result;

	}

} );

// File:src/math/interpolants/QuaternionLinearInterpolant.js

/**
 * Spherical linear unit quaternion interpolant.
 *
 * @author tschw
 */

THREE.QuaternionLinearInterpolant = function(
		parameterPositions, sampleValues, sampleSize, resultBuffer ) {

	THREE.Interpolant.call(
			this, parameterPositions, sampleValues, sampleSize, resultBuffer );

};

THREE.QuaternionLinearInterpolant.prototype =
		Object.assign( Object.create( THREE.Interpolant.prototype ), {

	constructor: THREE.QuaternionLinearInterpolant,

	interpolate_: function( i1, t0, t, t1 ) {

		var result = this.resultBuffer,
			values = this.sampleValues,
			stride = this.valueSize,

			offset = i1 * stride,

			alpha = ( t - t0 ) / ( t1 - t0 );

		for ( var end = offset + stride; offset !== end; offset += 4 ) {

			THREE.Quaternion.slerpFlat( result, 0,
					values, offset - stride, values, offset, alpha );

		}

		return result;

	}

} );

// File:src/core/Clock.js

/**
 * @author alteredq / http://alteredqualia.com/
 */

THREE.Clock = function ( autoStart ) {

	this.autoStart = ( autoStart !== undefined ) ? autoStart : true;

	this.startTime = 0;
	this.oldTime = 0;
	this.elapsedTime = 0;

	this.running = false;

};

THREE.Clock.prototype = {

	constructor: THREE.Clock,

	start: function () {

		this.startTime = ( performance || Date ).now();

		this.oldTime = this.startTime;
		this.running = true;

	},

	stop: function () {

		this.getElapsedTime();
		this.running = false;

	},

	getElapsedTime: function () {

		this.getDelta();
		return this.elapsedTime;

	},

	getDelta: function () {

		var diff = 0;

		if ( this.autoStart && ! this.running ) {

			this.start();

		}

		if ( this.running ) {

			var newTime = ( performance || Date ).now();

			diff = ( newTime - this.oldTime ) / 1000;
			this.oldTime = newTime;

			this.elapsedTime += diff;

		}

		return diff;

	}

};

// File:src/core/EventDispatcher.js

/**
 * https://github.com/mrdoob/eventdispatcher.js/
 */

THREE.EventDispatcher = function () {};

Object.assign( THREE.EventDispatcher.prototype, {

	addEventListener: function ( type, listener ) {

		if ( this._listeners === undefined ) this._listeners = {};

		var listeners = this._listeners;

		if ( listeners[ type ] === undefined ) {

			listeners[ type ] = [];

		}

		if ( listeners[ type ].indexOf( listener ) === - 1 ) {

			listeners[ type ].push( listener );

		}

	},

	hasEventListener: function ( type, listener ) {

		if ( this._listeners === undefined ) return false;

		var listeners = this._listeners;

		if ( listeners[ type ] !== undefined && listeners[ type ].indexOf( listener ) !== - 1 ) {

			return true;

		}

		return false;

	},

	removeEventListener: function ( type, listener ) {

		if ( this._listeners === undefined ) return;

		var listeners = this._listeners;
		var listenerArray = listeners[ type ];

		if ( listenerArray !== undefined ) {

			var index = listenerArray.indexOf( listener );

			if ( index !== - 1 ) {

				listenerArray.splice( index, 1 );

			}

		}

	},

	dispatchEvent: function ( event ) {

		if ( this._listeners === undefined ) return;

		var listeners = this._listeners;
		var listenerArray = listeners[ event.type ];

		if ( listenerArray !== undefined ) {

			event.target = this;

			var array = [], i = 0;
			var length = listenerArray.length;

			for ( i = 0; i < length; i ++ ) {

				array[ i ] = listenerArray[ i ];

			}

			for ( i = 0; i < length; i ++ ) {

				array[ i ].call( this, event );

			}

		}

	}

} );

// File:src/core/Layers.js

/**
 * @author mrdoob / http://mrdoob.com/
 */

THREE.Layers = function () {

	this.mask = 1;

};

THREE.Layers.prototype = {

	constructor: THREE.Layers,

	set: function ( channel ) {

		this.mask = 1 << channel;

	},

	enable: function ( channel ) {

		this.mask |= 1 << channel;

	},

	toggle: function ( channel ) {

		this.mask ^= 1 << channel;

	},

	disable: function ( channel ) {

		this.mask &= ~ ( 1 << channel );

	},

	test: function ( layers ) {

		return ( this.mask & layers.mask ) !== 0;

	}

};

// File:src/core/Raycaster.js

/**
 * @author mrdoob / http://mrdoob.com/
 * @author bhouston / http://clara.io/
 * @author stephomi / http://stephaneginier.com/
 */

( function ( THREE ) {

	THREE.Raycaster = function ( origin, direction, near, far ) {

		this.ray = new THREE.Ray( origin, direction );
		// direction is assumed to be normalized (for accurate distance calculations)

		this.near = near || 0;
		this.far = far || Infinity;

		this.params = {
			Mesh: {},
			Line: {},
			LOD: {},
			Points: { threshold: 1 },
			Sprite: {}
		};

		Object.defineProperties( this.params, {
			PointCloud: {
				get: function () {
					console.warn( 'THREE.Raycaster: params.PointCloud has been renamed to params.Points.' );
					return this.Points;
				}
			}
		} );

	};

	function ascSort( a, b ) {

		return a.distance - b.distance;

	}

	function intersectObject( object, raycaster, intersects, recursive ) {

		if ( object.visible === false ) return;

		object.raycast( raycaster, intersects );

		if ( recursive === true ) {

			var children = object.children;

			for ( var i = 0, l = children.length; i < l; i ++ ) {

				intersectObject( children[ i ], raycaster, intersects, true );

			}

		}

	}

	//

	THREE.Raycaster.prototype = {

		constructor: THREE.Raycaster,

		linePrecision: 1,

		set: function ( origin, direction ) {

			// direction is assumed to be normalized (for accurate distance calculations)

			this.ray.set( origin, direction );

		},

		setFromCamera: function ( coords, camera ) {

			if ( camera instanceof THREE.PerspectiveCamera ) {

				this.ray.origin.setFromMatrixPosition( camera.matrixWorld );
				this.ray.direction.set( coords.x, coords.y, 0.5 ).unproject( camera ).sub( this.ray.origin ).normalize();

			} else if ( camera instanceof THREE.OrthographicCamera ) {

				this.ray.origin.set( coords.x, coords.y, - 1 ).unproject( camera );
				this.ray.direction.set( 0, 0, - 1 ).transformDirection( camera.matrixWorld );

			} else {

				console.error( 'THREE.Raycaster: Unsupported camera type.' );

			}

		},

		intersectObject: function ( object, recursive ) {

			var intersects = [];

			intersectObject( object, this, intersects, recursive );

			intersects.sort( ascSort );

			return intersects;

		},

		intersectObjects: function ( objects, recursive ) {

			var intersects = [];

			if ( Array.isArray( objects ) === false ) {

				console.warn( 'THREE.Raycaster.intersectObjects: objects is not an Array.' );
				return intersects;

			}

			for ( var i = 0, l = objects.length; i < l; i ++ ) {

				intersectObject( objects[ i ], this, intersects, recursive );

			}

			intersects.sort( ascSort );

			return intersects;

		}

	};

}( THREE ) );

// File:src/core/Object3D.js

/**
 * @author mrdoob / http://mrdoob.com/
 * @author mikael emtinger / http://gomo.se/
 * @author alteredq / http://alteredqualia.com/
 * @author WestLangley / http://github.com/WestLangley
 * @author elephantatwork / www.elephantatwork.ch
 */

THREE.Object3D = function () {

	Object.defineProperty( this, 'id', { value: THREE.Object3DIdCount ++ } );

	this.uuid = THREE.Math.generateUUID();

	this.name = '';
	this.type = 'Object3D';

	this.parent = null;
	this.children = [];

	this.up = THREE.Object3D.DefaultUp.clone();

	var position = new THREE.Vector3();
	var rotation = new THREE.Euler();
	var quaternion = new THREE.Quaternion();
	var scale = new THREE.Vector3( 1, 1, 1 );

	function onRotationChange() {

		quaternion.setFromEuler( rotation, false );

	}

	function onQuaternionChange() {

		rotation.setFromQuaternion( quaternion, undefined, false );

	}

	rotation.onChange( onRotationChange );
	quaternion.onChange( onQuaternionChange );

	Object.defineProperties( this, {
		position: {
			enumerable: true,
			value: position
		},
		rotation: {
			enumerable: true,
			value: rotation
		},
		quaternion: {
			enumerable: true,
			value: quaternion
		},
		scale: {
			enumerable: true,
			value: scale
		},
		modelViewMatrix: {
			value: new THREE.Matrix4()
		},
		normalMatrix: {
			value: new THREE.Matrix3()
		}
	} );

	this.matrix = new THREE.Matrix4();
	this.matrixWorld = new THREE.Matrix4();

	this.matrixAutoUpdate = THREE.Object3D.DefaultMatrixAutoUpdate;
	this.matrixWorldNeedsUpdate = false;

	this.layers = new THREE.Layers();
	this.visible = true;

	this.castShadow = false;
	this.receiveShadow = false;

	this.frustumCulled = true;
	this.renderOrder = 0;

	this.userData = {};

};

THREE.Object3D.DefaultUp = new THREE.Vector3( 0, 1, 0 );
THREE.Object3D.DefaultMatrixAutoUpdate = true;

Object.assign( THREE.Object3D.prototype, THREE.EventDispatcher.prototype, {

	applyMatrix: function ( matrix ) {

		this.matrix.multiplyMatrices( matrix, this.matrix );

		this.matrix.decompose( this.position, this.quaternion, this.scale );

	},

	setRotationFromAxisAngle: function ( axis, angle ) {

		// assumes axis is normalized

		this.quaternion.setFromAxisAngle( axis, angle );

	},

	setRotationFromEuler: function ( euler ) {

		this.quaternion.setFromEuler( euler, true );

	},

	setRotationFromMatrix: function ( m ) {

		// assumes the upper 3x3 of m is a pure rotation matrix (i.e, unscaled)

		this.quaternion.setFromRotationMatrix( m );

	},

	setRotationFromQuaternion: function ( q ) {

		// assumes q is normalized

		this.quaternion.copy( q );

	},

	rotateOnAxis: function () {

		// rotate object on axis in object space
		// axis is assumed to be normalized

		var q1 = new THREE.Quaternion();

		return function ( axis, angle ) {

			q1.setFromAxisAngle( axis, angle );

			this.quaternion.multiply( q1 );

			return this;

		};

	}(),

	rotateX: function () {

		var v1 = new THREE.Vector3( 1, 0, 0 );

		return function ( angle ) {

			return this.rotateOnAxis( v1, angle );

		};

	}(),

	rotateY: function () {

		var v1 = new THREE.Vector3( 0, 1, 0 );

		return function ( angle ) {

			return this.rotateOnAxis( v1, angle );

		};

	}(),

	rotateZ: function () {

		var v1 = new THREE.Vector3( 0, 0, 1 );

		return function ( angle ) {

			return this.rotateOnAxis( v1, angle );

		};

	}(),

	translateOnAxis: function () {

		// translate object by distance along axis in object space
		// axis is assumed to be normalized

		var v1 = new THREE.Vector3();

		return function ( axis, distance ) {

			v1.copy( axis ).applyQuaternion( this.quaternion );

			this.position.add( v1.multiplyScalar( distance ) );

			return this;

		};

	}(),

	translateX: function () {

		var v1 = new THREE.Vector3( 1, 0, 0 );

		return function ( distance ) {

			return this.translateOnAxis( v1, distance );

		};

	}(),

	translateY: function () {

		var v1 = new THREE.Vector3( 0, 1, 0 );

		return function ( distance ) {

			return this.translateOnAxis( v1, distance );

		};

	}(),

	translateZ: function () {

		var v1 = new THREE.Vector3( 0, 0, 1 );

		return function ( distance ) {

			return this.translateOnAxis( v1, distance );

		};

	}(),

	localToWorld: function ( vector ) {

		return vector.applyMatrix4( this.matrixWorld );

	},

	worldToLocal: function () {

		var m1 = new THREE.Matrix4();

		return function ( vector ) {

			return vector.applyMatrix4( m1.getInverse( this.matrixWorld ) );

		};

	}(),

	lookAt: function () {

		// This routine does not support objects with rotated and/or translated parent(s)

		var m1 = new THREE.Matrix4();

		return function ( vector ) {

			m1.lookAt( vector, this.position, this.up );

			this.quaternion.setFromRotationMatrix( m1 );

		};

	}(),

	add: function ( object ) {

		if ( arguments.length > 1 ) {

			for ( var i = 0; i < arguments.length; i ++ ) {

				this.add( arguments[ i ] );

			}

			return this;

		}

		if ( object === this ) {

			console.error( "THREE.Object3D.add: object can't be added as a child of itself.", object );
			return this;

		}

		if ( object instanceof THREE.Object3D ) {

			if ( object.parent !== null ) {

				object.parent.remove( object );

			}

			object.parent = this;
			object.dispatchEvent( { type: 'added' } );

			this.children.push( object );

		} else {

			console.error( "THREE.Object3D.add: object not an instance of THREE.Object3D.", object );

		}

		return this;

	},

	remove: function ( object ) {

		if ( arguments.length > 1 ) {

			for ( var i = 0; i < arguments.length; i ++ ) {

				this.remove( arguments[ i ] );

			}

		}

		var index = this.children.indexOf( object );

		if ( index !== - 1 ) {

			object.parent = null;

			object.dispatchEvent( { type: 'removed' } );

			this.children.splice( index, 1 );

		}

	},

	getObjectById: function ( id ) {

		return this.getObjectByProperty( 'id', id );

	},

	getObjectByName: function ( name ) {

		return this.getObjectByProperty( 'name', name );

	},

	getObjectByProperty: function ( name, value ) {

		if ( this[ name ] === value ) return this;

		for ( var i = 0, l = this.children.length; i < l; i ++ ) {

			var child = this.children[ i ];
			var object = child.getObjectByProperty( name, value );

			if ( object !== undefined ) {

				return object;

			}

		}

		return undefined;

	},

	getWorldPosition: function ( optionalTarget ) {

		var result = optionalTarget || new THREE.Vector3();

		this.updateMatrixWorld( true );

		return result.setFromMatrixPosition( this.matrixWorld );

	},

	getWorldQuaternion: function () {

		var position = new THREE.Vector3();
		var scale = new THREE.Vector3();

		return function ( optionalTarget ) {

			var result = optionalTarget || new THREE.Quaternion();

			this.updateMatrixWorld( true );

			this.matrixWorld.decompose( position, result, scale );

			return result;

		};

	}(),

	getWorldRotation: function () {

		var quaternion = new THREE.Quaternion();

		return function ( optionalTarget ) {

			var result = optionalTarget || new THREE.Euler();

			this.getWorldQuaternion( quaternion );

			return result.setFromQuaternion( quaternion, this.rotation.order, false );

		};

	}(),

	getWorldScale: function () {

		var position = new THREE.Vector3();
		var quaternion = new THREE.Quaternion();

		return function ( optionalTarget ) {

			var result = optionalTarget || new THREE.Vector3();

			this.updateMatrixWorld( true );

			this.matrixWorld.decompose( position, quaternion, result );

			return result;

		};

	}(),

	getWorldDirection: function () {

		var quaternion = new THREE.Quaternion();

		return function ( optionalTarget ) {

			var result = optionalTarget || new THREE.Vector3();

			this.getWorldQuaternion( quaternion );

			return result.set( 0, 0, 1 ).applyQuaternion( quaternion );

		};

	}(),

	raycast: function () {},

	traverse: function ( callback ) {

		callback( this );

		var children = this.children;

		for ( var i = 0, l = children.length; i < l; i ++ ) {

			children[ i ].traverse( callback );

		}

	},

	traverseVisible: function ( callback ) {

		if ( this.visible === false ) return;

		callback( this );

		var children = this.children;

		for ( var i = 0, l = children.length; i < l; i ++ ) {

			children[ i ].traverseVisible( callback );

		}

	},

	traverseAncestors: function ( callback ) {

		var parent = this.parent;

		if ( parent !== null ) {

			callback( parent );

			parent.traverseAncestors( callback );

		}

	},

	updateMatrix: function () {

		this.matrix.compose( this.position, this.quaternion, this.scale );

		this.matrixWorldNeedsUpdate = true;

	},

	updateMatrixWorld: function ( force ) {

		if ( this.matrixAutoUpdate === true ) this.updateMatrix();

		if ( this.matrixWorldNeedsUpdate === true || force === true ) {

			if ( this.parent === null ) {

				this.matrixWorld.copy( this.matrix );

			} else {

				this.matrixWorld.multiplyMatrices( this.parent.matrixWorld, this.matrix );

			}

			this.matrixWorldNeedsUpdate = false;

			force = true;

		}

		// update children

		for ( var i = 0, l = this.children.length; i < l; i ++ ) {

			this.children[ i ].updateMatrixWorld( force );

		}

	},

	toJSON: function ( meta ) {

		// meta is '' when called from JSON.stringify
		var isRootObject = ( meta === undefined || meta === '' );

		var output = {};

		// meta is a hash used to collect geometries, materials.
		// not providing it implies that this is the root object
		// being serialized.
		if ( isRootObject ) {

			// initialize meta obj
			meta = {
				geometries: {},
				materials: {},
				textures: {},
				images: {}
			};

			output.metadata = {
				version: 4.4,
				type: 'Object',
				generator: 'Object3D.toJSON'
			};

		}

		// standard Object3D serialization

		var object = {};

		object.uuid = this.uuid;
		object.type = this.type;

		if ( this.name !== '' ) object.name = this.name;
		if ( JSON.stringify( this.userData ) !== '{}' ) object.userData = this.userData;
		if ( this.castShadow === true ) object.castShadow = true;
		if ( this.receiveShadow === true ) object.receiveShadow = true;
		if ( this.visible === false ) object.visible = false;

		object.matrix = this.matrix.toArray();

		//

		if ( this.geometry !== undefined ) {

			if ( meta.geometries[ this.geometry.uuid ] === undefined ) {

				meta.geometries[ this.geometry.uuid ] = this.geometry.toJSON( meta );

			}

			object.geometry = this.geometry.uuid;

		}

		if ( this.material !== undefined ) {

			if ( meta.materials[ this.material.uuid ] === undefined ) {

				meta.materials[ this.material.uuid ] = this.material.toJSON( meta );

			}

			object.material = this.material.uuid;

		}

		//

		if ( this.children.length > 0 ) {

			object.children = [];

			for ( var i = 0; i < this.children.length; i ++ ) {

				object.children.push( this.children[ i ].toJSON( meta ).object );

			}

		}

		if ( isRootObject ) {

			var geometries = extractFromCache( meta.geometries );
			var materials = extractFromCache( meta.materials );
			var textures = extractFromCache( meta.textures );
			var images = extractFromCache( meta.images );

			if ( geometries.length > 0 ) output.geometries = geometries;
			if ( materials.length > 0 ) output.materials = materials;
			if ( textures.length > 0 ) output.textures = textures;
			if ( images.length > 0 ) output.images = images;

		}

		output.object = object;

		return output;

		// extract data from the cache hash
		// remove metadata on each item
		// and return as array
		function extractFromCache ( cache ) {

			var values = [];
			for ( var key in cache ) {

				var data = cache[ key ];
				delete data.metadata;
				values.push( data );

			}
			return values;

		}

	},

	clone: function ( recursive ) {

		return new this.constructor().copy( this, recursive );

	},

	copy: function ( source, recursive ) {

		if ( recursive === undefined ) recursive = true;

		this.name = source.name;

		this.up.copy( source.up );

		this.position.copy( source.position );
		this.quaternion.copy( source.quaternion );
		this.scale.copy( source.scale );

		this.matrix.copy( source.matrix );
		this.matrixWorld.copy( source.matrixWorld );

		this.matrixAutoUpdate = source.matrixAutoUpdate;
		this.matrixWorldNeedsUpdate = source.matrixWorldNeedsUpdate;

		this.visible = source.visible;

		this.castShadow = source.castShadow;
		this.receiveShadow = source.receiveShadow;

		this.frustumCulled = source.frustumCulled;
		this.renderOrder = source.renderOrder;

		this.userData = JSON.parse( JSON.stringify( source.userData ) );

		if ( recursive === true ) {

			for ( var i = 0; i < source.children.length; i ++ ) {

				var child = source.children[ i ];
				this.add( child.clone() );

			}

		}

		return this;

	}

} );

THREE.Object3DIdCount = 0;

// File:src/core/Face3.js

/**
 * @author mrdoob / http://mrdoob.com/
 * @author alteredq / http://alteredqualia.com/
 */

THREE.Face3 = function ( a, b, c, normal, color, materialIndex ) {

	this.a = a;
	this.b = b;
	this.c = c;

	this.normal = normal instanceof THREE.Vector3 ? normal : new THREE.Vector3();
	this.vertexNormals = Array.isArray( normal ) ? normal : [];

	this.color = color instanceof THREE.Color ? color : new THREE.Color();
	this.vertexColors = Array.isArray( color ) ? color : [];

	this.materialIndex = materialIndex !== undefined ? materialIndex : 0;

};

THREE.Face3.prototype = {

	constructor: THREE.Face3,

	clone: function () {

		return new this.constructor().copy( this );

	},

	copy: function ( source ) {

		this.a = source.a;
		this.b = source.b;
		this.c = source.c;

		this.normal.copy( source.normal );
		this.color.copy( source.color );

		this.materialIndex = source.materialIndex;

		for ( var i = 0, il = source.vertexNormals.length; i < il; i ++ ) {

			this.vertexNormals[ i ] = source.vertexNormals[ i ].clone();

		}

		for ( var i = 0, il = source.vertexColors.length; i < il; i ++ ) {

			this.vertexColors[ i ] = source.vertexColors[ i ].clone();

		}

		return this;

	}

};

// File:src/core/BufferAttribute.js

/**
 * @author mrdoob / http://mrdoob.com/
 */

THREE.BufferAttribute = function ( array, itemSize, normalized ) {

	this.uuid = THREE.Math.generateUUID();

	this.array = array;
	this.itemSize = itemSize;

	this.dynamic = false;
	this.updateRange = { offset: 0, count: - 1 };

	this.version = 0;
	this.normalized = normalized === true;

};

THREE.BufferAttribute.prototype = {

	constructor: THREE.BufferAttribute,

	get count() {

		return this.array.length / this.itemSize;

	},

	set needsUpdate( value ) {

		if ( value === true ) this.version ++;

	},

	setDynamic: function ( value ) {

		this.dynamic = value;

		return this;

	},

	copy: function ( source ) {

		this.array = new source.array.constructor( source.array );
		this.itemSize = source.itemSize;

		this.dynamic = source.dynamic;

		return this;

	},

	copyAt: function ( index1, attribute, index2 ) {

		index1 *= this.itemSize;
		index2 *= attribute.itemSize;

		for ( var i = 0, l = this.itemSize; i < l; i ++ ) {

			this.array[ index1 + i ] = attribute.array[ index2 + i ];

		}

		return this;

	},

	copyArray: function ( array ) {

		this.array.set( array );

		return this;

	},

	copyColorsArray: function ( colors ) {

		var array = this.array, offset = 0;

		for ( var i = 0, l = colors.length; i < l; i ++ ) {

			var color = colors[ i ];

			if ( color === undefined ) {

				console.warn( 'THREE.BufferAttribute.copyColorsArray(): color is undefined', i );
				color = new THREE.Color();

			}

			array[ offset ++ ] = color.r;
			array[ offset ++ ] = color.g;
			array[ offset ++ ] = color.b;

		}

		return this;

	},

	copyIndicesArray: function ( indices ) {

		var array = this.array, offset = 0;

		for ( var i = 0, l = indices.length; i < l; i ++ ) {

			var index = indices[ i ];

			array[ offset ++ ] = index.a;
			array[ offset ++ ] = index.b;
			array[ offset ++ ] = index.c;

		}

		return this;

	},

	copyVector2sArray: function ( vectors ) {

		var array = this.array, offset = 0;

		for ( var i = 0, l = vectors.length; i < l; i ++ ) {

			var vector = vectors[ i ];

			if ( vector === undefined ) {

				console.warn( 'THREE.BufferAttribute.copyVector2sArray(): vector is undefined', i );
				vector = new THREE.Vector2();

			}

			array[ offset ++ ] = vector.x;
			array[ offset ++ ] = vector.y;

		}

		return this;

	},

	copyVector3sArray: function ( vectors ) {

		var array = this.array, offset = 0;

		for ( var i = 0, l = vectors.length; i < l; i ++ ) {

			var vector = vectors[ i ];

			if ( vector === undefined ) {

				console.warn( 'THREE.BufferAttribute.copyVector3sArray(): vector is undefined', i );
				vector = new THREE.Vector3();

			}

			array[ offset ++ ] = vector.x;
			array[ offset ++ ] = vector.y;
			array[ offset ++ ] = vector.z;

		}

		return this;

	},

	copyVector4sArray: function ( vectors ) {

		var array = this.array, offset = 0;

		for ( var i = 0, l = vectors.length; i < l; i ++ ) {

			var vector = vectors[ i ];

			if ( vector === undefined ) {

				console.warn( 'THREE.BufferAttribute.copyVector4sArray(): vector is undefined', i );
				vector = new THREE.Vector4();

			}

			array[ offset ++ ] = vector.x;
			array[ offset ++ ] = vector.y;
			array[ offset ++ ] = vector.z;
			array[ offset ++ ] = vector.w;

		}

		return this;

	},

	set: function ( value, offset ) {

		if ( offset === undefined ) offset = 0;

		this.array.set( value, offset );

		return this;

	},

	getX: function ( index ) {

		return this.array[ index * this.itemSize ];

	},

	setX: function ( index, x ) {

		this.array[ index * this.itemSize ] = x;

		return this;

	},

	getY: function ( index ) {

		return this.array[ index * this.itemSize + 1 ];

	},

	setY: function ( index, y ) {

		this.array[ index * this.itemSize + 1 ] = y;

		return this;

	},

	getZ: function ( index ) {

		return this.array[ index * this.itemSize + 2 ];

	},

	setZ: function ( index, z ) {

		this.array[ index * this.itemSize + 2 ] = z;

		return this;

	},

	getW: function ( index ) {

		return this.array[ index * this.itemSize + 3 ];

	},

	setW: function ( index, w ) {

		this.array[ index * this.itemSize + 3 ] = w;

		return this;

	},

	setXY: function ( index, x, y ) {

		index *= this.itemSize;

		this.array[ index + 0 ] = x;
		this.array[ index + 1 ] = y;

		return this;

	},

	setXYZ: function ( index, x, y, z ) {

		index *= this.itemSize;

		this.array[ index + 0 ] = x;
		this.array[ index + 1 ] = y;
		this.array[ index + 2 ] = z;

		return this;

	},

	setXYZW: function ( index, x, y, z, w ) {

		index *= this.itemSize;

		this.array[ index + 0 ] = x;
		this.array[ index + 1 ] = y;
		this.array[ index + 2 ] = z;
		this.array[ index + 3 ] = w;

		return this;

	},

	clone: function () {

		return new this.constructor().copy( this );

	}

};

//

THREE.Int8Attribute = function ( array, itemSize ) {

	return new THREE.BufferAttribute( new Int8Array( array ), itemSize );

};

THREE.Uint8Attribute = function ( array, itemSize ) {

	return new THREE.BufferAttribute( new Uint8Array( array ), itemSize );

};

THREE.Uint8ClampedAttribute = function ( array, itemSize ) {

	return new THREE.BufferAttribute( new Uint8ClampedArray( array ), itemSize );

};

THREE.Int16Attribute = function ( array, itemSize ) {

	return new THREE.BufferAttribute( new Int16Array( array ), itemSize );

};

THREE.Uint16Attribute = function ( array, itemSize ) {

	return new THREE.BufferAttribute( new Uint16Array( array ), itemSize );

};

THREE.Int32Attribute = function ( array, itemSize ) {

	return new THREE.BufferAttribute( new Int32Array( array ), itemSize );

};

THREE.Uint32Attribute = function ( array, itemSize ) {

	return new THREE.BufferAttribute( new Uint32Array( array ), itemSize );

};

THREE.Float32Attribute = function ( array, itemSize ) {

	return new THREE.BufferAttribute( new Float32Array( array ), itemSize );

};

THREE.Float64Attribute = function ( array, itemSize ) {

	return new THREE.BufferAttribute( new Float64Array( array ), itemSize );

};


// Deprecated

THREE.DynamicBufferAttribute = function ( array, itemSize ) {

	console.warn( 'THREE.DynamicBufferAttribute has been removed. Use new THREE.BufferAttribute().setDynamic( true ) instead.' );
	return new THREE.BufferAttribute( array, itemSize ).setDynamic( true );

};

// File:src/core/InstancedBufferAttribute.js

/**
 * @author benaadams / https://twitter.com/ben_a_adams
 */

THREE.InstancedBufferAttribute = function ( array, itemSize, meshPerAttribute ) {

	THREE.BufferAttribute.call( this, array, itemSize );

	this.meshPerAttribute = meshPerAttribute || 1;

};

THREE.InstancedBufferAttribute.prototype = Object.create( THREE.BufferAttribute.prototype );
THREE.InstancedBufferAttribute.prototype.constructor = THREE.InstancedBufferAttribute;

THREE.InstancedBufferAttribute.prototype.copy = function ( source ) {

	THREE.BufferAttribute.prototype.copy.call( this, source );

	this.meshPerAttribute = source.meshPerAttribute;

	return this;

};

// File:src/core/InterleavedBuffer.js

/**
 * @author benaadams / https://twitter.com/ben_a_adams
 */

THREE.InterleavedBuffer = function ( array, stride ) {

	this.uuid = THREE.Math.generateUUID();

	this.array = array;
	this.stride = stride;

	this.dynamic = false;
	this.updateRange = { offset: 0, count: - 1 };

	this.version = 0;

};

THREE.InterleavedBuffer.prototype = {

	constructor: THREE.InterleavedBuffer,

	get length () {

		return this.array.length;

	},

	get count () {

		return this.array.length / this.stride;

	},

	set needsUpdate( value ) {

		if ( value === true ) this.version ++;

	},

	setDynamic: function ( value ) {

		this.dynamic = value;

		return this;

	},

	copy: function ( source ) {

		this.array = new source.array.constructor( source.array );
		this.stride = source.stride;
		this.dynamic = source.dynamic;

		return this;

	},

	copyAt: function ( index1, attribute, index2 ) {

		index1 *= this.stride;
		index2 *= attribute.stride;

		for ( var i = 0, l = this.stride; i < l; i ++ ) {

			this.array[ index1 + i ] = attribute.array[ index2 + i ];

		}

		return this;

	},

	set: function ( value, offset ) {

		if ( offset === undefined ) offset = 0;

		this.array.set( value, offset );

		return this;

	},

	clone: function () {

		return new this.constructor().copy( this );

	}

};

// File:src/core/InstancedInterleavedBuffer.js

/**
 * @author benaadams / https://twitter.com/ben_a_adams
 */

THREE.InstancedInterleavedBuffer = function ( array, stride, meshPerAttribute ) {

	THREE.InterleavedBuffer.call( this, array, stride );

	this.meshPerAttribute = meshPerAttribute || 1;

};

THREE.InstancedInterleavedBuffer.prototype = Object.create( THREE.InterleavedBuffer.prototype );
THREE.InstancedInterleavedBuffer.prototype.constructor = THREE.InstancedInterleavedBuffer;

THREE.InstancedInterleavedBuffer.prototype.copy = function ( source ) {

	THREE.InterleavedBuffer.prototype.copy.call( this, source );

	this.meshPerAttribute = source.meshPerAttribute;

	return this;

};

// File:src/core/InterleavedBufferAttribute.js

/**
 * @author benaadams / https://twitter.com/ben_a_adams
 */

THREE.InterleavedBufferAttribute = function ( interleavedBuffer, itemSize, offset ) {

	this.uuid = THREE.Math.generateUUID();

	this.data = interleavedBuffer;
	this.itemSize = itemSize;
	this.offset = offset;

};


THREE.InterleavedBufferAttribute.prototype = {

	constructor: THREE.InterleavedBufferAttribute,

	get length() {

		console.warn( 'THREE.BufferAttribute: .length has been deprecated. Please use .count.' );
		return this.array.length;

	},

	get count() {

		return this.data.count;

	},

	setX: function ( index, x ) {

		this.data.array[ index * this.data.stride + this.offset ] = x;

		return this;

	},

	setY: function ( index, y ) {

		this.data.array[ index * this.data.stride + this.offset + 1 ] = y;

		return this;

	},

	setZ: function ( index, z ) {

		this.data.array[ index * this.data.stride + this.offset + 2 ] = z;

		return this;

	},

	setW: function ( index, w ) {

		this.data.array[ index * this.data.stride + this.offset + 3 ] = w;

		return this;

	},

	getX: function ( index ) {

		return this.data.array[ index * this.data.stride + this.offset ];

	},

	getY: function ( index ) {

		return this.data.array[ index * this.data.stride + this.offset + 1 ];

	},

	getZ: function ( index ) {

		return this.data.array[ index * this.data.stride + this.offset + 2 ];

	},

	getW: function ( index ) {

		return this.data.array[ index * this.data.stride + this.offset + 3 ];

	},

	setXY: function ( index, x, y ) {

		index = index * this.data.stride + this.offset;

		this.data.array[ index + 0 ] = x;
		this.data.array[ index + 1 ] = y;

		return this;

	},

	setXYZ: function ( index, x, y, z ) {

		index = index * this.data.stride + this.offset;

		this.data.array[ index + 0 ] = x;
		this.data.array[ index + 1 ] = y;
		this.data.array[ index + 2 ] = z;

		return this;

	},

	setXYZW: function ( index, x, y, z, w ) {

		index = index * this.data.stride + this.offset;

		this.data.array[ index + 0 ] = x;
		this.data.array[ index + 1 ] = y;
		this.data.array[ index + 2 ] = z;
		this.data.array[ index + 3 ] = w;

		return this;

	}

};

// File:src/core/Geometry.js

/**
 * @author mrdoob / http://mrdoob.com/
 * @author kile / http://kile.stravaganza.org/
 * @author alteredq / http://alteredqualia.com/
 * @author mikael emtinger / http://gomo.se/
 * @author zz85 / http://www.lab4games.net/zz85/blog
 * @author bhouston / http://clara.io
 */

THREE.Geometry = function () {

	Object.defineProperty( this, 'id', { value: THREE.GeometryIdCount ++ } );

	this.uuid = THREE.Math.generateUUID();

	this.name = '';
	this.type = 'Geometry';

	this.vertices = [];
	this.colors = [];
	this.faces = [];
	this.faceVertexUvs = [ [] ];

	this.morphTargets = [];
	this.morphNormals = [];

	this.skinWeights = [];
	this.skinIndices = [];

	this.lineDistances = [];

	this.boundingBox = null;
	this.boundingSphere = null;

	// update flags

	this.verticesNeedUpdate = false;
	this.elementsNeedUpdate = false;
	this.uvsNeedUpdate = false;
	this.normalsNeedUpdate = false;
	this.colorsNeedUpdate = false;
	this.lineDistancesNeedUpdate = false;
	this.groupsNeedUpdate = false;

};

Object.assign( THREE.Geometry.prototype, THREE.EventDispatcher.prototype, {

	applyMatrix: function ( matrix ) {

		var normalMatrix = new THREE.Matrix3().getNormalMatrix( matrix );

		for ( var i = 0, il = this.vertices.length; i < il; i ++ ) {

			var vertex = this.vertices[ i ];
			vertex.applyMatrix4( matrix );

		}

		for ( var i = 0, il = this.faces.length; i < il; i ++ ) {

			var face = this.faces[ i ];
			face.normal.applyMatrix3( normalMatrix ).normalize();

			for ( var j = 0, jl = face.vertexNormals.length; j < jl; j ++ ) {

				face.vertexNormals[ j ].applyMatrix3( normalMatrix ).normalize();

			}

		}

		if ( this.boundingBox !== null ) {

			this.computeBoundingBox();

		}

		if ( this.boundingSphere !== null ) {

			this.computeBoundingSphere();

		}

		this.verticesNeedUpdate = true;
		this.normalsNeedUpdate = true;

		return this;

	},

	rotateX: function () {

		// rotate geometry around world x-axis

		var m1;

		return function rotateX( angle ) {

			if ( m1 === undefined ) m1 = new THREE.Matrix4();

			m1.makeRotationX( angle );

			this.applyMatrix( m1 );

			return this;

		};

	}(),

	rotateY: function () {

		// rotate geometry around world y-axis

		var m1;

		return function rotateY( angle ) {

			if ( m1 === undefined ) m1 = new THREE.Matrix4();

			m1.makeRotationY( angle );

			this.applyMatrix( m1 );

			return this;

		};

	}(),

	rotateZ: function () {

		// rotate geometry around world z-axis

		var m1;

		return function rotateZ( angle ) {

			if ( m1 === undefined ) m1 = new THREE.Matrix4();

			m1.makeRotationZ( angle );

			this.applyMatrix( m1 );

			return this;

		};

	}(),

	translate: function () {

		// translate geometry

		var m1;

		return function translate( x, y, z ) {

			if ( m1 === undefined ) m1 = new THREE.Matrix4();

			m1.makeTranslation( x, y, z );

			this.applyMatrix( m1 );

			return this;

		};

	}(),

	scale: function () {

		// scale geometry

		var m1;

		return function scale( x, y, z ) {

			if ( m1 === undefined ) m1 = new THREE.Matrix4();

			m1.makeScale( x, y, z );

			this.applyMatrix( m1 );

			return this;

		};

	}(),

	lookAt: function () {

		var obj;

		return function lookAt( vector ) {

			if ( obj === undefined ) obj = new THREE.Object3D();

			obj.lookAt( vector );

			obj.updateMatrix();

			this.applyMatrix( obj.matrix );

		};

	}(),

	fromBufferGeometry: function ( geometry ) {

		var scope = this;

		var indices = geometry.index !== null ? geometry.index.array : undefined;
		var attributes = geometry.attributes;

		var positions = attributes.position.array;
		var normals = attributes.normal !== undefined ? attributes.normal.array : undefined;
		var colors = attributes.color !== undefined ? attributes.color.array : undefined;
		var uvs = attributes.uv !== undefined ? attributes.uv.array : undefined;
		var uvs2 = attributes.uv2 !== undefined ? attributes.uv2.array : undefined;

		if ( uvs2 !== undefined ) this.faceVertexUvs[ 1 ] = [];

		var tempNormals = [];
		var tempUVs = [];
		var tempUVs2 = [];

		for ( var i = 0, j = 0; i < positions.length; i += 3, j += 2 ) {

			scope.vertices.push( new THREE.Vector3( positions[ i ], positions[ i + 1 ], positions[ i + 2 ] ) );

			if ( normals !== undefined ) {

				tempNormals.push( new THREE.Vector3( normals[ i ], normals[ i + 1 ], normals[ i + 2 ] ) );

			}

			if ( colors !== undefined ) {

				scope.colors.push( new THREE.Color( colors[ i ], colors[ i + 1 ], colors[ i + 2 ] ) );

			}

			if ( uvs !== undefined ) {

				tempUVs.push( new THREE.Vector2( uvs[ j ], uvs[ j + 1 ] ) );

			}

			if ( uvs2 !== undefined ) {

				tempUVs2.push( new THREE.Vector2( uvs2[ j ], uvs2[ j + 1 ] ) );

			}

		}

		function addFace( a, b, c, materialIndex ) {

			var vertexNormals = normals !== undefined ? [ tempNormals[ a ].clone(), tempNormals[ b ].clone(), tempNormals[ c ].clone() ] : [];
			var vertexColors = colors !== undefined ? [ scope.colors[ a ].clone(), scope.colors[ b ].clone(), scope.colors[ c ].clone() ] : [];

			var face = new THREE.Face3( a, b, c, vertexNormals, vertexColors, materialIndex );

			scope.faces.push( face );

			if ( uvs !== undefined ) {

				scope.faceVertexUvs[ 0 ].push( [ tempUVs[ a ].clone(), tempUVs[ b ].clone(), tempUVs[ c ].clone() ] );

			}

			if ( uvs2 !== undefined ) {

				scope.faceVertexUvs[ 1 ].push( [ tempUVs2[ a ].clone(), tempUVs2[ b ].clone(), tempUVs2[ c ].clone() ] );

			}

		}

		if ( indices !== undefined ) {

			var groups = geometry.groups;

			if ( groups.length > 0 ) {

				for ( var i = 0; i < groups.length; i ++ ) {

					var group = groups[ i ];

					var start = group.start;
					var count = group.count;

					for ( var j = start, jl = start + count; j < jl; j += 3 ) {

						addFace( indices[ j ], indices[ j + 1 ], indices[ j + 2 ], group.materialIndex  );

					}

				}

			} else {

				for ( var i = 0; i < indices.length; i += 3 ) {

					addFace( indices[ i ], indices[ i + 1 ], indices[ i + 2 ] );

				}

			}

		} else {

			for ( var i = 0; i < positions.length / 3; i += 3 ) {

				addFace( i, i + 1, i + 2 );

			}

		}

		this.computeFaceNormals();

		if ( geometry.boundingBox !== null ) {

			this.boundingBox = geometry.boundingBox.clone();

		}

		if ( geometry.boundingSphere !== null ) {

			this.boundingSphere = geometry.boundingSphere.clone();

		}

		return this;

	},

	center: function () {

		this.computeBoundingBox();

		var offset = this.boundingBox.center().negate();

		this.translate( offset.x, offset.y, offset.z );

		return offset;

	},

	normalize: function () {

		this.computeBoundingSphere();

		var center = this.boundingSphere.center;
		var radius = this.boundingSphere.radius;

		var s = radius === 0 ? 1 : 1.0 / radius;

		var matrix = new THREE.Matrix4();
		matrix.set(
			s, 0, 0, - s * center.x,
			0, s, 0, - s * center.y,
			0, 0, s, - s * center.z,
			0, 0, 0, 1
		);

		this.applyMatrix( matrix );

		return this;

	},

	computeFaceNormals: function () {

		var cb = new THREE.Vector3(), ab = new THREE.Vector3();

		for ( var f = 0, fl = this.faces.length; f < fl; f ++ ) {

			var face = this.faces[ f ];

			var vA = this.vertices[ face.a ];
			var vB = this.vertices[ face.b ];
			var vC = this.vertices[ face.c ];

			cb.subVectors( vC, vB );
			ab.subVectors( vA, vB );
			cb.cross( ab );

			cb.normalize();

			face.normal.copy( cb );

		}

	},

	computeVertexNormals: function ( areaWeighted ) {

		if ( areaWeighted === undefined ) areaWeighted = true;

		var v, vl, f, fl, face, vertices;

		vertices = new Array( this.vertices.length );

		for ( v = 0, vl = this.vertices.length; v < vl; v ++ ) {

			vertices[ v ] = new THREE.Vector3();

		}

		if ( areaWeighted ) {

			// vertex normals weighted by triangle areas
			// http://www.iquilezles.org/www/articles/normals/normals.htm

			var vA, vB, vC;
			var cb = new THREE.Vector3(), ab = new THREE.Vector3();

			for ( f = 0, fl = this.faces.length; f < fl; f ++ ) {

				face = this.faces[ f ];

				vA = this.vertices[ face.a ];
				vB = this.vertices[ face.b ];
				vC = this.vertices[ face.c ];

				cb.subVectors( vC, vB );
				ab.subVectors( vA, vB );
				cb.cross( ab );

				vertices[ face.a ].add( cb );
				vertices[ face.b ].add( cb );
				vertices[ face.c ].add( cb );

			}

		} else {

			for ( f = 0, fl = this.faces.length; f < fl; f ++ ) {

				face = this.faces[ f ];

				vertices[ face.a ].add( face.normal );
				vertices[ face.b ].add( face.normal );
				vertices[ face.c ].add( face.normal );

			}

		}

		for ( v = 0, vl = this.vertices.length; v < vl; v ++ ) {

			vertices[ v ].normalize();

		}

		for ( f = 0, fl = this.faces.length; f < fl; f ++ ) {

			face = this.faces[ f ];

			var vertexNormals = face.vertexNormals;

			if ( vertexNormals.length === 3 ) {

				vertexNormals[ 0 ].copy( vertices[ face.a ] );
				vertexNormals[ 1 ].copy( vertices[ face.b ] );
				vertexNormals[ 2 ].copy( vertices[ face.c ] );

			} else {

				vertexNormals[ 0 ] = vertices[ face.a ].clone();
				vertexNormals[ 1 ] = vertices[ face.b ].clone();
				vertexNormals[ 2 ] = vertices[ face.c ].clone();

			}

		}

		if ( this.faces.length > 0 ) {

			this.normalsNeedUpdate = true;

		}

	},

	computeMorphNormals: function () {

		var i, il, f, fl, face;

		// save original normals
		// - create temp variables on first access
		//   otherwise just copy (for faster repeated calls)

		for ( f = 0, fl = this.faces.length; f < fl; f ++ ) {

			face = this.faces[ f ];

			if ( ! face.__originalFaceNormal ) {

				face.__originalFaceNormal = face.normal.clone();

			} else {

				face.__originalFaceNormal.copy( face.normal );

			}

			if ( ! face.__originalVertexNormals ) face.__originalVertexNormals = [];

			for ( i = 0, il = face.vertexNormals.length; i < il; i ++ ) {

				if ( ! face.__originalVertexNormals[ i ] ) {

					face.__originalVertexNormals[ i ] = face.vertexNormals[ i ].clone();

				} else {

					face.__originalVertexNormals[ i ].copy( face.vertexNormals[ i ] );

				}

			}

		}

		// use temp geometry to compute face and vertex normals for each morph

		var tmpGeo = new THREE.Geometry();
		tmpGeo.faces = this.faces;

		for ( i = 0, il = this.morphTargets.length; i < il; i ++ ) {

			// create on first access

			if ( ! this.morphNormals[ i ] ) {

				this.morphNormals[ i ] = {};
				this.morphNormals[ i ].faceNormals = [];
				this.morphNormals[ i ].vertexNormals = [];

				var dstNormalsFace = this.morphNormals[ i ].faceNormals;
				var dstNormalsVertex = this.morphNormals[ i ].vertexNormals;

				var faceNormal, vertexNormals;

				for ( f = 0, fl = this.faces.length; f < fl; f ++ ) {

					faceNormal = new THREE.Vector3();
					vertexNormals = { a: new THREE.Vector3(), b: new THREE.Vector3(), c: new THREE.Vector3() };

					dstNormalsFace.push( faceNormal );
					dstNormalsVertex.push( vertexNormals );

				}

			}

			var morphNormals = this.morphNormals[ i ];

			// set vertices to morph target

			tmpGeo.vertices = this.morphTargets[ i ].vertices;

			// compute morph normals

			tmpGeo.computeFaceNormals();
			tmpGeo.computeVertexNormals();

			// store morph normals

			var faceNormal, vertexNormals;

			for ( f = 0, fl = this.faces.length; f < fl; f ++ ) {

				face = this.faces[ f ];

				faceNormal = morphNormals.faceNormals[ f ];
				vertexNormals = morphNormals.vertexNormals[ f ];

				faceNormal.copy( face.normal );

				vertexNormals.a.copy( face.vertexNormals[ 0 ] );
				vertexNormals.b.copy( face.vertexNormals[ 1 ] );
				vertexNormals.c.copy( face.vertexNormals[ 2 ] );

			}

		}

		// restore original normals

		for ( f = 0, fl = this.faces.length; f < fl; f ++ ) {

			face = this.faces[ f ];

			face.normal = face.__originalFaceNormal;
			face.vertexNormals = face.__originalVertexNormals;

		}

	},

	computeTangents: function () {

		console.warn( 'THREE.Geometry: .computeTangents() has been removed.' );

	},

	computeLineDistances: function () {

		var d = 0;
		var vertices = this.vertices;

		for ( var i = 0, il = vertices.length; i < il; i ++ ) {

			if ( i > 0 ) {

				d += vertices[ i ].distanceTo( vertices[ i - 1 ] );

			}

			this.lineDistances[ i ] = d;

		}

	},

	computeBoundingBox: function () {

		if ( this.boundingBox === null ) {

			this.boundingBox = new THREE.Box3();

		}

		this.boundingBox.setFromPoints( this.vertices );

	},

	computeBoundingSphere: function () {

		if ( this.boundingSphere === null ) {

			this.boundingSphere = new THREE.Sphere();

		}

		this.boundingSphere.setFromPoints( this.vertices );

	},

	merge: function ( geometry, matrix, materialIndexOffset ) {

		if ( geometry instanceof THREE.Geometry === false ) {

			console.error( 'THREE.Geometry.merge(): geometry not an instance of THREE.Geometry.', geometry );
			return;

		}

		var normalMatrix,
		vertexOffset = this.vertices.length,
		vertices1 = this.vertices,
		vertices2 = geometry.vertices,
		faces1 = this.faces,
		faces2 = geometry.faces,
		uvs1 = this.faceVertexUvs[ 0 ],
		uvs2 = geometry.faceVertexUvs[ 0 ];

		if ( materialIndexOffset === undefined ) materialIndexOffset = 0;

		if ( matrix !== undefined ) {

			normalMatrix = new THREE.Matrix3().getNormalMatrix( matrix );

		}

		// vertices

		for ( var i = 0, il = vertices2.length; i < il; i ++ ) {

			var vertex = vertices2[ i ];

			var vertexCopy = vertex.clone();

			if ( matrix !== undefined ) vertexCopy.applyMatrix4( matrix );

			vertices1.push( vertexCopy );

		}

		// faces

		for ( i = 0, il = faces2.length; i < il; i ++ ) {

			var face = faces2[ i ], faceCopy, normal, color,
			faceVertexNormals = face.vertexNormals,
			faceVertexColors = face.vertexColors;

			faceCopy = new THREE.Face3( face.a + vertexOffset, face.b + vertexOffset, face.c + vertexOffset );
			faceCopy.normal.copy( face.normal );

			if ( normalMatrix !== undefined ) {

				faceCopy.normal.applyMatrix3( normalMatrix ).normalize();

			}

			for ( var j = 0, jl = faceVertexNormals.length; j < jl; j ++ ) {

				normal = faceVertexNormals[ j ].clone();

				if ( normalMatrix !== undefined ) {

					normal.applyMatrix3( normalMatrix ).normalize();

				}

				faceCopy.vertexNormals.push( normal );

			}

			faceCopy.color.copy( face.color );

			for ( var j = 0, jl = faceVertexColors.length; j < jl; j ++ ) {

				color = faceVertexColors[ j ];
				faceCopy.vertexColors.push( color.clone() );

			}

			faceCopy.materialIndex = face.materialIndex + materialIndexOffset;

			faces1.push( faceCopy );

		}

		// uvs

		for ( i = 0, il = uvs2.length; i < il; i ++ ) {

			var uv = uvs2[ i ], uvCopy = [];

			if ( uv === undefined ) {

				continue;

			}

			for ( var j = 0, jl = uv.length; j < jl; j ++ ) {

				uvCopy.push( uv[ j ].clone() );

			}

			uvs1.push( uvCopy );

		}

	},

	mergeMesh: function ( mesh ) {

		if ( mesh instanceof THREE.Mesh === false ) {

			console.error( 'THREE.Geometry.mergeMesh(): mesh not an instance of THREE.Mesh.', mesh );
			return;

		}

		mesh.matrixAutoUpdate && mesh.updateMatrix();

		this.merge( mesh.geometry, mesh.matrix );

	},

	/*
	 * Checks for duplicate vertices with hashmap.
	 * Duplicated vertices are removed
	 * and faces' vertices are updated.
	 */

	mergeVertices: function () {

		var verticesMap = {}; // Hashmap for looking up vertices by position coordinates (and making sure they are unique)
		var unique = [], changes = [];

		var v, key;
		var precisionPoints = 4; // number of decimal points, e.g. 4 for epsilon of 0.0001
		var precision = Math.pow( 10, precisionPoints );
		var i, il, face;
		var indices, j, jl;

		for ( i = 0, il = this.vertices.length; i < il; i ++ ) {

			v = this.vertices[ i ];
			key = Math.round( v.x * precision ) + '_' + Math.round( v.y * precision ) + '_' + Math.round( v.z * precision );

			if ( verticesMap[ key ] === undefined ) {

				verticesMap[ key ] = i;
				unique.push( this.vertices[ i ] );
				changes[ i ] = unique.length - 1;

			} else {

				//console.log('Duplicate vertex found. ', i, ' could be using ', verticesMap[key]);
				changes[ i ] = changes[ verticesMap[ key ] ];

			}

		}


		// if faces are completely degenerate after merging vertices, we
		// have to remove them from the geometry.
		var faceIndicesToRemove = [];

		for ( i = 0, il = this.faces.length; i < il; i ++ ) {

			face = this.faces[ i ];

			face.a = changes[ face.a ];
			face.b = changes[ face.b ];
			face.c = changes[ face.c ];

			indices = [ face.a, face.b, face.c ];

			var dupIndex = - 1;

			// if any duplicate vertices are found in a Face3
			// we have to remove the face as nothing can be saved
			for ( var n = 0; n < 3; n ++ ) {

				if ( indices[ n ] === indices[ ( n + 1 ) % 3 ] ) {

					dupIndex = n;
					faceIndicesToRemove.push( i );
					break;

				}

			}

		}

		for ( i = faceIndicesToRemove.length - 1; i >= 0; i -- ) {

			var idx = faceIndicesToRemove[ i ];

			this.faces.splice( idx, 1 );

			for ( j = 0, jl = this.faceVertexUvs.length; j < jl; j ++ ) {

				this.faceVertexUvs[ j ].splice( idx, 1 );

			}

		}

		// Use unique set of vertices

		var diff = this.vertices.length - unique.length;
		this.vertices = unique;
		return diff;

	},

	sortFacesByMaterialIndex: function () {

		var faces = this.faces;
		var length = faces.length;

		// tag faces

		for ( var i = 0; i < length; i ++ ) {

			faces[ i ]._id = i;

		}

		// sort faces

		function materialIndexSort( a, b ) {

			return a.materialIndex - b.materialIndex;

		}

		faces.sort( materialIndexSort );

		// sort uvs

		var uvs1 = this.faceVertexUvs[ 0 ];
		var uvs2 = this.faceVertexUvs[ 1 ];

		var newUvs1, newUvs2;

		if ( uvs1 && uvs1.length === length ) newUvs1 = [];
		if ( uvs2 && uvs2.length === length ) newUvs2 = [];

		for ( var i = 0; i < length; i ++ ) {

			var id = faces[ i ]._id;

			if ( newUvs1 ) newUvs1.push( uvs1[ id ] );
			if ( newUvs2 ) newUvs2.push( uvs2[ id ] );

		}

		if ( newUvs1 ) this.faceVertexUvs[ 0 ] = newUvs1;
		if ( newUvs2 ) this.faceVertexUvs[ 1 ] = newUvs2;

	},

	toJSON: function () {

		var data = {
			metadata: {
				version: 4.4,
				type: 'Geometry',
				generator: 'Geometry.toJSON'
			}
		};

		// standard Geometry serialization

		data.uuid = this.uuid;
		data.type = this.type;
		if ( this.name !== '' ) data.name = this.name;

		if ( this.parameters !== undefined ) {

			var parameters = this.parameters;

			for ( var key in parameters ) {

				if ( parameters[ key ] !== undefined ) data[ key ] = parameters[ key ];

			}

			return data;

		}

		var vertices = [];

		for ( var i = 0; i < this.vertices.length; i ++ ) {

			var vertex = this.vertices[ i ];
			vertices.push( vertex.x, vertex.y, vertex.z );

		}

		var faces = [];
		var normals = [];
		var normalsHash = {};
		var colors = [];
		var colorsHash = {};
		var uvs = [];
		var uvsHash = {};

		for ( var i = 0; i < this.faces.length; i ++ ) {

			var face = this.faces[ i ];

			var hasMaterial = true;
			var hasFaceUv = false; // deprecated
			var hasFaceVertexUv = this.faceVertexUvs[ 0 ][ i ] !== undefined;
			var hasFaceNormal = face.normal.length() > 0;
			var hasFaceVertexNormal = face.vertexNormals.length > 0;
			var hasFaceColor = face.color.r !== 1 || face.color.g !== 1 || face.color.b !== 1;
			var hasFaceVertexColor = face.vertexColors.length > 0;

			var faceType = 0;

			faceType = setBit( faceType, 0, 0 ); // isQuad
			faceType = setBit( faceType, 1, hasMaterial );
			faceType = setBit( faceType, 2, hasFaceUv );
			faceType = setBit( faceType, 3, hasFaceVertexUv );
			faceType = setBit( faceType, 4, hasFaceNormal );
			faceType = setBit( faceType, 5, hasFaceVertexNormal );
			faceType = setBit( faceType, 6, hasFaceColor );
			faceType = setBit( faceType, 7, hasFaceVertexColor );

			faces.push( faceType );
			faces.push( face.a, face.b, face.c );
			faces.push( face.materialIndex );

			if ( hasFaceVertexUv ) {

				var faceVertexUvs = this.faceVertexUvs[ 0 ][ i ];

				faces.push(
					getUvIndex( faceVertexUvs[ 0 ] ),
					getUvIndex( faceVertexUvs[ 1 ] ),
					getUvIndex( faceVertexUvs[ 2 ] )
				);

			}

			if ( hasFaceNormal ) {

				faces.push( getNormalIndex( face.normal ) );

			}

			if ( hasFaceVertexNormal ) {

				var vertexNormals = face.vertexNormals;

				faces.push(
					getNormalIndex( vertexNormals[ 0 ] ),
					getNormalIndex( vertexNormals[ 1 ] ),
					getNormalIndex( vertexNormals[ 2 ] )
				);

			}

			if ( hasFaceColor ) {

				faces.push( getColorIndex( face.color ) );

			}

			if ( hasFaceVertexColor ) {

				var vertexColors = face.vertexColors;

				faces.push(
					getColorIndex( vertexColors[ 0 ] ),
					getColorIndex( vertexColors[ 1 ] ),
					getColorIndex( vertexColors[ 2 ] )
				);

			}

		}

		function setBit( value, position, enabled ) {

			return enabled ? value | ( 1 << position ) : value & ( ~ ( 1 << position ) );

		}

		function getNormalIndex( normal ) {

			var hash = normal.x.toString() + normal.y.toString() + normal.z.toString();

			if ( normalsHash[ hash ] !== undefined ) {

				return normalsHash[ hash ];

			}

			normalsHash[ hash ] = normals.length / 3;
			normals.push( normal.x, normal.y, normal.z );

			return normalsHash[ hash ];

		}

		function getColorIndex( color ) {

			var hash = color.r.toString() + color.g.toString() + color.b.toString();

			if ( colorsHash[ hash ] !== undefined ) {

				return colorsHash[ hash ];

			}

			colorsHash[ hash ] = colors.length;
			colors.push( color.getHex() );

			return colorsHash[ hash ];

		}

		function getUvIndex( uv ) {

			var hash = uv.x.toString() + uv.y.toString();

			if ( uvsHash[ hash ] !== undefined ) {

				return uvsHash[ hash ];

			}

			uvsHash[ hash ] = uvs.length / 2;
			uvs.push( uv.x, uv.y );

			return uvsHash[ hash ];

		}

		data.data = {};

		data.data.vertices = vertices;
		data.data.normals = normals;
		if ( colors.length > 0 ) data.data.colors = colors;
		if ( uvs.length > 0 ) data.data.uvs = [ uvs ]; // temporal backward compatibility
		data.data.faces = faces;

		return data;

	},

	clone: function () {

		/*
		// Handle primitives

		var parameters = this.parameters;

		if ( parameters !== undefined ) {

			var values = [];

			for ( var key in parameters ) {

				values.push( parameters[ key ] );

			}

			var geometry = Object.create( this.constructor.prototype );
			this.constructor.apply( geometry, values );
			return geometry;

		}

		return new this.constructor().copy( this );
		*/

		return new THREE.Geometry().copy( this );

	},

	copy: function ( source ) {

		this.vertices = [];
		this.faces = [];
		this.faceVertexUvs = [ [] ];

		var vertices = source.vertices;

		for ( var i = 0, il = vertices.length; i < il; i ++ ) {

			this.vertices.push( vertices[ i ].clone() );

		}

		var faces = source.faces;

		for ( var i = 0, il = faces.length; i < il; i ++ ) {

			this.faces.push( faces[ i ].clone() );

		}

		for ( var i = 0, il = source.faceVertexUvs.length; i < il; i ++ ) {

			var faceVertexUvs = source.faceVertexUvs[ i ];

			if ( this.faceVertexUvs[ i ] === undefined ) {

				this.faceVertexUvs[ i ] = [];

			}

			for ( var j = 0, jl = faceVertexUvs.length; j < jl; j ++ ) {

				var uvs = faceVertexUvs[ j ], uvsCopy = [];

				for ( var k = 0, kl = uvs.length; k < kl; k ++ ) {

					var uv = uvs[ k ];

					uvsCopy.push( uv.clone() );

				}

				this.faceVertexUvs[ i ].push( uvsCopy );

			}

		}

		return this;

	},

	dispose: function () {

		this.dispatchEvent( { type: 'dispose' } );

	}

} );

THREE.GeometryIdCount = 0;

// File:src/core/DirectGeometry.js

/**
 * @author mrdoob / http://mrdoob.com/
 */

THREE.DirectGeometry = function () {

	Object.defineProperty( this, 'id', { value: THREE.GeometryIdCount ++ } );

	this.uuid = THREE.Math.generateUUID();

	this.name = '';
	this.type = 'DirectGeometry';

	this.indices = [];
	this.vertices = [];
	this.normals = [];
	this.colors = [];
	this.uvs = [];
	this.uvs2 = [];

	this.groups = [];

	this.morphTargets = {};

	this.skinWeights = [];
	this.skinIndices = [];

	// this.lineDistances = [];

	this.boundingBox = null;
	this.boundingSphere = null;

	// update flags

	this.verticesNeedUpdate = false;
	this.normalsNeedUpdate = false;
	this.colorsNeedUpdate = false;
	this.uvsNeedUpdate = false;
	this.groupsNeedUpdate = false;

};

Object.assign( THREE.DirectGeometry.prototype, THREE.EventDispatcher.prototype, {

	computeBoundingBox: THREE.Geometry.prototype.computeBoundingBox,
	computeBoundingSphere: THREE.Geometry.prototype.computeBoundingSphere,

	computeFaceNormals: function () {

		console.warn( 'THREE.DirectGeometry: computeFaceNormals() is not a method of this type of geometry.' );

	},

	computeVertexNormals: function () {

		console.warn( 'THREE.DirectGeometry: computeVertexNormals() is not a method of this type of geometry.' );

	},

	computeGroups: function ( geometry ) {

		var group;
		var groups = [];
		var materialIndex;

		var faces = geometry.faces;

		for ( var i = 0; i < faces.length; i ++ ) {

			var face = faces[ i ];

			// materials

			if ( face.materialIndex !== materialIndex ) {

				materialIndex = face.materialIndex;

				if ( group !== undefined ) {

					group.count = ( i * 3 ) - group.start;
					groups.push( group );

				}

				group = {
					start: i * 3,
					materialIndex: materialIndex
				};

			}

		}

		if ( group !== undefined ) {

			group.count = ( i * 3 ) - group.start;
			groups.push( group );

		}

		this.groups = groups;

	},

	fromGeometry: function ( geometry ) {

		var faces = geometry.faces;
		var vertices = geometry.vertices;
		var faceVertexUvs = geometry.faceVertexUvs;

		var hasFaceVertexUv = faceVertexUvs[ 0 ] && faceVertexUvs[ 0 ].length > 0;
		var hasFaceVertexUv2 = faceVertexUvs[ 1 ] && faceVertexUvs[ 1 ].length > 0;

		// morphs

		var morphTargets = geometry.morphTargets;
		var morphTargetsLength = morphTargets.length;

		var morphTargetsPosition;

		if ( morphTargetsLength > 0 ) {

			morphTargetsPosition = [];

			for ( var i = 0; i < morphTargetsLength; i ++ ) {

				morphTargetsPosition[ i ] = [];

			}

			this.morphTargets.position = morphTargetsPosition;

		}

		var morphNormals = geometry.morphNormals;
		var morphNormalsLength = morphNormals.length;

		var morphTargetsNormal;

		if ( morphNormalsLength > 0 ) {

			morphTargetsNormal = [];

			for ( var i = 0; i < morphNormalsLength; i ++ ) {

				morphTargetsNormal[ i ] = [];

			}

			this.morphTargets.normal = morphTargetsNormal;

		}

		// skins

		var skinIndices = geometry.skinIndices;
		var skinWeights = geometry.skinWeights;

		var hasSkinIndices = skinIndices.length === vertices.length;
		var hasSkinWeights = skinWeights.length === vertices.length;

		//

		for ( var i = 0; i < faces.length; i ++ ) {

			var face = faces[ i ];

			this.vertices.push( vertices[ face.a ], vertices[ face.b ], vertices[ face.c ] );

			var vertexNormals = face.vertexNormals;

			if ( vertexNormals.length === 3 ) {

				this.normals.push( vertexNormals[ 0 ], vertexNormals[ 1 ], vertexNormals[ 2 ] );

			} else {

				var normal = face.normal;

				this.normals.push( normal, normal, normal );

			}

			var vertexColors = face.vertexColors;

			if ( vertexColors.length === 3 ) {

				this.colors.push( vertexColors[ 0 ], vertexColors[ 1 ], vertexColors[ 2 ] );

			} else {

				var color = face.color;

				this.colors.push( color, color, color );

			}

			if ( hasFaceVertexUv === true ) {

				var vertexUvs = faceVertexUvs[ 0 ][ i ];

				if ( vertexUvs !== undefined ) {

					this.uvs.push( vertexUvs[ 0 ], vertexUvs[ 1 ], vertexUvs[ 2 ] );

				} else {

					console.warn( 'THREE.DirectGeometry.fromGeometry(): Undefined vertexUv ', i );

					this.uvs.push( new THREE.Vector2(), new THREE.Vector2(), new THREE.Vector2() );

				}

			}

			if ( hasFaceVertexUv2 === true ) {

				var vertexUvs = faceVertexUvs[ 1 ][ i ];

				if ( vertexUvs !== undefined ) {

					this.uvs2.push( vertexUvs[ 0 ], vertexUvs[ 1 ], vertexUvs[ 2 ] );

				} else {

					console.warn( 'THREE.DirectGeometry.fromGeometry(): Undefined vertexUv2 ', i );

					this.uvs2.push( new THREE.Vector2(), new THREE.Vector2(), new THREE.Vector2() );

				}

			}

			// morphs

			for ( var j = 0; j < morphTargetsLength; j ++ ) {

				var morphTarget = morphTargets[ j ].vertices;

				morphTargetsPosition[ j ].push( morphTarget[ face.a ], morphTarget[ face.b ], morphTarget[ face.c ] );

			}

			for ( var j = 0; j < morphNormalsLength; j ++ ) {

				var morphNormal = morphNormals[ j ].vertexNormals[ i ];

				morphTargetsNormal[ j ].push( morphNormal.a, morphNormal.b, morphNormal.c );

			}

			// skins

			if ( hasSkinIndices ) {

				this.skinIndices.push( skinIndices[ face.a ], skinIndices[ face.b ], skinIndices[ face.c ] );

			}

			if ( hasSkinWeights ) {

				this.skinWeights.push( skinWeights[ face.a ], skinWeights[ face.b ], skinWeights[ face.c ] );

			}

		}

		this.computeGroups( geometry );

		this.verticesNeedUpdate = geometry.verticesNeedUpdate;
		this.normalsNeedUpdate = geometry.normalsNeedUpdate;
		this.colorsNeedUpdate = geometry.colorsNeedUpdate;
		this.uvsNeedUpdate = geometry.uvsNeedUpdate;
		this.groupsNeedUpdate = geometry.groupsNeedUpdate;

		return this;

	},

	dispose: function () {

		this.dispatchEvent( { type: 'dispose' } );

	}

} );

// File:src/core/BufferGeometry.js

/**
 * @author alteredq / http://alteredqualia.com/
 * @author mrdoob / http://mrdoob.com/
 */

THREE.BufferGeometry = function () {

	Object.defineProperty( this, 'id', { value: THREE.GeometryIdCount ++ } );

	this.uuid = THREE.Math.generateUUID();

	this.name = '';
	this.type = 'BufferGeometry';

	this.index = null;
	this.attributes = {};

	this.morphAttributes = {};

	this.groups = [];

	this.boundingBox = null;
	this.boundingSphere = null;

	this.drawRange = { start: 0, count: Infinity };

};

Object.assign( THREE.BufferGeometry.prototype, THREE.EventDispatcher.prototype, {

	getIndex: function () {

		return this.index;

	},

	setIndex: function ( index ) {

		this.index = index;

	},

	addAttribute: function ( name, attribute ) {

		if ( attribute instanceof THREE.BufferAttribute === false && attribute instanceof THREE.InterleavedBufferAttribute === false ) {

			console.warn( 'THREE.BufferGeometry: .addAttribute() now expects ( name, attribute ).' );

			this.addAttribute( name, new THREE.BufferAttribute( arguments[ 1 ], arguments[ 2 ] ) );

			return;

		}

		if ( name === 'index' ) {

			console.warn( 'THREE.BufferGeometry.addAttribute: Use .setIndex() for index attribute.' );
			this.setIndex( attribute );

			return;

		}

		this.attributes[ name ] = attribute;

		return this;

	},

	getAttribute: function ( name ) {

		return this.attributes[ name ];

	},

	removeAttribute: function ( name ) {

		delete this.attributes[ name ];

		return this;

	},

	addGroup: function ( start, count, materialIndex ) {

		this.groups.push( {

			start: start,
			count: count,
			materialIndex: materialIndex !== undefined ? materialIndex : 0

		} );

	},

	clearGroups: function () {

		this.groups = [];

	},

	setDrawRange: function ( start, count ) {

		this.drawRange.start = start;
		this.drawRange.count = count;

	},

	applyMatrix: function ( matrix ) {

		var position = this.attributes.position;

		if ( position !== undefined ) {

			matrix.applyToVector3Array( position.array );
			position.needsUpdate = true;

		}

		var normal = this.attributes.normal;

		if ( normal !== undefined ) {

			var normalMatrix = new THREE.Matrix3().getNormalMatrix( matrix );

			normalMatrix.applyToVector3Array( normal.array );
			normal.needsUpdate = true;

		}

		if ( this.boundingBox !== null ) {

			this.computeBoundingBox();

		}

		if ( this.boundingSphere !== null ) {

			this.computeBoundingSphere();

		}

		return this;

	},

	rotateX: function () {

		// rotate geometry around world x-axis

		var m1;

		return function rotateX( angle ) {

			if ( m1 === undefined ) m1 = new THREE.Matrix4();

			m1.makeRotationX( angle );

			this.applyMatrix( m1 );

			return this;

		};

	}(),

	rotateY: function () {

		// rotate geometry around world y-axis

		var m1;

		return function rotateY( angle ) {

			if ( m1 === undefined ) m1 = new THREE.Matrix4();

			m1.makeRotationY( angle );

			this.applyMatrix( m1 );

			return this;

		};

	}(),

	rotateZ: function () {

		// rotate geometry around world z-axis

		var m1;

		return function rotateZ( angle ) {

			if ( m1 === undefined ) m1 = new THREE.Matrix4();

			m1.makeRotationZ( angle );

			this.applyMatrix( m1 );

			return this;

		};

	}(),

	translate: function () {

		// translate geometry

		var m1;

		return function translate( x, y, z ) {

			if ( m1 === undefined ) m1 = new THREE.Matrix4();

			m1.makeTranslation( x, y, z );

			this.applyMatrix( m1 );

			return this;

		};

	}(),

	scale: function () {

		// scale geometry

		var m1;

		return function scale( x, y, z ) {

			if ( m1 === undefined ) m1 = new THREE.Matrix4();

			m1.makeScale( x, y, z );

			this.applyMatrix( m1 );

			return this;

		};

	}(),

	lookAt: function () {

		var obj;

		return function lookAt( vector ) {

			if ( obj === undefined ) obj = new THREE.Object3D();

			obj.lookAt( vector );

			obj.updateMatrix();

			this.applyMatrix( obj.matrix );

		};

	}(),

	center: function () {

		this.computeBoundingBox();

		var offset = this.boundingBox.center().negate();

		this.translate( offset.x, offset.y, offset.z );

		return offset;

	},

	setFromObject: function ( object ) {

		// console.log( 'THREE.BufferGeometry.setFromObject(). Converting', object, this );

		var geometry = object.geometry;

		if ( object instanceof THREE.Points || object instanceof THREE.Line ) {

			var positions = new THREE.Float32Attribute( geometry.vertices.length * 3, 3 );
			var colors = new THREE.Float32Attribute( geometry.colors.length * 3, 3 );

			this.addAttribute( 'position', positions.copyVector3sArray( geometry.vertices ) );
			this.addAttribute( 'color', colors.copyColorsArray( geometry.colors ) );

			if ( geometry.lineDistances && geometry.lineDistances.length === geometry.vertices.length ) {

				var lineDistances = new THREE.Float32Attribute( geometry.lineDistances.length, 1 );

				this.addAttribute( 'lineDistance', lineDistances.copyArray( geometry.lineDistances ) );

			}

			if ( geometry.boundingSphere !== null ) {

				this.boundingSphere = geometry.boundingSphere.clone();

			}

			if ( geometry.boundingBox !== null ) {

				this.boundingBox = geometry.boundingBox.clone();

			}

		} else if ( object instanceof THREE.Mesh ) {

			if ( geometry instanceof THREE.Geometry ) {

				this.fromGeometry( geometry );

			}

		}

		return this;

	},

	updateFromObject: function ( object ) {

		var geometry = object.geometry;

		if ( object instanceof THREE.Mesh ) {

			var direct = geometry.__directGeometry;

			if ( direct === undefined ) {

				return this.fromGeometry( geometry );

			}

			direct.verticesNeedUpdate = geometry.verticesNeedUpdate;
			direct.normalsNeedUpdate = geometry.normalsNeedUpdate;
			direct.colorsNeedUpdate = geometry.colorsNeedUpdate;
			direct.uvsNeedUpdate = geometry.uvsNeedUpdate;
			direct.groupsNeedUpdate = geometry.groupsNeedUpdate;

			geometry.verticesNeedUpdate = false;
			geometry.normalsNeedUpdate = false;
			geometry.colorsNeedUpdate = false;
			geometry.uvsNeedUpdate = false;
			geometry.groupsNeedUpdate = false;

			geometry = direct;

		}

		if ( geometry.verticesNeedUpdate === true ) {

			var attribute = this.attributes.position;

			if ( attribute !== undefined ) {

				attribute.copyVector3sArray( geometry.vertices );
				attribute.needsUpdate = true;

			}

			geometry.verticesNeedUpdate = false;

		}

		if ( geometry.normalsNeedUpdate === true ) {

			var attribute = this.attributes.normal;

			if ( attribute !== undefined ) {

				attribute.copyVector3sArray( geometry.normals );
				attribute.needsUpdate = true;

			}

			geometry.normalsNeedUpdate = false;

		}

		if ( geometry.colorsNeedUpdate === true ) {

			var attribute = this.attributes.color;

			if ( attribute !== undefined ) {

				attribute.copyColorsArray( geometry.colors );
				attribute.needsUpdate = true;

			}

			geometry.colorsNeedUpdate = false;

		}

		if ( geometry.uvsNeedUpdate ) {

			var attribute = this.attributes.uv;

			if ( attribute !== undefined ) {

				attribute.copyVector2sArray( geometry.uvs );
				attribute.needsUpdate = true;

			}

			geometry.uvsNeedUpdate = false;

		}

		if ( geometry.lineDistancesNeedUpdate ) {

			var attribute = this.attributes.lineDistance;

			if ( attribute !== undefined ) {

				attribute.copyArray( geometry.lineDistances );
				attribute.needsUpdate = true;

			}

			geometry.lineDistancesNeedUpdate = false;

		}

		if ( geometry.groupsNeedUpdate ) {

			geometry.computeGroups( object.geometry );
			this.groups = geometry.groups;

			geometry.groupsNeedUpdate = false;

		}

		return this;

	},

	fromGeometry: function ( geometry ) {

		geometry.__directGeometry = new THREE.DirectGeometry().fromGeometry( geometry );

		return this.fromDirectGeometry( geometry.__directGeometry );

	},

	fromDirectGeometry: function ( geometry ) {

		var positions = new Float32Array( geometry.vertices.length * 3 );
		this.addAttribute( 'position', new THREE.BufferAttribute( positions, 3 ).copyVector3sArray( geometry.vertices ) );

		if ( geometry.normals.length > 0 ) {

			var normals = new Float32Array( geometry.normals.length * 3 );
			this.addAttribute( 'normal', new THREE.BufferAttribute( normals, 3 ).copyVector3sArray( geometry.normals ) );

		}

		if ( geometry.colors.length > 0 ) {

			var colors = new Float32Array( geometry.colors.length * 3 );
			this.addAttribute( 'color', new THREE.BufferAttribute( colors, 3 ).copyColorsArray( geometry.colors ) );

		}

		if ( geometry.uvs.length > 0 ) {

			var uvs = new Float32Array( geometry.uvs.length * 2 );
			this.addAttribute( 'uv', new THREE.BufferAttribute( uvs, 2 ).copyVector2sArray( geometry.uvs ) );

		}

		if ( geometry.uvs2.length > 0 ) {

			var uvs2 = new Float32Array( geometry.uvs2.length * 2 );
			this.addAttribute( 'uv2', new THREE.BufferAttribute( uvs2, 2 ).copyVector2sArray( geometry.uvs2 ) );

		}

		if ( geometry.indices.length > 0 ) {

			var TypeArray = geometry.vertices.length > 65535 ? Uint32Array : Uint16Array;
			var indices = new TypeArray( geometry.indices.length * 3 );
			this.setIndex( new THREE.BufferAttribute( indices, 1 ).copyIndicesArray( geometry.indices ) );

		}

		// groups

		this.groups = geometry.groups;

		// morphs

		for ( var name in geometry.morphTargets ) {

			var array = [];
			var morphTargets = geometry.morphTargets[ name ];

			for ( var i = 0, l = morphTargets.length; i < l; i ++ ) {

				var morphTarget = morphTargets[ i ];

				var attribute = new THREE.Float32Attribute( morphTarget.length * 3, 3 );

				array.push( attribute.copyVector3sArray( morphTarget ) );

			}

			this.morphAttributes[ name ] = array;

		}

		// skinning

		if ( geometry.skinIndices.length > 0 ) {

			var skinIndices = new THREE.Float32Attribute( geometry.skinIndices.length * 4, 4 );
			this.addAttribute( 'skinIndex', skinIndices.copyVector4sArray( geometry.skinIndices ) );

		}

		if ( geometry.skinWeights.length > 0 ) {

			var skinWeights = new THREE.Float32Attribute( geometry.skinWeights.length * 4, 4 );
			this.addAttribute( 'skinWeight', skinWeights.copyVector4sArray( geometry.skinWeights ) );

		}

		//

		if ( geometry.boundingSphere !== null ) {

			this.boundingSphere = geometry.boundingSphere.clone();

		}

		if ( geometry.boundingBox !== null ) {

			this.boundingBox = geometry.boundingBox.clone();

		}

		return this;

	},

	computeBoundingBox: function () {

		if ( this.boundingBox === null ) {

			this.boundingBox = new THREE.Box3();

		}

		var positions = this.attributes.position.array;

		if ( positions !== undefined ) {

			this.boundingBox.setFromArray( positions );

		} else {

			this.boundingBox.makeEmpty();

		}

		if ( isNaN( this.boundingBox.min.x ) || isNaN( this.boundingBox.min.y ) || isNaN( this.boundingBox.min.z ) ) {

			console.error( 'THREE.BufferGeometry.computeBoundingBox: Computed min/max have NaN values. The "position" attribute is likely to have NaN values.', this );

		}

	},

	computeBoundingSphere: function () {

		var box = new THREE.Box3();
		var vector = new THREE.Vector3();

		return function computeBoundingSphere() {

			if ( this.boundingSphere === null ) {

				this.boundingSphere = new THREE.Sphere();

			}

			var positions = this.attributes.position.array;

			if ( positions ) {

				var center = this.boundingSphere.center;

				box.setFromArray( positions );
				box.center( center );

				// hoping to find a boundingSphere with a radius smaller than the
				// boundingSphere of the boundingBox: sqrt(3) smaller in the best case

				var maxRadiusSq = 0;

				for ( var i = 0, il = positions.length; i < il; i += 3 ) {

					vector.fromArray( positions, i );
					maxRadiusSq = Math.max( maxRadiusSq, center.distanceToSquared( vector ) );

				}

				this.boundingSphere.radius = Math.sqrt( maxRadiusSq );

				if ( isNaN( this.boundingSphere.radius ) ) {

					console.error( 'THREE.BufferGeometry.computeBoundingSphere(): Computed radius is NaN. The "position" attribute is likely to have NaN values.', this );

				}

			}

		};

	}(),

	computeFaceNormals: function () {

		// backwards compatibility

	},

	computeVertexNormals: function () {

		var index = this.index;
		var attributes = this.attributes;
		var groups = this.groups;

		if ( attributes.position ) {

			var positions = attributes.position.array;

			if ( attributes.normal === undefined ) {

				this.addAttribute( 'normal', new THREE.BufferAttribute( new Float32Array( positions.length ), 3 ) );

			} else {

				// reset existing normals to zero

				var array = attributes.normal.array;

				for ( var i = 0, il = array.length; i < il; i ++ ) {

					array[ i ] = 0;

				}

			}

			var normals = attributes.normal.array;

			var vA, vB, vC,

			pA = new THREE.Vector3(),
			pB = new THREE.Vector3(),
			pC = new THREE.Vector3(),

			cb = new THREE.Vector3(),
			ab = new THREE.Vector3();

			// indexed elements

			if ( index ) {

				var indices = index.array;

				if ( groups.length === 0 ) {

					this.addGroup( 0, indices.length );

				}

				for ( var j = 0, jl = groups.length; j < jl; ++ j ) {

					var group = groups[ j ];

					var start = group.start;
					var count = group.count;

					for ( var i = start, il = start + count; i < il; i += 3 ) {

						vA = indices[ i + 0 ] * 3;
						vB = indices[ i + 1 ] * 3;
						vC = indices[ i + 2 ] * 3;

						pA.fromArray( positions, vA );
						pB.fromArray( positions, vB );
						pC.fromArray( positions, vC );

						cb.subVectors( pC, pB );
						ab.subVectors( pA, pB );
						cb.cross( ab );

						normals[ vA ] += cb.x;
						normals[ vA + 1 ] += cb.y;
						normals[ vA + 2 ] += cb.z;

						normals[ vB ] += cb.x;
						normals[ vB + 1 ] += cb.y;
						normals[ vB + 2 ] += cb.z;

						normals[ vC ] += cb.x;
						normals[ vC + 1 ] += cb.y;
						normals[ vC + 2 ] += cb.z;

					}

				}

			} else {

				// non-indexed elements (unconnected triangle soup)

				for ( var i = 0, il = positions.length; i < il; i += 9 ) {

					pA.fromArray( positions, i );
					pB.fromArray( positions, i + 3 );
					pC.fromArray( positions, i + 6 );

					cb.subVectors( pC, pB );
					ab.subVectors( pA, pB );
					cb.cross( ab );

					normals[ i ] = cb.x;
					normals[ i + 1 ] = cb.y;
					normals[ i + 2 ] = cb.z;

					normals[ i + 3 ] = cb.x;
					normals[ i + 4 ] = cb.y;
					normals[ i + 5 ] = cb.z;

					normals[ i + 6 ] = cb.x;
					normals[ i + 7 ] = cb.y;
					normals[ i + 8 ] = cb.z;

				}

			}

			this.normalizeNormals();

			attributes.normal.needsUpdate = true;

		}

	},

	merge: function ( geometry, offset ) {

		if ( geometry instanceof THREE.BufferGeometry === false ) {

			console.error( 'THREE.BufferGeometry.merge(): geometry not an instance of THREE.BufferGeometry.', geometry );
			return;

		}

		if ( offset === undefined ) offset = 0;

		var attributes = this.attributes;

		for ( var key in attributes ) {

			if ( geometry.attributes[ key ] === undefined ) continue;

			var attribute1 = attributes[ key ];
			var attributeArray1 = attribute1.array;

			var attribute2 = geometry.attributes[ key ];
			var attributeArray2 = attribute2.array;

			var attributeSize = attribute2.itemSize;

			for ( var i = 0, j = attributeSize * offset; i < attributeArray2.length; i ++, j ++ ) {

				attributeArray1[ j ] = attributeArray2[ i ];

			}

		}

		return this;

	},

	normalizeNormals: function () {

		var normals = this.attributes.normal.array;

		var x, y, z, n;

		for ( var i = 0, il = normals.length; i < il; i += 3 ) {

			x = normals[ i ];
			y = normals[ i + 1 ];
			z = normals[ i + 2 ];

			n = 1.0 / Math.sqrt( x * x + y * y + z * z );

			normals[ i ] *= n;
			normals[ i + 1 ] *= n;
			normals[ i + 2 ] *= n;

		}

	},

	toNonIndexed: function () {

		if ( this.index === null ) {

			console.warn( 'THREE.BufferGeometry.toNonIndexed(): Geometry is already non-indexed.' );
			return this;

		}

		var geometry2 = new THREE.BufferGeometry();

		var indices = this.index.array;
		var attributes = this.attributes;

		for ( var name in attributes ) {

			var attribute = attributes[ name ];

			var array = attribute.array;
			var itemSize = attribute.itemSize;

			var array2 = new array.constructor( indices.length * itemSize );

			var index = 0, index2 = 0;

			for ( var i = 0, l = indices.length; i < l; i ++ ) {

				index = indices[ i ] * itemSize;

				for ( var j = 0; j < itemSize; j ++ ) {

					array2[ index2 ++ ] = array[ index ++ ];

				}

			}

			geometry2.addAttribute( name, new THREE.BufferAttribute( array2, itemSize ) );

		}

		return geometry2;

	},

	toJSON: function () {

		var data = {
			metadata: {
				version: 4.4,
				type: 'BufferGeometry',
				generator: 'BufferGeometry.toJSON'
			}
		};

		// standard BufferGeometry serialization

		data.uuid = this.uuid;
		data.type = this.type;
		if ( this.name !== '' ) data.name = this.name;

		if ( this.parameters !== undefined ) {

			var parameters = this.parameters;

			for ( var key in parameters ) {

				if ( parameters[ key ] !== undefined ) data[ key ] = parameters[ key ];

			}

			return data;

		}

		data.data = { attributes: {} };

		var index = this.index;

		if ( index !== null ) {

			var array = Array.prototype.slice.call( index.array );

			data.data.index = {
				type: index.array.constructor.name,
				array: array
			};

		}

		var attributes = this.attributes;

		for ( var key in attributes ) {

			var attribute = attributes[ key ];

			var array = Array.prototype.slice.call( attribute.array );

			data.data.attributes[ key ] = {
				itemSize: attribute.itemSize,
				type: attribute.array.constructor.name,
				array: array,
				normalized: attribute.normalized
			};

		}

		var groups = this.groups;

		if ( groups.length > 0 ) {

			data.data.groups = JSON.parse( JSON.stringify( groups ) );

		}

		var boundingSphere = this.boundingSphere;

		if ( boundingSphere !== null ) {

			data.data.boundingSphere = {
				center: boundingSphere.center.toArray(),
				radius: boundingSphere.radius
			};

		}

		return data;

	},

	clone: function () {

		/*
		// Handle primitives

		var parameters = this.parameters;

		if ( parameters !== undefined ) {

			var values = [];

			for ( var key in parameters ) {

				values.push( parameters[ key ] );

			}

			var geometry = Object.create( this.constructor.prototype );
			this.constructor.apply( geometry, values );
			return geometry;

		}

		return new this.constructor().copy( this );
		*/

		return new THREE.BufferGeometry().copy( this );

	},

	copy: function ( source ) {

		var index = source.index;

		if ( index !== null ) {

			this.setIndex( index.clone() );

		}

		var attributes = source.attributes;

		for ( var name in attributes ) {

			var attribute = attributes[ name ];
			this.addAttribute( name, attribute.clone() );

		}

		var groups = source.groups;

		for ( var i = 0, l = groups.length; i < l; i ++ ) {

			var group = groups[ i ];
			this.addGroup( group.start, group.count, group.materialIndex );

		}

		return this;

	},

	dispose: function () {

		this.dispatchEvent( { type: 'dispose' } );

	}

} );

THREE.BufferGeometry.MaxIndex = 65535;

// File:src/core/InstancedBufferGeometry.js

/**
 * @author benaadams / https://twitter.com/ben_a_adams
 */

THREE.InstancedBufferGeometry = function () {

	THREE.BufferGeometry.call( this );

	this.type = 'InstancedBufferGeometry';
	this.maxInstancedCount = undefined;

};

THREE.InstancedBufferGeometry.prototype = Object.create( THREE.BufferGeometry.prototype );
THREE.InstancedBufferGeometry.prototype.constructor = THREE.InstancedBufferGeometry;

THREE.InstancedBufferGeometry.prototype.addGroup = function ( start, count, instances ) {

	this.groups.push( {

		start: start,
		count: count,
		instances: instances

	} );

};

THREE.InstancedBufferGeometry.prototype.copy = function ( source ) {

	var index = source.index;

	if ( index !== null ) {

		this.setIndex( index.clone() );

	}

	var attributes = source.attributes;

	for ( var name in attributes ) {

		var attribute = attributes[ name ];
		this.addAttribute( name, attribute.clone() );

	}

	var groups = source.groups;

	for ( var i = 0, l = groups.length; i < l; i ++ ) {

		var group = groups[ i ];
		this.addGroup( group.start, group.count, group.instances );

	}

	return this;

};

// File:src/core/Uniform.js

/**
 * @author mrdoob / http://mrdoob.com/
 */

THREE.Uniform = function ( value ) {

	if ( typeof value === 'string' ) {

		console.warn( 'THREE.Uniform: Type parameter is no longer needed.' );
		value = arguments[ 1 ];

	}

	this.value = value;

	this.dynamic = false;

};

THREE.Uniform.prototype = {

	constructor: THREE.Uniform,

	onUpdate: function ( callback ) {

		this.dynamic = true;
		this.onUpdateCallback = callback;

		return this;

	}

};

// File:src/animation/AnimationAction.js

/**
 *
 * Action provided by AnimationMixer for scheduling clip playback on specific
 * objects.
 *
 * @author Ben Houston / http://clara.io/
 * @author David Sarno / http://lighthaus.us/
 * @author tschw
 *
 */

THREE.AnimationAction = function() {

	throw new Error( "THREE.AnimationAction: " +
			"Use mixer.clipAction for construction." );

};

THREE.AnimationAction._new =
		function AnimationAction( mixer, clip, localRoot ) {

	this._mixer = mixer;
	this._clip = clip;
	this._localRoot = localRoot || null;

	var tracks = clip.tracks,
		nTracks = tracks.length,
		interpolants = new Array( nTracks );

	var interpolantSettings = {
			endingStart: 	THREE.ZeroCurvatureEnding,
			endingEnd:		THREE.ZeroCurvatureEnding
	};

	for ( var i = 0; i !== nTracks; ++ i ) {

		var interpolant = tracks[ i ].createInterpolant( null );
		interpolants[ i ] = interpolant;
		interpolant.settings = interpolantSettings;

	}

	this._interpolantSettings = interpolantSettings;

	this._interpolants = interpolants;	// bound by the mixer

	// inside: PropertyMixer (managed by the mixer)
	this._propertyBindings = new Array( nTracks );

	this._cacheIndex = null;			// for the memory manager
	this._byClipCacheIndex = null;		// for the memory manager

	this._timeScaleInterpolant = null;
	this._weightInterpolant = null;

	this.loop = THREE.LoopRepeat;
	this._loopCount = -1;

	// global mixer time when the action is to be started
	// it's set back to 'null' upon start of the action
	this._startTime = null;

	// scaled local time of the action
	// gets clamped or wrapped to 0..clip.duration according to loop
	this.time = 0;

	this.timeScale = 1;
	this._effectiveTimeScale = 1;

	this.weight = 1;
	this._effectiveWeight = 1;

	this.repetitions = Infinity; 		// no. of repetitions when looping

	this.paused = false;				// false -> zero effective time scale
	this.enabled = true;				// true -> zero effective weight

	this.clampWhenFinished 	= false;	// keep feeding the last frame?

	this.zeroSlopeAtStart 	= true;		// for smooth interpolation w/o separate
	this.zeroSlopeAtEnd		= true;		// clips for start, loop and end

};

THREE.AnimationAction._new.prototype = {

	constructor: THREE.AnimationAction._new,

	// State & Scheduling

	play: function() {

		this._mixer._activateAction( this );

		return this;

	},

	stop: function() {

		this._mixer._deactivateAction( this );

		return this.reset();

	},

	reset: function() {

		this.paused = false;
		this.enabled = true;

		this.time = 0;			// restart clip
		this._loopCount = -1;	// forget previous loops
		this._startTime = null;	// forget scheduling

		return this.stopFading().stopWarping();

	},

	isRunning: function() {

		var start = this._startTime;

		return this.enabled && ! this.paused && this.timeScale !== 0 &&
				this._startTime === null && this._mixer._isActiveAction( this );

	},

	// return true when play has been called
	isScheduled: function() {

		return this._mixer._isActiveAction( this );

	},

	startAt: function( time ) {

		this._startTime = time;

		return this;

	},

	setLoop: function( mode, repetitions ) {

		this.loop = mode;
		this.repetitions = repetitions;

		return this;

	},

	// Weight

	// set the weight stopping any scheduled fading
	// although .enabled = false yields an effective weight of zero, this
	// method does *not* change .enabled, because it would be confusing
	setEffectiveWeight: function( weight ) {

		this.weight = weight;

		// note: same logic as when updated at runtime
		this._effectiveWeight = this.enabled ? weight : 0;

		return this.stopFading();

	},

	// return the weight considering fading and .enabled
	getEffectiveWeight: function() {

		return this._effectiveWeight;

	},

	fadeIn: function( duration ) {

		return this._scheduleFading( duration, 0, 1 );

	},

	fadeOut: function( duration ) {

		return this._scheduleFading( duration, 1, 0 );

	},

	crossFadeFrom: function( fadeOutAction, duration, warp ) {

		var mixer = this._mixer;

		fadeOutAction.fadeOut( duration );
		this.fadeIn( duration );

		if( warp ) {

			var fadeInDuration = this._clip.duration,
				fadeOutDuration = fadeOutAction._clip.duration,

				startEndRatio = fadeOutDuration / fadeInDuration,
				endStartRatio = fadeInDuration / fadeOutDuration;

			fadeOutAction.warp( 1.0, startEndRatio, duration );
			this.warp( endStartRatio, 1.0, duration );

		}

		return this;

	},

	crossFadeTo: function( fadeInAction, duration, warp ) {

		return fadeInAction.crossFadeFrom( this, duration, warp );

	},

	stopFading: function() {

		var weightInterpolant = this._weightInterpolant;

		if ( weightInterpolant !== null ) {

			this._weightInterpolant = null;
			this._mixer._takeBackControlInterpolant( weightInterpolant );

		}

		return this;

	},

	// Time Scale Control

	// set the weight stopping any scheduled warping
	// although .paused = true yields an effective time scale of zero, this
	// method does *not* change .paused, because it would be confusing
	setEffectiveTimeScale: function( timeScale ) {

		this.timeScale = timeScale;
		this._effectiveTimeScale = this.paused ? 0 :timeScale;

		return this.stopWarping();

	},

	// return the time scale considering warping and .paused
	getEffectiveTimeScale: function() {

		return this._effectiveTimeScale;

	},

	setDuration: function( duration ) {

		this.timeScale = this._clip.duration / duration;

		return this.stopWarping();

	},

	syncWith: function( action ) {

		this.time = action.time;
		this.timeScale = action.timeScale;

		return this.stopWarping();

	},

	halt: function( duration ) {

		return this.warp( this._effectiveTimeScale, 0, duration );

	},

	warp: function( startTimeScale, endTimeScale, duration ) {

		var mixer = this._mixer, now = mixer.time,
			interpolant = this._timeScaleInterpolant,

			timeScale = this.timeScale;

		if ( interpolant === null ) {

			interpolant = mixer._lendControlInterpolant(),
			this._timeScaleInterpolant = interpolant;

		}

		var times = interpolant.parameterPositions,
			values = interpolant.sampleValues;

		times[ 0 ] = now;
		times[ 1 ] = now + duration;

		values[ 0 ] = startTimeScale / timeScale;
		values[ 1 ] = endTimeScale / timeScale;

		return this;

	},

	stopWarping: function() {

		var timeScaleInterpolant = this._timeScaleInterpolant;

		if ( timeScaleInterpolant !== null ) {

			this._timeScaleInterpolant = null;
			this._mixer._takeBackControlInterpolant( timeScaleInterpolant );

		}

		return this;

	},

	// Object Accessors

	getMixer: function() {

		return this._mixer;

	},

	getClip: function() {

		return this._clip;

	},

	getRoot: function() {

		return this._localRoot || this._mixer._root;

	},

	// Interna

	_update: function( time, deltaTime, timeDirection, accuIndex ) {
		// called by the mixer

		var startTime = this._startTime;

		if ( startTime !== null ) {

			// check for scheduled start of action

			var timeRunning = ( time - startTime ) * timeDirection;
			if ( timeRunning < 0 || timeDirection === 0 ) {

				return; // yet to come / don't decide when delta = 0

			}

			// start

			this._startTime = null; // unschedule
			deltaTime = timeDirection * timeRunning;

		}

		// apply time scale and advance time

		deltaTime *= this._updateTimeScale( time );
		var clipTime = this._updateTime( deltaTime );

		// note: _updateTime may disable the action resulting in
		// an effective weight of 0

		var weight = this._updateWeight( time );

		if ( weight > 0 ) {

			var interpolants = this._interpolants;
			var propertyMixers = this._propertyBindings;

			for ( var j = 0, m = interpolants.length; j !== m; ++ j ) {

				interpolants[ j ].evaluate( clipTime );
				propertyMixers[ j ].accumulate( accuIndex, weight );

			}

		}

	},

	_updateWeight: function( time ) {

		var weight = 0;

		if ( this.enabled ) {

			weight = this.weight;
			var interpolant = this._weightInterpolant;

			if ( interpolant !== null ) {

				var interpolantValue = interpolant.evaluate( time )[ 0 ];

				weight *= interpolantValue;

				if ( time > interpolant.parameterPositions[ 1 ] ) {

					this.stopFading();

					if ( interpolantValue === 0 ) {

						// faded out, disable
						this.enabled = false;

					}

				}

			}

		}

		this._effectiveWeight = weight;
		return weight;

	},

	_updateTimeScale: function( time ) {

		var timeScale = 0;

		if ( ! this.paused ) {

			timeScale = this.timeScale;

			var interpolant = this._timeScaleInterpolant;

			if ( interpolant !== null ) {

				var interpolantValue = interpolant.evaluate( time )[ 0 ];

				timeScale *= interpolantValue;

				if ( time > interpolant.parameterPositions[ 1 ] ) {

					this.stopWarping();

					if ( timeScale === 0 ) {

						// motion has halted, pause
						this.paused = true;

					} else {

						// warp done - apply final time scale
						this.timeScale = timeScale;

					}

				}

			}

		}

		this._effectiveTimeScale = timeScale;
		return timeScale;

	},

	_updateTime: function( deltaTime ) {

		var time = this.time + deltaTime;

		if ( deltaTime === 0 ) return time;

		var duration = this._clip.duration,

			loop = this.loop,
			loopCount = this._loopCount;

		if ( loop === THREE.LoopOnce ) {

			if ( loopCount === -1 ) {
				// just started

				this.loopCount = 0;
				this._setEndings( true, true, false );

			}

			handle_stop: {

				if ( time >= duration ) {

					time = duration;

				} else if ( time < 0 ) {

					time = 0;

				} else break handle_stop;

				if ( this.clampWhenFinished ) this.paused = true;
				else this.enabled = false;

				this._mixer.dispatchEvent( {
					type: 'finished', action: this,
					direction: deltaTime < 0 ? -1 : 1
				} );

			}

		} else { // repetitive Repeat or PingPong

			var pingPong = ( loop === THREE.LoopPingPong );

			if ( loopCount === -1 ) {
				// just started

				if ( deltaTime >= 0 ) {

					loopCount = 0;

					this._setEndings(
							true, this.repetitions === 0, pingPong );

				} else {

					// when looping in reverse direction, the initial
					// transition through zero counts as a repetition,
					// so leave loopCount at -1

					this._setEndings(
							this.repetitions === 0, true, pingPong );

				}

			}

			if ( time >= duration || time < 0 ) {
				// wrap around

				var loopDelta = Math.floor( time / duration ); // signed
				time -= duration * loopDelta;

				loopCount += Math.abs( loopDelta );

				var pending = this.repetitions - loopCount;

				if ( pending < 0 ) {
					// have to stop (switch state, clamp time, fire event)

					if ( this.clampWhenFinished ) this.paused = true;
					else this.enabled = false;

					time = deltaTime > 0 ? duration : 0;

					this._mixer.dispatchEvent( {
						type: 'finished', action: this,
						direction: deltaTime > 0 ? 1 : -1
					} );

				} else {
					// keep running

					if ( pending === 0 ) {
						// entering the last round

						var atStart = deltaTime < 0;
						this._setEndings( atStart, ! atStart, pingPong );

					} else {

						this._setEndings( false, false, pingPong );

					}

					this._loopCount = loopCount;

					this._mixer.dispatchEvent( {
						type: 'loop', action: this, loopDelta: loopDelta
					} );

				}

			}

			if ( pingPong && ( loopCount & 1 ) === 1 ) {
				// invert time for the "pong round"

				this.time = time;
				return duration - time;

			}

		}

		this.time = time;
		return time;

	},

	_setEndings: function( atStart, atEnd, pingPong ) {

		var settings = this._interpolantSettings;

		if ( pingPong ) {

			settings.endingStart 	= THREE.ZeroSlopeEnding;
			settings.endingEnd		= THREE.ZeroSlopeEnding;

		} else {

			// assuming for LoopOnce atStart == atEnd == true

			if ( atStart ) {

				settings.endingStart = this.zeroSlopeAtStart ?
						THREE.ZeroSlopeEnding : THREE.ZeroCurvatureEnding;

			} else {

				settings.endingStart = THREE.WrapAroundEnding;

			}

			if ( atEnd ) {

				settings.endingEnd = this.zeroSlopeAtEnd ?
						THREE.ZeroSlopeEnding : THREE.ZeroCurvatureEnding;

			} else {

				settings.endingEnd 	 = THREE.WrapAroundEnding;

			}

		}

	},

	_scheduleFading: function( duration, weightNow, weightThen ) {

		var mixer = this._mixer, now = mixer.time,
			interpolant = this._weightInterpolant;

		if ( interpolant === null ) {

			interpolant = mixer._lendControlInterpolant(),
			this._weightInterpolant = interpolant;

		}

		var times = interpolant.parameterPositions,
			values = interpolant.sampleValues;

		times[ 0 ] = now; 				values[ 0 ] = weightNow;
		times[ 1 ] = now + duration;	values[ 1 ] = weightThen;

		return this;

	}

};


// File:src/animation/AnimationClip.js

/**
 *
 * Reusable set of Tracks that represent an animation.
 *
 * @author Ben Houston / http://clara.io/
 * @author David Sarno / http://lighthaus.us/
 */

THREE.AnimationClip = function ( name, duration, tracks ) {

	this.name = name;
	this.tracks = tracks;
	this.duration = ( duration !== undefined ) ? duration : -1;

	this.uuid = THREE.Math.generateUUID();

	// this means it should figure out its duration by scanning the tracks
	if ( this.duration < 0 ) {

		this.resetDuration();

	}

	// maybe only do these on demand, as doing them here could potentially slow down loading
	// but leaving these here during development as this ensures a lot of testing of these functions
	this.trim();
	this.optimize();

};

THREE.AnimationClip.prototype = {

	constructor: THREE.AnimationClip,

	resetDuration: function() {

		var tracks = this.tracks,
			duration = 0;

		for ( var i = 0, n = tracks.length; i !== n; ++ i ) {

			var track = this.tracks[ i ];

			duration = Math.max(
					duration, track.times[ track.times.length - 1 ] );

		}

		this.duration = duration;

	},

	trim: function() {

		for ( var i = 0; i < this.tracks.length; i ++ ) {

			this.tracks[ i ].trim( 0, this.duration );

		}

		return this;

	},

	optimize: function() {

		for ( var i = 0; i < this.tracks.length; i ++ ) {

			this.tracks[ i ].optimize();

		}

		return this;

	}

};

// Static methods:

Object.assign( THREE.AnimationClip, {

	parse: function( json ) {

		var tracks = [],
			jsonTracks = json.tracks,
			frameTime = 1.0 / ( json.fps || 1.0 );

		for ( var i = 0, n = jsonTracks.length; i !== n; ++ i ) {

			tracks.push( THREE.KeyframeTrack.parse( jsonTracks[ i ] ).scale( frameTime ) );

		}

		return new THREE.AnimationClip( json.name, json.duration, tracks );

	},


	toJSON: function( clip ) {

		var tracks = [],
			clipTracks = clip.tracks;

		var json = {

			'name': clip.name,
			'duration': clip.duration,
			'tracks': tracks

		};

		for ( var i = 0, n = clipTracks.length; i !== n; ++ i ) {

			tracks.push( THREE.KeyframeTrack.toJSON( clipTracks[ i ] ) );

		}

		return json;

	},


	CreateFromMorphTargetSequence: function( name, morphTargetSequence, fps, noLoop ) {

		var numMorphTargets = morphTargetSequence.length;
		var tracks = [];

		for ( var i = 0; i < numMorphTargets; i ++ ) {

			var times = [];
			var values = [];

			times.push(
					( i + numMorphTargets - 1 ) % numMorphTargets,
					i,
					( i + 1 ) % numMorphTargets );

			values.push( 0, 1, 0 );

			var order = THREE.AnimationUtils.getKeyframeOrder( times );
			times = THREE.AnimationUtils.sortedArray( times, 1, order );
			values = THREE.AnimationUtils.sortedArray( values, 1, order );

			// if there is a key at the first frame, duplicate it as the
			// last frame as well for perfect loop.
			if ( ! noLoop && times[ 0 ] === 0 ) {

				times.push( numMorphTargets );
				values.push( values[ 0 ] );

			}

			tracks.push(
					new THREE.NumberKeyframeTrack(
						'.morphTargetInfluences[' + morphTargetSequence[ i ].name + ']',
						times, values
					).scale( 1.0 / fps ) );
		}

		return new THREE.AnimationClip( name, -1, tracks );

	},

	findByName: function( objectOrClipArray, name ) {

		var clipArray = objectOrClipArray;

		if ( ! Array.isArray( objectOrClipArray ) ) {

			var o = objectOrClipArray;
			clipArray = o.geometry && o.geometry.animations || o.animations;

		}

		for ( var i = 0; i < clipArray.length; i ++ ) {

			if ( clipArray[ i ].name === name ) {

				return clipArray[ i ];

			}
		}

		return null;

	},

	CreateClipsFromMorphTargetSequences: function( morphTargets, fps, noLoop ) {

		var animationToMorphTargets = {};

		// tested with https://regex101.com/ on trick sequences
		// such flamingo_flyA_003, flamingo_run1_003, crdeath0059
		var pattern = /^([\w-]*?)([\d]+)$/;

		// sort morph target names into animation groups based
		// patterns like Walk_001, Walk_002, Run_001, Run_002
		for ( var i = 0, il = morphTargets.length; i < il; i ++ ) {

			var morphTarget = morphTargets[ i ];
			var parts = morphTarget.name.match( pattern );

			if ( parts && parts.length > 1 ) {

				var name = parts[ 1 ];

				var animationMorphTargets = animationToMorphTargets[ name ];
				if ( ! animationMorphTargets ) {

					animationToMorphTargets[ name ] = animationMorphTargets = [];

				}

				animationMorphTargets.push( morphTarget );

			}

		}

		var clips = [];

		for ( var name in animationToMorphTargets ) {

			clips.push( THREE.AnimationClip.CreateFromMorphTargetSequence( name, animationToMorphTargets[ name ], fps, noLoop ) );

		}

		return clips;

	},

	// parse the animation.hierarchy format
	parseAnimation: function( animation, bones, nodeName ) {

		if ( ! animation ) {

			console.error( "  no animation in JSONLoader data" );
			return null;

		}

		var addNonemptyTrack = function(
				trackType, trackName, animationKeys, propertyName, destTracks ) {

			// only return track if there are actually keys.
			if ( animationKeys.length !== 0 ) {

				var times = [];
				var values = [];

				THREE.AnimationUtils.flattenJSON(
						animationKeys, times, values, propertyName );

				// empty keys are filtered out, so check again
				if ( times.length !== 0 ) {

					destTracks.push( new trackType( trackName, times, values ) );

				}

			}

		};

		var tracks = [];

		var clipName = animation.name || 'default';
		// automatic length determination in AnimationClip.
		var duration = animation.length || -1;
		var fps = animation.fps || 30;

		var hierarchyTracks = animation.hierarchy || [];

		for ( var h = 0; h < hierarchyTracks.length; h ++ ) {

			var animationKeys = hierarchyTracks[ h ].keys;

			// skip empty tracks
			if ( ! animationKeys || animationKeys.length === 0 ) continue;

			// process morph targets in a way exactly compatible
			// with AnimationHandler.init( animation )
			if ( animationKeys[0].morphTargets ) {

				// figure out all morph targets used in this track
				var morphTargetNames = {};
				for ( var k = 0; k < animationKeys.length; k ++ ) {

					if ( animationKeys[k].morphTargets ) {

						for ( var m = 0; m < animationKeys[k].morphTargets.length; m ++ ) {

							morphTargetNames[ animationKeys[k].morphTargets[m] ] = -1;
						}

					}

				}

				// create a track for each morph target with all zero
				// morphTargetInfluences except for the keys in which
				// the morphTarget is named.
				for ( var morphTargetName in morphTargetNames ) {

					var times = [];
					var values = [];

					for ( var m = 0;
							m !== animationKeys[k].morphTargets.length; ++ m ) {

						var animationKey = animationKeys[k];

						times.push( animationKey.time );
						values.push( ( animationKey.morphTarget === morphTargetName ) ? 1 : 0 );

					}

					tracks.push( new THREE.NumberKeyframeTrack(
							'.morphTargetInfluence[' + morphTargetName + ']', times, values ) );

				}

				duration = morphTargetNames.length * ( fps || 1.0 );

			} else {
				// ...assume skeletal animation

				var boneName = '.bones[' + bones[ h ].name + ']';

				addNonemptyTrack(
						THREE.VectorKeyframeTrack, boneName + '.position',
						animationKeys, 'pos', tracks );

				addNonemptyTrack(
						THREE.QuaternionKeyframeTrack, boneName + '.quaternion',
						animationKeys, 'rot', tracks );

				addNonemptyTrack(
						THREE.VectorKeyframeTrack, boneName + '.scale',
						animationKeys, 'scl', tracks );

			}

		}

		if ( tracks.length === 0 ) {

			return null;

		}

		var clip = new THREE.AnimationClip( clipName, duration, tracks );

		return clip;

	}

} );

// File:src/animation/AnimationMixer.js

/**
 *
 * Player for AnimationClips.
 *
 *
 * @author Ben Houston / http://clara.io/
 * @author David Sarno / http://lighthaus.us/
 * @author tschw
 */

THREE.AnimationMixer = function( root ) {

	this._root = root;
	this._initMemoryManager();
	this._accuIndex = 0;

	this.time = 0;

	this.timeScale = 1.0;

};

Object.assign( THREE.AnimationMixer.prototype, THREE.EventDispatcher.prototype, {

	// return an action for a clip optionally using a custom root target
	// object (this method allocates a lot of dynamic memory in case a
	// previously unknown clip/root combination is specified)
	clipAction: function( clip, optionalRoot ) {

		var root = optionalRoot || this._root,
			rootUuid = root.uuid,

			clipObject = typeof clip === 'string' ?
					THREE.AnimationClip.findByName( root, clip ) : clip,

			clipUuid = clipObject !== null ? clipObject.uuid : clip,

			actionsForClip = this._actionsByClip[ clipUuid ],
			prototypeAction = null;

		if ( actionsForClip !== undefined ) {

			var existingAction =
					actionsForClip.actionByRoot[ rootUuid ];

			if ( existingAction !== undefined ) {

				return existingAction;

			}

			// we know the clip, so we don't have to parse all
			// the bindings again but can just copy
			prototypeAction = actionsForClip.knownActions[ 0 ];

			// also, take the clip from the prototype action
			if ( clipObject === null )
				clipObject = prototypeAction._clip;

		}

		// clip must be known when specified via string
		if ( clipObject === null ) return null;

		// allocate all resources required to run it
		var newAction = new THREE.
				AnimationMixer._Action( this, clipObject, optionalRoot );

		this._bindAction( newAction, prototypeAction );

		// and make the action known to the memory manager
		this._addInactiveAction( newAction, clipUuid, rootUuid );

		return newAction;

	},

	// get an existing action
	existingAction: function( clip, optionalRoot ) {

		var root = optionalRoot || this._root,
			rootUuid = root.uuid,

			clipObject = typeof clip === 'string' ?
					THREE.AnimationClip.findByName( root, clip ) : clip,

			clipUuid = clipObject ? clipObject.uuid : clip,

			actionsForClip = this._actionsByClip[ clipUuid ];

		if ( actionsForClip !== undefined ) {

			return actionsForClip.actionByRoot[ rootUuid ] || null;

		}

		return null;

	},

	// deactivates all previously scheduled actions
	stopAllAction: function() {

		var actions = this._actions,
			nActions = this._nActiveActions,
			bindings = this._bindings,
			nBindings = this._nActiveBindings;

		this._nActiveActions = 0;
		this._nActiveBindings = 0;

		for ( var i = 0; i !== nActions; ++ i ) {

			actions[ i ].reset();

		}

		for ( var i = 0; i !== nBindings; ++ i ) {

			bindings[ i ].useCount = 0;

		}

		return this;

	},

	// advance the time and update apply the animation
	update: function( deltaTime ) {

		deltaTime *= this.timeScale;

		var actions = this._actions,
			nActions = this._nActiveActions,

			time = this.time += deltaTime,
			timeDirection = Math.sign( deltaTime ),

			accuIndex = this._accuIndex ^= 1;

		// run active actions

		for ( var i = 0; i !== nActions; ++ i ) {

			var action = actions[ i ];

			if ( action.enabled ) {

				action._update( time, deltaTime, timeDirection, accuIndex );

			}

		}

		// update scene graph

		var bindings = this._bindings,
			nBindings = this._nActiveBindings;

		for ( var i = 0; i !== nBindings; ++ i ) {

			bindings[ i ].apply( accuIndex );

		}

		return this;

	},

	// return this mixer's root target object
	getRoot: function() {

		return this._root;

	},

	// free all resources specific to a particular clip
	uncacheClip: function( clip ) {

		var actions = this._actions,
			clipUuid = clip.uuid,
			actionsByClip = this._actionsByClip,
			actionsForClip = actionsByClip[ clipUuid ];

		if ( actionsForClip !== undefined ) {

			// note: just calling _removeInactiveAction would mess up the
			// iteration state and also require updating the state we can
			// just throw away

			var actionsToRemove = actionsForClip.knownActions;

			for ( var i = 0, n = actionsToRemove.length; i !== n; ++ i ) {

				var action = actionsToRemove[ i ];

				this._deactivateAction( action );

				var cacheIndex = action._cacheIndex,
					lastInactiveAction = actions[ actions.length - 1 ];

				action._cacheIndex = null;
				action._byClipCacheIndex = null;

				lastInactiveAction._cacheIndex = cacheIndex;
				actions[ cacheIndex ] = lastInactiveAction;
				actions.pop();

				this._removeInactiveBindingsForAction( action );

			}

			delete actionsByClip[ clipUuid ];

		}

	},

	// free all resources specific to a particular root target object
	uncacheRoot: function( root ) {

		var rootUuid = root.uuid,
			actionsByClip = this._actionsByClip;

		for ( var clipUuid in actionsByClip ) {

			var actionByRoot = actionsByClip[ clipUuid ].actionByRoot,
				action = actionByRoot[ rootUuid ];

			if ( action !== undefined ) {

				this._deactivateAction( action );
				this._removeInactiveAction( action );

			}

		}

		var bindingsByRoot = this._bindingsByRootAndName,
			bindingByName = bindingsByRoot[ rootUuid ];

		if ( bindingByName !== undefined ) {

			for ( var trackName in bindingByName ) {

				var binding = bindingByName[ trackName ];
				binding.restoreOriginalState();
				this._removeInactiveBinding( binding );

			}

		}

	},

	// remove a targeted clip from the cache
	uncacheAction: function( clip, optionalRoot ) {

		var action = this.existingAction( clip, optionalRoot );

		if ( action !== null ) {

			this._deactivateAction( action );
			this._removeInactiveAction( action );

		}

	}

} );

THREE.AnimationMixer._Action = THREE.AnimationAction._new;

// Implementation details:

Object.assign( THREE.AnimationMixer.prototype, {

	_bindAction: function( action, prototypeAction ) {

		var root = action._localRoot || this._root,
			tracks = action._clip.tracks,
			nTracks = tracks.length,
			bindings = action._propertyBindings,
			interpolants = action._interpolants,
			rootUuid = root.uuid,
			bindingsByRoot = this._bindingsByRootAndName,
			bindingsByName = bindingsByRoot[ rootUuid ];

		if ( bindingsByName === undefined ) {

			bindingsByName = {};
			bindingsByRoot[ rootUuid ] = bindingsByName;

		}

		for ( var i = 0; i !== nTracks; ++ i ) {

			var track = tracks[ i ],
				trackName = track.name,
				binding = bindingsByName[ trackName ];

			if ( binding !== undefined ) {

				bindings[ i ] = binding;

			} else {

				binding = bindings[ i ];

				if ( binding !== undefined ) {

					// existing binding, make sure the cache knows

					if ( binding._cacheIndex === null ) {

						++ binding.referenceCount;
						this._addInactiveBinding( binding, rootUuid, trackName );

					}

					continue;

				}

				var path = prototypeAction && prototypeAction.
						_propertyBindings[ i ].binding.parsedPath;

				binding = new THREE.PropertyMixer(
						THREE.PropertyBinding.create( root, trackName, path ),
						track.ValueTypeName, track.getValueSize() );

				++ binding.referenceCount;
				this._addInactiveBinding( binding, rootUuid, trackName );

				bindings[ i ] = binding;

			}

			interpolants[ i ].resultBuffer = binding.buffer;

		}

	},

	_activateAction: function( action ) {

		if ( ! this._isActiveAction( action ) ) {

			if ( action._cacheIndex === null ) {

				// this action has been forgotten by the cache, but the user
				// appears to be still using it -> rebind

				var rootUuid = ( action._localRoot || this._root ).uuid,
					clipUuid = action._clip.uuid,
					actionsForClip = this._actionsByClip[ clipUuid ];

				this._bindAction( action,
						actionsForClip && actionsForClip.knownActions[ 0 ] );

				this._addInactiveAction( action, clipUuid, rootUuid );

			}

			var bindings = action._propertyBindings;

			// increment reference counts / sort out state
			for ( var i = 0, n = bindings.length; i !== n; ++ i ) {

				var binding = bindings[ i ];

				if ( binding.useCount ++ === 0 ) {

					this._lendBinding( binding );
					binding.saveOriginalState();

				}

			}

			this._lendAction( action );

		}

	},

	_deactivateAction: function( action ) {

		if ( this._isActiveAction( action ) ) {

			var bindings = action._propertyBindings;

			// decrement reference counts / sort out state
			for ( var i = 0, n = bindings.length; i !== n; ++ i ) {

				var binding = bindings[ i ];

				if ( -- binding.useCount === 0 ) {

					binding.restoreOriginalState();
					this._takeBackBinding( binding );

				}

			}

			this._takeBackAction( action );

		}

	},

	// Memory manager

	_initMemoryManager: function() {

		this._actions = []; // 'nActiveActions' followed by inactive ones
		this._nActiveActions = 0;

		this._actionsByClip = {};
		// inside:
		// {
		// 		knownActions: Array< _Action >	- used as prototypes
		// 		actionByRoot: _Action			- lookup
		// }


		this._bindings = []; // 'nActiveBindings' followed by inactive ones
		this._nActiveBindings = 0;

		this._bindingsByRootAndName = {}; // inside: Map< name, PropertyMixer >


		this._controlInterpolants = []; // same game as above
		this._nActiveControlInterpolants = 0;

		var scope = this;

		this.stats = {

			actions: {
				get total() { return scope._actions.length; },
				get inUse() { return scope._nActiveActions; }
			},
			bindings: {
				get total() { return scope._bindings.length; },
				get inUse() { return scope._nActiveBindings; }
			},
			controlInterpolants: {
				get total() { return scope._controlInterpolants.length; },
				get inUse() { return scope._nActiveControlInterpolants; }
			}

		};

	},

	// Memory management for _Action objects

	_isActiveAction: function( action ) {

		var index = action._cacheIndex;
		return index !== null && index < this._nActiveActions;

	},

	_addInactiveAction: function( action, clipUuid, rootUuid ) {

		var actions = this._actions,
			actionsByClip = this._actionsByClip,
			actionsForClip = actionsByClip[ clipUuid ];

		if ( actionsForClip === undefined ) {

			actionsForClip = {

				knownActions: [ action ],
				actionByRoot: {}

			};

			action._byClipCacheIndex = 0;

			actionsByClip[ clipUuid ] = actionsForClip;

		} else {

			var knownActions = actionsForClip.knownActions;

			action._byClipCacheIndex = knownActions.length;
			knownActions.push( action );

		}

		action._cacheIndex = actions.length;
		actions.push( action );

		actionsForClip.actionByRoot[ rootUuid ] = action;

	},

	_removeInactiveAction: function( action ) {

		var actions = this._actions,
			lastInactiveAction = actions[ actions.length - 1 ],
			cacheIndex = action._cacheIndex;

		lastInactiveAction._cacheIndex = cacheIndex;
		actions[ cacheIndex ] = lastInactiveAction;
		actions.pop();

		action._cacheIndex = null;


		var clipUuid = action._clip.uuid,
			actionsByClip = this._actionsByClip,
			actionsForClip = actionsByClip[ clipUuid ],
			knownActionsForClip = actionsForClip.knownActions,

			lastKnownAction =
				knownActionsForClip[ knownActionsForClip.length - 1 ],

			byClipCacheIndex = action._byClipCacheIndex;

		lastKnownAction._byClipCacheIndex = byClipCacheIndex;
		knownActionsForClip[ byClipCacheIndex ] = lastKnownAction;
		knownActionsForClip.pop();

		action._byClipCacheIndex = null;


		var actionByRoot = actionsForClip.actionByRoot,
			rootUuid = ( actions._localRoot || this._root ).uuid;

		delete actionByRoot[ rootUuid ];

		if ( knownActionsForClip.length === 0 ) {

			delete actionsByClip[ clipUuid ];

		}

		this._removeInactiveBindingsForAction( action );

	},

	_removeInactiveBindingsForAction: function( action ) {

		var bindings = action._propertyBindings;
		for ( var i = 0, n = bindings.length; i !== n; ++ i ) {

			var binding = bindings[ i ];

			if ( -- binding.referenceCount === 0 ) {

				this._removeInactiveBinding( binding );

			}

		}

	},

	_lendAction: function( action ) {

		// [ active actions |  inactive actions  ]
		// [  active actions >| inactive actions ]
		//                 s        a
		//                  <-swap->
		//                 a        s

		var actions = this._actions,
			prevIndex = action._cacheIndex,

			lastActiveIndex = this._nActiveActions ++,

			firstInactiveAction = actions[ lastActiveIndex ];

		action._cacheIndex = lastActiveIndex;
		actions[ lastActiveIndex ] = action;

		firstInactiveAction._cacheIndex = prevIndex;
		actions[ prevIndex ] = firstInactiveAction;

	},

	_takeBackAction: function( action ) {

		// [  active actions  | inactive actions ]
		// [ active actions |< inactive actions  ]
		//        a        s
		//         <-swap->
		//        s        a

		var actions = this._actions,
			prevIndex = action._cacheIndex,

			firstInactiveIndex = -- this._nActiveActions,

			lastActiveAction = actions[ firstInactiveIndex ];

		action._cacheIndex = firstInactiveIndex;
		actions[ firstInactiveIndex ] = action;

		lastActiveAction._cacheIndex = prevIndex;
		actions[ prevIndex ] = lastActiveAction;

	},

	// Memory management for PropertyMixer objects

	_addInactiveBinding: function( binding, rootUuid, trackName ) {

		var bindingsByRoot = this._bindingsByRootAndName,
			bindingByName = bindingsByRoot[ rootUuid ],

			bindings = this._bindings;

		if ( bindingByName === undefined ) {

			bindingByName = {};
			bindingsByRoot[ rootUuid ] = bindingByName;

		}

		bindingByName[ trackName ] = binding;

		binding._cacheIndex = bindings.length;
		bindings.push( binding );

	},

	_removeInactiveBinding: function( binding ) {

		var bindings = this._bindings,
			propBinding = binding.binding,
			rootUuid = propBinding.rootNode.uuid,
			trackName = propBinding.path,
			bindingsByRoot = this._bindingsByRootAndName,
			bindingByName = bindingsByRoot[ rootUuid ],

			lastInactiveBinding = bindings[ bindings.length - 1 ],
			cacheIndex = binding._cacheIndex;

		lastInactiveBinding._cacheIndex = cacheIndex;
		bindings[ cacheIndex ] = lastInactiveBinding;
		bindings.pop();

		delete bindingByName[ trackName ];

		remove_empty_map: {

			for ( var _ in bindingByName ) break remove_empty_map;

			delete bindingsByRoot[ rootUuid ];

		}

	},

	_lendBinding: function( binding ) {

		var bindings = this._bindings,
			prevIndex = binding._cacheIndex,

			lastActiveIndex = this._nActiveBindings ++,

			firstInactiveBinding = bindings[ lastActiveIndex ];

		binding._cacheIndex = lastActiveIndex;
		bindings[ lastActiveIndex ] = binding;

		firstInactiveBinding._cacheIndex = prevIndex;
		bindings[ prevIndex ] = firstInactiveBinding;

	},

	_takeBackBinding: function( binding ) {

		var bindings = this._bindings,
			prevIndex = binding._cacheIndex,

			firstInactiveIndex = -- this._nActiveBindings,

			lastActiveBinding = bindings[ firstInactiveIndex ];

		binding._cacheIndex = firstInactiveIndex;
		bindings[ firstInactiveIndex ] = binding;

		lastActiveBinding._cacheIndex = prevIndex;
		bindings[ prevIndex ] = lastActiveBinding;

	},


	// Memory management of Interpolants for weight and time scale

	_lendControlInterpolant: function() {

		var interpolants = this._controlInterpolants,
			lastActiveIndex = this._nActiveControlInterpolants ++,
			interpolant = interpolants[ lastActiveIndex ];

		if ( interpolant === undefined ) {

			interpolant = new THREE.LinearInterpolant(
					new Float32Array( 2 ), new Float32Array( 2 ),
						1, this._controlInterpolantsResultBuffer );

			interpolant.__cacheIndex = lastActiveIndex;
			interpolants[ lastActiveIndex ] = interpolant;

		}

		return interpolant;

	},

	_takeBackControlInterpolant: function( interpolant ) {

		var interpolants = this._controlInterpolants,
			prevIndex = interpolant.__cacheIndex,

			firstInactiveIndex = -- this._nActiveControlInterpolants,

			lastActiveInterpolant = interpolants[ firstInactiveIndex ];

		interpolant.__cacheIndex = firstInactiveIndex;
		interpolants[ firstInactiveIndex ] = interpolant;

		lastActiveInterpolant.__cacheIndex = prevIndex;
		interpolants[ prevIndex ] = lastActiveInterpolant;

	},

	_controlInterpolantsResultBuffer: new Float32Array( 1 )

} );

// File:src/animation/AnimationObjectGroup.js

/**
 *
 * A group of objects that receives a shared animation state.
 *
 * Usage:
 *
 * 	-	Add objects you would otherwise pass as 'root' to the
 * 		constructor or the .clipAction method of AnimationMixer.
 *
 * 	-	Instead pass this object as 'root'.
 *
 * 	-	You can also add and remove objects later when the mixer
 * 		is running.
 *
 * Note:
 *
 *  	Objects of this class appear as one object to the mixer,
 *  	so cache control of the individual objects must be done
 *  	on the group.
 *
 * Limitation:
 *
 * 	- 	The animated properties must be compatible among the
 * 		all objects in the group.
 *
 *  -	A single property can either be controlled through a
 *  	target group or directly, but not both.
 *
 * @author tschw
 */

THREE.AnimationObjectGroup = function( var_args ) {

	this.uuid = THREE.Math.generateUUID();

	// cached objects followed by the active ones
	this._objects = Array.prototype.slice.call( arguments );

	this.nCachedObjects_ = 0;			// threshold
	// note: read by PropertyBinding.Composite

	var indices = {};
	this._indicesByUUID = indices;		// for bookkeeping

	for ( var i = 0, n = arguments.length; i !== n; ++ i ) {

		indices[ arguments[ i ].uuid ] = i;

	}

	this._paths = [];					// inside: string
	this._parsedPaths = [];				// inside: { we don't care, here }
	this._bindings = []; 				// inside: Array< PropertyBinding >
	this._bindingsIndicesByPath = {}; 	// inside: indices in these arrays

	var scope = this;

	this.stats = {

		objects: {
			get total() { return scope._objects.length; },
			get inUse() { return this.total - scope.nCachedObjects_;  }
		},

		get bindingsPerObject() { return scope._bindings.length; }

	};

};

THREE.AnimationObjectGroup.prototype = {

	constructor: THREE.AnimationObjectGroup,

	add: function( var_args ) {

		var objects = this._objects,
			nObjects = objects.length,
			nCachedObjects = this.nCachedObjects_,
			indicesByUUID = this._indicesByUUID,
			paths = this._paths,
			parsedPaths = this._parsedPaths,
			bindings = this._bindings,
			nBindings = bindings.length;

		for ( var i = 0, n = arguments.length; i !== n; ++ i ) {

			var object = arguments[ i ],
				uuid = object.uuid,
				index = indicesByUUID[ uuid ];

			if ( index === undefined ) {

				// unknown object -> add it to the ACTIVE region

				index = nObjects ++;
				indicesByUUID[ uuid ] = index;
				objects.push( object );

				// accounting is done, now do the same for all bindings

				for ( var j = 0, m = nBindings; j !== m; ++ j ) {

					bindings[ j ].push(
							new THREE.PropertyBinding(
								object, paths[ j ], parsedPaths[ j ] ) );

				}

			} else if ( index < nCachedObjects ) {

				var knownObject = objects[ index ];

				// move existing object to the ACTIVE region

				var firstActiveIndex = -- nCachedObjects,
					lastCachedObject = objects[ firstActiveIndex ];

				indicesByUUID[ lastCachedObject.uuid ] = index;
				objects[ index ] = lastCachedObject;

				indicesByUUID[ uuid ] = firstActiveIndex;
				objects[ firstActiveIndex ] = object;

				// accounting is done, now do the same for all bindings

				for ( var j = 0, m = nBindings; j !== m; ++ j ) {

					var bindingsForPath = bindings[ j ],
						lastCached = bindingsForPath[ firstActiveIndex ],
						binding = bindingsForPath[ index ];

					bindingsForPath[ index ] = lastCached;

					if ( binding === undefined ) {

						// since we do not bother to create new bindings
						// for objects that are cached, the binding may
						// or may not exist

						binding = new THREE.PropertyBinding(
								object, paths[ j ], parsedPaths[ j ] );

					}

					bindingsForPath[ firstActiveIndex ] = binding;

				}

			} else if ( objects[ index ] !== knownObject) {

				console.error( "Different objects with the same UUID " +
						"detected. Clean the caches or recreate your " +
						"infrastructure when reloading scenes..." );

			} // else the object is already where we want it to be

		} // for arguments

		this.nCachedObjects_ = nCachedObjects;

	},

	remove: function( var_args ) {

		var objects = this._objects,
			nObjects = objects.length,
			nCachedObjects = this.nCachedObjects_,
			indicesByUUID = this._indicesByUUID,
			bindings = this._bindings,
			nBindings = bindings.length;

		for ( var i = 0, n = arguments.length; i !== n; ++ i ) {

			var object = arguments[ i ],
				uuid = object.uuid,
				index = indicesByUUID[ uuid ];

			if ( index !== undefined && index >= nCachedObjects ) {

				// move existing object into the CACHED region

				var lastCachedIndex = nCachedObjects ++,
					firstActiveObject = objects[ lastCachedIndex ];

				indicesByUUID[ firstActiveObject.uuid ] = index;
				objects[ index ] = firstActiveObject;

				indicesByUUID[ uuid ] = lastCachedIndex;
				objects[ lastCachedIndex ] = object;

				// accounting is done, now do the same for all bindings

				for ( var j = 0, m = nBindings; j !== m; ++ j ) {

					var bindingsForPath = bindings[ j ],
						firstActive = bindingsForPath[ lastCachedIndex ],
						binding = bindingsForPath[ index ];

					bindingsForPath[ index ] = firstActive;
					bindingsForPath[ lastCachedIndex ] = binding;

				}

			}

		} // for arguments

		this.nCachedObjects_ = nCachedObjects;

	},

	// remove & forget
	uncache: function( var_args ) {

		var objects = this._objects,
			nObjects = objects.length,
			nCachedObjects = this.nCachedObjects_,
			indicesByUUID = this._indicesByUUID,
			bindings = this._bindings,
			nBindings = bindings.length;

		for ( var i = 0, n = arguments.length; i !== n; ++ i ) {

			var object = arguments[ i ],
				uuid = object.uuid,
				index = indicesByUUID[ uuid ];

			if ( index !== undefined ) {

				delete indicesByUUID[ uuid ];

				if ( index < nCachedObjects ) {

					// object is cached, shrink the CACHED region

					var firstActiveIndex = -- nCachedObjects,
						lastCachedObject = objects[ firstActiveIndex ],
						lastIndex = -- nObjects,
						lastObject = objects[ lastIndex ];

					// last cached object takes this object's place
					indicesByUUID[ lastCachedObject.uuid ] = index;
					objects[ index ] = lastCachedObject;

					// last object goes to the activated slot and pop
					indicesByUUID[ lastObject.uuid ] = firstActiveIndex;
					objects[ firstActiveIndex ] = lastObject;
					objects.pop();

					// accounting is done, now do the same for all bindings

					for ( var j = 0, m = nBindings; j !== m; ++ j ) {

						var bindingsForPath = bindings[ j ],
							lastCached = bindingsForPath[ firstActiveIndex ],
							last = bindingsForPath[ lastIndex ];

						bindingsForPath[ index ] = lastCached;
						bindingsForPath[ firstActiveIndex ] = last;
						bindingsForPath.pop();

					}

				} else {

					// object is active, just swap with the last and pop

					var lastIndex = -- nObjects,
						lastObject = objects[ lastIndex ];

					indicesByUUID[ lastObject.uuid ] = index;
					objects[ index ] = lastObject;
					objects.pop();

					// accounting is done, now do the same for all bindings

					for ( var j = 0, m = nBindings; j !== m; ++ j ) {

						var bindingsForPath = bindings[ j ];

						bindingsForPath[ index ] = bindingsForPath[ lastIndex ];
						bindingsForPath.pop();

					}

				} // cached or active

			} // if object is known

		} // for arguments

		this.nCachedObjects_ = nCachedObjects;

	},

	// Internal interface used by befriended PropertyBinding.Composite:

	subscribe_: function( path, parsedPath ) {
		// returns an array of bindings for the given path that is changed
		// according to the contained objects in the group

		var indicesByPath = this._bindingsIndicesByPath,
			index = indicesByPath[ path ],
			bindings = this._bindings;

		if ( index !== undefined ) return bindings[ index ];

		var paths = this._paths,
			parsedPaths = this._parsedPaths,
			objects = this._objects,
			nObjects = objects.length,
			nCachedObjects = this.nCachedObjects_,
			bindingsForPath = new Array( nObjects );

		index = bindings.length;

		indicesByPath[ path ] = index;

		paths.push( path );
		parsedPaths.push( parsedPath );
		bindings.push( bindingsForPath );

		for ( var i = nCachedObjects,
				n = objects.length; i !== n; ++ i ) {

			var object = objects[ i ];

			bindingsForPath[ i ] =
					new THREE.PropertyBinding( object, path, parsedPath );

		}

		return bindingsForPath;

	},

	unsubscribe_: function( path ) {
		// tells the group to forget about a property path and no longer
		// update the array previously obtained with 'subscribe_'

		var indicesByPath = this._bindingsIndicesByPath,
			index = indicesByPath[ path ];

		if ( index !== undefined ) {

			var paths = this._paths,
				parsedPaths = this._parsedPaths,
				bindings = this._bindings,
				lastBindingsIndex = bindings.length - 1,
				lastBindings = bindings[ lastBindingsIndex ],
				lastBindingsPath = path[ lastBindingsIndex ];

			indicesByPath[ lastBindingsPath ] = index;

			bindings[ index ] = lastBindings;
			bindings.pop();

			parsedPaths[ index ] = parsedPaths[ lastBindingsIndex ];
			parsedPaths.pop();

			paths[ index ] = paths[ lastBindingsIndex ];
			paths.pop();

		}

	}

};


// File:src/animation/AnimationUtils.js

/**
 * @author tschw
 * @author Ben Houston / http://clara.io/
 * @author David Sarno / http://lighthaus.us/
 */

THREE.AnimationUtils = {

	// same as Array.prototype.slice, but also works on typed arrays
	arraySlice: function( array, from, to ) {

		if ( THREE.AnimationUtils.isTypedArray( array ) ) {

			return new array.constructor( array.subarray( from, to ) );

		}

		return array.slice( from, to );

	},

	// converts an array to a specific type
	convertArray: function( array, type, forceClone ) {

		if ( ! array || // let 'undefined' and 'null' pass
				! forceClone && array.constructor === type ) return array;

		if ( typeof type.BYTES_PER_ELEMENT === 'number' ) {

			return new type( array ); // create typed array

		}

		return Array.prototype.slice.call( array ); // create Array

	},

	isTypedArray: function( object ) {

		return ArrayBuffer.isView( object ) &&
				! ( object instanceof DataView );

	},

	// returns an array by which times and values can be sorted
	getKeyframeOrder: function( times ) {

		function compareTime( i, j ) {

			return times[ i ] - times[ j ];

		}

		var n = times.length;
		var result = new Array( n );
		for ( var i = 0; i !== n; ++ i ) result[ i ] = i;

		result.sort( compareTime );

		return result;

	},

	// uses the array previously returned by 'getKeyframeOrder' to sort data
	sortedArray: function( values, stride, order ) {

		var nValues = values.length;
		var result = new values.constructor( nValues );

		for ( var i = 0, dstOffset = 0; dstOffset !== nValues; ++ i ) {

			var srcOffset = order[ i ] * stride;

			for ( var j = 0; j !== stride; ++ j ) {

				result[ dstOffset ++ ] = values[ srcOffset + j ];

			}

		}

		return result;

	},

	// function for parsing AOS keyframe formats
	flattenJSON: function( jsonKeys, times, values, valuePropertyName ) {

		var i = 1, key = jsonKeys[ 0 ];

		while ( key !== undefined && key[ valuePropertyName ] === undefined ) {

			key = jsonKeys[ i ++ ];

		}

		if ( key === undefined ) return; // no data

		var value = key[ valuePropertyName ];
		if ( value === undefined ) return; // no data

		if ( Array.isArray( value ) ) {

			do {

				value = key[ valuePropertyName ];

				if ( value !== undefined ) {

					times.push( key.time );
					values.push.apply( values, value ); // push all elements

				}

				key = jsonKeys[ i ++ ];

			} while ( key !== undefined );

		} else if ( value.toArray !== undefined ) {
			// ...assume THREE.Math-ish

			do {

				value = key[ valuePropertyName ];

				if ( value !== undefined ) {

					times.push( key.time );
					value.toArray( values, values.length );

				}

				key = jsonKeys[ i ++ ];

			} while ( key !== undefined );

		} else {
			// otherwise push as-is

			do {

				value = key[ valuePropertyName ];

				if ( value !== undefined ) {

					times.push( key.time );
					values.push( value );

				}

				key = jsonKeys[ i ++ ];

			} while ( key !== undefined );

		}

	}

};

// File:src/animation/KeyframeTrack.js

/**
 *
 * A timed sequence of keyframes for a specific property.
 *
 *
 * @author Ben Houston / http://clara.io/
 * @author David Sarno / http://lighthaus.us/
 * @author tschw
 */

THREE.KeyframeTrack = function ( name, times, values, interpolation ) {

	if( name === undefined ) throw new Error( "track name is undefined" );

	if( times === undefined || times.length === 0 ) {

		throw new Error( "no keyframes in track named " + name );

	}

	this.name = name;

	this.times = THREE.AnimationUtils.convertArray( times, this.TimeBufferType );
	this.values = THREE.AnimationUtils.convertArray( values, this.ValueBufferType );

	this.setInterpolation( interpolation || this.DefaultInterpolation );

	this.validate();
	this.optimize();

};

THREE.KeyframeTrack.prototype = {

	constructor: THREE.KeyframeTrack,

	TimeBufferType: Float32Array,
	ValueBufferType: Float32Array,

	DefaultInterpolation: THREE.InterpolateLinear,

	InterpolantFactoryMethodDiscrete: function( result ) {

		return new THREE.DiscreteInterpolant(
				this.times, this.values, this.getValueSize(), result );

	},

	InterpolantFactoryMethodLinear: function( result ) {

		return new THREE.LinearInterpolant(
				this.times, this.values, this.getValueSize(), result );

	},

	InterpolantFactoryMethodSmooth: function( result ) {

		return new THREE.CubicInterpolant(
				this.times, this.values, this.getValueSize(), result );

	},

	setInterpolation: function( interpolation ) {

		var factoryMethod;

		switch ( interpolation ) {

			case THREE.InterpolateDiscrete:

				factoryMethod = this.InterpolantFactoryMethodDiscrete;

				break;

			case THREE.InterpolateLinear:

				factoryMethod = this.InterpolantFactoryMethodLinear;

				break;

			case THREE.InterpolateSmooth:

				factoryMethod = this.InterpolantFactoryMethodSmooth;

				break;

		}

		if ( factoryMethod === undefined ) {

			var message = "unsupported interpolation for " +
					this.ValueTypeName + " keyframe track named " + this.name;

			if ( this.createInterpolant === undefined ) {

				// fall back to default, unless the default itself is messed up
				if ( interpolation !== this.DefaultInterpolation ) {

					this.setInterpolation( this.DefaultInterpolation );

				} else {

					throw new Error( message ); // fatal, in this case

				}

			}

			console.warn( message );
			return;

		}

		this.createInterpolant = factoryMethod;

	},

	getInterpolation: function() {

		switch ( this.createInterpolant ) {

			case this.InterpolantFactoryMethodDiscrete:

				return THREE.InterpolateDiscrete;

			case this.InterpolantFactoryMethodLinear:

				return THREE.InterpolateLinear;

			case this.InterpolantFactoryMethodSmooth:

				return THREE.InterpolateSmooth;

		}

	},

	getValueSize: function() {

		return this.values.length / this.times.length;

	},

	// move all keyframes either forwards or backwards in time
	shift: function( timeOffset ) {

		if( timeOffset !== 0.0 ) {

			var times = this.times;

			for( var i = 0, n = times.length; i !== n; ++ i ) {

				times[ i ] += timeOffset;

			}

		}

		return this;

	},

	// scale all keyframe times by a factor (useful for frame <-> seconds conversions)
	scale: function( timeScale ) {

		if( timeScale !== 1.0 ) {

			var times = this.times;

			for( var i = 0, n = times.length; i !== n; ++ i ) {

				times[ i ] *= timeScale;

			}

		}

		return this;

	},

	// removes keyframes before and after animation without changing any values within the range [startTime, endTime].
	// IMPORTANT: We do not shift around keys to the start of the track time, because for interpolated keys this will change their values
	trim: function( startTime, endTime ) {

		var times = this.times,
			nKeys = times.length,
			from = 0,
			to = nKeys - 1;

		while ( from !== nKeys && times[ from ] < startTime ) ++ from;
		while ( to !== -1 && times[ to ] > endTime ) -- to;

		++ to; // inclusive -> exclusive bound

		if( from !== 0 || to !== nKeys ) {

			// empty tracks are forbidden, so keep at least one keyframe
			if ( from >= to ) to = Math.max( to , 1 ), from = to - 1;

			var stride = this.getValueSize();
			this.times = THREE.AnimationUtils.arraySlice( times, from, to );
			this.values = THREE.AnimationUtils.
					arraySlice( this.values, from * stride, to * stride );

		}

		return this;

	},

	// ensure we do not get a GarbageInGarbageOut situation, make sure tracks are at least minimally viable
	validate: function() {

		var valid = true;

		var valueSize = this.getValueSize();
		if ( valueSize - Math.floor( valueSize ) !== 0 ) {

			console.error( "invalid value size in track", this );
			valid = false;

		}

		var times = this.times,
			values = this.values,

			nKeys = times.length;

		if( nKeys === 0 ) {

			console.error( "track is empty", this );
			valid = false;

		}

		var prevTime = null;

		for( var i = 0; i !== nKeys; i ++ ) {

			var currTime = times[ i ];

			if ( typeof currTime === 'number' && isNaN( currTime ) ) {

				console.error( "time is not a valid number", this, i, currTime );
				valid = false;
				break;

			}

			if( prevTime !== null && prevTime > currTime ) {

				console.error( "out of order keys", this, i, currTime, prevTime );
				valid = false;
				break;

			}

			prevTime = currTime;

		}

		if ( values !== undefined ) {

			if ( THREE.AnimationUtils.isTypedArray( values ) ) {

				for ( var i = 0, n = values.length; i !== n; ++ i ) {

					var value = values[ i ];

					if ( isNaN( value ) ) {

						console.error( "value is not a valid number", this, i, value );
						valid = false;
						break;

					}

				}

			}

		}

		return valid;

	},

	// removes equivalent sequential keys as common in morph target sequences
	// (0,0,0,0,1,1,1,0,0,0,0,0,0,0) --> (0,0,1,1,0,0)
	optimize: function() {

		var times = this.times,
			values = this.values,
			stride = this.getValueSize(),

			writeIndex = 1;

		for( var i = 1, n = times.length - 1; i <= n; ++ i ) {

			var keep = false;

			var time = times[ i ];
			var timeNext = times[ i + 1 ];

			// remove adjacent keyframes scheduled at the same time

			if ( time !== timeNext && ( i !== 1 || time !== time[ 0 ] ) ) {

				// remove unnecessary keyframes same as their neighbors
				var offset = i * stride,
					offsetP = offset - stride,
					offsetN = offset + stride;

				for ( var j = 0; j !== stride; ++ j ) {

					var value = values[ offset + j ];

					if ( value !== values[ offsetP + j ] ||
							value !== values[ offsetN + j ] ) {

						keep = true;
						break;

					}

				}

			}

			// in-place compaction

			if ( keep ) {

				if ( i !== writeIndex ) {

					times[ writeIndex ] = times[ i ];

					var readOffset = i * stride,
						writeOffset = writeIndex * stride;

					for ( var j = 0; j !== stride; ++ j ) {

						values[ writeOffset + j ] = values[ readOffset + j ];

					}


				}

				++ writeIndex;

			}

		}

		if ( writeIndex !== times.length ) {

			this.times = THREE.AnimationUtils.arraySlice( times, 0, writeIndex );
			this.values = THREE.AnimationUtils.arraySlice( values, 0, writeIndex * stride );

		}

		return this;

	}

};

// Static methods:

Object.assign( THREE.KeyframeTrack, {

	// Serialization (in static context, because of constructor invocation
	// and automatic invocation of .toJSON):

	parse: function( json ) {

		if( json.type === undefined ) {

			throw new Error( "track type undefined, can not parse" );

		}

		var trackType = THREE.KeyframeTrack._getTrackTypeForValueTypeName( json.type );

		if ( json.times === undefined ) {

			var times = [], values = [];

			THREE.AnimationUtils.flattenJSON( json.keys, times, values, 'value' );

			json.times = times;
			json.values = values;

		}

		// derived classes can define a static parse method
		if ( trackType.parse !== undefined ) {

			return trackType.parse( json );

		} else {

			// by default, we asssume a constructor compatible with the base
			return new trackType(
					json.name, json.times, json.values, json.interpolation );

		}

	},

	toJSON: function( track ) {

		var trackType = track.constructor;

		var json;

		// derived classes can define a static toJSON method
		if ( trackType.toJSON !== undefined ) {

			json = trackType.toJSON( track );

		} else {

			// by default, we assume the data can be serialized as-is
			json = {

				'name': track.name,
				'times': THREE.AnimationUtils.convertArray( track.times, Array ),
				'values': THREE.AnimationUtils.convertArray( track.values, Array )

			};

			var interpolation = track.getInterpolation();

			if ( interpolation !== track.DefaultInterpolation ) {

				json.interpolation = interpolation;

			}

		}

		json.type = track.ValueTypeName; // mandatory

		return json;

	},

	_getTrackTypeForValueTypeName: function( typeName ) {

		switch( typeName.toLowerCase() ) {

			case "scalar":
			case "double":
			case "float":
			case "number":
			case "integer":

				return THREE.NumberKeyframeTrack;

			case "vector":
			case "vector2":
			case "vector3":
			case "vector4":

				return THREE.VectorKeyframeTrack;

			case "color":

				return THREE.ColorKeyframeTrack;

			case "quaternion":

				return THREE.QuaternionKeyframeTrack;

			case "bool":
			case "boolean":

				return THREE.BooleanKeyframeTrack;

			case "string":

				return THREE.StringKeyframeTrack;

		}

		throw new Error( "Unsupported typeName: " + typeName );

	}

} );

// File:src/animation/PropertyBinding.js

/**
 *
 * A reference to a real property in the scene graph.
 *
 *
 * @author Ben Houston / http://clara.io/
 * @author David Sarno / http://lighthaus.us/
 * @author tschw
 */

THREE.PropertyBinding = function ( rootNode, path, parsedPath ) {

	this.path = path;
	this.parsedPath = parsedPath ||
			THREE.PropertyBinding.parseTrackName( path );

	this.node = THREE.PropertyBinding.findNode(
			rootNode, this.parsedPath.nodeName ) || rootNode;

	this.rootNode = rootNode;

};

THREE.PropertyBinding.prototype = {

	constructor: THREE.PropertyBinding,

	getValue: function getValue_unbound( targetArray, offset ) {

		this.bind();
		this.getValue( targetArray, offset );

		// Note: This class uses a State pattern on a per-method basis:
		// 'bind' sets 'this.getValue' / 'setValue' and shadows the
		// prototype version of these methods with one that represents
		// the bound state. When the property is not found, the methods
		// become no-ops.

	},

	setValue: function getValue_unbound( sourceArray, offset ) {

		this.bind();
		this.setValue( sourceArray, offset );

	},

	// create getter / setter pair for a property in the scene graph
	bind: function() {

		var targetObject = this.node,
			parsedPath = this.parsedPath,

			objectName = parsedPath.objectName,
			propertyName = parsedPath.propertyName,
			propertyIndex = parsedPath.propertyIndex;

		if ( ! targetObject ) {

			targetObject = THREE.PropertyBinding.findNode(
					this.rootNode, parsedPath.nodeName ) || this.rootNode;

			this.node = targetObject;

		}

		// set fail state so we can just 'return' on error
		this.getValue = this._getValue_unavailable;
		this.setValue = this._setValue_unavailable;

 		// ensure there is a value node
		if ( ! targetObject ) {

			console.error( "  trying to update node for track: " + this.path + " but it wasn't found." );
			return;

		}

		if( objectName ) {

			var objectIndex = parsedPath.objectIndex;

			// special cases were we need to reach deeper into the hierarchy to get the face materials....
			switch ( objectName ) {

				case 'materials':

					if( ! targetObject.material ) {

						console.error( '  can not bind to material as node does not have a material', this );
						return;

					}

					if( ! targetObject.material.materials ) {

						console.error( '  can not bind to material.materials as node.material does not have a materials array', this );
						return;

					}

					targetObject = targetObject.material.materials;

					break;

				case 'bones':

					if( ! targetObject.skeleton ) {

						console.error( '  can not bind to bones as node does not have a skeleton', this );
						return;

					}

					// potential future optimization: skip this if propertyIndex is already an integer
					// and convert the integer string to a true integer.

					targetObject = targetObject.skeleton.bones;

					// support resolving morphTarget names into indices.
					for ( var i = 0; i < targetObject.length; i ++ ) {

						if ( targetObject[i].name === objectIndex ) {

							objectIndex = i;
							break;

						}

					}

					break;

				default:

					if ( targetObject[ objectName ] === undefined ) {

						console.error( '  can not bind to objectName of node, undefined', this );
						return;

					}

					targetObject = targetObject[ objectName ];

			}


			if ( objectIndex !== undefined ) {

				if( targetObject[ objectIndex ] === undefined ) {

					console.error( "  trying to bind to objectIndex of objectName, but is undefined:", this, targetObject );
					return;

				}

				targetObject = targetObject[ objectIndex ];

			}

		}

		// resolve property
		var nodeProperty = targetObject[ propertyName ];

		if ( ! nodeProperty ) {

			var nodeName = parsedPath.nodeName;

			console.error( "  trying to update property for track: " + nodeName +
					'.' + propertyName + " but it wasn't found.", targetObject );
			return;

		}

		// determine versioning scheme
		var versioning = this.Versioning.None;

		if ( targetObject.needsUpdate !== undefined ) { // material

			versioning = this.Versioning.NeedsUpdate;
			this.targetObject = targetObject;

		} else if ( targetObject.matrixWorldNeedsUpdate !== undefined ) { // node transform

			versioning = this.Versioning.MatrixWorldNeedsUpdate;
			this.targetObject = targetObject;

		}

		// determine how the property gets bound
		var bindingType = this.BindingType.Direct;

		if ( propertyIndex !== undefined ) {
			// access a sub element of the property array (only primitives are supported right now)

			if ( propertyName === "morphTargetInfluences" ) {
				// potential optimization, skip this if propertyIndex is already an integer, and convert the integer string to a true integer.

				// support resolving morphTarget names into indices.
				if ( ! targetObject.geometry ) {

					console.error( '  can not bind to morphTargetInfluences becasuse node does not have a geometry', this );
					return;

				}

				if ( ! targetObject.geometry.morphTargets ) {

					console.error( '  can not bind to morphTargetInfluences becasuse node does not have a geometry.morphTargets', this );
					return;

				}

				for ( var i = 0; i < this.node.geometry.morphTargets.length; i ++ ) {

					if ( targetObject.geometry.morphTargets[i].name === propertyIndex ) {

						propertyIndex = i;
						break;

					}

				}

			}

			bindingType = this.BindingType.ArrayElement;

			this.resolvedProperty = nodeProperty;
			this.propertyIndex = propertyIndex;

		} else if ( nodeProperty.fromArray !== undefined && nodeProperty.toArray !== undefined ) {
			// must use copy for Object3D.Euler/Quaternion

			bindingType = this.BindingType.HasFromToArray;

			this.resolvedProperty = nodeProperty;

		} else if ( nodeProperty.length !== undefined ) {

			bindingType = this.BindingType.EntireArray;

			this.resolvedProperty = nodeProperty;

		} else {

			this.propertyName = propertyName;

		}

		// select getter / setter
		this.getValue = this.GetterByBindingType[ bindingType ];
		this.setValue = this.SetterByBindingTypeAndVersioning[ bindingType ][ versioning ];

	},

	unbind: function() {

		this.node = null;

		// back to the prototype version of getValue / setValue
		// note: avoiding to mutate the shape of 'this' via 'delete'
		this.getValue = this._getValue_unbound;
		this.setValue = this._setValue_unbound;

	}

};

Object.assign( THREE.PropertyBinding.prototype, { // prototype, continued

	// these are used to "bind" a nonexistent property
	_getValue_unavailable: function() {},
	_setValue_unavailable: function() {},

	// initial state of these methods that calls 'bind'
	_getValue_unbound: THREE.PropertyBinding.prototype.getValue,
	_setValue_unbound: THREE.PropertyBinding.prototype.setValue,

	BindingType: {
		Direct: 0,
		EntireArray: 1,
		ArrayElement: 2,
		HasFromToArray: 3
	},

	Versioning: {
		None: 0,
		NeedsUpdate: 1,
		MatrixWorldNeedsUpdate: 2
	},

	GetterByBindingType: [

		function getValue_direct( buffer, offset ) {

			buffer[ offset ] = this.node[ this.propertyName ];

		},

		function getValue_array( buffer, offset ) {

			var source = this.resolvedProperty;

			for ( var i = 0, n = source.length; i !== n; ++ i ) {

				buffer[ offset ++ ] = source[ i ];

			}

		},

		function getValue_arrayElement( buffer, offset ) {

			buffer[ offset ] = this.resolvedProperty[ this.propertyIndex ];

		},

		function getValue_toArray( buffer, offset ) {

			this.resolvedProperty.toArray( buffer, offset );

		}

	],

	SetterByBindingTypeAndVersioning: [

		[
			// Direct

			function setValue_direct( buffer, offset ) {

				this.node[ this.propertyName ] = buffer[ offset ];

			},

			function setValue_direct_setNeedsUpdate( buffer, offset ) {

				this.node[ this.propertyName ] = buffer[ offset ];
				this.targetObject.needsUpdate = true;

			},

			function setValue_direct_setMatrixWorldNeedsUpdate( buffer, offset ) {

				this.node[ this.propertyName ] = buffer[ offset ];
				this.targetObject.matrixWorldNeedsUpdate = true;

			}

		], [

			// EntireArray

			function setValue_array( buffer, offset ) {

				var dest = this.resolvedProperty;

				for ( var i = 0, n = dest.length; i !== n; ++ i ) {

					dest[ i ] = buffer[ offset ++ ];

				}

			},

			function setValue_array_setNeedsUpdate( buffer, offset ) {

				var dest = this.resolvedProperty;

				for ( var i = 0, n = dest.length; i !== n; ++ i ) {

					dest[ i ] = buffer[ offset ++ ];

				}

				this.targetObject.needsUpdate = true;

			},

			function setValue_array_setMatrixWorldNeedsUpdate( buffer, offset ) {

				var dest = this.resolvedProperty;

				for ( var i = 0, n = dest.length; i !== n; ++ i ) {

					dest[ i ] = buffer[ offset ++ ];

				}

				this.targetObject.matrixWorldNeedsUpdate = true;

			}

		], [

			// ArrayElement

			function setValue_arrayElement( buffer, offset ) {

				this.resolvedProperty[ this.propertyIndex ] = buffer[ offset ];

			},

			function setValue_arrayElement_setNeedsUpdate( buffer, offset ) {

				this.resolvedProperty[ this.propertyIndex ] = buffer[ offset ];
				this.targetObject.needsUpdate = true;

			},

			function setValue_arrayElement_setMatrixWorldNeedsUpdate( buffer, offset ) {

				this.resolvedProperty[ this.propertyIndex ] = buffer[ offset ];
				this.targetObject.matrixWorldNeedsUpdate = true;

			}

		], [

			// HasToFromArray

			function setValue_fromArray( buffer, offset ) {

				this.resolvedProperty.fromArray( buffer, offset );

			},

			function setValue_fromArray_setNeedsUpdate( buffer, offset ) {

				this.resolvedProperty.fromArray( buffer, offset );
				this.targetObject.needsUpdate = true;

			},

			function setValue_fromArray_setMatrixWorldNeedsUpdate( buffer, offset ) {

				this.resolvedProperty.fromArray( buffer, offset );
				this.targetObject.matrixWorldNeedsUpdate = true;

			}

		]

	]

} );

THREE.PropertyBinding.Composite =
		function( targetGroup, path, optionalParsedPath ) {

	var parsedPath = optionalParsedPath ||
			THREE.PropertyBinding.parseTrackName( path );

	this._targetGroup = targetGroup;
	this._bindings = targetGroup.subscribe_( path, parsedPath );

};

THREE.PropertyBinding.Composite.prototype = {

	constructor: THREE.PropertyBinding.Composite,

	getValue: function( array, offset ) {

		this.bind(); // bind all binding

		var firstValidIndex = this._targetGroup.nCachedObjects_,
			binding = this._bindings[ firstValidIndex ];

		// and only call .getValue on the first
		if ( binding !== undefined ) binding.getValue( array, offset );

	},

	setValue: function( array, offset ) {

		var bindings = this._bindings;

		for ( var i = this._targetGroup.nCachedObjects_,
				n = bindings.length; i !== n; ++ i ) {

			bindings[ i ].setValue( array, offset );

		}

	},

	bind: function() {

		var bindings = this._bindings;

		for ( var i = this._targetGroup.nCachedObjects_,
				n = bindings.length; i !== n; ++ i ) {

			bindings[ i ].bind();

		}

	},

	unbind: function() {

		var bindings = this._bindings;

		for ( var i = this._targetGroup.nCachedObjects_,
				n = bindings.length; i !== n; ++ i ) {

			bindings[ i ].unbind();

		}

	}

};

THREE.PropertyBinding.create = function( root, path, parsedPath ) {

	if ( ! ( root instanceof THREE.AnimationObjectGroup ) ) {

		return new THREE.PropertyBinding( root, path, parsedPath );

	} else {

		return new THREE.PropertyBinding.Composite( root, path, parsedPath );

	}

};

THREE.PropertyBinding.parseTrackName = function( trackName ) {

	// matches strings in the form of:
	//    nodeName.property
	//    nodeName.property[accessor]
	//    nodeName.material.property[accessor]
	//    uuid.property[accessor]
	//    uuid.objectName[objectIndex].propertyName[propertyIndex]
	//    parentName/nodeName.property
	//    parentName/parentName/nodeName.property[index]
	//	  .bone[Armature.DEF_cog].position
	// created and tested via https://regex101.com/#javascript

	var re = /^(([\w]+\/)*)([\w-\d]+)?(\.([\w]+)(\[([\w\d\[\]\_.:\- ]+)\])?)?(\.([\w.]+)(\[([\w\d\[\]\_. ]+)\])?)$/;
	var matches = re.exec(trackName);

	if( ! matches ) {
		throw new Error( "cannot parse trackName at all: " + trackName );
	}

    if (matches.index === re.lastIndex) {
        re.lastIndex++;
    }

	var results = {
		// directoryName: matches[1], // (tschw) currently unused
		nodeName: matches[3], 	// allowed to be null, specified root node.
		objectName: matches[5],
		objectIndex: matches[7],
		propertyName: matches[9],
		propertyIndex: matches[11]	// allowed to be null, specifies that the whole property is set.
	};

	if( results.propertyName === null || results.propertyName.length === 0 ) {
		throw new Error( "can not parse propertyName from trackName: " + trackName );
	}

	return results;

};

THREE.PropertyBinding.findNode = function( root, nodeName ) {

	if( ! nodeName || nodeName === "" || nodeName === "root" || nodeName === "." || nodeName === -1 || nodeName === root.name || nodeName === root.uuid ) {

		return root;

	}

	// search into skeleton bones.
	if( root.skeleton ) {

		var searchSkeleton = function( skeleton ) {

			for( var i = 0; i < skeleton.bones.length; i ++ ) {

				var bone = skeleton.bones[i];

				if( bone.name === nodeName ) {

					return bone;

				}
			}

			return null;

		};

		var bone = searchSkeleton( root.skeleton );

		if( bone ) {

			return bone;

		}
	}

	// search into node subtree.
	if( root.children ) {

		var searchNodeSubtree = function( children ) {

			for( var i = 0; i < children.length; i ++ ) {

				var childNode = children[i];

				if( childNode.name === nodeName || childNode.uuid === nodeName ) {

					return childNode;

				}

				var result = searchNodeSubtree( childNode.children );

				if( result ) return result;

			}

			return null;

		};

		var subTreeNode = searchNodeSubtree( root.children );

		if( subTreeNode ) {

			return subTreeNode;

		}

	}

	return null;

};

// File:src/animation/PropertyMixer.js

/**
 *
 * Buffered scene graph property that allows weighted accumulation.
 *
 *
 * @author Ben Houston / http://clara.io/
 * @author David Sarno / http://lighthaus.us/
 * @author tschw
 */

THREE.PropertyMixer = function ( binding, typeName, valueSize ) {

	this.binding = binding;
	this.valueSize = valueSize;

	var bufferType = Float64Array,
		mixFunction;

	switch ( typeName ) {

		case 'quaternion':			mixFunction = this._slerp;		break;

		case 'string':
		case 'bool':

			bufferType = Array,		mixFunction = this._select;		break;

		default:					mixFunction = this._lerp;

	}

	this.buffer = new bufferType( valueSize * 4 );
	// layout: [ incoming | accu0 | accu1 | orig ]
	//
	// interpolators can use .buffer as their .result
	// the data then goes to 'incoming'
	//
	// 'accu0' and 'accu1' are used frame-interleaved for
	// the cumulative result and are compared to detect
	// changes
	//
	// 'orig' stores the original state of the property

	this._mixBufferRegion = mixFunction;

	this.cumulativeWeight = 0;

	this.useCount = 0;
	this.referenceCount = 0;

};

THREE.PropertyMixer.prototype = {

	constructor: THREE.PropertyMixer,

	// accumulate data in the 'incoming' region into 'accu<i>'
	accumulate: function( accuIndex, weight ) {

		// note: happily accumulating nothing when weight = 0, the caller knows
		// the weight and shouldn't have made the call in the first place

		var buffer = this.buffer,
			stride = this.valueSize,
			offset = accuIndex * stride + stride,

			currentWeight = this.cumulativeWeight;

		if ( currentWeight === 0 ) {

			// accuN := incoming * weight

			for ( var i = 0; i !== stride; ++ i ) {

				buffer[ offset + i ] = buffer[ i ];

			}

			currentWeight = weight;

		} else {

			// accuN := accuN + incoming * weight

			currentWeight += weight;
			var mix = weight / currentWeight;
			this._mixBufferRegion( buffer, offset, 0, mix, stride );

		}

		this.cumulativeWeight = currentWeight;

	},

	// apply the state of 'accu<i>' to the binding when accus differ
	apply: function( accuIndex ) {

		var stride = this.valueSize,
			buffer = this.buffer,
			offset = accuIndex * stride + stride,

			weight = this.cumulativeWeight,

			binding = this.binding;

		this.cumulativeWeight = 0;

		if ( weight < 1 ) {

			// accuN := accuN + original * ( 1 - cumulativeWeight )

			var originalValueOffset = stride * 3;

			this._mixBufferRegion(
					buffer, offset, originalValueOffset, 1 - weight, stride );

		}

		for ( var i = stride, e = stride + stride; i !== e; ++ i ) {

			if ( buffer[ i ] !== buffer[ i + stride ] ) {

				// value has changed -> update scene graph

				binding.setValue( buffer, offset );
				break;

			}

		}

	},

	// remember the state of the bound property and copy it to both accus
	saveOriginalState: function() {

		var binding = this.binding;

		var buffer = this.buffer,
			stride = this.valueSize,

			originalValueOffset = stride * 3;

		binding.getValue( buffer, originalValueOffset );

		// accu[0..1] := orig -- initially detect changes against the original
		for ( var i = stride, e = originalValueOffset; i !== e; ++ i ) {

			buffer[ i ] = buffer[ originalValueOffset + ( i % stride ) ];

		}

		this.cumulativeWeight = 0;

	},

	// apply the state previously taken via 'saveOriginalState' to the binding
	restoreOriginalState: function() {

		var originalValueOffset = this.valueSize * 3;
		this.binding.setValue( this.buffer, originalValueOffset );

	},


	// mix functions

	_select: function( buffer, dstOffset, srcOffset, t, stride ) {

		if ( t >= 0.5 ) {

			for ( var i = 0; i !== stride; ++ i ) {

				buffer[ dstOffset + i ] = buffer[ srcOffset + i ];

			}

		}

	},

	_slerp: function( buffer, dstOffset, srcOffset, t, stride ) {

		THREE.Quaternion.slerpFlat( buffer, dstOffset,
				buffer, dstOffset, buffer, srcOffset, t );

	},

	_lerp: function( buffer, dstOffset, srcOffset, t, stride ) {

		var s = 1 - t;

		for ( var i = 0; i !== stride; ++ i ) {

			var j = dstOffset + i;

			buffer[ j ] = buffer[ j ] * s + buffer[ srcOffset + i ] * t;

		}

	}

};

// File:src/animation/tracks/BooleanKeyframeTrack.js

/**
 *
 * A Track of Boolean keyframe values.
 *
 *
 * @author Ben Houston / http://clara.io/
 * @author David Sarno / http://lighthaus.us/
 * @author tschw
 */

THREE.BooleanKeyframeTrack = function ( name, times, values ) {

	THREE.KeyframeTrack.call( this, name, times, values );

};

THREE.BooleanKeyframeTrack.prototype =
		Object.assign( Object.create( THREE.KeyframeTrack.prototype ), {

	constructor: THREE.BooleanKeyframeTrack,

	ValueTypeName: 'bool',
	ValueBufferType: Array,

	DefaultInterpolation: THREE.InterpolateDiscrete,

	InterpolantFactoryMethodLinear: undefined,
	InterpolantFactoryMethodSmooth: undefined

	// Note: Actually this track could have a optimized / compressed
	// representation of a single value and a custom interpolant that
	// computes "firstValue ^ isOdd( index )".

} );

// File:src/animation/tracks/ColorKeyframeTrack.js

/**
 *
 * A Track of keyframe values that represent color.
 *
 *
 * @author Ben Houston / http://clara.io/
 * @author David Sarno / http://lighthaus.us/
 * @author tschw
 */

THREE.ColorKeyframeTrack = function ( name, times, values, interpolation ) {

	THREE.KeyframeTrack.call( this, name, times, values, interpolation );

};

THREE.ColorKeyframeTrack.prototype =
		Object.assign( Object.create( THREE.KeyframeTrack.prototype ), {

	constructor: THREE.ColorKeyframeTrack,

	ValueTypeName: 'color'

	// ValueBufferType is inherited

	// DefaultInterpolation is inherited


	// Note: Very basic implementation and nothing special yet.
	// However, this is the place for color space parameterization.

} );

// File:src/animation/tracks/NumberKeyframeTrack.js

/**
 *
 * A Track of numeric keyframe values.
 *
 * @author Ben Houston / http://clara.io/
 * @author David Sarno / http://lighthaus.us/
 * @author tschw
 */

THREE.NumberKeyframeTrack = function ( name, times, values, interpolation ) {

	THREE.KeyframeTrack.call( this, name, times, values, interpolation );

};

THREE.NumberKeyframeTrack.prototype =
		Object.assign( Object.create( THREE.KeyframeTrack.prototype ), {

	constructor: THREE.NumberKeyframeTrack,

	ValueTypeName: 'number',

	// ValueBufferType is inherited

	// DefaultInterpolation is inherited

} );

// File:src/animation/tracks/QuaternionKeyframeTrack.js

/**
 *
 * A Track of quaternion keyframe values.
 *
 * @author Ben Houston / http://clara.io/
 * @author David Sarno / http://lighthaus.us/
 * @author tschw
 */

THREE.QuaternionKeyframeTrack = function ( name, times, values, interpolation ) {

	THREE.KeyframeTrack.call( this, name, times, values, interpolation );

};

THREE.QuaternionKeyframeTrack.prototype =
		Object.assign( Object.create( THREE.KeyframeTrack.prototype ), {

	constructor: THREE.QuaternionKeyframeTrack,

	ValueTypeName: 'quaternion',

	// ValueBufferType is inherited

	DefaultInterpolation: THREE.InterpolateLinear,

	InterpolantFactoryMethodLinear: function( result ) {

		return new THREE.QuaternionLinearInterpolant(
				this.times, this.values, this.getValueSize(), result );

	},

	InterpolantFactoryMethodSmooth: undefined // not yet implemented

} );

// File:src/animation/tracks/StringKeyframeTrack.js

/**
 *
 * A Track that interpolates Strings
 *
 *
 * @author Ben Houston / http://clara.io/
 * @author David Sarno / http://lighthaus.us/
 * @author tschw
 */

THREE.StringKeyframeTrack = function ( name, times, values, interpolation ) {

	THREE.KeyframeTrack.call( this, name, times, values, interpolation );

};

THREE.StringKeyframeTrack.prototype =
		Object.assign( Object.create( THREE.KeyframeTrack.prototype ), {

	constructor: THREE.StringKeyframeTrack,

	ValueTypeName: 'string',
	ValueBufferType: Array,

	DefaultInterpolation: THREE.InterpolateDiscrete,

	InterpolantFactoryMethodLinear: undefined,

	InterpolantFactoryMethodSmooth: undefined

} );

// File:src/animation/tracks/VectorKeyframeTrack.js

/**
 *
 * A Track of vectored keyframe values.
 *
 *
 * @author Ben Houston / http://clara.io/
 * @author David Sarno / http://lighthaus.us/
 * @author tschw
 */

THREE.VectorKeyframeTrack = function ( name, times, values, interpolation ) {

	THREE.KeyframeTrack.call( this, name, times, values, interpolation );

};

THREE.VectorKeyframeTrack.prototype =
		Object.assign( Object.create( THREE.KeyframeTrack.prototype ), {

	constructor: THREE.VectorKeyframeTrack,

	ValueTypeName: 'vector'

	// ValueBufferType is inherited

	// DefaultInterpolation is inherited

} );

// File:src/audio/Audio.js

/**
 * @author mrdoob / http://mrdoob.com/
 * @author Reece Aaron Lecrivain / http://reecenotes.com/
 */

THREE.Audio = function ( listener ) {

	THREE.Object3D.call( this );

	this.type = 'Audio';

	this.context = listener.context;
	this.source = this.context.createBufferSource();
	this.source.onended = this.onEnded.bind( this );

	this.gain = this.context.createGain();
	this.gain.connect( listener.getInput() );

	this.autoplay = false;

	this.startTime = 0;
	this.playbackRate = 1;
	this.isPlaying = false;
	this.hasPlaybackControl = true;
	this.sourceType = 'empty';

	this.filters = [];

};

THREE.Audio.prototype = Object.assign( Object.create( THREE.Object3D.prototype ), {

	constructor: THREE.Audio,

	getOutput: function () {

		return this.gain;

	},

	setNodeSource: function ( audioNode ) {

		this.hasPlaybackControl = false;
		this.sourceType = 'audioNode';
		this.source = audioNode;
		this.connect();

		return this;

	},

	setBuffer: function ( audioBuffer ) {

		this.source.buffer = audioBuffer;
		this.sourceType = 'buffer';

		if ( this.autoplay ) this.play();

		return this;

	},

	play: function () {

		if ( this.isPlaying === true ) {

			console.warn( 'THREE.Audio: Audio is already playing.' );
			return;

		}

		if ( this.hasPlaybackControl === false ) {

			console.warn( 'THREE.Audio: this Audio has no playback control.' );
			return;

		}

		var source = this.context.createBufferSource();

		source.buffer = this.source.buffer;
		source.loop = this.source.loop;
		source.onended = this.source.onended;
		source.start( 0, this.startTime );
		source.playbackRate.value = this.playbackRate;

		this.isPlaying = true;

		this.source = source;

		return this.connect();

	},

	pause: function () {

		if ( this.hasPlaybackControl === false ) {

			console.warn( 'THREE.Audio: this Audio has no playback control.' );
			return;

		}

		this.source.stop();
		this.startTime = this.context.currentTime;

		return this;

	},

	stop: function () {

		if ( this.hasPlaybackControl === false ) {

			console.warn( 'THREE.Audio: this Audio has no playback control.' );
			return;

		}

		this.source.stop();
		this.startTime = 0;

		return this;

	},

	connect: function () {

		if ( this.filters.length > 0 ) {

			this.source.connect( this.filters[ 0 ] );

			for ( var i = 1, l = this.filters.length; i < l; i ++ ) {

				this.filters[ i - 1 ].connect( this.filters[ i ] );

			}

			this.filters[ this.filters.length - 1 ].connect( this.getOutput() );

		} else {

			this.source.connect( this.getOutput() );

		}

		return this;

	},

	disconnect: function () {

		if ( this.filters.length > 0 ) {

			this.source.disconnect( this.filters[ 0 ] );

			for ( var i = 1, l = this.filters.length; i < l; i ++ ) {

				this.filters[ i - 1 ].disconnect( this.filters[ i ] );

			}

			this.filters[ this.filters.length - 1 ].disconnect( this.getOutput() );

		} else {

			this.source.disconnect( this.getOutput() );

		}

		return this;

	},

	getFilters: function () {

		return this.filters;

	},

	setFilters: function ( value ) {

		if ( ! value ) value = [];

		if ( this.isPlaying === true ) {

			this.disconnect();
			this.filters = value;
			this.connect();

		} else {

			this.filters = value;

		}

		return this;

	},

	getFilter: function () {

		return this.getFilters()[ 0 ];

	},

	setFilter: function ( filter ) {

		return this.setFilters( filter ? [ filter ] : [] );

	},

	setPlaybackRate: function ( value ) {

		if ( this.hasPlaybackControl === false ) {

			console.warn( 'THREE.Audio: this Audio has no playback control.' );
			return;

		}

		this.playbackRate = value;

		if ( this.isPlaying === true ) {

			this.source.playbackRate.value = this.playbackRate;

		}

		return this;

	},

	getPlaybackRate: function () {

		return this.playbackRate;

	},

	onEnded: function () {

		this.isPlaying = false;

	},

	getLoop: function () {

		if ( this.hasPlaybackControl === false ) {

			console.warn( 'THREE.Audio: this Audio has no playback control.' );
			return false;

		}

		return this.source.loop;

	},

	setLoop: function ( value ) {

		if ( this.hasPlaybackControl === false ) {

			console.warn( 'THREE.Audio: this Audio has no playback control.' );
			return;

		}

		this.source.loop = value;

	},

	getVolume: function () {

		return this.gain.gain.value;

	},


	setVolume: function ( value ) {

		this.gain.gain.value = value;

		return this;

	}

} );

// File:src/audio/AudioAnalyser.js

/**
 * @author mrdoob / http://mrdoob.com/
 */

THREE.AudioAnalyser = function ( audio, fftSize ) {

	this.analyser = audio.context.createAnalyser();
	this.analyser.fftSize = fftSize !== undefined ? fftSize : 2048;

	this.data = new Uint8Array( this.analyser.frequencyBinCount );

	audio.getOutput().connect( this.analyser );

};

Object.assign( THREE.AudioAnalyser.prototype, {

	getFrequencyData: function () {

		this.analyser.getByteFrequencyData( this.data );

		return this.data;

	},

	getAverageFrequency: function () {

		var value = 0, data = this.getFrequencyData();

		for ( var i = 0; i < data.length; i ++ ) {

			value += data[ i ];

		}

		return value / data.length;

	}

} );

// File:src/audio/AudioContext.js

/**
 * @author mrdoob / http://mrdoob.com/
 */

Object.defineProperty( THREE, 'AudioContext', {

	get: ( function () {

		var context;

		return function () {

			if ( context === undefined ) {

				context = new ( window.AudioContext || window.webkitAudioContext )();

			}

			return context;

		};

	} )()

} );

// File:src/audio/PositionalAudio.js

/**
 * @author mrdoob / http://mrdoob.com/
 */

THREE.PositionalAudio = function ( listener ) {

	THREE.Audio.call( this, listener );

	this.panner = this.context.createPanner();
	this.panner.connect( this.gain );

};

THREE.PositionalAudio.prototype = Object.assign( Object.create( THREE.Audio.prototype ), {

	constructor: THREE.PositionalAudio,

	getOutput: function () {

		return this.panner;

	},

	getRefDistance: function () {

		return this.panner.refDistance;

	},

	setRefDistance: function ( value ) {

		this.panner.refDistance = value;

	},

	getRolloffFactor: function () {

		return this.panner.rolloffFactor;

	},

	setRolloffFactor: function ( value ) {

		this.panner.rolloffFactor = value;

	},

	getDistanceModel: function () {

		return this.panner.distanceModel;

	},

	setDistanceModel: function ( value ) {

		this.panner.distanceModel = value;

	},

	getMaxDistance: function () {

		return this.panner.maxDistance;

	},

	setMaxDistance: function ( value ) {

		this.panner.maxDistance = value;

	},

	updateMatrixWorld: ( function () {

		var position = new THREE.Vector3();

		return function updateMatrixWorld( force ) {

			THREE.Object3D.prototype.updateMatrixWorld.call( this, force );

			position.setFromMatrixPosition( this.matrixWorld );

			this.panner.setPosition( position.x, position.y, position.z );

		};

	} )()


} );

// File:src/audio/AudioListener.js

/**
 * @author mrdoob / http://mrdoob.com/
 */

THREE.AudioListener = function () {

	THREE.Object3D.call( this );

	this.type = 'AudioListener';

	this.context = THREE.AudioContext;

	this.gain = this.context.createGain();
	this.gain.connect( this.context.destination );

	this.filter = null;

};

THREE.AudioListener.prototype = Object.assign( Object.create( THREE.Object3D.prototype ), {

	constructor: THREE.AudioListener,

	getInput: function () {

		return this.gain;

	},

	removeFilter: function ( ) {

		if ( this.filter !== null ) {

			this.gain.disconnect( this.filter );
			this.filter.disconnect( this.context.destination );
			this.gain.connect( this.context.destination );
			this.filter = null;

		}

	},

	getFilter: function () {

		return this.filter;

	},

	setFilter: function ( value ) {

		if ( this.filter !== null ) {

			this.gain.disconnect( this.filter );
			this.filter.disconnect( this.context.destination );

		} else {

			this.gain.disconnect( this.context.destination );

		}

		this.filter = value;
		this.gain.connect( this.filter );
		this.filter.connect( this.context.destination );

	},

	getMasterVolume: function () {

		return this.gain.gain.value;

	},

	setMasterVolume: function ( value ) {

		this.gain.gain.value = value;

	},

	updateMatrixWorld: ( function () {

		var position = new THREE.Vector3();
		var quaternion = new THREE.Quaternion();
		var scale = new THREE.Vector3();

		var orientation = new THREE.Vector3();

		return function updateMatrixWorld( force ) {

			THREE.Object3D.prototype.updateMatrixWorld.call( this, force );

			var listener = this.context.listener;
			var up = this.up;

			this.matrixWorld.decompose( position, quaternion, scale );

			orientation.set( 0, 0, - 1 ).applyQuaternion( quaternion );

			listener.setPosition( position.x, position.y, position.z );
			listener.setOrientation( orientation.x, orientation.y, orientation.z, up.x, up.y, up.z );

		};

	} )()

} );

// File:src/cameras/Camera.js

/**
 * @author mrdoob / http://mrdoob.com/
 * @author mikael emtinger / http://gomo.se/
 * @author WestLangley / http://github.com/WestLangley
*/

THREE.Camera = function () {

	THREE.Object3D.call( this );

	this.type = 'Camera';

	this.matrixWorldInverse = new THREE.Matrix4();
	this.projectionMatrix = new THREE.Matrix4();

};

THREE.Camera.prototype = Object.create( THREE.Object3D.prototype );
THREE.Camera.prototype.constructor = THREE.Camera;

THREE.Camera.prototype.getWorldDirection = function () {

	var quaternion = new THREE.Quaternion();

	return function ( optionalTarget ) {

		var result = optionalTarget || new THREE.Vector3();

		this.getWorldQuaternion( quaternion );

		return result.set( 0, 0, - 1 ).applyQuaternion( quaternion );

	};

}();

THREE.Camera.prototype.lookAt = function () {

	// This routine does not support cameras with rotated and/or translated parent(s)

	var m1 = new THREE.Matrix4();

	return function ( vector ) {

		m1.lookAt( this.position, vector, this.up );

		this.quaternion.setFromRotationMatrix( m1 );

	};

}();

THREE.Camera.prototype.clone = function () {

	return new this.constructor().copy( this );

};

THREE.Camera.prototype.copy = function ( source ) {

	THREE.Object3D.prototype.copy.call( this, source );

	this.matrixWorldInverse.copy( source.matrixWorldInverse );
	this.projectionMatrix.copy( source.projectionMatrix );

	return this;

};

// File:src/cameras/CubeCamera.js

/**
 * Camera for rendering cube maps
 *	- renders scene into axis-aligned cube
 *
 * @author alteredq / http://alteredqualia.com/
 */

THREE.CubeCamera = function ( near, far, cubeResolution ) {

	THREE.Object3D.call( this );

	this.type = 'CubeCamera';

	var fov = 90, aspect = 1;

	var cameraPX = new THREE.PerspectiveCamera( fov, aspect, near, far );
	cameraPX.up.set( 0, - 1, 0 );
	cameraPX.lookAt( new THREE.Vector3( 1, 0, 0 ) );
	this.add( cameraPX );

	var cameraNX = new THREE.PerspectiveCamera( fov, aspect, near, far );
	cameraNX.up.set( 0, - 1, 0 );
	cameraNX.lookAt( new THREE.Vector3( - 1, 0, 0 ) );
	this.add( cameraNX );

	var cameraPY = new THREE.PerspectiveCamera( fov, aspect, near, far );
	cameraPY.up.set( 0, 0, 1 );
	cameraPY.lookAt( new THREE.Vector3( 0, 1, 0 ) );
	this.add( cameraPY );

	var cameraNY = new THREE.PerspectiveCamera( fov, aspect, near, far );
	cameraNY.up.set( 0, 0, - 1 );
	cameraNY.lookAt( new THREE.Vector3( 0, - 1, 0 ) );
	this.add( cameraNY );

	var cameraPZ = new THREE.PerspectiveCamera( fov, aspect, near, far );
	cameraPZ.up.set( 0, - 1, 0 );
	cameraPZ.lookAt( new THREE.Vector3( 0, 0, 1 ) );
	this.add( cameraPZ );

	var cameraNZ = new THREE.PerspectiveCamera( fov, aspect, near, far );
	cameraNZ.up.set( 0, - 1, 0 );
	cameraNZ.lookAt( new THREE.Vector3( 0, 0, - 1 ) );
	this.add( cameraNZ );

	var options = { format: THREE.RGBFormat, magFilter: THREE.LinearFilter, minFilter: THREE.LinearFilter };

	this.renderTarget = new THREE.WebGLRenderTargetCube( cubeResolution, cubeResolution, options );

	this.updateCubeMap = function ( renderer, scene ) {

		if ( this.parent === null ) this.updateMatrixWorld();

		var renderTarget = this.renderTarget;
		var generateMipmaps = renderTarget.texture.generateMipmaps;

		renderTarget.texture.generateMipmaps = false;

		renderTarget.activeCubeFace = 0;
		renderer.render( scene, cameraPX, renderTarget );

		renderTarget.activeCubeFace = 1;
		renderer.render( scene, cameraNX, renderTarget );

		renderTarget.activeCubeFace = 2;
		renderer.render( scene, cameraPY, renderTarget );

		renderTarget.activeCubeFace = 3;
		renderer.render( scene, cameraNY, renderTarget );

		renderTarget.activeCubeFace = 4;
		renderer.render( scene, cameraPZ, renderTarget );

		renderTarget.texture.generateMipmaps = generateMipmaps;

		renderTarget.activeCubeFace = 5;
		renderer.render( scene, cameraNZ, renderTarget );

		renderer.setRenderTarget( null );

	};

};

THREE.CubeCamera.prototype = Object.create( THREE.Object3D.prototype );
THREE.CubeCamera.prototype.constructor = THREE.CubeCamera;

// File:src/cameras/OrthographicCamera.js

/**
 * @author alteredq / http://alteredqualia.com/
 */

THREE.OrthographicCamera = function ( left, right, top, bottom, near, far ) {

	THREE.Camera.call( this );

	this.type = 'OrthographicCamera';

	this.zoom = 1;

	this.left = left;
	this.right = right;
	this.top = top;
	this.bottom = bottom;

	this.near = ( near !== undefined ) ? near : 0.1;
	this.far = ( far !== undefined ) ? far : 2000;

	this.updateProjectionMatrix();

};

THREE.OrthographicCamera.prototype = Object.assign( Object.create( THREE.Camera.prototype ), {

	constructor: THREE.OrthographicCamera,

	copy: function ( source ) {

		THREE.Camera.prototype.copy.call( this, source );

		this.left = source.left;
		this.right = source.right;
		this.top = source.top;
		this.bottom = source.bottom;
		this.near = source.near;
		this.far = source.far;

		this.zoom = source.zoom;

		return this;

	},

	updateProjectionMatrix: function () {

		var dx = ( this.right - this.left ) / ( 2 * this.zoom );
		var dy = ( this.top - this.bottom ) / ( 2 * this.zoom );
		var cx = ( this.right + this.left ) / 2;
		var cy = ( this.top + this.bottom ) / 2;

		this.projectionMatrix.makeOrthographic( cx - dx, cx + dx, cy + dy, cy - dy, this.near, this.far );

	},

	toJSON: function ( meta ) {

		var data = THREE.Object3D.prototype.toJSON.call( this, meta );

		data.object.zoom = this.zoom;
		data.object.left = this.left;
		data.object.right = this.right;
		data.object.top = this.top;
		data.object.bottom = this.bottom;
		data.object.near = this.near;
		data.object.far = this.far;

		return data;

	}

} );

// File:src/cameras/PerspectiveCamera.js

/**
 * @author mrdoob / http://mrdoob.com/
 * @author greggman / http://games.greggman.com/
 * @author zz85 / http://www.lab4games.net/zz85/blog
 * @author tschw
 */

THREE.PerspectiveCamera = function( fov, aspect, near, far ) {

	THREE.Camera.call( this );

	this.type = 'PerspectiveCamera';

	this.fov = fov !== undefined ? fov : 50;
	this.zoom = 1;

	this.near = near !== undefined ? near : 0.1;
	this.far = far !== undefined ? far : 2000;
	this.focus = 10;

	this.aspect = aspect !== undefined ? aspect : 1;
	this.view = null;

	this.filmGauge = 35;	// width of the film (default in millimeters)
	this.filmOffset = 0;	// horizontal film offset (same unit as gauge)

	this.updateProjectionMatrix();

};

THREE.PerspectiveCamera.prototype = Object.assign( Object.create( THREE.Camera.prototype ), {

	constructor: THREE.PerspectiveCamera,

	copy: function ( source ) {

		THREE.Camera.prototype.copy.call( this, source );

		this.fov = source.fov;
		this.zoom = source.zoom;

		this.near = source.near;
		this.far = source.far;
		this.focus = source.focus;

		this.aspect = source.aspect;
		this.view = source.view === null ? null : Object.assign( {}, source.view );

		this.filmGauge = source.filmGauge;
		this.filmOffset = source.filmOffset;

		return this;

	},

	/**
	 * Sets the FOV by focal length in respect to the current .filmGauge.
	 *
	 * The default film gauge is 35, so that the focal length can be specified for
	 * a 35mm (full frame) camera.
	 *
	 * Values for focal length and film gauge must have the same unit.
	 */
	setFocalLength: function ( focalLength ) {

		// see http://www.bobatkins.com/photography/technical/field_of_view.html
		var vExtentSlope = 0.5 * this.getFilmHeight() / focalLength;

		this.fov = THREE.Math.RAD2DEG * 2 * Math.atan( vExtentSlope );
		this.updateProjectionMatrix();

	},

	/**
	 * Calculates the focal length from the current .fov and .filmGauge.
	 */
	getFocalLength: function () {

		var vExtentSlope = Math.tan( THREE.Math.DEG2RAD * 0.5 * this.fov );

		return 0.5 * this.getFilmHeight() / vExtentSlope;

	},

	getEffectiveFOV: function () {

		return THREE.Math.RAD2DEG * 2 * Math.atan(
				Math.tan( THREE.Math.DEG2RAD * 0.5 * this.fov ) / this.zoom );

	},

	getFilmWidth: function () {

		// film not completely covered in portrait format (aspect < 1)
		return this.filmGauge * Math.min( this.aspect, 1 );

	},

	getFilmHeight: function () {

		// film not completely covered in landscape format (aspect > 1)
		return this.filmGauge / Math.max( this.aspect, 1 );

	},

	/**
	 * Sets an offset in a larger frustum. This is useful for multi-window or
	 * multi-monitor/multi-machine setups.
	 *
	 * For example, if you have 3x2 monitors and each monitor is 1920x1080 and
	 * the monitors are in grid like this
	 *
	 *   +---+---+---+
	 *   | A | B | C |
	 *   +---+---+---+
	 *   | D | E | F |
	 *   +---+---+---+
	 *
	 * then for each monitor you would call it like this
	 *
	 *   var w = 1920;
	 *   var h = 1080;
	 *   var fullWidth = w * 3;
	 *   var fullHeight = h * 2;
	 *
	 *   --A--
	 *   camera.setOffset( fullWidth, fullHeight, w * 0, h * 0, w, h );
	 *   --B--
	 *   camera.setOffset( fullWidth, fullHeight, w * 1, h * 0, w, h );
	 *   --C--
	 *   camera.setOffset( fullWidth, fullHeight, w * 2, h * 0, w, h );
	 *   --D--
	 *   camera.setOffset( fullWidth, fullHeight, w * 0, h * 1, w, h );
	 *   --E--
	 *   camera.setOffset( fullWidth, fullHeight, w * 1, h * 1, w, h );
	 *   --F--
	 *   camera.setOffset( fullWidth, fullHeight, w * 2, h * 1, w, h );
	 *
	 *   Note there is no reason monitors have to be the same size or in a grid.
	 */
	setViewOffset: function ( fullWidth, fullHeight, x, y, width, height ) {

		this.aspect = fullWidth / fullHeight;

		this.view = {
			fullWidth: fullWidth,
			fullHeight: fullHeight,
			offsetX: x,
			offsetY: y,
			width: width,
			height: height
		};

		this.updateProjectionMatrix();

	},

	clearViewOffset: function() {

		this.view = null;
		this.updateProjectionMatrix();

	},

	updateProjectionMatrix: function () {

		var near = this.near,
			top = near * Math.tan(
					THREE.Math.DEG2RAD * 0.5 * this.fov ) / this.zoom,
			height = 2 * top,
			width = this.aspect * height,
			left = - 0.5 * width,
			view = this.view;

		if ( view !== null ) {

			var fullWidth = view.fullWidth,
				fullHeight = view.fullHeight;

			left += view.offsetX * width / fullWidth;
			top -= view.offsetY * height / fullHeight;
			width *= view.width / fullWidth;
			height *= view.height / fullHeight;

		}

		var skew = this.filmOffset;
		if ( skew !== 0 ) left += near * skew / this.getFilmWidth();

		this.projectionMatrix.makeFrustum(
				left, left + width, top - height, top, near, this.far );

	},

	toJSON: function ( meta ) {

		var data = THREE.Object3D.prototype.toJSON.call( this, meta );

		data.object.fov = this.fov;
		data.object.zoom = this.zoom;

		data.object.near = this.near;
		data.object.far = this.far;
		data.object.focus = this.focus;

		data.object.aspect = this.aspect;

		if ( this.view !== null ) data.object.view = Object.assign( {}, this.view );

		data.object.filmGauge = this.filmGauge;
		data.object.filmOffset = this.filmOffset;

		return data;

	}

} );

// File:src/cameras/StereoCamera.js

/**
 * @author mrdoob / http://mrdoob.com/
 */

THREE.StereoCamera = function () {

	this.type = 'StereoCamera';

	this.aspect = 1;

	this.cameraL = new THREE.PerspectiveCamera();
	this.cameraL.layers.enable( 1 );
	this.cameraL.matrixAutoUpdate = false;

	this.cameraR = new THREE.PerspectiveCamera();
	this.cameraR.layers.enable( 2 );
	this.cameraR.matrixAutoUpdate = false;

};

Object.assign( THREE.StereoCamera.prototype, {

	update: ( function () {

		var focus, fov, aspect, near, far;

		var eyeRight = new THREE.Matrix4();
		var eyeLeft = new THREE.Matrix4();

		return function update ( camera ) {

			var needsUpdate = focus !== camera.focus || fov !== camera.fov ||
												aspect !== camera.aspect * this.aspect || near !== camera.near ||
												far !== camera.far;

			if ( needsUpdate ) {

				focus = camera.focus;
				fov = camera.fov;
				aspect = camera.aspect * this.aspect;
				near = camera.near;
				far = camera.far;

				// Off-axis stereoscopic effect based on
				// http://paulbourke.net/stereographics/stereorender/

				var projectionMatrix = camera.projectionMatrix.clone();
				var eyeSep = 0.064 / 2;
				var eyeSepOnProjection = eyeSep * near / focus;
				var ymax = near * Math.tan( THREE.Math.DEG2RAD * fov * 0.5 );
				var xmin, xmax;

				// translate xOffset

				eyeLeft.elements[ 12 ] = - eyeSep;
				eyeRight.elements[ 12 ] = eyeSep;

				// for left eye

				xmin = - ymax * aspect + eyeSepOnProjection;
				xmax = ymax * aspect + eyeSepOnProjection;

				projectionMatrix.elements[ 0 ] = 2 * near / ( xmax - xmin );
				projectionMatrix.elements[ 8 ] = ( xmax + xmin ) / ( xmax - xmin );

				this.cameraL.projectionMatrix.copy( projectionMatrix );

				// for right eye

				xmin = - ymax * aspect - eyeSepOnProjection;
				xmax = ymax * aspect - eyeSepOnProjection;

				projectionMatrix.elements[ 0 ] = 2 * near / ( xmax - xmin );
				projectionMatrix.elements[ 8 ] = ( xmax + xmin ) / ( xmax - xmin );

				this.cameraR.projectionMatrix.copy( projectionMatrix );

			}

			this.cameraL.matrixWorld.copy( camera.matrixWorld ).multiply( eyeLeft );
			this.cameraR.matrixWorld.copy( camera.matrixWorld ).multiply( eyeRight );

		};

	} )()

} );

// File:src/lights/Light.js

/**
 * @author mrdoob / http://mrdoob.com/
 * @author alteredq / http://alteredqualia.com/
 */

THREE.Light = function ( color, intensity ) {

	THREE.Object3D.call( this );

	this.type = 'Light';

	this.color = new THREE.Color( color );
	this.intensity = intensity !== undefined ? intensity : 1;

	this.receiveShadow = undefined;

};

THREE.Light.prototype = Object.assign( Object.create( THREE.Object3D.prototype ), {

	constructor: THREE.Light,

	copy: function ( source ) {

		THREE.Object3D.prototype.copy.call( this, source );

		this.color.copy( source.color );
		this.intensity = source.intensity;

		return this;

	},

	toJSON: function ( meta ) {

		var data = THREE.Object3D.prototype.toJSON.call( this, meta );

		data.object.color = this.color.getHex();
		data.object.intensity = this.intensity;

		if ( this.groundColor !== undefined ) data.object.groundColor = this.groundColor.getHex();

		if ( this.distance !== undefined ) data.object.distance = this.distance;
		if ( this.angle !== undefined ) data.object.angle = this.angle;
		if ( this.decay !== undefined ) data.object.decay = this.decay;
		if ( this.penumbra !== undefined ) data.object.penumbra = this.penumbra;

		return data;

	}

} );

// File:src/lights/LightShadow.js

/**
 * @author mrdoob / http://mrdoob.com/
 */

THREE.LightShadow = function ( camera ) {

	this.camera = camera;

	this.bias = 0;
	this.radius = 1;

	this.mapSize = new THREE.Vector2( 512, 512 );

	this.map = null;
	this.matrix = new THREE.Matrix4();

};

Object.assign( THREE.LightShadow.prototype, {

	copy: function ( source ) {

		this.camera = source.camera.clone();

		this.bias = source.bias;
		this.radius = source.radius;

		this.mapSize.copy( source.mapSize );

		return this;

	},

	clone: function () {

		return new this.constructor().copy( this );

	}

} );

// File:src/lights/AmbientLight.js

/**
 * @author mrdoob / http://mrdoob.com/
 */

THREE.AmbientLight = function ( color, intensity ) {

	THREE.Light.call( this, color, intensity );

	this.type = 'AmbientLight';

	this.castShadow = undefined;

};

THREE.AmbientLight.prototype = Object.assign( Object.create( THREE.Light.prototype ), {

	constructor: THREE.AmbientLight

} );

// File:src/lights/DirectionalLight.js

/**
 * @author mrdoob / http://mrdoob.com/
 * @author alteredq / http://alteredqualia.com/
 */

THREE.DirectionalLight = function ( color, intensity ) {

	THREE.Light.call( this, color, intensity );

	this.type = 'DirectionalLight';

	this.position.set( 0, 1, 0 );
	this.updateMatrix();

	this.target = new THREE.Object3D();

	this.shadow = new THREE.DirectionalLightShadow();

};

THREE.DirectionalLight.prototype = Object.assign( Object.create( THREE.Light.prototype ), {

	constructor: THREE.DirectionalLight,

	copy: function ( source ) {

		THREE.Light.prototype.copy.call( this, source );

		this.target = source.target.clone();

		this.shadow = source.shadow.clone();

		return this;

	}

} );

// File:src/lights/DirectionalLightShadow.js

/**
 * @author mrdoob / http://mrdoob.com/
 */

THREE.DirectionalLightShadow = function ( light ) {

	THREE.LightShadow.call( this, new THREE.OrthographicCamera( - 5, 5, 5, - 5, 0.5, 500 ) );

};

THREE.DirectionalLightShadow.prototype = Object.assign( Object.create( THREE.LightShadow.prototype ), {

	constructor: THREE.DirectionalLightShadow

} );

// File:src/lights/HemisphereLight.js

/**
 * @author alteredq / http://alteredqualia.com/
 */

THREE.HemisphereLight = function ( skyColor, groundColor, intensity ) {

	THREE.Light.call( this, skyColor, intensity );

	this.type = 'HemisphereLight';

	this.castShadow = undefined;

	this.position.set( 0, 1, 0 );
	this.updateMatrix();

	this.groundColor = new THREE.Color( groundColor );

};

THREE.HemisphereLight.prototype = Object.assign( Object.create( THREE.Light.prototype ), {

	constructor: THREE.HemisphereLight,

	copy: function ( source ) {

		THREE.Light.prototype.copy.call( this, source );

		this.groundColor.copy( source.groundColor );

		return this;

	}

} );

// File:src/lights/PointLight.js

/**
 * @author mrdoob / http://mrdoob.com/
 */


THREE.PointLight = function ( color, intensity, distance, decay ) {

	THREE.Light.call( this, color, intensity );

	this.type = 'PointLight';

	Object.defineProperty( this, 'power', {
		get: function () {
			// intensity = power per solid angle.
			// ref: equation (15) from http://www.frostbite.com/wp-content/uploads/2014/11/course_notes_moving_frostbite_to_pbr.pdf
			return this.intensity * 4 * Math.PI;

		},
		set: function ( power ) {
			// intensity = power per solid angle.
			// ref: equation (15) from http://www.frostbite.com/wp-content/uploads/2014/11/course_notes_moving_frostbite_to_pbr.pdf
			this.intensity = power / ( 4 * Math.PI );
		}
	} );

	this.distance = ( distance !== undefined ) ? distance : 0;
	this.decay = ( decay !== undefined ) ? decay : 1;	// for physically correct lights, should be 2.

	this.shadow = new THREE.LightShadow( new THREE.PerspectiveCamera( 90, 1, 0.5, 500 ) );

};

THREE.PointLight.prototype = Object.assign( Object.create( THREE.Light.prototype ), {

	constructor: THREE.PointLight,

	copy: function ( source ) {

		THREE.Light.prototype.copy.call( this, source );

		this.distance = source.distance;
		this.decay = source.decay;

		this.shadow = source.shadow.clone();

		return this;

	}

} );

// File:src/lights/SpotLight.js

/**
 * @author alteredq / http://alteredqualia.com/
 */

THREE.SpotLight = function ( color, intensity, distance, angle, penumbra, decay ) {

	THREE.Light.call( this, color, intensity );

	this.type = 'SpotLight';

	this.position.set( 0, 1, 0 );
	this.updateMatrix();

	this.target = new THREE.Object3D();

	Object.defineProperty( this, 'power', {
		get: function () {
			// intensity = power per solid angle.
			// ref: equation (17) from http://www.frostbite.com/wp-content/uploads/2014/11/course_notes_moving_frostbite_to_pbr.pdf
			return this.intensity * Math.PI;
		},
		set: function ( power ) {
			// intensity = power per solid angle.
			// ref: equation (17) from http://www.frostbite.com/wp-content/uploads/2014/11/course_notes_moving_frostbite_to_pbr.pdf
			this.intensity = power / Math.PI;
		}
	} );

	this.distance = ( distance !== undefined ) ? distance : 0;
	this.angle = ( angle !== undefined ) ? angle : Math.PI / 3;
	this.penumbra = ( penumbra !== undefined ) ? penumbra : 0;
	this.decay = ( decay !== undefined ) ? decay : 1;	// for physically correct lights, should be 2.

	this.shadow = new THREE.SpotLightShadow();

};

THREE.SpotLight.prototype = Object.assign( Object.create( THREE.Light.prototype ), {

	constructor: THREE.SpotLight,

	copy: function ( source ) {

		THREE.Light.prototype.copy.call( this, source );

		this.distance = source.distance;
		this.angle = source.angle;
		this.penumbra = source.penumbra;
		this.decay = source.decay;

		this.target = source.target.clone();

		this.shadow = source.shadow.clone();

		return this;

	}

} );

// File:src/lights/SpotLightShadow.js

/**
 * @author mrdoob / http://mrdoob.com/
 */

THREE.SpotLightShadow = function () {

	THREE.LightShadow.call( this, new THREE.PerspectiveCamera( 50, 1, 0.5, 500 ) );

};

THREE.SpotLightShadow.prototype = Object.assign( Object.create( THREE.LightShadow.prototype ), {

	constructor: THREE.SpotLightShadow,

	update: function ( light ) {

		var fov = THREE.Math.RAD2DEG * 2 * light.angle;
		var aspect = this.mapSize.width / this.mapSize.height;
		var far = light.distance || 500;

		var camera = this.camera;

		if ( fov !== camera.fov || aspect !== camera.aspect || far !== camera.far ) {

			camera.fov = fov;
			camera.aspect = aspect;
			camera.far = far;
			camera.updateProjectionMatrix();

		}

	}

} );

// File:src/loaders/AudioLoader.js

/**
 * @author Reece Aaron Lecrivain / http://reecenotes.com/
 */

THREE.AudioLoader = function ( manager ) {

	this.manager = ( manager !== undefined ) ? manager : THREE.DefaultLoadingManager;

};

THREE.AudioLoader.prototype = {

	constructor: THREE.AudioLoader,

	load: function ( url, onLoad, onProgress, onError ) {

		var loader = new THREE.XHRLoader( this.manager );
		loader.setResponseType( 'arraybuffer' );
		loader.load( url, function ( buffer ) {

			var context = THREE.AudioContext;

			context.decodeAudioData( buffer, function ( audioBuffer ) {

				onLoad( audioBuffer );

			} );

		}, onProgress, onError );

	}

};

// File:src/loaders/Cache.js

/**
 * @author mrdoob / http://mrdoob.com/
 */

THREE.Cache = {

	enabled: false,

	files: {},

	add: function ( key, file ) {

		if ( this.enabled === false ) return;

		// console.log( 'THREE.Cache', 'Adding key:', key );

		this.files[ key ] = file;

	},

	get: function ( key ) {

		if ( this.enabled === false ) return;

		// console.log( 'THREE.Cache', 'Checking key:', key );

		return this.files[ key ];

	},

	remove: function ( key ) {

		delete this.files[ key ];

	},

	clear: function () {

		this.files = {};

	}

};

// File:src/loaders/Loader.js

/**
 * @author alteredq / http://alteredqualia.com/
 */

THREE.Loader = function () {

	this.onLoadStart = function () {};
	this.onLoadProgress = function () {};
	this.onLoadComplete = function () {};

};

THREE.Loader.prototype = {

	constructor: THREE.Loader,

	crossOrigin: undefined,

	extractUrlBase: function ( url ) {

		var parts = url.split( '/' );

		if ( parts.length === 1 ) return './';

		parts.pop();

		return parts.join( '/' ) + '/';

	},

	initMaterials: function ( materials, texturePath, crossOrigin ) {

		var array = [];

		for ( var i = 0; i < materials.length; ++ i ) {

			array[ i ] = this.createMaterial( materials[ i ], texturePath, crossOrigin );

		}

		return array;

	},

	createMaterial: ( function () {

		var color, textureLoader, materialLoader;

		return function ( m, texturePath, crossOrigin ) {

			if ( color === undefined ) color = new THREE.Color();
			if ( textureLoader === undefined ) textureLoader = new THREE.TextureLoader();
			if ( materialLoader === undefined ) materialLoader = new THREE.MaterialLoader();

			// convert from old material format

			var textures = {};

			function loadTexture( path, repeat, offset, wrap, anisotropy ) {

				var fullPath = texturePath + path;
				var loader = THREE.Loader.Handlers.get( fullPath );

				var texture;

				if ( loader !== null ) {

					texture = loader.load( fullPath );

				} else {

					textureLoader.setCrossOrigin( crossOrigin );
					texture = textureLoader.load( fullPath );

				}

				if ( repeat !== undefined ) {

					texture.repeat.fromArray( repeat );

					if ( repeat[ 0 ] !== 1 ) texture.wrapS = THREE.RepeatWrapping;
					if ( repeat[ 1 ] !== 1 ) texture.wrapT = THREE.RepeatWrapping;

				}

				if ( offset !== undefined ) {

					texture.offset.fromArray( offset );

				}

				if ( wrap !== undefined ) {

					if ( wrap[ 0 ] === 'repeat' ) texture.wrapS = THREE.RepeatWrapping;
					if ( wrap[ 0 ] === 'mirror' ) texture.wrapS = THREE.MirroredRepeatWrapping;

					if ( wrap[ 1 ] === 'repeat' ) texture.wrapT = THREE.RepeatWrapping;
					if ( wrap[ 1 ] === 'mirror' ) texture.wrapT = THREE.MirroredRepeatWrapping;

				}

				if ( anisotropy !== undefined ) {

					texture.anisotropy = anisotropy;

				}

				var uuid = THREE.Math.generateUUID();

				textures[ uuid ] = texture;

				return uuid;

			}

			//

			var json = {
				uuid: THREE.Math.generateUUID(),
				type: 'MeshLambertMaterial'
			};

			for ( var name in m ) {

				var value = m[ name ];

				switch ( name ) {
					case 'DbgColor':
					case 'DbgIndex':
					case 'opticalDensity':
					case 'illumination':
						break;
					case 'DbgName':
						json.name = value;
						break;
					case 'blending':
						json.blending = THREE[ value ];
						break;
					case 'colorAmbient':
					case 'mapAmbient':
						console.warn( 'THREE.Loader.createMaterial:', name, 'is no longer supported.' );
						break;
					case 'colorDiffuse':
						json.color = color.fromArray( value ).getHex();
						break;
					case 'colorSpecular':
						json.specular = color.fromArray( value ).getHex();
						break;
					case 'colorEmissive':
						json.emissive = color.fromArray( value ).getHex();
						break;
					case 'specularCoef':
						json.shininess = value;
						break;
					case 'shading':
						if ( value.toLowerCase() === 'basic' ) json.type = 'MeshBasicMaterial';
						if ( value.toLowerCase() === 'phong' ) json.type = 'MeshPhongMaterial';
						break;
					case 'mapDiffuse':
						json.map = loadTexture( value, m.mapDiffuseRepeat, m.mapDiffuseOffset, m.mapDiffuseWrap, m.mapDiffuseAnisotropy );
						break;
					case 'mapDiffuseRepeat':
					case 'mapDiffuseOffset':
					case 'mapDiffuseWrap':
					case 'mapDiffuseAnisotropy':
						break;
					case 'mapLight':
						json.lightMap = loadTexture( value, m.mapLightRepeat, m.mapLightOffset, m.mapLightWrap, m.mapLightAnisotropy );
						break;
					case 'mapLightRepeat':
					case 'mapLightOffset':
					case 'mapLightWrap':
					case 'mapLightAnisotropy':
						break;
					case 'mapAO':
						json.aoMap = loadTexture( value, m.mapAORepeat, m.mapAOOffset, m.mapAOWrap, m.mapAOAnisotropy );
						break;
					case 'mapAORepeat':
					case 'mapAOOffset':
					case 'mapAOWrap':
					case 'mapAOAnisotropy':
						break;
					case 'mapBump':
						json.bumpMap = loadTexture( value, m.mapBumpRepeat, m.mapBumpOffset, m.mapBumpWrap, m.mapBumpAnisotropy );
						break;
					case 'mapBumpScale':
						json.bumpScale = value;
						break;
					case 'mapBumpRepeat':
					case 'mapBumpOffset':
					case 'mapBumpWrap':
					case 'mapBumpAnisotropy':
						break;
					case 'mapNormal':
						json.normalMap = loadTexture( value, m.mapNormalRepeat, m.mapNormalOffset, m.mapNormalWrap, m.mapNormalAnisotropy );
						break;
					case 'mapNormalFactor':
						json.normalScale = [ value, value ];
						break;
					case 'mapNormalRepeat':
					case 'mapNormalOffset':
					case 'mapNormalWrap':
					case 'mapNormalAnisotropy':
						break;
					case 'mapSpecular':
						json.specularMap = loadTexture( value, m.mapSpecularRepeat, m.mapSpecularOffset, m.mapSpecularWrap, m.mapSpecularAnisotropy );
						break;
					case 'mapSpecularRepeat':
					case 'mapSpecularOffset':
					case 'mapSpecularWrap':
					case 'mapSpecularAnisotropy':
						break;
					case 'mapAlpha':
						json.alphaMap = loadTexture( value, m.mapAlphaRepeat, m.mapAlphaOffset, m.mapAlphaWrap, m.mapAlphaAnisotropy );
						break;
					case 'mapAlphaRepeat':
					case 'mapAlphaOffset':
					case 'mapAlphaWrap':
					case 'mapAlphaAnisotropy':
						break;
					case 'flipSided':
						json.side = THREE.BackSide;
						break;
					case 'doubleSided':
						json.side = THREE.DoubleSide;
						break;
					case 'transparency':
						console.warn( 'THREE.Loader.createMaterial: transparency has been renamed to opacity' );
						json.opacity = value;
						break;
					case 'depthTest':
					case 'depthWrite':
					case 'colorWrite':
					case 'opacity':
					case 'reflectivity':
					case 'transparent':
					case 'visible':
					case 'wireframe':
						json[ name ] = value;
						break;
					case 'vertexColors':
						if ( value === true ) json.vertexColors = THREE.VertexColors;
						if ( value === 'face' ) json.vertexColors = THREE.FaceColors;
						break;
					default:
						console.error( 'THREE.Loader.createMaterial: Unsupported', name, value );
						break;
				}

			}

			if ( json.type === 'MeshBasicMaterial' ) delete json.emissive;
			if ( json.type !== 'MeshPhongMaterial' ) delete json.specular;

			if ( json.opacity < 1 ) json.transparent = true;

			materialLoader.setTextures( textures );

			return materialLoader.parse( json );

		};

	} )()

};

THREE.Loader.Handlers = {

	handlers: [],

	add: function ( regex, loader ) {

		this.handlers.push( regex, loader );

	},

	get: function ( file ) {

		var handlers = this.handlers;

		for ( var i = 0, l = handlers.length; i < l; i += 2 ) {

			var regex = handlers[ i ];
			var loader  = handlers[ i + 1 ];

			if ( regex.test( file ) ) {

				return loader;

			}

		}

		return null;

	}

};

// File:src/loaders/XHRLoader.js

/**
 * @author mrdoob / http://mrdoob.com/
 */

THREE.XHRLoader = function ( manager ) {

	this.manager = ( manager !== undefined ) ? manager : THREE.DefaultLoadingManager;

};

THREE.XHRLoader.prototype = {

	constructor: THREE.XHRLoader,

	load: function ( url, onLoad, onProgress, onError ) {

		if ( this.path !== undefined ) url = this.path + url;

		var scope = this;

		var cached = THREE.Cache.get( url );

		if ( cached !== undefined ) {

			if ( onLoad ) {

				setTimeout( function () {

					onLoad( cached );

				}, 0 );

			}

			return cached;

		}

		var request = new XMLHttpRequest();
		request.overrideMimeType( 'text/plain' );
		request.open( 'GET', url, true );

		request.addEventListener( 'load', function ( event ) {

			var response = event.target.response;

			THREE.Cache.add( url, response );

			if ( this.status === 200 ) {

				if ( onLoad ) onLoad( response );

				scope.manager.itemEnd( url );

			} else if ( this.status === 0 ) {

				// Some browsers return HTTP Status 0 when using non-http protocol
				// e.g. 'file://' or 'data://'. Handle as success.

				console.warn( 'THREE.XHRLoader: HTTP Status 0 received.' );

				if ( onLoad ) onLoad( response );

				scope.manager.itemEnd( url );

			} else {

				if ( onError ) onError( event );

				scope.manager.itemError( url );

			}

		}, false );

		if ( onProgress !== undefined ) {

			request.addEventListener( 'progress', function ( event ) {

				onProgress( event );

			}, false );

		}

		request.addEventListener( 'error', function ( event ) {

			if ( onError ) onError( event );

			scope.manager.itemError( url );

		}, false );

		if ( this.responseType !== undefined ) request.responseType = this.responseType;
		if ( this.withCredentials !== undefined ) request.withCredentials = this.withCredentials;

		request.send( null );

		scope.manager.itemStart( url );

		return request;

	},

	setPath: function ( value ) {

		this.path = value;

	},

	setResponseType: function ( value ) {

		this.responseType = value;

	},

	setWithCredentials: function ( value ) {

		this.withCredentials = value;

	}

};

// File:src/loaders/FontLoader.js

/**
 * @author mrdoob / http://mrdoob.com/
 */

THREE.FontLoader = function ( manager ) {

	this.manager = ( manager !== undefined ) ? manager : THREE.DefaultLoadingManager;

};

THREE.FontLoader.prototype = {

	constructor: THREE.FontLoader,

	load: function ( url, onLoad, onProgress, onError ) {

		var scope = this;

		var loader = new THREE.XHRLoader( this.manager );
		loader.load( url, function ( text ) {

			var json;

			try {

				json = JSON.parse( text );

			} catch ( e ) {

				console.warn( 'THREE.FontLoader: typeface.js support is being deprecated. Use typeface.json instead.' );
				json = JSON.parse( text.substring( 65, text.length - 2 ) );

			}

			var font = scope.parse( json );

			if ( onLoad ) onLoad( font );

		}, onProgress, onError );

	},

	parse: function ( json ) {

		return new THREE.Font( json );

	}

};

// File:src/loaders/ImageLoader.js

/**
 * @author mrdoob / http://mrdoob.com/
 */

THREE.ImageLoader = function ( manager ) {

	this.manager = ( manager !== undefined ) ? manager : THREE.DefaultLoadingManager;

};

THREE.ImageLoader.prototype = {

	constructor: THREE.ImageLoader,

	load: function ( url, onLoad, onProgress, onError ) {

		if ( this.path !== undefined ) url = this.path + url;

		var scope = this;

		var cached = THREE.Cache.get( url );

		if ( cached !== undefined ) {

			scope.manager.itemStart( url );

			if ( onLoad ) {

				setTimeout( function () {

					onLoad( cached );

					scope.manager.itemEnd( url );

				}, 0 );

			} else {

				scope.manager.itemEnd( url );

			}

			return cached;

		}

		var image = document.createElement( 'img' );

		image.addEventListener( 'load', function ( event ) {

			THREE.Cache.add( url, this );

			if ( onLoad ) onLoad( this );

			scope.manager.itemEnd( url );

		}, false );

		if ( onProgress !== undefined ) {

			image.addEventListener( 'progress', function ( event ) {

				onProgress( event );

			}, false );

		}

		image.addEventListener( 'error', function ( event ) {

			if ( onError ) onError( event );

			scope.manager.itemError( url );

		}, false );

		if ( this.crossOrigin !== undefined ) image.crossOrigin = this.crossOrigin;

		scope.manager.itemStart( url );

		image.src = url;

		return image;

	},

	setCrossOrigin: function ( value ) {

		this.crossOrigin = value;

	},

	setPath: function ( value ) {

		this.path = value;

	}

};

// File:src/loaders/JSONLoader.js

/**
 * @author mrdoob / http://mrdoob.com/
 * @author alteredq / http://alteredqualia.com/
 */

THREE.JSONLoader = function ( manager ) {

	if ( typeof manager === 'boolean' ) {

		console.warn( 'THREE.JSONLoader: showStatus parameter has been removed from constructor.' );
		manager = undefined;

	}

	this.manager = ( manager !== undefined ) ? manager : THREE.DefaultLoadingManager;

	this.withCredentials = false;

};

THREE.JSONLoader.prototype = {

	constructor: THREE.JSONLoader,

	// Deprecated

	get statusDomElement () {

		if ( this._statusDomElement === undefined ) {

			this._statusDomElement = document.createElement( 'div' );

		}

		console.warn( 'THREE.JSONLoader: .statusDomElement has been removed.' );
		return this._statusDomElement;

	},

	load: function( url, onLoad, onProgress, onError ) {

		var scope = this;

		var texturePath = this.texturePath && ( typeof this.texturePath === "string" ) ? this.texturePath : THREE.Loader.prototype.extractUrlBase( url );

		var loader = new THREE.XHRLoader( this.manager );
		loader.setWithCredentials( this.withCredentials );
		loader.load( url, function ( text ) {

			var json = JSON.parse( text );
			var metadata = json.metadata;

			if ( metadata !== undefined ) {

				var type = metadata.type;

				if ( type !== undefined ) {

					if ( type.toLowerCase() === 'object' ) {

						console.error( 'THREE.JSONLoader: ' + url + ' should be loaded with THREE.ObjectLoader instead.' );
						return;

					}

					if ( type.toLowerCase() === 'scene' ) {

						console.error( 'THREE.JSONLoader: ' + url + ' should be loaded with THREE.SceneLoader instead.' );
						return;

					}

				}

			}

			var object = scope.parse( json, texturePath );
			onLoad( object.geometry, object.materials );

		}, onProgress, onError );

	},

	setTexturePath: function ( value ) {

		this.texturePath = value;

	},

	parse: function ( json, texturePath ) {

		var geometry = new THREE.Geometry(),
		scale = ( json.scale !== undefined ) ? 1.0 / json.scale : 1.0;

		parseModel( scale );

		parseSkin();
		parseMorphing( scale );
		parseAnimations();

		geometry.computeFaceNormals();
		geometry.computeBoundingSphere();

		function parseModel( scale ) {

			function isBitSet( value, position ) {

				return value & ( 1 << position );

			}

			var i, j, fi,

			offset, zLength,

		colorIndex, normalIndex, uvIndex, materialIndex,

			type,
			isQuad,
			hasMaterial,
			hasFaceVertexUv,
			hasFaceNormal, hasFaceVertexNormal,
			hasFaceColor, hasFaceVertexColor,

		vertex, face, faceA, faceB, hex, normal,

			uvLayer, uv, u, v,

			faces = json.faces,
			vertices = json.vertices,
			normals = json.normals,
			colors = json.colors,

			nUvLayers = 0;

			if ( json.uvs !== undefined ) {

				// disregard empty arrays

				for ( i = 0; i < json.uvs.length; i ++ ) {

					if ( json.uvs[ i ].length ) nUvLayers ++;

				}

				for ( i = 0; i < nUvLayers; i ++ ) {

					geometry.faceVertexUvs[ i ] = [];

				}

			}

			offset = 0;
			zLength = vertices.length;

			while ( offset < zLength ) {

				vertex = new THREE.Vector3();

				vertex.x = vertices[ offset ++ ] * scale;
				vertex.y = vertices[ offset ++ ] * scale;
				vertex.z = vertices[ offset ++ ] * scale;

				geometry.vertices.push( vertex );

			}

			offset = 0;
			zLength = faces.length;

			while ( offset < zLength ) {

				type = faces[ offset ++ ];


				isQuad              = isBitSet( type, 0 );
				hasMaterial         = isBitSet( type, 1 );
				hasFaceVertexUv     = isBitSet( type, 3 );
				hasFaceNormal       = isBitSet( type, 4 );
				hasFaceVertexNormal = isBitSet( type, 5 );
				hasFaceColor	     = isBitSet( type, 6 );
				hasFaceVertexColor  = isBitSet( type, 7 );

				// console.log("type", type, "bits", isQuad, hasMaterial, hasFaceVertexUv, hasFaceNormal, hasFaceVertexNormal, hasFaceColor, hasFaceVertexColor);

				if ( isQuad ) {

					faceA = new THREE.Face3();
					faceA.a = faces[ offset ];
					faceA.b = faces[ offset + 1 ];
					faceA.c = faces[ offset + 3 ];

					faceB = new THREE.Face3();
					faceB.a = faces[ offset + 1 ];
					faceB.b = faces[ offset + 2 ];
					faceB.c = faces[ offset + 3 ];

					offset += 4;

					if ( hasMaterial ) {

						materialIndex = faces[ offset ++ ];
						faceA.materialIndex = materialIndex;
						faceB.materialIndex = materialIndex;

					}

					// to get face <=> uv index correspondence

					fi = geometry.faces.length;

					if ( hasFaceVertexUv ) {

						for ( i = 0; i < nUvLayers; i ++ ) {

							uvLayer = json.uvs[ i ];

							geometry.faceVertexUvs[ i ][ fi ] = [];
							geometry.faceVertexUvs[ i ][ fi + 1 ] = [];

							for ( j = 0; j < 4; j ++ ) {

								uvIndex = faces[ offset ++ ];

								u = uvLayer[ uvIndex * 2 ];
								v = uvLayer[ uvIndex * 2 + 1 ];

								uv = new THREE.Vector2( u, v );

								if ( j !== 2 ) geometry.faceVertexUvs[ i ][ fi ].push( uv );
								if ( j !== 0 ) geometry.faceVertexUvs[ i ][ fi + 1 ].push( uv );

							}

						}

					}

					if ( hasFaceNormal ) {

						normalIndex = faces[ offset ++ ] * 3;

						faceA.normal.set(
							normals[ normalIndex ++ ],
							normals[ normalIndex ++ ],
							normals[ normalIndex ]
						);

						faceB.normal.copy( faceA.normal );

					}

					if ( hasFaceVertexNormal ) {

						for ( i = 0; i < 4; i ++ ) {

							normalIndex = faces[ offset ++ ] * 3;

							normal = new THREE.Vector3(
								normals[ normalIndex ++ ],
								normals[ normalIndex ++ ],
								normals[ normalIndex ]
							);


							if ( i !== 2 ) faceA.vertexNormals.push( normal );
							if ( i !== 0 ) faceB.vertexNormals.push( normal );

						}

					}


					if ( hasFaceColor ) {

						colorIndex = faces[ offset ++ ];
						hex = colors[ colorIndex ];

						faceA.color.setHex( hex );
						faceB.color.setHex( hex );

					}


					if ( hasFaceVertexColor ) {

						for ( i = 0; i < 4; i ++ ) {

							colorIndex = faces[ offset ++ ];
							hex = colors[ colorIndex ];

							if ( i !== 2 ) faceA.vertexColors.push( new THREE.Color( hex ) );
							if ( i !== 0 ) faceB.vertexColors.push( new THREE.Color( hex ) );

						}

					}

					geometry.faces.push( faceA );
					geometry.faces.push( faceB );

				} else {

					face = new THREE.Face3();
					face.a = faces[ offset ++ ];
					face.b = faces[ offset ++ ];
					face.c = faces[ offset ++ ];

					if ( hasMaterial ) {

						materialIndex = faces[ offset ++ ];
						face.materialIndex = materialIndex;

					}

					// to get face <=> uv index correspondence

					fi = geometry.faces.length;

					if ( hasFaceVertexUv ) {

						for ( i = 0; i < nUvLayers; i ++ ) {

							uvLayer = json.uvs[ i ];

							geometry.faceVertexUvs[ i ][ fi ] = [];

							for ( j = 0; j < 3; j ++ ) {

								uvIndex = faces[ offset ++ ];

								u = uvLayer[ uvIndex * 2 ];
								v = uvLayer[ uvIndex * 2 + 1 ];

								uv = new THREE.Vector2( u, v );

								geometry.faceVertexUvs[ i ][ fi ].push( uv );

							}

						}

					}

					if ( hasFaceNormal ) {

						normalIndex = faces[ offset ++ ] * 3;

						face.normal.set(
							normals[ normalIndex ++ ],
							normals[ normalIndex ++ ],
							normals[ normalIndex ]
						);

					}

					if ( hasFaceVertexNormal ) {

						for ( i = 0; i < 3; i ++ ) {

							normalIndex = faces[ offset ++ ] * 3;

							normal = new THREE.Vector3(
								normals[ normalIndex ++ ],
								normals[ normalIndex ++ ],
								normals[ normalIndex ]
							);

							face.vertexNormals.push( normal );

						}

					}


					if ( hasFaceColor ) {

						colorIndex = faces[ offset ++ ];
						face.color.setHex( colors[ colorIndex ] );

					}


					if ( hasFaceVertexColor ) {

						for ( i = 0; i < 3; i ++ ) {

							colorIndex = faces[ offset ++ ];
							face.vertexColors.push( new THREE.Color( colors[ colorIndex ] ) );

						}

					}

					geometry.faces.push( face );

				}

			}

		}

		function parseSkin() {

			var influencesPerVertex = ( json.influencesPerVertex !== undefined ) ? json.influencesPerVertex : 2;

			if ( json.skinWeights ) {

				for ( var i = 0, l = json.skinWeights.length; i < l; i += influencesPerVertex ) {

					var x =                               json.skinWeights[ i ];
					var y = ( influencesPerVertex > 1 ) ? json.skinWeights[ i + 1 ] : 0;
					var z = ( influencesPerVertex > 2 ) ? json.skinWeights[ i + 2 ] : 0;
					var w = ( influencesPerVertex > 3 ) ? json.skinWeights[ i + 3 ] : 0;

					geometry.skinWeights.push( new THREE.Vector4( x, y, z, w ) );

				}

			}

			if ( json.skinIndices ) {

				for ( var i = 0, l = json.skinIndices.length; i < l; i += influencesPerVertex ) {

					var a =                               json.skinIndices[ i ];
					var b = ( influencesPerVertex > 1 ) ? json.skinIndices[ i + 1 ] : 0;
					var c = ( influencesPerVertex > 2 ) ? json.skinIndices[ i + 2 ] : 0;
					var d = ( influencesPerVertex > 3 ) ? json.skinIndices[ i + 3 ] : 0;

					geometry.skinIndices.push( new THREE.Vector4( a, b, c, d ) );

				}

			}

			geometry.bones = json.bones;

			if ( geometry.bones && geometry.bones.length > 0 && ( geometry.skinWeights.length !== geometry.skinIndices.length || geometry.skinIndices.length !== geometry.vertices.length ) ) {

				console.warn( 'When skinning, number of vertices (' + geometry.vertices.length + '), skinIndices (' +
					geometry.skinIndices.length + '), and skinWeights (' + geometry.skinWeights.length + ') should match.' );

			}

		}

		function parseMorphing( scale ) {

			if ( json.morphTargets !== undefined ) {

				for ( var i = 0, l = json.morphTargets.length; i < l; i ++ ) {

					geometry.morphTargets[ i ] = {};
					geometry.morphTargets[ i ].name = json.morphTargets[ i ].name;
					geometry.morphTargets[ i ].vertices = [];

					var dstVertices = geometry.morphTargets[ i ].vertices;
					var srcVertices = json.morphTargets[ i ].vertices;

					for ( var v = 0, vl = srcVertices.length; v < vl; v += 3 ) {

						var vertex = new THREE.Vector3();
						vertex.x = srcVertices[ v ] * scale;
						vertex.y = srcVertices[ v + 1 ] * scale;
						vertex.z = srcVertices[ v + 2 ] * scale;

						dstVertices.push( vertex );

					}

				}

			}

			if ( json.morphColors !== undefined && json.morphColors.length > 0 ) {

				console.warn( 'THREE.JSONLoader: "morphColors" no longer supported. Using them as face colors.' );

				var faces = geometry.faces;
				var morphColors = json.morphColors[ 0 ].colors;

				for ( var i = 0, l = faces.length; i < l; i ++ ) {

					faces[ i ].color.fromArray( morphColors, i * 3 );

				}

			}

		}

		function parseAnimations() {

			var outputAnimations = [];

			// parse old style Bone/Hierarchy animations
			var animations = [];

			if ( json.animation !== undefined ) {

				animations.push( json.animation );

			}

			if ( json.animations !== undefined ) {

				if ( json.animations.length ) {

					animations = animations.concat( json.animations );

				} else {

					animations.push( json.animations );

				}

			}

			for ( var i = 0; i < animations.length; i ++ ) {

				var clip = THREE.AnimationClip.parseAnimation( animations[ i ], geometry.bones );
				if ( clip ) outputAnimations.push( clip );

			}

			// parse implicit morph animations
			if ( geometry.morphTargets ) {

				// TODO: Figure out what an appropraite FPS is for morph target animations -- defaulting to 10, but really it is completely arbitrary.
				var morphAnimationClips = THREE.AnimationClip.CreateClipsFromMorphTargetSequences( geometry.morphTargets, 10 );
				outputAnimations = outputAnimations.concat( morphAnimationClips );

			}

			if ( outputAnimations.length > 0 ) geometry.animations = outputAnimations;

		}

		if ( json.materials === undefined || json.materials.length === 0 ) {

			return { geometry: geometry };

		} else {

			var materials = THREE.Loader.prototype.initMaterials( json.materials, texturePath, this.crossOrigin );

			return { geometry: geometry, materials: materials };

		}

	}

};

// File:src/loaders/LoadingManager.js

/**
 * @author mrdoob / http://mrdoob.com/
 */

THREE.LoadingManager = function ( onLoad, onProgress, onError ) {

	var scope = this;

	var isLoading = false, itemsLoaded = 0, itemsTotal = 0;

	this.onStart = undefined;
	this.onLoad = onLoad;
	this.onProgress = onProgress;
	this.onError = onError;

	this.itemStart = function ( url ) {

		itemsTotal ++;

		if ( isLoading === false ) {

			if ( scope.onStart !== undefined ) {

				scope.onStart( url, itemsLoaded, itemsTotal );

			}

		}

		isLoading = true;

	};

	this.itemEnd = function ( url ) {

		itemsLoaded ++;

		if ( scope.onProgress !== undefined ) {

			scope.onProgress( url, itemsLoaded, itemsTotal );

		}

		if ( itemsLoaded === itemsTotal ) {

			isLoading = false;

			if ( scope.onLoad !== undefined ) {

				scope.onLoad();

			}

		}

	};

	this.itemError = function ( url ) {

		if ( scope.onError !== undefined ) {

			scope.onError( url );

		}

	};

};

THREE.DefaultLoadingManager = new THREE.LoadingManager();

// File:src/loaders/BufferGeometryLoader.js

/**
 * @author mrdoob / http://mrdoob.com/
 */

THREE.BufferGeometryLoader = function ( manager ) {

	this.manager = ( manager !== undefined ) ? manager : THREE.DefaultLoadingManager;

};

THREE.BufferGeometryLoader.prototype = {

	constructor: THREE.BufferGeometryLoader,

	load: function ( url, onLoad, onProgress, onError ) {

		var scope = this;

		var loader = new THREE.XHRLoader( scope.manager );
		loader.load( url, function ( text ) {

			onLoad( scope.parse( JSON.parse( text ) ) );

		}, onProgress, onError );

	},

	parse: function ( json ) {

		var geometry = new THREE.BufferGeometry();

		var index = json.data.index;

		var TYPED_ARRAYS = {
			'Int8Array': Int8Array,
			'Uint8Array': Uint8Array,
			'Uint8ClampedArray': Uint8ClampedArray,
			'Int16Array': Int16Array,
			'Uint16Array': Uint16Array,
			'Int32Array': Int32Array,
			'Uint32Array': Uint32Array,
			'Float32Array': Float32Array,
			'Float64Array': Float64Array
		};

		if ( index !== undefined ) {

			var typedArray = new TYPED_ARRAYS[ index.type ]( index.array );
			geometry.setIndex( new THREE.BufferAttribute( typedArray, 1 ) );

		}

		var attributes = json.data.attributes;

		for ( var key in attributes ) {

			var attribute = attributes[ key ];
			var typedArray = new TYPED_ARRAYS[ attribute.type ]( attribute.array );

			geometry.addAttribute( key, new THREE.BufferAttribute( typedArray, attribute.itemSize, attribute.normalized ) );

		}

		var groups = json.data.groups || json.data.drawcalls || json.data.offsets;

		if ( groups !== undefined ) {

			for ( var i = 0, n = groups.length; i !== n; ++ i ) {

				var group = groups[ i ];

				geometry.addGroup( group.start, group.count, group.materialIndex );

			}

		}

		var boundingSphere = json.data.boundingSphere;

		if ( boundingSphere !== undefined ) {

			var center = new THREE.Vector3();

			if ( boundingSphere.center !== undefined ) {

				center.fromArray( boundingSphere.center );

			}

			geometry.boundingSphere = new THREE.Sphere( center, boundingSphere.radius );

		}

		return geometry;

	}

};

// File:src/loaders/MaterialLoader.js

/**
 * @author mrdoob / http://mrdoob.com/
 */

THREE.MaterialLoader = function ( manager ) {

	this.manager = ( manager !== undefined ) ? manager : THREE.DefaultLoadingManager;
	this.textures = {};

};

THREE.MaterialLoader.prototype = {

	constructor: THREE.MaterialLoader,

	load: function ( url, onLoad, onProgress, onError ) {

		var scope = this;

		var loader = new THREE.XHRLoader( scope.manager );
		loader.load( url, function ( text ) {

			onLoad( scope.parse( JSON.parse( text ) ) );

		}, onProgress, onError );

	},

	setTextures: function ( value ) {

		this.textures = value;

	},

	getTexture: function ( name ) {

		var textures = this.textures;

		if ( textures[ name ] === undefined ) {

			console.warn( 'THREE.MaterialLoader: Undefined texture', name );

		}

		return textures[ name ];

	},

	parse: function ( json ) {

		var material = new THREE[ json.type ];

		if ( json.uuid !== undefined ) material.uuid = json.uuid;
		if ( json.name !== undefined ) material.name = json.name;
		if ( json.color !== undefined ) material.color.setHex( json.color );
		if ( json.roughness !== undefined ) material.roughness = json.roughness;
		if ( json.metalness !== undefined ) material.metalness = json.metalness;
		if ( json.emissive !== undefined ) material.emissive.setHex( json.emissive );
		if ( json.specular !== undefined ) material.specular.setHex( json.specular );
		if ( json.shininess !== undefined ) material.shininess = json.shininess;
		if ( json.uniforms !== undefined ) material.uniforms = json.uniforms;
		if ( json.vertexShader !== undefined ) material.vertexShader = json.vertexShader;
		if ( json.fragmentShader !== undefined ) material.fragmentShader = json.fragmentShader;
		if ( json.vertexColors !== undefined ) material.vertexColors = json.vertexColors;
		if ( json.shading !== undefined ) material.shading = json.shading;
		if ( json.blending !== undefined ) material.blending = json.blending;
		if ( json.side !== undefined ) material.side = json.side;
		if ( json.opacity !== undefined ) material.opacity = json.opacity;
		if ( json.transparent !== undefined ) material.transparent = json.transparent;
		if ( json.alphaTest !== undefined ) material.alphaTest = json.alphaTest;
		if ( json.depthTest !== undefined ) material.depthTest = json.depthTest;
		if ( json.depthWrite !== undefined ) material.depthWrite = json.depthWrite;
		if ( json.colorWrite !== undefined ) material.colorWrite = json.colorWrite;
		if ( json.wireframe !== undefined ) material.wireframe = json.wireframe;
		if ( json.wireframeLinewidth !== undefined ) material.wireframeLinewidth = json.wireframeLinewidth;

		// for PointsMaterial
		if ( json.size !== undefined ) material.size = json.size;
		if ( json.sizeAttenuation !== undefined ) material.sizeAttenuation = json.sizeAttenuation;

		// maps

		if ( json.map !== undefined ) material.map = this.getTexture( json.map );

		if ( json.alphaMap !== undefined ) {

			material.alphaMap = this.getTexture( json.alphaMap );
			material.transparent = true;

		}

		if ( json.bumpMap !== undefined ) material.bumpMap = this.getTexture( json.bumpMap );
		if ( json.bumpScale !== undefined ) material.bumpScale = json.bumpScale;

		if ( json.normalMap !== undefined ) material.normalMap = this.getTexture( json.normalMap );
		if ( json.normalScale !== undefined ) {

			var normalScale = json.normalScale;

			if ( Array.isArray( normalScale ) === false ) {

				// Blender exporter used to export a scalar. See #7459

				normalScale = [ normalScale, normalScale ];

			}

			material.normalScale = new THREE.Vector2().fromArray( normalScale );

		}

		if ( json.displacementMap !== undefined ) material.displacementMap = this.getTexture( json.displacementMap );
		if ( json.displacementScale !== undefined ) material.displacementScale = json.displacementScale;
		if ( json.displacementBias !== undefined ) material.displacementBias = json.displacementBias;

		if ( json.roughnessMap !== undefined ) material.roughnessMap = this.getTexture( json.roughnessMap );
		if ( json.metalnessMap !== undefined ) material.metalnessMap = this.getTexture( json.metalnessMap );

		if ( json.emissiveMap !== undefined ) material.emissiveMap = this.getTexture( json.emissiveMap );
		if ( json.emissiveIntensity !== undefined ) material.emissiveIntensity = json.emissiveIntensity;

		if ( json.specularMap !== undefined ) material.specularMap = this.getTexture( json.specularMap );

		if ( json.envMap !== undefined ) {

			material.envMap = this.getTexture( json.envMap );
			material.combine = THREE.MultiplyOperation;

		}

		if ( json.reflectivity ) material.reflectivity = json.reflectivity;

		if ( json.lightMap !== undefined ) material.lightMap = this.getTexture( json.lightMap );
		if ( json.lightMapIntensity !== undefined ) material.lightMapIntensity = json.lightMapIntensity;

		if ( json.aoMap !== undefined ) material.aoMap = this.getTexture( json.aoMap );
		if ( json.aoMapIntensity !== undefined ) material.aoMapIntensity = json.aoMapIntensity;

		// MultiMaterial

		if ( json.materials !== undefined ) {

			for ( var i = 0, l = json.materials.length; i < l; i ++ ) {

				material.materials.push( this.parse( json.materials[ i ] ) );

			}

		}

		return material;

	}

};

// File:src/loaders/ObjectLoader.js

/**
 * @author mrdoob / http://mrdoob.com/
 */

THREE.ObjectLoader = function ( manager ) {

	this.manager = ( manager !== undefined ) ? manager : THREE.DefaultLoadingManager;
	this.texturePath = '';

};

THREE.ObjectLoader.prototype = {

	constructor: THREE.ObjectLoader,

	load: function ( url, onLoad, onProgress, onError ) {

		if ( this.texturePath === '' ) {

			this.texturePath = url.substring( 0, url.lastIndexOf( '/' ) + 1 );

		}

		var scope = this;

		var loader = new THREE.XHRLoader( scope.manager );
		loader.load( url, function ( text ) {

			scope.parse( JSON.parse( text ), onLoad );

		}, onProgress, onError );

	},

	setTexturePath: function ( value ) {

		this.texturePath = value;

	},

	setCrossOrigin: function ( value ) {

		this.crossOrigin = value;

	},

	parse: function ( json, onLoad ) {

		var geometries = this.parseGeometries( json.geometries );

		var images = this.parseImages( json.images, function () {

			if ( onLoad !== undefined ) onLoad( object );

		} );

		var textures  = this.parseTextures( json.textures, images );
		var materials = this.parseMaterials( json.materials, textures );

		var object = this.parseObject( json.object, geometries, materials );

		if ( json.animations ) {

			object.animations = this.parseAnimations( json.animations );

		}

		if ( json.images === undefined || json.images.length === 0 ) {

			if ( onLoad !== undefined ) onLoad( object );

		}

		return object;

	},

	parseGeometries: function ( json ) {

		var geometries = {};

		if ( json !== undefined ) {

			var geometryLoader = new THREE.JSONLoader();
			var bufferGeometryLoader = new THREE.BufferGeometryLoader();

			for ( var i = 0, l = json.length; i < l; i ++ ) {

				var geometry;
				var data = json[ i ];

				switch ( data.type ) {

					case 'PlaneGeometry':
					case 'PlaneBufferGeometry':

						geometry = new THREE[ data.type ](
							data.width,
							data.height,
							data.widthSegments,
							data.heightSegments
						);

						break;

					case 'BoxGeometry':
					case 'BoxBufferGeometry':
					case 'CubeGeometry': // backwards compatible

						geometry = new THREE[ data.type ](
							data.width,
							data.height,
							data.depth,
							data.widthSegments,
							data.heightSegments,
							data.depthSegments
						);

						break;

					case 'CircleGeometry':
					case 'CircleBufferGeometry':

						geometry = new THREE[ data.type ](
							data.radius,
							data.segments,
							data.thetaStart,
							data.thetaLength
						);

						break;

					case 'CylinderGeometry':
					case 'CylinderBufferGeometry':

						geometry = new THREE[ data.type ](
							data.radiusTop,
							data.radiusBottom,
							data.height,
							data.radialSegments,
							data.heightSegments,
							data.openEnded,
							data.thetaStart,
							data.thetaLength
						);

						break;

					case 'ConeGeometry':
					case 'ConeBufferGeometry':

						geometry = new THREE [ data.type ](
							data.radius,
							data.height,
							data.radialSegments,
							data.heightSegments,
							data.openEnded,
							data.thetaStart,
							data.thetaLength
						);

						break;

					case 'SphereGeometry':
					case 'SphereBufferGeometry':

						geometry = new THREE[ data.type ](
							data.radius,
							data.widthSegments,
							data.heightSegments,
							data.phiStart,
							data.phiLength,
							data.thetaStart,
							data.thetaLength
						);

						break;

					case 'DodecahedronGeometry':
					case 'IcosahedronGeometry':
					case 'OctahedronGeometry':
					case 'TetrahedronGeometry':

						geometry = new THREE[ data.type ](
							data.radius,
							data.detail
						);

						break;

					case 'RingGeometry':
					case 'RingBufferGeometry':

						geometry = new THREE[ data.type ](
							data.innerRadius,
							data.outerRadius,
							data.thetaSegments,
							data.phiSegments,
							data.thetaStart,
							data.thetaLength
						);

						break;

					case 'TorusGeometry':
					case 'TorusBufferGeometry':

						geometry = new THREE[ data.type ](
							data.radius,
							data.tube,
							data.radialSegments,
							data.tubularSegments,
							data.arc
						);

						break;

					case 'TorusKnotGeometry':
					case 'TorusKnotBufferGeometry':

						geometry = new THREE[ data.type ](
							data.radius,
							data.tube,
							data.tubularSegments,
							data.radialSegments,
							data.p,
							data.q
						);

						break;

					case 'LatheGeometry':
					case 'LatheBufferGeometry':

						geometry = new THREE[ data.type ](
							data.points,
							data.segments,
							data.phiStart,
							data.phiLength
						);

						break;

					case 'BufferGeometry':

						geometry = bufferGeometryLoader.parse( data );

						break;

					case 'Geometry':

						geometry = geometryLoader.parse( data.data, this.texturePath ).geometry;

						break;

					default:

						console.warn( 'THREE.ObjectLoader: Unsupported geometry type "' + data.type + '"' );

						continue;

				}

				geometry.uuid = data.uuid;

				if ( data.name !== undefined ) geometry.name = data.name;

				geometries[ data.uuid ] = geometry;

			}

		}

		return geometries;

	},

	parseMaterials: function ( json, textures ) {

		var materials = {};

		if ( json !== undefined ) {

			var loader = new THREE.MaterialLoader();
			loader.setTextures( textures );

			for ( var i = 0, l = json.length; i < l; i ++ ) {

				var material = loader.parse( json[ i ] );
				materials[ material.uuid ] = material;

			}

		}

		return materials;

	},

	parseAnimations: function ( json ) {

		var animations = [];

		for ( var i = 0; i < json.length; i ++ ) {

			var clip = THREE.AnimationClip.parse( json[ i ] );

			animations.push( clip );

		}

		return animations;

	},

	parseImages: function ( json, onLoad ) {

		var scope = this;
		var images = {};

		function loadImage( url ) {

			scope.manager.itemStart( url );

			return loader.load( url, function () {

				scope.manager.itemEnd( url );

			} );

		}

		if ( json !== undefined && json.length > 0 ) {

			var manager = new THREE.LoadingManager( onLoad );

			var loader = new THREE.ImageLoader( manager );
			loader.setCrossOrigin( this.crossOrigin );

			for ( var i = 0, l = json.length; i < l; i ++ ) {

				var image = json[ i ];
				var path = /^(\/\/)|([a-z]+:(\/\/)?)/i.test( image.url ) ? image.url : scope.texturePath + image.url;

				images[ image.uuid ] = loadImage( path );

			}

		}

		return images;

	},

	parseTextures: function ( json, images ) {

		function parseConstant( value ) {

			if ( typeof( value ) === 'number' ) return value;

			console.warn( 'THREE.ObjectLoader.parseTexture: Constant should be in numeric form.', value );

			return THREE[ value ];

		}

		var textures = {};

		if ( json !== undefined ) {

			for ( var i = 0, l = json.length; i < l; i ++ ) {

				var data = json[ i ];

				if ( data.image === undefined ) {

					console.warn( 'THREE.ObjectLoader: No "image" specified for', data.uuid );

				}

				if ( images[ data.image ] === undefined ) {

					console.warn( 'THREE.ObjectLoader: Undefined image', data.image );

				}

				var texture = new THREE.Texture( images[ data.image ] );
				texture.needsUpdate = true;

				texture.uuid = data.uuid;

				if ( data.name !== undefined ) texture.name = data.name;
				if ( data.mapping !== undefined ) texture.mapping = parseConstant( data.mapping );
				if ( data.offset !== undefined ) texture.offset = new THREE.Vector2( data.offset[ 0 ], data.offset[ 1 ] );
				if ( data.repeat !== undefined ) texture.repeat = new THREE.Vector2( data.repeat[ 0 ], data.repeat[ 1 ] );
				if ( data.minFilter !== undefined ) texture.minFilter = parseConstant( data.minFilter );
				if ( data.magFilter !== undefined ) texture.magFilter = parseConstant( data.magFilter );
				if ( data.anisotropy !== undefined ) texture.anisotropy = data.anisotropy;
				if ( Array.isArray( data.wrap ) ) {

					texture.wrapS = parseConstant( data.wrap[ 0 ] );
					texture.wrapT = parseConstant( data.wrap[ 1 ] );

				}

				textures[ data.uuid ] = texture;

			}

		}

		return textures;

	},

	parseObject: function () {

		var matrix = new THREE.Matrix4();

		return function ( data, geometries, materials ) {

			var object;

			function getGeometry( name ) {

				if ( geometries[ name ] === undefined ) {

					console.warn( 'THREE.ObjectLoader: Undefined geometry', name );

				}

				return geometries[ name ];

			}

			function getMaterial( name ) {

				if ( name === undefined ) return undefined;

				if ( materials[ name ] === undefined ) {

					console.warn( 'THREE.ObjectLoader: Undefined material', name );

				}

				return materials[ name ];

			}

			switch ( data.type ) {

				case 'Scene':

					object = new THREE.Scene();

					break;

				case 'PerspectiveCamera':

					object = new THREE.PerspectiveCamera( data.fov, data.aspect, data.near, data.far );

					if ( data.focus !== undefined ) object.focus = data.focus;
					if ( data.zoom !== undefined ) object.zoom = data.zoom;
					if ( data.filmGauge !== undefined ) object.filmGauge = data.filmGauge;
					if ( data.filmOffset !== undefined ) object.filmOffset = data.filmOffset;
					if ( data.view !== undefined ) object.view = Object.assign( {}, data.view );

					break;

				case 'OrthographicCamera':

					object = new THREE.OrthographicCamera( data.left, data.right, data.top, data.bottom, data.near, data.far );

					break;

				case 'AmbientLight':

					object = new THREE.AmbientLight( data.color, data.intensity );

					break;

				case 'DirectionalLight':

					object = new THREE.DirectionalLight( data.color, data.intensity );

					break;

				case 'PointLight':

					object = new THREE.PointLight( data.color, data.intensity, data.distance, data.decay );

					break;

				case 'SpotLight':

					object = new THREE.SpotLight( data.color, data.intensity, data.distance, data.angle, data.penumbra, data.decay );

					break;

				case 'HemisphereLight':

					object = new THREE.HemisphereLight( data.color, data.groundColor, data.intensity );

					break;

				case 'Mesh':

					var geometry = getGeometry( data.geometry );
					var material = getMaterial( data.material );

					if ( geometry.bones && geometry.bones.length > 0 ) {

						object = new THREE.SkinnedMesh( geometry, material );

					} else {

						object = new THREE.Mesh( geometry, material );

					}

					break;

				case 'LOD':

					object = new THREE.LOD();

					break;

				case 'Line':

					object = new THREE.Line( getGeometry( data.geometry ), getMaterial( data.material ), data.mode );

					break;

				case 'PointCloud':
				case 'Points':

					object = new THREE.Points( getGeometry( data.geometry ), getMaterial( data.material ) );

					break;

				case 'Sprite':

					object = new THREE.Sprite( getMaterial( data.material ) );

					break;

				case 'Group':

					object = new THREE.Group();

					break;

				default:

					object = new THREE.Object3D();

			}

			object.uuid = data.uuid;

			if ( data.name !== undefined ) object.name = data.name;
			if ( data.matrix !== undefined ) {

				matrix.fromArray( data.matrix );
				matrix.decompose( object.position, object.quaternion, object.scale );

			} else {

				if ( data.position !== undefined ) object.position.fromArray( data.position );
				if ( data.rotation !== undefined ) object.rotation.fromArray( data.rotation );
				if ( data.scale !== undefined ) object.scale.fromArray( data.scale );

			}

			if ( data.castShadow !== undefined ) object.castShadow = data.castShadow;
			if ( data.receiveShadow !== undefined ) object.receiveShadow = data.receiveShadow;

			if ( data.visible !== undefined ) object.visible = data.visible;
			if ( data.userData !== undefined ) object.userData = data.userData;

			if ( data.children !== undefined ) {

				for ( var child in data.children ) {

					object.add( this.parseObject( data.children[ child ], geometries, materials ) );

				}

			}

			if ( data.type === 'LOD' ) {

				var levels = data.levels;

				for ( var l = 0; l < levels.length; l ++ ) {

					var level = levels[ l ];
					var child = object.getObjectByProperty( 'uuid', level.object );

					if ( child !== undefined ) {

						object.addLevel( child, level.distance );

					}

				}

			}

			return object;

		};

	}()

};

// File:src/loaders/TextureLoader.js

/**
 * @author mrdoob / http://mrdoob.com/
 */

THREE.TextureLoader = function ( manager ) {

	this.manager = ( manager !== undefined ) ? manager : THREE.DefaultLoadingManager;

};

THREE.TextureLoader.prototype = {

	constructor: THREE.TextureLoader,

	load: function ( url, onLoad, onProgress, onError ) {

		var texture = new THREE.Texture();

		var loader = new THREE.ImageLoader( this.manager );
		loader.setCrossOrigin( this.crossOrigin );
		loader.setPath( this.path );
		loader.load( url, function ( image ) {

			texture.image = image;
			texture.needsUpdate = true;

			if ( onLoad !== undefined ) {

				onLoad( texture );

			}

		}, onProgress, onError );

		return texture;

	},

	setCrossOrigin: function ( value ) {

		this.crossOrigin = value;

	},

	setPath: function ( value ) {

		this.path = value;

	}

};

// File:src/loaders/CubeTextureLoader.js

/**
 * @author mrdoob / http://mrdoob.com/
 */

THREE.CubeTextureLoader = function ( manager ) {

	this.manager = ( manager !== undefined ) ? manager : THREE.DefaultLoadingManager;

};

THREE.CubeTextureLoader.prototype = {

	constructor: THREE.CubeTextureLoader,

	load: function ( urls, onLoad, onProgress, onError ) {

		var texture = new THREE.CubeTexture();

		var loader = new THREE.ImageLoader( this.manager );
		loader.setCrossOrigin( this.crossOrigin );
		loader.setPath( this.path );

		var loaded = 0;

		function loadTexture( i ) {

			loader.load( urls[ i ], function ( image ) {

				texture.images[ i ] = image;

				loaded ++;

				if ( loaded === 6 ) {

					texture.needsUpdate = true;

					if ( onLoad ) onLoad( texture );

				}

			}, undefined, onError );

		}

		for ( var i = 0; i < urls.length; ++ i ) {

			loadTexture( i );

		}

		return texture;

	},

	setCrossOrigin: function ( value ) {

		this.crossOrigin = value;

	},

	setPath: function ( value ) {

		this.path = value;

	}

};

// File:src/loaders/BinaryTextureLoader.js

/**
 * @author Nikos M. / https://github.com/foo123/
 *
 * Abstract Base class to load generic binary textures formats (rgbe, hdr, ...)
 */

THREE.DataTextureLoader = THREE.BinaryTextureLoader = function ( manager ) {

	this.manager = ( manager !== undefined ) ? manager : THREE.DefaultLoadingManager;

	// override in sub classes
	this._parser = null;

};

THREE.BinaryTextureLoader.prototype = {

	constructor: THREE.BinaryTextureLoader,

	load: function ( url, onLoad, onProgress, onError ) {

		var scope = this;

		var texture = new THREE.DataTexture();

		var loader = new THREE.XHRLoader( this.manager );
		loader.setResponseType( 'arraybuffer' );

		loader.load( url, function ( buffer ) {

			var texData = scope._parser( buffer );

			if ( ! texData ) return;

			if ( undefined !== texData.image ) {

				texture.image = texData.image;

			} else if ( undefined !== texData.data ) {

				texture.image.width = texData.width;
				texture.image.height = texData.height;
				texture.image.data = texData.data;

			}

			texture.wrapS = undefined !== texData.wrapS ? texData.wrapS : THREE.ClampToEdgeWrapping;
			texture.wrapT = undefined !== texData.wrapT ? texData.wrapT : THREE.ClampToEdgeWrapping;

			texture.magFilter = undefined !== texData.magFilter ? texData.magFilter : THREE.LinearFilter;
			texture.minFilter = undefined !== texData.minFilter ? texData.minFilter : THREE.LinearMipMapLinearFilter;

			texture.anisotropy = undefined !== texData.anisotropy ? texData.anisotropy : 1;

			if ( undefined !== texData.format ) {

				texture.format = texData.format;

			}
			if ( undefined !== texData.type ) {

				texture.type = texData.type;

			}

			if ( undefined !== texData.mipmaps ) {

				texture.mipmaps = texData.mipmaps;

			}

			if ( 1 === texData.mipmapCount ) {

				texture.minFilter = THREE.LinearFilter;

			}

			texture.needsUpdate = true;

			if ( onLoad ) onLoad( texture, texData );

		}, onProgress, onError );


		return texture;

	}

};

// File:src/loaders/CompressedTextureLoader.js

/**
 * @author mrdoob / http://mrdoob.com/
 *
 * Abstract Base class to block based textures loader (dds, pvr, ...)
 */

THREE.CompressedTextureLoader = function ( manager ) {

	this.manager = ( manager !== undefined ) ? manager : THREE.DefaultLoadingManager;

	// override in sub classes
	this._parser = null;

};


THREE.CompressedTextureLoader.prototype = {

	constructor: THREE.CompressedTextureLoader,

	load: function ( url, onLoad, onProgress, onError ) {

		var scope = this;

		var images = [];

		var texture = new THREE.CompressedTexture();
		texture.image = images;

		var loader = new THREE.XHRLoader( this.manager );
		loader.setPath( this.path );
		loader.setResponseType( 'arraybuffer' );

		function loadTexture( i ) {

			loader.load( url[ i ], function ( buffer ) {

				var texDatas = scope._parser( buffer, true );

				images[ i ] = {
					width: texDatas.width,
					height: texDatas.height,
					format: texDatas.format,
					mipmaps: texDatas.mipmaps
				};

				loaded += 1;

				if ( loaded === 6 ) {

					if ( texDatas.mipmapCount === 1 )
						texture.minFilter = THREE.LinearFilter;

					texture.format = texDatas.format;
					texture.needsUpdate = true;

					if ( onLoad ) onLoad( texture );

				}

			}, onProgress, onError );

		}

		if ( Array.isArray( url ) ) {

			var loaded = 0;

			for ( var i = 0, il = url.length; i < il; ++ i ) {

				loadTexture( i );

			}

		} else {

			// compressed cubemap texture stored in a single DDS file

			loader.load( url, function ( buffer ) {

				var texDatas = scope._parser( buffer, true );

				if ( texDatas.isCubemap ) {

					var faces = texDatas.mipmaps.length / texDatas.mipmapCount;

					for ( var f = 0; f < faces; f ++ ) {

						images[ f ] = { mipmaps : [] };

						for ( var i = 0; i < texDatas.mipmapCount; i ++ ) {

							images[ f ].mipmaps.push( texDatas.mipmaps[ f * texDatas.mipmapCount + i ] );
							images[ f ].format = texDatas.format;
							images[ f ].width = texDatas.width;
							images[ f ].height = texDatas.height;

						}

					}

				} else {

					texture.image.width = texDatas.width;
					texture.image.height = texDatas.height;
					texture.mipmaps = texDatas.mipmaps;

				}

				if ( texDatas.mipmapCount === 1 ) {

					texture.minFilter = THREE.LinearFilter;

				}

				texture.format = texDatas.format;
				texture.needsUpdate = true;

				if ( onLoad ) onLoad( texture );

			}, onProgress, onError );

		}

		return texture;

	},

	setPath: function ( value ) {

		this.path = value;

	}

};

// File:src/materials/Material.js

/**
 * @author mrdoob / http://mrdoob.com/
 * @author alteredq / http://alteredqualia.com/
 */

THREE.Material = function () {

	Object.defineProperty( this, 'id', { value: THREE.MaterialIdCount ++ } );

	this.uuid = THREE.Math.generateUUID();

	this.name = '';
	this.type = 'Material';

	this.fog = true;
	this.lights = true;

	this.blending = THREE.NormalBlending;
	this.side = THREE.FrontSide;
	this.shading = THREE.SmoothShading; // THREE.FlatShading, THREE.SmoothShading
	this.vertexColors = THREE.NoColors; // THREE.NoColors, THREE.VertexColors, THREE.FaceColors

	this.opacity = 1;
	this.transparent = false;

	this.blendSrc = THREE.SrcAlphaFactor;
	this.blendDst = THREE.OneMinusSrcAlphaFactor;
	this.blendEquation = THREE.AddEquation;
	this.blendSrcAlpha = null;
	this.blendDstAlpha = null;
	this.blendEquationAlpha = null;

	this.depthFunc = THREE.LessEqualDepth;
	this.depthTest = true;
	this.depthWrite = true;

	this.clippingPlanes = null;
	this.clipShadows = false;

	this.colorWrite = true;

	this.precision = null; // override the renderer's default precision for this material

	this.polygonOffset = false;
	this.polygonOffsetFactor = 0;
	this.polygonOffsetUnits = 0;

	this.alphaTest = 0;
	this.premultipliedAlpha = false;

	this.overdraw = 0; // Overdrawn pixels (typically between 0 and 1) for fixing antialiasing gaps in CanvasRenderer

	this.visible = true;

	this._needsUpdate = true;

};

THREE.Material.prototype = {

	constructor: THREE.Material,

	get needsUpdate() {

		return this._needsUpdate;

	},

	set needsUpdate( value ) {

		if ( value === true ) this.update();
		this._needsUpdate = value;

	},

	setValues: function ( values ) {

		if ( values === undefined ) return;

		for ( var key in values ) {

			var newValue = values[ key ];

			if ( newValue === undefined ) {

				console.warn( "THREE.Material: '" + key + "' parameter is undefined." );
				continue;

			}

			var currentValue = this[ key ];

			if ( currentValue === undefined ) {

				console.warn( "THREE." + this.type + ": '" + key + "' is not a property of this material." );
				continue;

			}

			if ( currentValue instanceof THREE.Color ) {

				currentValue.set( newValue );

			} else if ( currentValue instanceof THREE.Vector3 && newValue instanceof THREE.Vector3 ) {

				currentValue.copy( newValue );

			} else if ( key === 'overdraw' ) {

				// ensure overdraw is backwards-compatible with legacy boolean type
				this[ key ] = Number( newValue );

			} else {

				this[ key ] = newValue;

			}

		}

	},

	toJSON: function ( meta ) {

		var isRoot = meta === undefined;

		if ( isRoot ) {

			meta = {
				textures: {},
				images: {}
			};

		}

		var data = {
			metadata: {
				version: 4.4,
				type: 'Material',
				generator: 'Material.toJSON'
			}
		};

		// standard Material serialization
		data.uuid = this.uuid;
		data.type = this.type;

		if ( this.name !== '' ) data.name = this.name;

		if ( this.color instanceof THREE.Color ) data.color = this.color.getHex();
```
