## Supplementary Dataset 5 for "A 3D molecular atlas of the chick embryonic heart": TrackballControls.js

```
/**
 * @author Eberhard Graether / http://egraether.com/
 * @author Mark Lundin 	/ http://mark-lundin.com
 * @author Bill Hill / http://www.emouseatlas.org
 */

THREE.TrackballControls = function ( object, domElement ) {

	var _this = this;
	var STATE = { NONE: -1, ROTATE: 0, ZOOM: 1, PAN: 2, TOUCH_ROTATE: 3, TOUCH_ZOOM: 4, TOUCH_PAN: 5 };

	this.object = object;
	this.domElement = ( domElement !== undefined ) ? domElement : document;

	// API

	this.enabled = true;

	this.screen = { left: 0, top: 0, width: 0, height: 0 };

	this.rotateSpeed = 1.0;
	this.zoomSpeed = 1.2;
	this.panSpeed = 0.3;

	this.noRotate = false;
	this.noZoom = false;
	this.noPan = false;
	this.noRoll = false;

	this.staticMoving = false;
	this.dynamicDampingFactor = 0.2;

	this.minDistance = 0;
	this.maxDistance = Infinity;

	this.keys = [ 65 /*A*/, 83 /*S*/, 68 /*D*/ ];

	// internals

	this.target = new THREE.Vector3();

	var EPS = 0.000001;

	var lastPosition = new THREE.Vector3();

	var _state = STATE.NONE,
	_prevState = STATE.NONE,

	_eye = new THREE.Vector3(),

	_rotateStart = new THREE.Vector3(),
	_rotateEnd = new THREE.Vector3(),

	_zoomStart = new THREE.Vector2(),
	_zoomEnd = new THREE.Vector2(),

	_touchZoomDistanceStart = 0,
	_touchZoomDistanceEnd = 0,

	_panStart = new THREE.Vector2(),
	_panEnd = new THREE.Vector2();

	// for reset

	this.target0 = this.target.clone();
	this.position0 = this.object.position.clone();
	this.up0 = this.object.up.clone();

	// events

	var changeEvent = { type: 'change' };
	var startEvent = { type: 'start'};
	var endEvent = { type: 'end'};


	// methods

	this.handleResize = function () {

		if ( this.domElement === document ) {

			this.screen.left = 0;
			this.screen.top = 0;
			this.screen.width = window.innerWidth;
			this.screen.height = window.innerHeight;

		} else {

			var box = this.domElement.getBoundingClientRect();
			// adjustments come from similar code in the jquery offset() function
			var d = this.domElement.ownerDocument.documentElement;
			this.screen.left = box.left + window.pageXOffset - d.clientLeft;
			this.screen.top = box.top + window.pageYOffset - d.clientTop;
			this.screen.width = box.width;
			this.screen.height = box.height;

		}

	};

	this.handleEvent = function ( event ) {

		if ( typeof this[ event.type ] == 'function' ) {

			this[ event.type ]( event );

		}

	};

	this.getMouseOnScreen = function ( pageX, pageY, vector ) {

		return vector.set(
			( pageX - _this.screen.left ) / _this.screen.width,
			( pageY - _this.screen.top ) / _this.screen.height
		);

	};

	this.getMouseProjectionOnBall = (function(){

		var objectUp = new THREE.Vector3(),
		    mouseOnBall = new THREE.Vector3();


		return function ( pageX, pageY, projection ) {

			mouseOnBall.set(
				( pageX - _this.screen.width * 0.5 - _this.screen.left ) / (_this.screen.width*.5),
				( _this.screen.height * 0.5 + _this.screen.top - pageY ) / (_this.screen.height*.5),
				0.0
			);

			var length = mouseOnBall.length();

			if ( _this.noRoll ) {

				if ( length < Math.SQRT1_2 ) {

					mouseOnBall.z = Math.sqrt( 1.0 - length*length );

				} else {

					mouseOnBall.z = .5 / length;
					
				}

			} else if ( length > 1.0 ) {

				mouseOnBall.normalize();

			} else {

				mouseOnBall.z = Math.sqrt( 1.0 - length * length );

			}

			_eye.copy( _this.object.position ).sub( _this.target );

			projection.copy( _this.object.up ).setLength( mouseOnBall.y )
			projection.add( objectUp.copy( _this.object.up ).cross( _eye ).setLength( mouseOnBall.x ) );
			projection.add( _eye.setLength( mouseOnBall.z ) );

			return projection;
		}

	}());

	this.rotateCamera = (function(){

		var axis = new THREE.Vector3(),
			quaternion = new THREE.Quaternion();


		return function () {

			var angle = Math.acos( _rotateStart.dot( _rotateEnd ) / _rotateStart.length() / _rotateEnd.length() );

			if ( angle ) {

				axis.crossVectors( _rotateStart, _rotateEnd ).normalize();

				angle *= _this.rotateSpeed;

				quaternion.setFromAxisAngle( axis, -angle );

				_eye.applyQuaternion( quaternion );
				_this.object.up.applyQuaternion( quaternion );

				_rotateEnd.applyQuaternion( quaternion );

				if ( _this.staticMoving ) {

					_rotateStart.copy( _rotateEnd );

				} else {

					quaternion.setFromAxisAngle( axis, angle * ( _this.dynamicDampingFactor - 1.0 ) );
					_rotateStart.applyQuaternion( quaternion );

				}

			}
		}

	}());

	this.zoomCamera = function () {

		if ( _state === STATE.TOUCH_ZOOM ) {

			var factor = _touchZoomDistanceStart / _touchZoomDistanceEnd;
			_touchZoomDistanceStart = _touchZoomDistanceEnd;
			_eye.multiplyScalar( factor );

		} else {

			var factor = 1.0 + ( _zoomEnd.y - _zoomStart.y ) * _this.zoomSpeed;

			if ( factor !== 1.0 && factor > 0.0 ) {

				_eye.multiplyScalar( factor );

				if ( _this.staticMoving ) {

					_zoomStart.copy( _zoomEnd );

				} else {

					_zoomStart.y += ( _zoomEnd.y - _zoomStart.y ) * this.dynamicDampingFactor;

				}

			}

		}

	};

	this.panCamera = (function(){

		var mouseChange = new THREE.Vector2(),
			objectUp = new THREE.Vector3(),
			pan = new THREE.Vector3();

		return function () {

			mouseChange.copy( _panEnd ).sub( _panStart );

			if ( mouseChange.lengthSq() ) {

				mouseChange.multiplyScalar( _eye.length() * _this.panSpeed );

				var u = new THREE.Vector3(),
				    v = new THREE.Vector3();
				u.crossVectors( _eye, _this.object.up );
				v.crossVectors( _eye, u ).setLength( -(mouseChange.y) );
				u.setLength( mouseChange.x );
				pan.addVectors( u, v );

				_this.object.position.add( pan );
				_this.target.add( pan );

				if ( _this.staticMoving ) {

					_panStart.copy( _panEnd );

				} else {

					_panStart.add( mouseChange.subVectors( _panEnd, _panStart ).multiplyScalar( _this.dynamicDampingFactor ) );

				}

			}
		}

	}());

	this.checkDistances = function () {

		if ( !_this.noZoom || !_this.noPan ) {

			if ( _eye.lengthSq() > _this.maxDistance * _this.maxDistance ) {

				_this.object.position.addVectors( _this.target, _eye.setLength( _this.maxDistance ) );

			}

			if ( _eye.lengthSq() < _this.minDistance * _this.minDistance ) {

				_this.object.position.addVectors( _this.target, _eye.setLength( _this.minDistance ) );

			}

		}

	};

	this.update = function () {

		_eye.subVectors( _this.object.position, _this.target );

		if ( !_this.noRotate ) {

			_this.rotateCamera();

		}

		if ( !_this.noZoom ) {

			_this.zoomCamera();

		}

		if ( !_this.noPan ) {

			_this.panCamera();

		}

		_this.object.position.addVectors( _this.target, _eye );

		_this.checkDistances();

		_this.object.lookAt( _this.target );

		if ( lastPosition.distanceToSquared( _this.object.position ) > EPS ) {

			_this.dispatchEvent( changeEvent );

			lastPosition.copy( _this.object.position );

		}

	};

	this.reset = function () {

		_state = STATE.NONE;
		_prevState = STATE.NONE;

		_this.target.copy( _this.target0 );
		_this.object.position.copy( _this.position0 );
		_this.object.up.copy( _this.up0 );

		_eye.subVectors( _this.object.position, _this.target );

		_this.object.lookAt( _this.target );

		_this.dispatchEvent( changeEvent );

		lastPosition.copy( _this.object.position );

	};

	// listeners

	function keydown( event ) {

		if ( _this.enabled === false ) return;

		window.removeEventListener( 'keydown', keydown );

		_prevState = _state;

		if ( _state !== STATE.NONE ) {

			return;

		} else if ( event.keyCode === _this.keys[ STATE.ROTATE ] && !_this.noRotate ) {

			_state = STATE.ROTATE;

		} else if ( event.keyCode === _this.keys[ STATE.ZOOM ] && !_this.noZoom ) {

			_state = STATE.ZOOM;

		} else if ( event.keyCode === _this.keys[ STATE.PAN ] && !_this.noPan ) {

			_state = STATE.PAN;

		}

	}

	function keyup( event ) {

		if ( _this.enabled === false ) return;

		_state = _prevState;

		window.addEventListener( 'keydown', keydown, false );

	}

	function mousedown( event ) {

		if ( _this.enabled === false ) return;

		event.preventDefault();
		event.stopPropagation();

		if ( _state === STATE.NONE ) {

			_state = event.button;

		}

		if ( _state === STATE.ROTATE && !_this.noRotate ) {

			_this.getMouseProjectionOnBall( event.pageX, event.pageY, _rotateStart );
			_rotateEnd.copy(_rotateStart)

		} else if ( _state === STATE.ZOOM && !_this.noZoom ) {

			_this.getMouseOnScreen( event.pageX, event.pageY, _zoomStart );
			_zoomEnd.copy(_zoomStart);

		} else if ( _state === STATE.PAN && !_this.noPan ) {

			_this.getMouseOnScreen( event.pageX, event.pageY, _panStart );
			_panEnd.copy(_panStart)

		}

		document.addEventListener( 'mousemove', mousemove, false );
		document.addEventListener( 'mouseup', mouseup, false );
		_this.dispatchEvent( startEvent );


	}

	function mousemove( event ) {

		if ( _this.enabled === false ) return;

		event.preventDefault();
		event.stopPropagation();

		if ( _state === STATE.ROTATE && !_this.noRotate ) {

			_this.getMouseProjectionOnBall( event.pageX, event.pageY, _rotateEnd );

		} else if ( _state === STATE.ZOOM && !_this.noZoom ) {

			_this.getMouseOnScreen( event.pageX, event.pageY, _zoomEnd );

		} else if ( _state === STATE.PAN && !_this.noPan ) {

			_this.getMouseOnScreen( event.pageX, event.pageY, _panEnd );

		}

	}

	function mouseup( event ) {

		if ( _this.enabled === false ) return;

		event.preventDefault();
		event.stopPropagation();

		_state = STATE.NONE;

		document.removeEventListener( 'mousemove', mousemove );
		document.removeEventListener( 'mouseup', mouseup );
		_this.dispatchEvent( endEvent );

	}

	function mousewheel( event ) {

		if ( _this.enabled === false ) return;

		event.preventDefault();
		event.stopPropagation();

		var delta = 0;

		if ( event.wheelDelta ) { // WebKit / Opera / Explorer 9

			delta = event.wheelDelta / 40;

		} else if ( event.detail ) { // Firefox

			delta = - event.detail / 3;

		}

		_zoomStart.y += delta * 0.01;
		_this.dispatchEvent( startEvent );
		_this.dispatchEvent( endEvent );

	}

	function touchstart( event ) {

		if ( _this.enabled === false ) return;

		switch ( event.touches.length ) {

			case 1:
				_state = STATE.TOUCH_ROTATE;
				_rotateEnd.copy( _this.getMouseProjectionOnBall( event.touches[ 0 ].pageX, event.touches[ 0 ].pageY, _rotateStart ));
				break;

			case 2:
				_state = STATE.TOUCH_ZOOM;
				var dx = event.touches[ 0 ].pageX - event.touches[ 1 ].pageX;
				var dy = event.touches[ 0 ].pageY - event.touches[ 1 ].pageY;
				_touchZoomDistanceEnd = _touchZoomDistanceStart = Math.sqrt( dx * dx + dy * dy );
				break;

			case 3:
				_state = STATE.TOUCH_PAN;
				_panEnd.copy( _this.getMouseOnScreen( event.touches[ 0 ].pageX, event.touches[ 0 ].pageY, _panStart ));
				break;

			default:
				_state = STATE.NONE;

		}
		_this.dispatchEvent( startEvent );


	}

	function touchmove( event ) {

		if ( _this.enabled === false ) return;

		event.preventDefault();
		event.stopPropagation();

		switch ( event.touches.length ) {

			case 1:
				_this.getMouseProjectionOnBall( event.touches[ 0 ].pageX, event.touches[ 0 ].pageY, _rotateEnd );
				break;

			case 2:
				var dx = event.touches[ 0 ].pageX - event.touches[ 1 ].pageX;
				var dy = event.touches[ 0 ].pageY - event.touches[ 1 ].pageY;
				_touchZoomDistanceEnd = Math.sqrt( dx * dx + dy * dy )
				break;

			case 3:
				_this.getMouseOnScreen( event.touches[ 0 ].pageX, event.touches[ 0 ].pageY, _panEnd );
				break;

			default:
				_state = STATE.NONE;

		}

	}

	function touchend( event ) {

		if ( _this.enabled === false ) return;

		switch ( event.touches.length ) {

			case 1:
				_rotateStart.copy( _this.getMouseProjectionOnBall( event.touches[ 0 ].pageX, event.touches[ 0 ].pageY, _rotateEnd ));
				break;

			case 2:
				_touchZoomDistanceStart = _touchZoomDistanceEnd = 0;
				break;

			case 3:
				_panStart.copy( _this.getMouseOnScreen( event.touches[ 0 ].pageX, event.touches[ 0 ].pageY, _panEnd ));
				break;

		}

		_state = STATE.NONE;
		_this.dispatchEvent( endEvent );

	}

	this.domElement.addEventListener( 'contextmenu', function ( event ) { event.preventDefault(); }, false );

	this.domElement.addEventListener( 'mousedown', mousedown, false );

	this.domElement.addEventListener( 'mousewheel', mousewheel, false );
	this.domElement.addEventListener( 'DOMMouseScroll', mousewheel, false ); // firefox

	this.domElement.addEventListener( 'touchstart', touchstart, false );
	this.domElement.addEventListener( 'touchend', touchend, false );
	this.domElement.addEventListener( 'touchmove', touchmove, false );

	window.addEventListener( 'keydown', keydown, false );
	window.addEventListener( 'keyup', keyup, false );

	this.handleResize();

	// force an update at start
	this.update();

};

THREE.TrackballControls.prototype = Object.create( THREE.EventDispatcher.prototype );
```
