## Supplementary Dataset 5 for "A 3D molecular atlas of the chick embryonic heart": README.txt

To view the 3D molecular atlas of the HH12 chick embryonic heart you can
use either Firefox [1], Chrome [2] or possibly another web browser. Of these
Firefox is the more straightforward as it allows you to simply load local
files without setting special options or using a local web server.
Chrome will allow you to load local media files if started using the
appropriate 'command line arguments'. For other web browsers you will
probably need to run a local web server.

*Firefox*
To load the viewer using Firefox either navigate in a file manager to the
folder containing this readme file and open the file

chick_heart_gene_expression.html

or run Firefox from a command line giving this file as an argument.

*Chrome*
It is a little more involved using Chrome to load the viewer, since by
default Chrome will not load local data. You can load the viewer using
Chrome, but the method varies slightly depending on the operating system
that you are using. Whatever your operating system you must make sure that
you are not currently running Chrome, since otherwise the existing
instance of Chrome will be used.

_Windows_
  1. Find the file path of your Chrome Installation, eg
    C:\Users\-your-user-name\AppData\Local\Google\Chrome\Application>
  2. Start the Chrome browser from the command line window with the
    additional argument ‘–allow-file-access-from-files’, eg
  C:\Users\-your-user-name\AppData\Local\Google\Chrome\Application\chrome.exe --allow-file-access-from-files
  3. Find the file path of this readme file, eg
    C:\Temp\ChickHeartAtlas\HH12
  4. Load the viewer application using Chrome by giving the URL
    file:///C:/Temp/ChickHeartAtlas/HH12/chick_heart_gene_expression.html
  A further complication when using Chrome under Windows is that you may
  need to unblock the files from the zip (using the file manager) to prevent
  Windows from "protecting" you from them. Unblocking the zip file (prior to
  unzipping it) may avoid you needing to do this for each file that is loaded
  by the interface.

_Linux_
  In a terminal change to the directory containing this readme file
  then enter
  google-chrome --allow-file-access-from-files chick_heart_gene_expression.html

_Mac OS X_
  Within a terminal window (Terminal.App) change folder to the folder
  containing this readme file then type
    open /Applications/Google\ Chrome.app --args  --allow-file-access-from-file chick_heart_gene_expression.html

For each of these operating systems the command is a single line.

Chrome prevents you from loading local files (by default) for your security,
so when using Chrome as above avoid loading untrusted internet web pages
(helpfully Firefox allows local web pages to load local files).

*Simple Web Server*
If you have Python[3] installed on your computer then another option is to
run a simple web server. To do this (again using the command line) open a
terminal (console application), change to the folder containing the chick
heart data (HH12), then start the simple python web server with the command

python -m SimpleHTTPServer 8000

you should then be able to use any (modern) web browser to view the 3D atlas
using the URL

http://127.0.0.1:8000/gxp-pointcloud_chick_heart_local.html

  
*Files*
The files in this folder are:
chick_heart_gene_expression.html - Web page which starts the local viewer application.
chick_heart_gene_expression_local.html - Copy of chick_heart_gene_expression.html.
chick_heart_gene_expression_server.html - Viewer application for on a web server.
chickheart-pointclouds-help.html - Usage/help information used by the interface.
chickheart-pointclouds.json - Configuration for local application without sectioning.
chickheart-pointclouds-nosectioning.json - Copy of chickheart-pointclouds.json.
chickheart-pointclouds-sectioning.json - Configuration for a web server with sectioning enabled.
gxp-pointcloud.css - Style sheet for the application.
gxp-pointcloud.html - The actual viewer application linked from chick_heart_gene_expression.html on a server.
icons/ - Icons required by the application.
js/ - Javascript required by the application.
PointClouds/ - Point clouds for use by the application.
RefSurfaces/ - The reference model surface.
Wlz/ - Woolz objects for use by a WlzIIP3D server to enable sectioning.

[1] https://www.mozilla.org/
[2] https://www.google.com/chrome/
[3] https://www.python.org/
