## Supplementary figures and images for "A 3D molecular atlas of the chick embryonic heart"

### bigdot.png

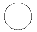

### cancel.png

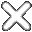

### dark.png

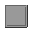

### eye.png

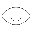

### gohome.png

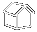

### help.png

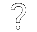

### light.png

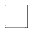

### load.png

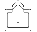

### minus.png

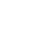

### noteye.png

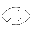

### notpointclouds.png

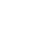

### notsectionclip.png

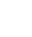

### plus.png

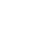

### pointclouds.png

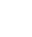

### redo.png

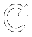

### save.png

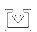

### section.png

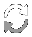

### sectionclip1.png

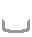

### sectionclip2.png

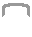

### sethome.png

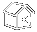

### share.png

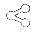

### smalldot.png

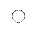

### swatch.png

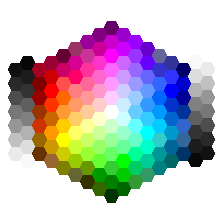

### togglegui.png

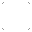

### undo.png

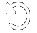
